# TMPRSS2, a SARS-CoV-2 internalization protease is downregulated in head and neck cancer patients

**DOI:** 10.1101/2020.06.16.154211

**Authors:** Andrea Sacconi, Sara Donzelli, Claudio Pulito, Stefano Ferrero, Aldo Morrone, Marta Rigoni, Fulvia Pimpinelli, Fabrizio Ensoli, Giuseppe Sanguineti, Raul Pellini, Nishant Agrawal, Evgeny Izumchenko, Gennaro Ciliberto, Aldo Giannì, Paola Muti, Sabrina Strano, Giovanni Blandino

## Abstract

**Objectives:** Two of the main target tissues of SARS-coronavirus 2 are the oral cavity pharynx-larynx epithelium, the main virus entry site, and the lung epithelium. The virus enters host cells through binding of the Spike protein to ACE2 receptor and subsequent S priming by the TMPRSS2 protease. Herein we aim to assess differences in both ACE2 and TMPRSS2 expression in normal tissues from oral cavity-pharynx-larynx and lung tissues as well as neoplastic tissues from the same histological areas. The information provided in this study may contribute to better understanding of SARS-coronavirus 2 ability to interact with different biological systems and contributes to cumulative knowledge on potential mechanisms to inhibit its diffusion.

**Materials and Methods:** The study has been conducted using The Cancer Genome Atlas (TCGA) and the Regina Elena Institute (IRE) databases and validated by experimental model in HNSCC and Lung cancer cells. Data from one COVID19 positive patient who was operated on for HNSCC was also included. We have analyzed 478 tumor samples and 44 normal samples from TCGA HNSCC cohort for whom both miRNA and mRNA sequencing was available. The dataset included 391 HPV- and 85 HPV+ cases, with 331 P53 mutated and 147 P53 wild type cases respectively. 352 out of 478 samples were male and 126 female. In IRE cohort we analyzed 66 tumor samples with matched normal sample for miRNA profiling and 23 tumor\normal matched samples for mRNA profiling. 45 out of 66 tumors from IRE cohort were male and 21 female, 38 were P53 mutated and 27 wild type. Most patients (63 of 66) in IRE cohort were HPV negative. Normalized TCGA HNSCC gene expression and miRNA expression data were obtained from Broad Institute TCGA Genome Data Analysis Center (http://gdac.broadinstitute.org/). mRNA expression data from IRE cohort used in this study has been deposited to NCBI’s Gene Expression Omnibus and is accessible through GEO series accession number GSE107591. In order to inference about potential molecular modulation of TMPRSS2, we also included miRNAs expression for the 66 IRE cohort matched tumor and normal samples from Agilent platform. DNA methylation data for TCGA tumors were obtained from Wanderer (http://maplab.imppc.org/wanderer/). We used miRWalk and miRNet web tools for miRNA-target interaction prediction and pathway enrichment analysis. The correlation and regression analyses as well as the miRNA and gene modulation and the survival analysis were conducted using Matlab R2019.

**Results:** TMPRSS2 expression in HNSCC was significantly reduced compared to the normal tissues and had a prognostic value in HNSCC patients. Reduction of TMPRSS2 expression was more evident in women than in men, in TP53 mutated versus wild TP53 tumors as well as in HPV negative patients compared to HPV positive counterparts. Functionally, we assessed the multivariate effect on TMPRSS2 in a single regression model. We observed that all variables had an independent effect on TMPRSS2 in HNSCC patients with HPV negative, TP53 mutated status and with elevated TP53-dependent Myc-target genes associated with low TMPRSS2 expression. Investigation of the molecular modulation of TMPRSS2 in both HNSCC and lung cancers revealed that expression of microRNAs targeting TMPRSS2 anti-correlated in both TCGA and IRE HNSCC datasets, while there was not evidence of TMPRSS2 promoter methylation in both tumor cohorts. Interestingly, the anti-correlation between microRNAs and TMPRSS2 expression was corroborated by testing this association in a SARS-CoV-2 positive HNSCC patient.

**Conclusions:** Collectively, these findings suggest that tumoral tissues, herein exemplified by HNSCC and lung cancers might be more resistant to SARS-CoV-2 infection due to reduced expression of TMPRSS2. The protective mechanism might occur, at least partially, through the aberrant activation of TMPRSS2 targeting microRNAs; thereby providing strong evidence on the role of non-coding RNA molecule in host viral infection. These observations may help to better assess the frailty of SARS-CoV-2 positive cancer patients.

## Introduction

Unlike other members of the *Coronaviridae* that circulate in the human population and cause only mild respiratory disease, severe acute respiratory syndrome coronavirus 2 (SARS-CoV-2) is a novel betacoronavirus which is transmitted from animals to humans and severely affects pulmonary respiration (1–5). SARS-CoV-2 enters host cells through the binding of Spike protein to ACE2 receptor and subsequent S protein priming performed by host proteases including TPMRSS2 (6–8).

HNSCC is the sixth leading cancer by incidence worldwide and the eighth most common cause of cancer death (9). Although in the past two decades new surgical and medical treatments have improved patients’ quality of life, the 5-year survival remains 40–50% of patients (10). HNSCC is typically characterized by a high incidence of local recurrences, which are the most common cause of death in HNSCC patients, occurring in 60% of the cases (11). The current standard therapies are surgical and systemic treatment followed by adjuvant radiotherapy (RT) with or without chemotherapy. Unfortunately, advances in treatments for HNSCC over the past two decades failed to substantially improve the overall disease outcome, with radio and chemo-resistance (intrinsic or acquired) remains one of the major challenges in the current therapy of HNSCC.

Here we sought to investigate the expression of both ACE2 and TMPRSS2 in head and neck cancer specimens. We found that unlike ACE2, whose expression was unchanged in HNSCC patients, TPMRSS2 expression was significantly reduced in tumor tissues compared to non-tumorous ones in both HNSCC TCGA and IRE datasets. Notably, TMPRSS2 downregulation associated with poorer survival in HNSCC patients with TP53 mutations, HPV negative status, aberrant MYC activation and low immune signature. Mechanistically, downregulation of TMPRSS2 significantly correlated with aberrant upregulation of specific microRNAs, which might target TMPRSS2 post-transcriptionally, thereby leading to its reduced expression levels in tumor tissues. microRNAs are an abundant class of small noncoding RNAs of approximately 22 nucleotides long. They act as negative regulators of gene expression at the post-transcriptional level, by binding their target mRNAs through imperfect base pairing with the respective 3’-untranslated region (3’-UTR). Deregulation of microRNAs leads to an altered expression of genes involved in many cell functions and cell fate regulation. Therefore, by regulating genes and pathways, microRNAs could contribute to modulate biological functions including potential effect on epithelial cell interaction with viruses and tumorigenesis (12, 13). We found that the expression of a group of microRNAs (miR-193b-3p; miR-503-5p; miR-455-5p; miR-31-3p; miR-193b-5p; miR-2355-5p) was anti-correlated to that of their target TMPRSS2. This anti-correlated expression was also evidenced in an HNSCC patient positive for SARS-CoV-2 infection.

## Results

### ACE2 and TMPRSS2 expression in HNSCC patients

To test our hypothesis, we first assessed the expression of both ACE2 and TMPRSS2 in TCGA HNSCC dataset. We found that while ACE2 expression level was comparable between non-tumorous versus malignant tissues, the TMPRSS2 expression was significantly downregulated in HNSCC patient samples (Fig. 1A-B). Interestingly, while female patients showed a more pronounced downregulation of TMPRSS2 than males (Fig. 1C), the level of ACE2 expression in female patients was upregulated (Suppl. Fig.1A).

**Figure 1.**
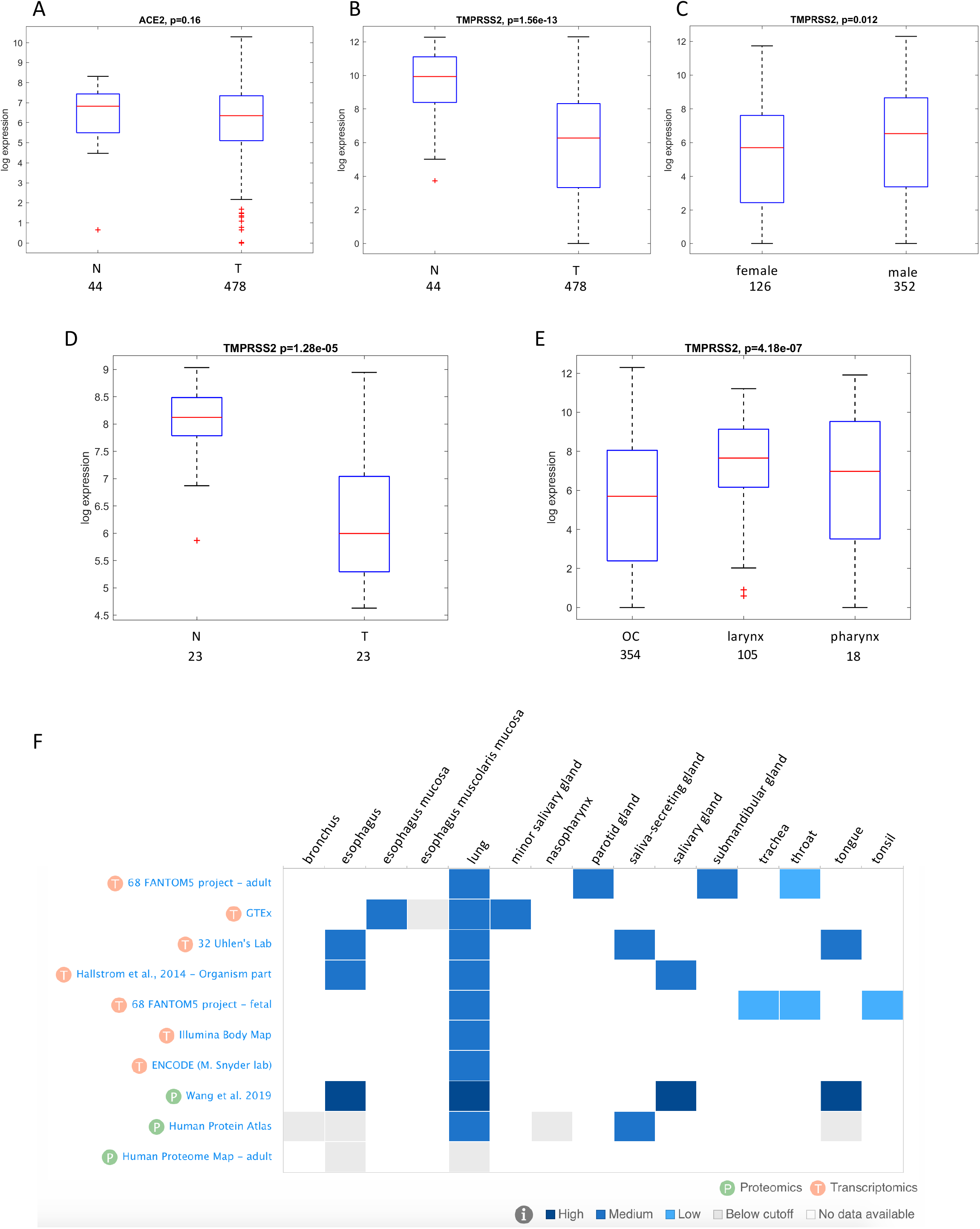
Distribution of ACE2 and TMPRSS2 gene expression in HNSCC patients. **A-B** Box-plot analysis representing ACE2 (A) and TMPRSS2 (B) gene expression levels in non-tumorous (N) and tumor (T) tissues from the HNSCC TCGA dataset. **C** Box-plot analysis representing TMPRSS2 gene expression levels in tumoral HNSCC TCGA samples according to the gender (female or male). **D** Box-plot analysis representing TMPRSS2 gene expression levels in non-tumorous (N) and tumor (T) tissues from the HNSCC cohort of IRCSS Regina Elena National Cancer Institute of Rome. **E** Box-plot analysis representing TMPRSS2 gene expression levels in tumoral HNSCC TCGA samples according to anatomical site. **F** Expression analysis of TMPRSS2 in the selected normal tissues and in the indicated studies, by using EMBL-EBI Expression Atlas public repository. The analysis includes both transcriptomics (T) and proteomics (P) data.

To validate these observations, we have assessed the expression of TMPRSS2 in an additional cohort of HNSCC patients enrolled at Regina Elena Cancer Institute (14). This cohort includes naïve HNSCC patients for which tumor and peritumoral tissues and resection margin specimens are available (15). Confirming our findings, TMPRSS2 expression was significantly reduced in tumor samples compared to non-tumorous tissues (Fig. 1D). Interestingly, expression of TMPRSS2 was significantly higher in tumors from larynx and pharynx compared to malignancies of the oral cavity (Fig. 1E). In non-tumorous tissues from either TCGA or IRE datasets no correlation between the TPMRSS2 expression and sex or histological site (oral cavity, larynx and pharynx) was detected (Suppl. Fig.1B).

To further assess the pattern of TMPRSS2 expression in normal tissues, we have used ATLAS, and showed data of transcript and protein expression of different tissue sites from which HNSCC develops (Fig. 1F). Lung tissue was included as a reference, as high expression of TMPRSS2 was reported in lung by RNA-Seq and proteomic analyses conducted by several studies (Fig. 1F). A widespread expression of TMPRSS2 was evidenced in lung and head neck tissues (Fig. 1F).

### TMPRSS2 expression is prognostic and associates with TP53 mutations and HPV status in HNSCC patients

As for many human cancers, TP53 is the most frequently mutated gene in HNSCC (9). In TCGA dataset, which includes 478 HNSCC molecularly well characterized cases, we found that patients carrying TP53 mutation exhibited a significantly lower level of TMPRSS2 expression compared to the patients with intact TP53 gene (Fig. 2A).

**Figure 2.**
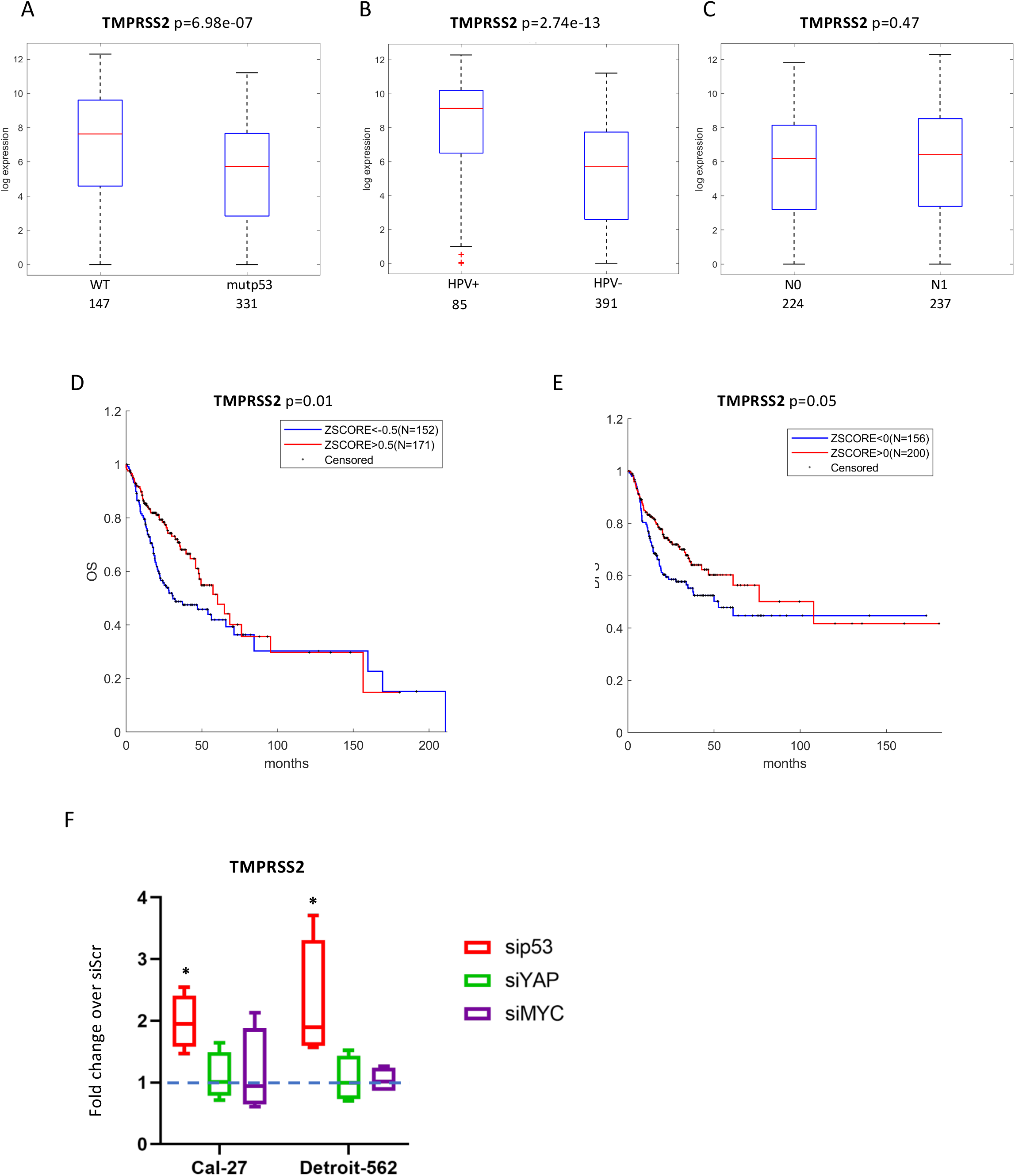
TMPRSS2 gene expression, HNSCC patient clinical variables and correlation with survival. **A-C** Box-plot analysis representing TMPRSS2 gene expression levels in tumoral HNSCC TCGA samples according to TP53 status (A), or HPV status (B), or N status (C). **D-E** Kaplan–Meier survival curves for TCGA HNSCC patients showing overall survival (OS) (D) and disease-free survival (DFS) (E), according to TMPRSS2 gene expression. **F** qRT-PCR analysis of TMPRSS2 expression levels in Cal-27 and Detroit-562 cell lines upon depletion of mutantp53 (sip53) or YAP (siYAP) or MYC (siMYC) compared to cells transduced with scramble molecules (value=1). *p-value < 0.05

Since the vast majority of TP53 mutated patients are HPV negative, we next looked at HPV negative cohort independently, and found that TMPRSS2 expression was substantially lower in HPV negative patients than in HPV positive ones (Fig. 2B). TMPRSS2 expression did not vary accordingly to N status (Fig. 2C). As it was previously reported that patients with HNSCC HPV negative and TP53 mutated cancer exhibit shorter overall survival (OS) and disease free survival (DFS) (16), we next performed a Kaplan Meyer analysis on data obtained from TCGA database, which revealed that lower TMPRSS2 expression associated with shorter OS and DFS in HNSCC patients (Fig. 2D-E).

We have next analysed the role of TMPRSS2 using two HNSCC cell lines (Cal-27 and Detroit-562) carrying TP53 mutations that exert gain of function activities. When p53 protein in these cell lines was depleted, expression of TMPRSS2 transcript was significantly up-regulated (Fig. 2F), suggesting that mutant p53 oncogenic protein may regulate (either directly or indirectly) TMPRSS2 expression in HNSCC cell lines. We also analysed Cal-27, and Detroit-562 cell lines depleted for YAP and MYC, two important co-factors of transcriptional activity of gain of function mutant p53 proteins (17). Unlike mutant p53, neither YAP nor MYC depletion did not affect the TMPRSS2 level in these cell lines (Fig. 2F). ACE2 expression was unaffected by mutant p53, YAP and MYC depletion in both HNSCC cell lines (Suppl. Fig 2). In summary, these observations indicate that low expression of TMPRSS2, in a context of TP53 mutations and HPV negative status is associated with poor prognosis in HNSCC patients.

### TMPRSS2 expression is associated with aberrant MYC activity and mutant p53 in HNSCC patients

MYC is a proto-oncogene that plays a crucial role in different steps of tumorigenesis (18). In HNSCC, aberrant MYC expression is associated with poor survival (19). In our study, at univariate levels we found that low TMPRSS2 expression was significantly associated with high levels of MYC in TCGA HNSCC patients (Fig. 3A).

**Figure 3.**
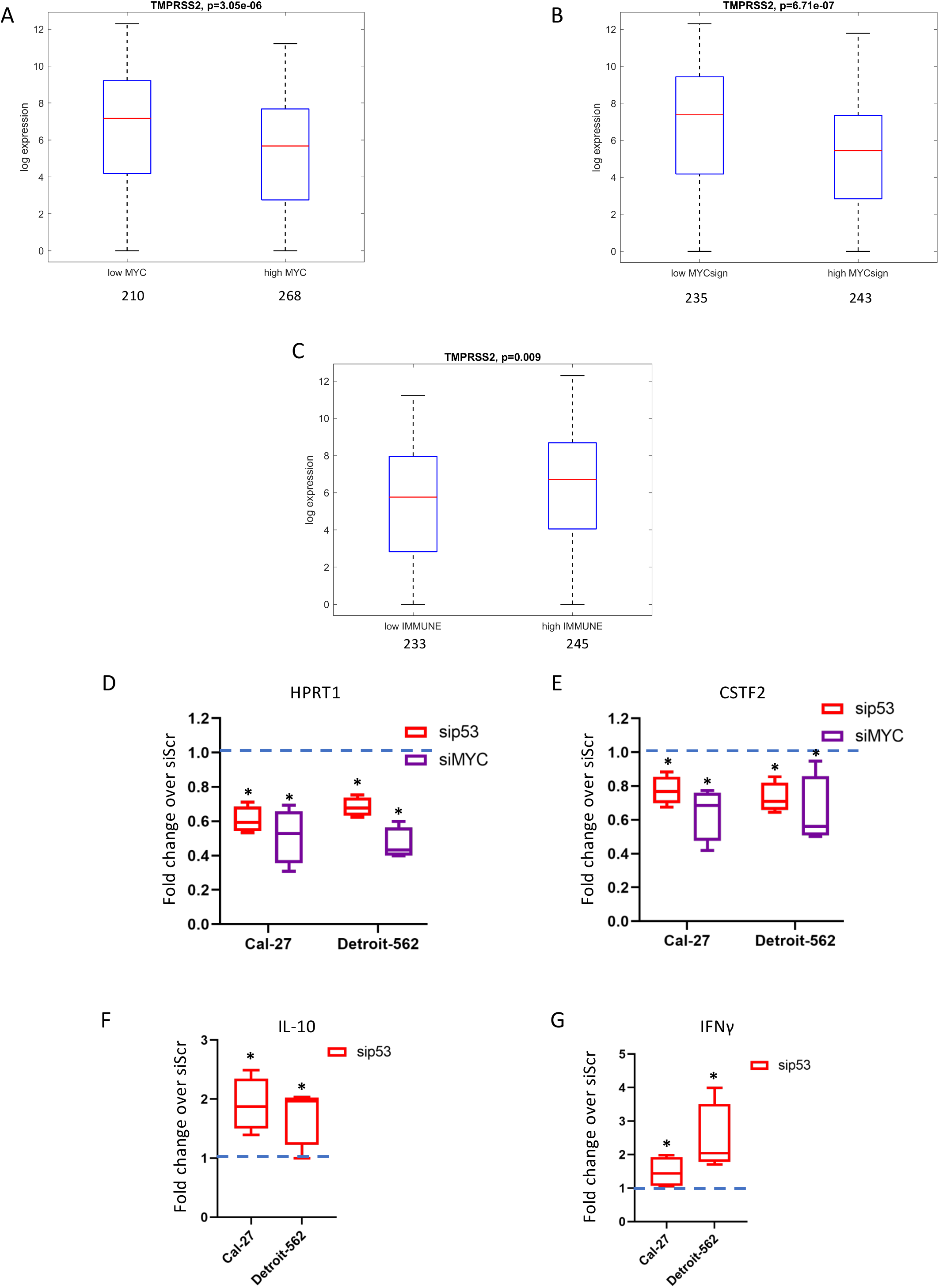
TMPRSS2 gene association with MYC signature and immune signature in HNSCC patients. **A** Box-plot analysis representing TMPRSS2 gene expression levels in tumoral HNSCC TCGA samples according to low or high expression values of MYC gene. **B** Box-plot analysis representing TMPRSS2 gene expression levels in tumoral HNSCC TCGA samples according to low or high expression values of 22-gene MYC signature. **C** Boxplot analysis representing TMPRSS2 gene expression levels in tumoral HNSCC TCGA samples according to low or high expression values of 17-gene immune signature. **D-E** qRT-PCR analysis of HPRT1 (D) and CSTF2 (E) expression levels in Cal-27 and Detroit-562 cell lines upon depletion of mutant p53 (sip53) or MYC (siMYC) compared to scramble control cells (value=1). **F-G** qRT-PCR analysis of IL-10 (F) and IFNγ (G) expression levels in Cal-27 and Detroit-562 cell lines upon depletion of mutant p53 (sip53) or MYC (siMYC) compared to scramble control cells (value=1). *p-value < 0.05

We have recently reported that MYC is a pivotal mediator of gain of function mutant p53 signalling in HNSCC (20). We have identified a mutantp53/MYC dependent signature whose aberrant activation (high expression levels) associated with shorter overall survival in HNSCC patients (Suppl. Fig. 3A) (20).

Notably, low expression of TMPRSS2 negatively associated with the high level of this previously reported 22-gene mutant p53/MYC signature (Fig. 3B). It has been extensively reported that mutant p53 proteins actively promote resistance to therapies, for example by inhibiting the onset of apoptosis (21). However, failure of patients to respond to anti-cancer therapies is also dependent on blockade of effector immune cell function by tumor cells (immune evasion) (22). It was reported that mutant p53 is associated with the inability of the immune system to recognize neoantigens, which should be abundant in these tumors characterized by high mutational burden (23). To explore whether mutant p53 is the driver of immune suppression in HNSCC, we evaluated the association between TP53 status and immune-related signatures using the TCGA dataset of HNSCC. In this study we have used an immune signature reported by Wood et al., due to its high stability in HNSCC specimens (Suppl. Fig.3B) (24). We found that low expression of TMPRSS2 significantly associates with low immune signature in TCGA HNSCC patients’ cohort (Fig. 3C). The transcriptional crosstalk between mutant p53, MYC/MYC signature and immune signature was assessed in two HNSCC cell lines, Cal-27 and Detroit-562 cells, As expected, both mutant p53 and MYC depletion reduced the expression of two MYC target genes, HPRT1 and CSTF2 (Fig. 3D-E). Interestingly, depletion of mutant p53 protein increased the expression of both IL-10 and IFNγ transcripts in Cal-27 and Detroit-562 cell lines (Fig. 3F-G).

We next conducted a linear regression analysis which includes the immune signature expression, HPV and TP53 status as well as MYC signature, to assess whether addition of the immune signature could improve the total variance seen in the model which did not include the immune signature.

We subsequently assessed the effect of MYC, MYC signature, HPV status (positive and negative), the wild-type and mutated TP53 and immune-signature on TMPRSS2 expression at univariate and multivariate levels. The univariate analysis showed that all the variables were able to significantly and independently modulate TMPRSS2 expression (Table 1, upper panel). When the same variables were included in one logic regression model, beside the immune signature, all other variables have significantly and independently contributed to the TMPRSS2 expression with a 15% of its total variance explained (Table 1, lower panel). In general, these findings reveal the association between TMPRSS2 downregulation with mutant p53 and MYC oncogenic activities in HNSCC patients, and indicate that the immune signature did not add any substantial effect when all other variables contribution was taken into consideration. The lack of independent effect of immune signature on TMPRSS2 expression could be related to the biological interdependence of the immune signature with TP53, which has a strong effect on inducing immune suppression (25).

**Table 1.**
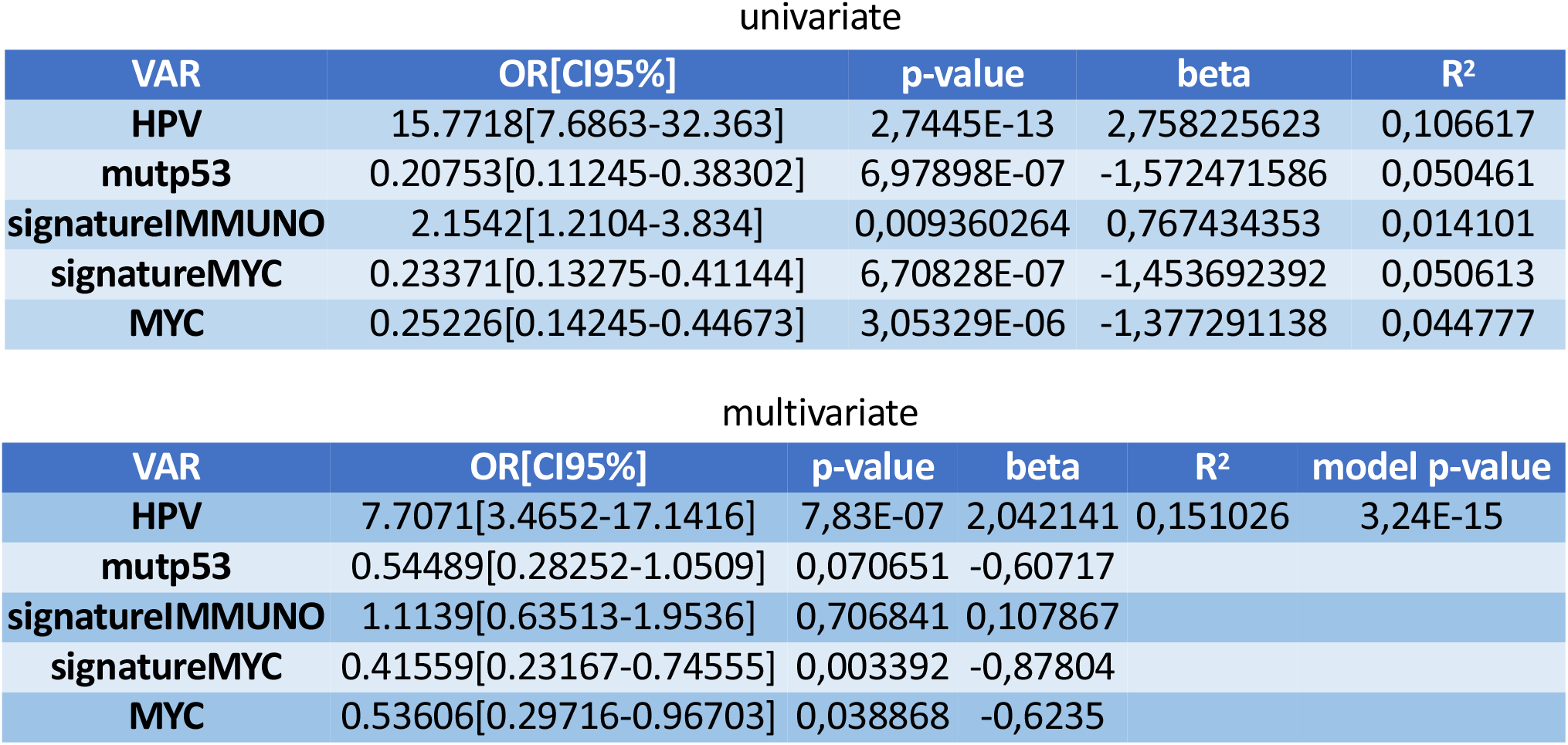
TMPRSS2 gene association with immune signature, MYC and MYC signature, HPV status and P53 mutation in HNSCC patients. Linear univariate and multivariate regression models were built from TCGA HNSCC patients considering TMPRSS2 as outcome variable. In the table is indicated the percentage of variance explained (R^2^) and the total p-value of the multivariate model.

### Epigenetic control of TMPRSS2 expression in HNSCC

To further investigate the molecular mechanisms underlying TMPRSS2 downregulation in HNSCC we have analyzed the extent of TMPRSS2 promoter methylation. To this end, we used a Wanderer tool that includes data of methylation specific sequencing for cases in TCGA databases (26). We compared the TMPRSS2 promoter methylation in tumor versus non-tumorous tissues in HNSCC TCGA dataset. As shown in Fig. 4A, the intensity ratio of methylated and unmethylated alleles (β value) was lower than 0.5, indicating that CpG islands either within or in the vicinity of TPMRSS2 promoter were unmethylated both in tumor and normal tissues. A similar unmethylated pattern for TMPRSS2 promoter was evidenced in both lung adenocarcinoma (LUAD) and lung squamous cell carcinoma (LUAS) specimens, where expression levels of TMPRSS2 were significantly downregulated (Fig. 4B-E).

**Figure 4.**
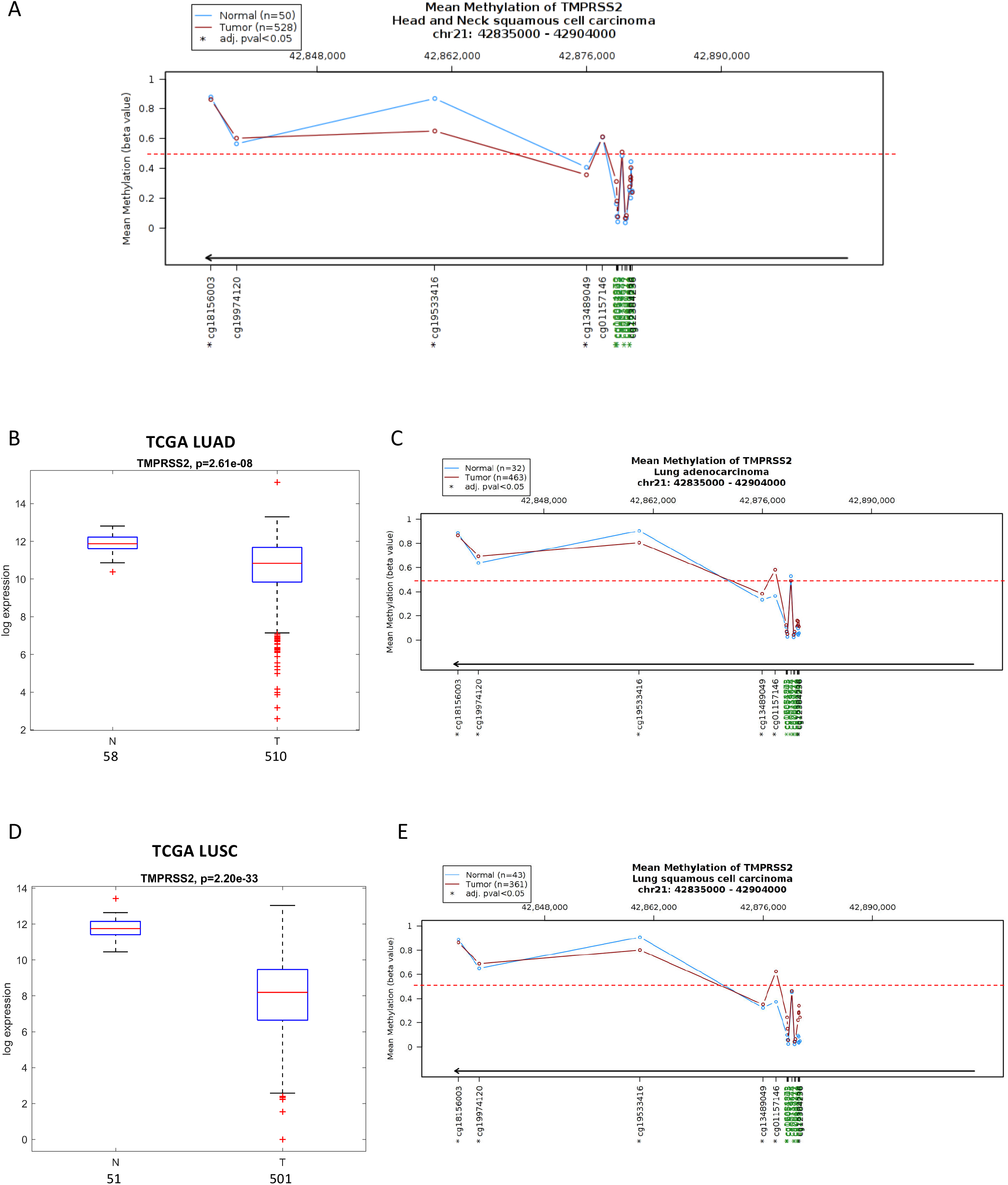
TMPRSS2 gene methylation status in normal and tumoral samples of HNSCC and lung cancer patients. **A** DNA methylation profile of TMPRSS2 gene in normal (blue line) and tumoral (red line) samples of HNSCC TCGA patients. **B** Box-plot analysis representing TMPRSS2 gene expression levels in non-tumorous (N) and tumor (T) tissues from the lung adenocarcinoma (LUAD) TCGA dataset. **C** DNA methylation profile of TMPRSS2 gene in normal (blue line) and tumoral (red line) samples of LUAD TCGA patients. **D** Box-plot analysis representing TMPRSS2 gene expression levels in non-tumorous (N) and tumor (T) tissues from the lung squamous cell carcinoma (LUSC) TCGA dataset. **E** DNA methylation profile of TMPRSS2 gene in normal (blue line) and tumoral (red line) samples of LUAD TCGA patients. DNA methylation profile has been performed by using Wanderer software. Beta values (β) are the estimate of methylation level using the ratio of intensities between methylated and unmethylated alleles. β values are between 0 and 1, with 0 being unmethylated and 1 fully methylated. The tool provides the detailed individual beta values of all the HumanMethylation450 probes inside or in the vicinity of the gene. Probes for the CpG island are indicated in green. Dotted red line indicates the threshold β value of 0.5

We next investigated whether TMPRSS2 downregulation in HNSCC patients could be due to selective targeting by microRNAs. Using miRWalk tool we searched for microRNAs that could putatively target the TMPRSS2 (Suppl. Table 1). We have selected a number of microRNAs candidates (miR-193b-3p; miR-503-5p; miR-455-5p; miR-31-3p; miR-193b-5p; miR-2355-5p) whose expression levels were inversely correlated with Pearson R=0.50402; 0.4383; R=0.42242; R=0.46154; R=0.42246; R=0.46644 compared to that of TMPRSS2 expression in TCGA HNSCC tumoral tissues (Fig. 5A). Notably, we found that all six selected microRNAs were significantly upregulated in tumoral tissues compared to the non-tumorous samples (Fig. 5B). This upregulation was confirmed in IRE cohort for all but miR-193-5p whose expression was unchanged and for miR-2355-5p that was not present on the arrays used to profile HNSCC matched tumor and non-tumorous samples (Fig. 5C).

**Figure 5.**
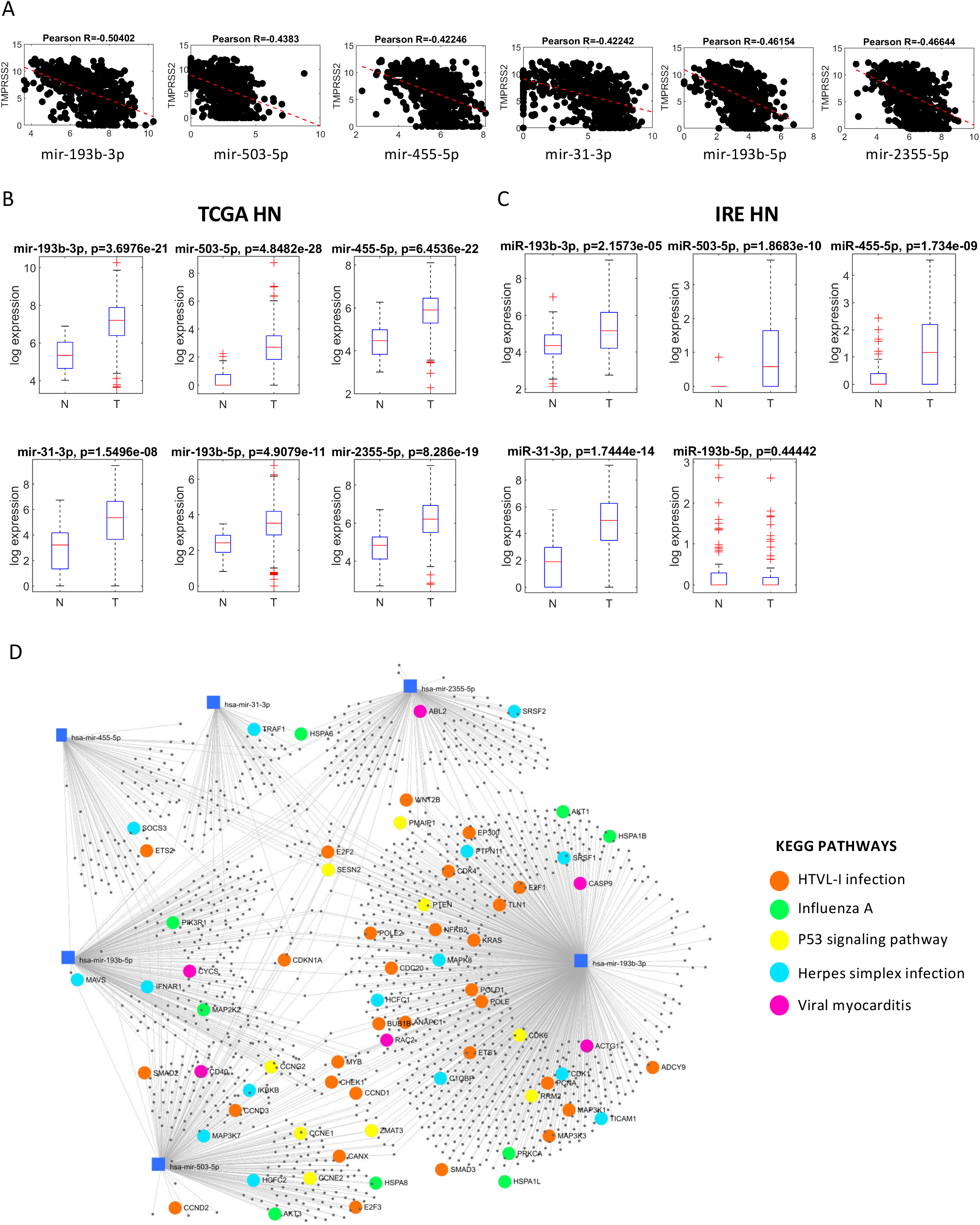
Expression levels of miRNAs predicted to target TMPRSS2 gene in HNSCC patients. **A** Graphs showing the correlation (Spearman coefficients) between indicated miRNAs and TMPRSS2 in the TCGA HNSCC dataset. **B** Box-plot analysis representing expression levels of miRNAs predicted to target TMPRSS2 in non-tumorous (N) and tumor (T) tissues from TCGA HNSCC patients. **C** Box-plot analysis representing expression levels of miRNAs predicted to target TMPRSS2 in non-tumorous (N), peri-tumor (PT) and tumor (T) tissues from IRCSS Regina Elena National Cancer Institute (IRE) HNSCC patients. **D** miRNA-centric network using the web tool miRNet. The network shows main validated miRNA-target interaction of the six selected miRNA signature. Genes involved in specific KEGG pathways are highlighted.

miRNet online based tool was used to identify the potential targets of miR-193b-3p, miR-503-5p, miR-455-5p, miR-31-3p, miR-193b-5p and miR-2355-5p (Suppl. Table 2). Subsequently, the identified list of microRNA targets was assessed using KEGG pathway enrichment analysis to reveal cell signaling pathways impacted by the aberrant activities of the selected panel of microRNAs (Table 2). In figure 5D we reported those pathways whose microRNA validated targets impinged on viral infections and p53 pathway.

**Table 2.** KEGG enriched pathways evaluated from validated miRNA-target interactions (miRNet). Total number of genes included in each pathway and number of genes represented in each pathway (hits) with respective p-values are shown.

| **Pathway** | **Total** | **Hits** | **Pvalue** |
| --- | --- | --- | --- |
| Cell cycle | 124 | 42 | 3.08e-15 |
| Small cell lung cancer | 80 | 22 | 9.85e-07 |
| Prostate cancer | 87 | 23 | 1.19e-06 |
| Pathways in cancer | 310 | 52 | 3.53e-06 |
| Focal adhesion | 200 | 38 | 4.18e-06 |
| p53 signaling pathway | 68 | 18 | 1.73e-05 |
| Pancreatic cancer | 69 | 18 | 2.15e-05 |
| HTLV-I infection | 199 | 36 | 2.37e-05 |
| Chronic myeloid leukemia | 73 | 18 | 4.9e-05 |
| Colorectal cancer | 49 | 14 | 6.03e-05 |
| Melanoma | 68 | 17 | 6.57e-05 |
| Adherens junction | 70 | 17 | 9.74e-05 |
| Non-small cell lung cancer | 52 | 14 | 0.000123 |
| Neurotrophin signaling pathway | 123 | 23 | 0.000443 |
| Glioma | 65 | 15 | 0.000453 |
| One carbon pool by folate | 19 | 7 | 0.00084 |
| Toxoplasmosis | 93 | 18 | 0.00121 |
| Fanconi anemia pathway | 39 | 10 | 0.00172 |
| Renal cell carcinoma | 60 | 13 | 0.00199 |
| Selenocompound metabolism | 12 | 5 | 0.00259 |
| Oocyte meiosis | 108 | 19 | 0.00287 |
| Bladder cancer | 29 | 8 | 0.00302 |
| Acute myeloid leukemia | 57 | 12 | 0.00378 |
| Progesterone-mediated oocyte maturation | 80 | 15 | 0.00417 |
| Glyoxylate and dicarboxylate metabolism | 19 | 6 | 0.00485 |
| Regulation of actin cytoskeleton | 182 | 27 | 0.00539 |
| Pyrimidine metabolism | 101 | 17 | 0.00741 |
| ErbB signaling pathway | 87 | 15 | 0.00927 |
| Thyroid cancer | 28 | 7 | 0.00975 |
| MAPK signaling pathway | 265 | 35 | 0.0108 |
| Influenza A | 107 | 17 | 0.0131 |
| Endometrial cancer | 44 | 9 | 0.014 |
| RNA transport | 126 | 19 | 0.0154 |
| Herpes simplex infection | 103 | 16 | 0.0193 |
| Protein processing in endoplasmic reticulum | 129 | 19 | 0.0194 |
| Glycine, serine and threonine metabolism | 33 | 7 | 0.0239 |
| Viral myocarditis | 26 | 6 | 0.0241 |
| Cysteine and methionine metabolism | 34 | 7 | 0.0279 |
| Jak-STAT signaling pathway | 99 | 15 | 0.0283 |
| mRNA surveillance pathway | 82 | 13 | 0.0285 |
| Epstein-Barr virus infection | 91 | 14 | 0.0297 |
| Valine, leucine and isoleucine degradation | 44 | 8 | 0.0384 |
| Pertussis | 52 | 9 | 0.0385 |
| mTOR signaling pathway | 45 | 8 | 0.0432 |
| Tight junction | 118 | 16 | 0.0585 |

To further confirm the negative correlation between the level of TMPRSS2 transcript and microRNAs we assessed the expression of miR-503-5p and TMPRSS2 in an HNSCC patient positive to COVID-19. 80 years old male patient underwent surgery in February 2020 for resection of squamous cell carcinoma (T4aN1G0R1). Two days after surgery, the patient developed pneumonia symptoms and was found positive for SARS-CoV-2 infection by nasopharyngeal swab (Fig. 6A-C). Excised FPPE tumoral tissue was assessed for viral gene expression and found negative to SARS-CoV-2 infection (Fig. 6D, upper panel). Matched tumoral and non-tumorous FPPE specimens from a male lung cancer patient who underwent surgical resection in 2011 were used as a reference negative control and found negative for expression of SARS-CoV-2 target gene expression (Fig. 6D medium and lower panels). Expression levels of TMPRSS2 were assessed by RT-PCR in COVID 19 HNSCC patient as well as in five lung cancer patient samples resected from 2011 to 2014 respectively (Fig. 6E). Matched non-tumorous specimens were also analyzed. Interestingly, we found that TMPRSS2 expression was significantly lower in both HNSCC-COVID19 patients and lung cancer tissues compared to matched non-tumorous counterparts. Unlike TMPRSS2 expression, miR-503-5p levels were significantly higher in tumors than in non-tumorous tissues (Fig. 6F).

**Figure 6.**
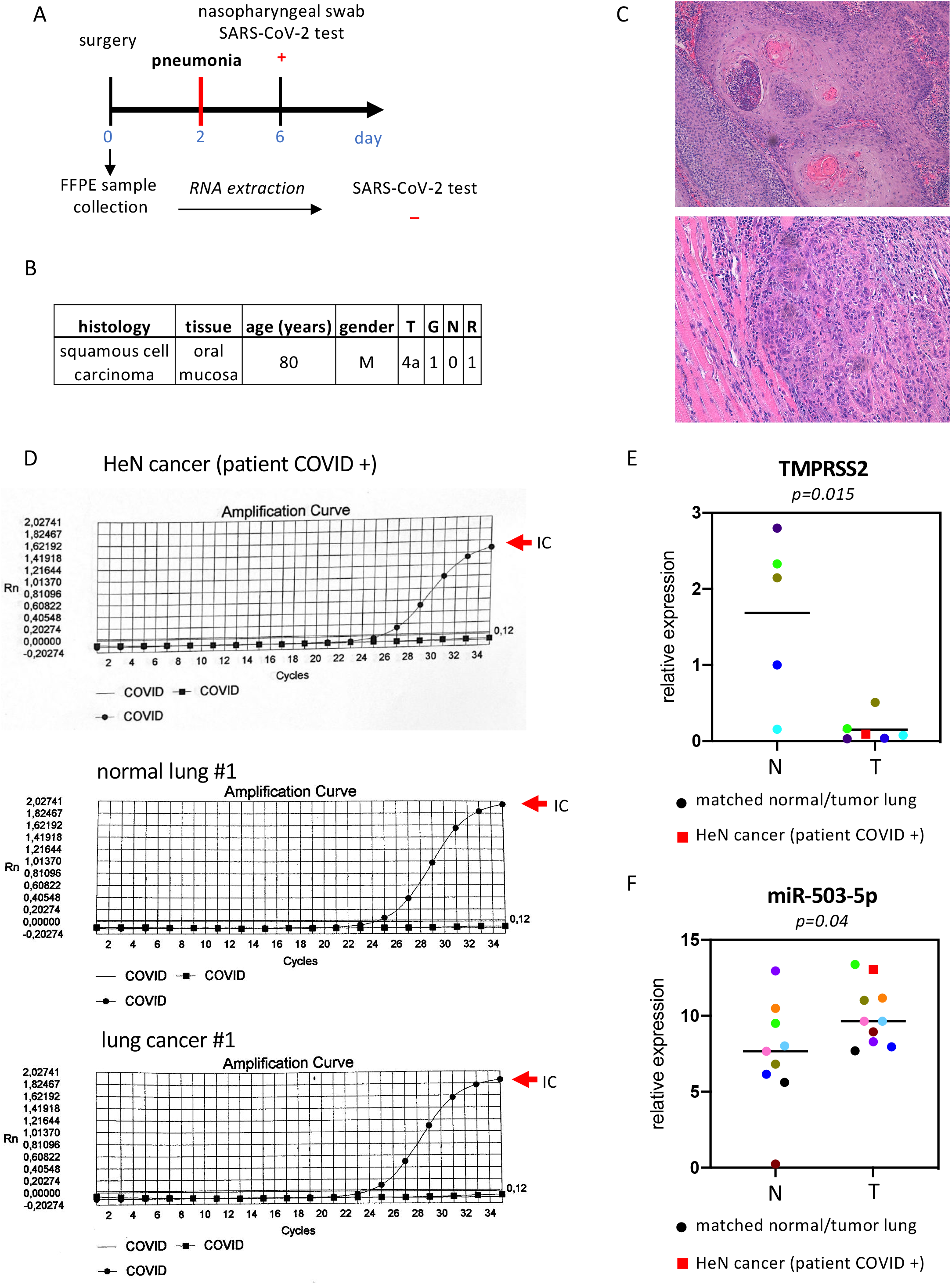
Case report: COVID-19 positive HNSCC patient. **A** Clinical history of COVID-19 positive HNSCC patient: at day 0 patient underwent surgical resection of the tumor that was formalin fixed and paraffin embedded. 2 days after surgery the patient developed pneumonia symptoms and nasopharyngeal swab SARS-CoV-2 test at day 6 was positive. SARS-CoV-2 test was also performed in FFPE tumoral tissue and resulted to be negative. **B** Clinical characteristics of COVID-19 positive HNSCC patient. **C** Well differentiated squamous cell carcinoma (G1) showing areas of keratinization with infiltration of muscular layer. **D** Real-time PCR curves from SARS-CoV-2 test run of RNA extracted from FFPE tumor tissue of HNSCC patient and from FFPE non-tumorous and tumor lung tissues of a lung cancer patient enrolled in 2011. The internal control was indicated by the red arrow. **E** qRT-PCR analysis of TMPRSS2 in 5 normal lung tissues (N) and 5 matched lung cancer tissues plus COVID-19 positive HNSCC tissue (T). **F** qRT-PCR analysis of miR-503-5p in 9 non-tumorous lung tissues (N) and 9 matched lung cancer tissues plus COVID-19 positive HNSCC tissue (T).

Both miR-31-3p (located on Chr 9) and miR-503-5p (located on Chr X) are hosted in long non-coding RNAs, named MIR31HG and MIR503HG, respectively (Fig. 7A). We found that the expression of both MIR31HG and MIR503HG is higher in HNSCC tumors than in non-tumorous tissues (Fig. 7B-C) similarly to miR-31-3p and miR-503-5p. Interestingly, HNSCC patients with high expression of these two microRNAs exhibit shorter disease free survival (Fig.7D-E).

**Figure 7.**
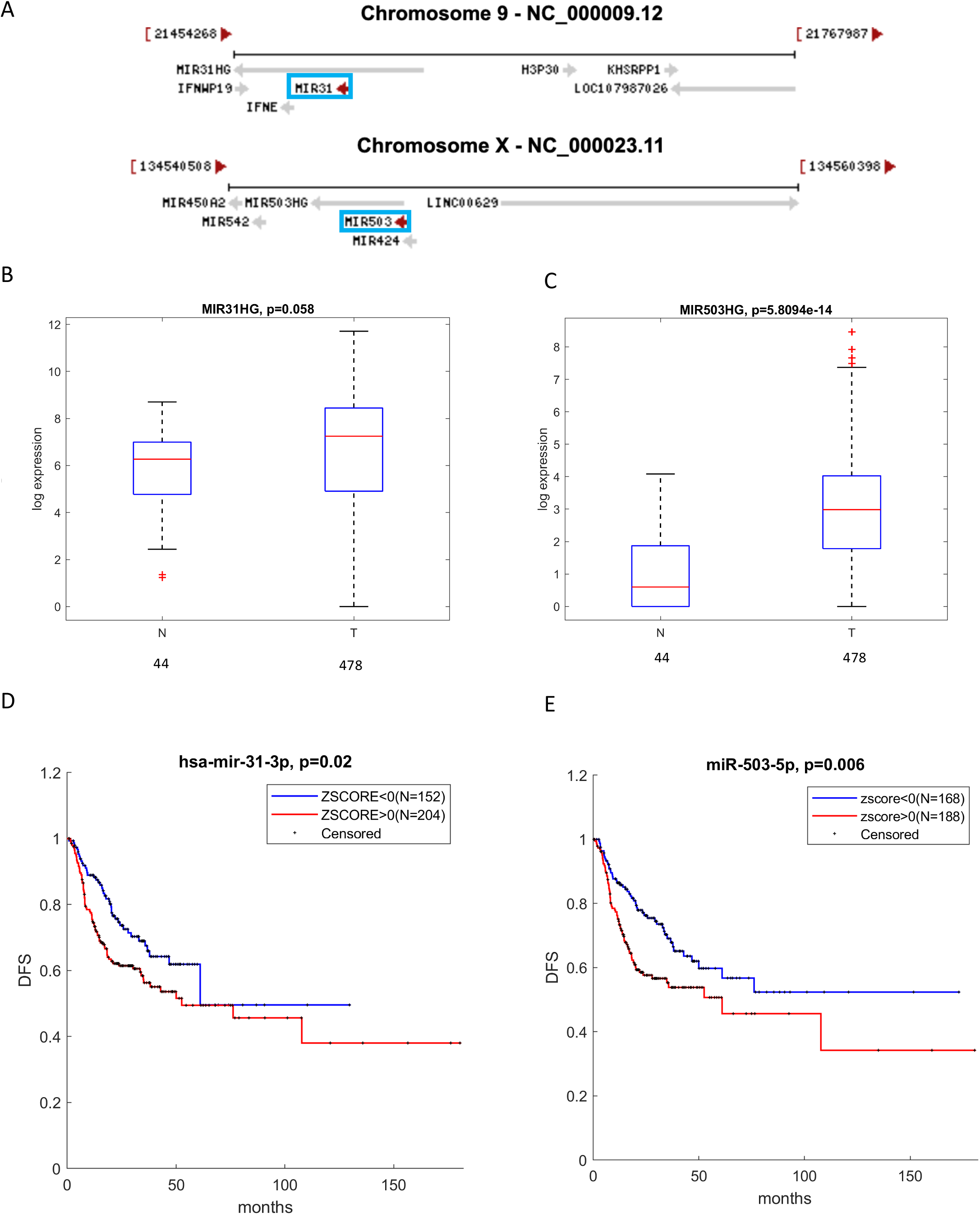
miR-31 and miR-503 locus in HNSCC patients. **A** Schematic representation of miR-31 and miR-503 genes localization. **B-C** Box-plot analysis representing expression levels of MIR31HG (B) and MIR503HG (C) in non-tumorous (N) and tumor (T) tissues from TCGA HNSCC patients. **E-F** Kaplan–Meier survival curves for TCGA HNSCC patients showing disease-free survival (DFS) according to miR-31-3p (D) or miR-503-5p (E) gene expression.

Collectively these findings suggest that downregulation of TMPRSS2 expression evidenced in HNSCC and LUAD/LUSC is not due to methylation of its regulatory regions. Our study further suggests that aberrant expression of microRNAs targeting TMPRSS2, may assemble a post-transcriptional regulatory network leading to a reduced expression of TMPRSS2 in HNSCC. This regulatory network might also include other non-coding RNA molecules such as LNC-RNAs.

## Discussion

The present work indicates that neoplastic tissue from SARS-CoV-2 target organs such as head and neck and lung, might be more resistant to SARS-CoV-2 infection due to reduced expression of TMPRSS2 (Fig. 8). The study was based on the bioinformatic and biostatistics analysis as well as on the subsequent validation of the results in cell models and, as a-proof-of-principle, in neoplastic tissue directly collected from a HNSCC patient affected by Covid19. Notably, we found that reduced expression of TMPRSS2 associates with HPV negative status and TP53 mutations, both of which are important determinants of poor survival in HNSCC patients. We have recently shown that MYC as oncogenic protein di per se and a MYC-dependent gene signature cooperate with gain of function TP53 mutations to foster HNSCC proliferation and to increase resistance to the treatment (20). Indeed, we found that depletion of mutant p53 proteins in HNSCC cell lines increased TMPRSS2 expression, suggesting that mutant p53 contributes either directly or indirectly to reduced TMPRSS2 expression. Unlike depletion of mutant p53, silencing of YAP and MYC, two oncogenic co-factors of gain of function mutant p53 proteins did not have any effect of TMPRSS2 expression (20, 27, 28). While these findings may entirely rule out a possibility that gain of function mutant p53-dependet transcriptional networks controls TMPRSS2 expression in HNSCC cells, our observations highlight that tumor cells carrying TP53 mutations may activate post-transcriptional events impinging the TMPRSS2 expression. Supporting such possibility, we have detected that the up-regulation of TMPRSS2 targeting microRNAs inversely correlated with expression of TMPRSS2 in HNSCC. We and others have previously reported that gain of function mutant p53 proteins are able to either up-regulate or down-regulate the expression of microRNAs (29, 30). The lack of methylation in regulatory regions of TMPRSS2 further supports the proposed working model (Fig. 8). Our findings derived from TCGA indicate that similar molecular mechanisms might underlie the reduced expression of TMPRSS2 in both LUAD and LUSC, as no evidence of promoter methylation was also evidenced for lung cancer patients.

**Figure 8.**
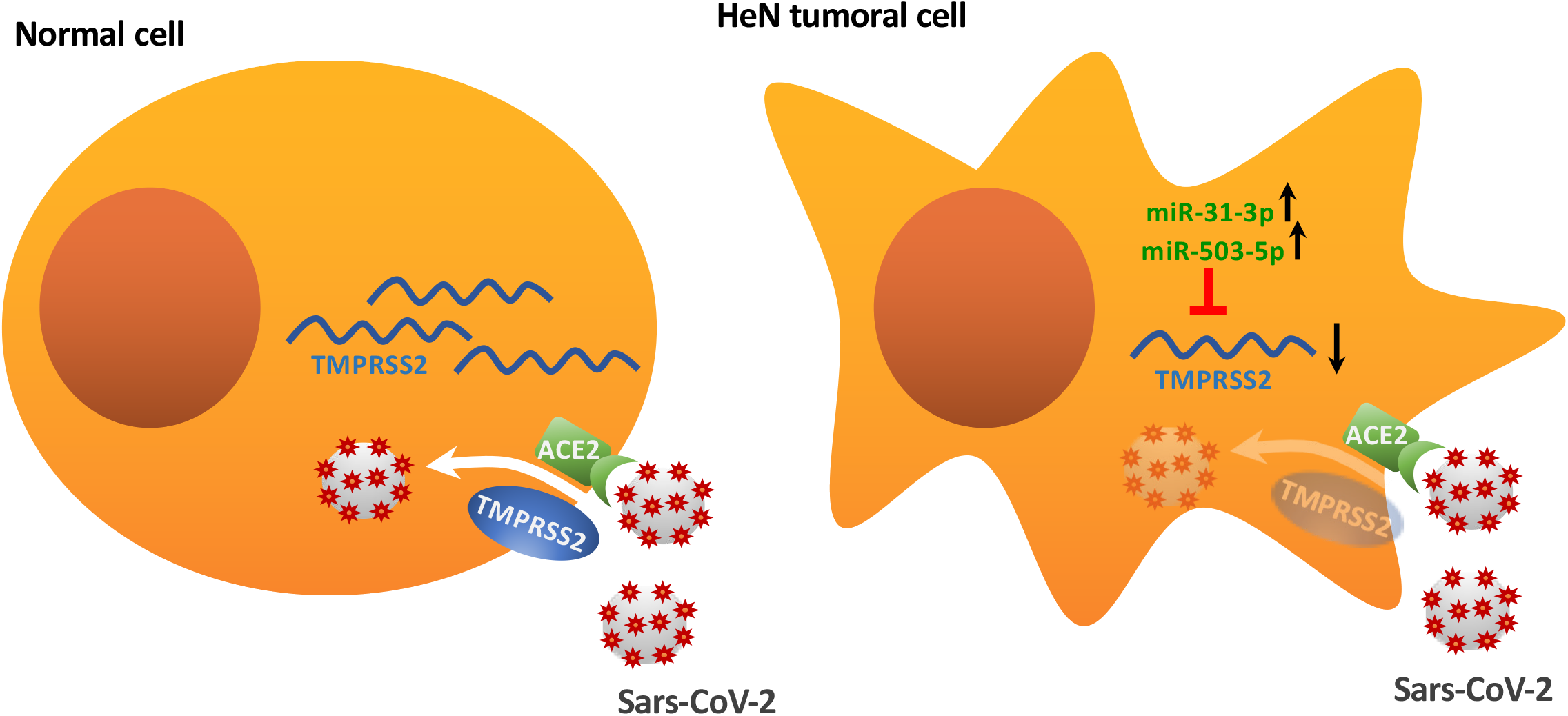
Schematic representation of the proposed molecular mechanism. In normal cell, TMPRSS2 mRNA translation leads to the production of the transmembrane-bound serine protease that allows the internalization of SARS-CoV-2 bound to the ACE2 receptor. On the contrary, in tumoral cell, in particular head and neck tumoral cell, TMPRSS2 mRNA levels are downregulated by the targeting activity of upregulated miRNAs, such as miR-31-3p and miR-503-5p. This might determine a reduction in the expression of the protease and a consequent inhibition of the internalization of SARS-CoV-2-ACE2 complex.

These findings further support our hypothesis that TMPRSS2 downregulation in HNSCC patients was associated with selective targeting of microRNAs, in particular those which putatively target TMPRSS2. Expression of six selected microRNAs (miR-193b-3p; miR-503-5p; miR-455-5p; miR-31-3p; miR-193b-5p; miR-2355-5p) significantly anti-correlated with expression of TMPRSS2 (Fig. 5B). It is particularly noteworthy to mention that part of the most significantly predicted pathways targeted by the six microRNAs, were related to respiratory virus infections (e.g., Influenza A) and other viral infections (e.g., Herpes Simplex). The observed microRNA-induced modulation of TMPRSS2 provides a new insight for potential assessment of agents capable of regulating the microRNA expression and induce TMPRSS2 downregulation, as SARS-CoV-2 infection prevention strategy.

Clinically, it is increasingly evident that cancer patients represent at least in part, the most vulnerable population target of SARS-CoV-2 infection. This is certainly due to many factors, including the aggressiveness of the type of tumor and the side effects of cancer treatment. Here we provide additional evidence suggesting that tumor tissues are less prone to SARS-CoV-2 infection than non-tumorous tissues, due to reduced expression of TMPRSS2. Furthermore, this reduction is more evident in HNSCC patients with shorter overall survival as well as those with HPV negative status and TP53 mutations.

## Materials and Methods

The study has been conducted using the TCGA and the Regina Elena Institute (IRE) databases and validated by experimental model in HNSCC and Lung cancer cells. We also included data from one COVID19 patients who underwent surgery for HNSCC.

The ethical committee of the Regina Elena National Cancer Institute and of University of Milan approved the study.

### Bioinformatic Analysis

#### Gene Expression Analysis

Analysis of 23 matched tumor and normal samples with gene expression profile from Affymetrix platform were background adjusted and quantile normalized. The gene expression values were obtained by using Robust Multiple-array Average (RMA) procedure.

MiRNAs expression for 66 matched tumor and normal samples from Agilent platform were analyzed as described in Ganci et al. (14).

mRNA expression data from IRE cohort used during the current study has been deposited in NCBI’s Gene Expression Omnibus and is accessible through GEO series accession number GSE107591 (https://www.ncbi.nlm.nih.gov/geo/query/acc.cgi?acc=GSE107591).

Normalized TCGA HNSC gene expression and miRNA expression of 478 tumor samples and 44 normal samples were obtained from Broad Institute TCGA Genome Data Analysis Center (http://gdac.broadinstitute.org/): Firehose stddata 2016_01_28. Broad Institute of MIT and Harvard. doi:10.7908/C11G0KM9

Significance of miRNA and gene modulation between expression values of normal and tumor samples was assessed by two-side paired or unpaired Student’s test and ANOVA test was used for comparisons among more than two groups. Significance was defined at the p<0.05 level.

A generalized linear model was fitted to evaluate linear regression of TMPRSS2 with immune signature, MYC-dependent gene signatures and clinical variables.

TMPRSS2 gene and protein expression data in normal tissues were obtained from EMBL-EBI Expression Atlas public repository (https://www.ebi.ac.uk/gxa/home).

#### Methylation Analysis

DNA methylation data of TCGA casuistry were obtained from Wanderer (http://maplab.imppc.org/wanderer/).

#### MiRNA target and Pathway analysis

We used miRWalk (http://mirwalk.umm.uni-heidelberg.de/) and miRNet (https://www.mirnet.ca/miRNet/home.xhtml) web tools for miRNA-target interaction prediction and pathway enrichment analysis. Spearman’s correlation coefficient was used to establish significance of negative association on patient samples for each predicted interaction.

#### Survival analysis

Disease-free survival (DFS) and overall survival (OS) were performed by using Kaplan-Meier analysis and the log-rank test was used to assess differences between curves. Patients with high and low signal intensity were defined by considering positive and negative z-score values, if not differently specified.

The correlation and regression analyses as well as the miRNA and gene modulation and the survival analysis were completely conducted with Matlab R2019.

### Cell cultures

Cal-27 (mutp53H193L) and Detroit-562 (mutp53R175H) cell lines were obtained from ATCC.

Cells were cultured in RPMI1640 (Cal27) and DMEM (Detroit 562) medium (Invitrogen-GIBCO) supplemented with 10% FBS, penicillin (100 U/mL), and streptomycin (100 mg/mL; Invitrogen-GIBCO). All cell lines were grown at 37° C in a balanced air humidified incubator with 5% CO2

### Cell transfection

The transfections were performed with Lipofectamine RNAiMax (Life Technologies). All experiments were conducted according to the manufacturer’s recommendations. siRNAs were purchased from Eurofins MWG and sequences are as follows: si-SCR: 5’-AAGUUCAGCGUGUCCGGGGAG-3’; si-MYC:5’GCCACAGCAUACAUCCUGU-3’; si-YAP: 5’-GACAUCUUCUGGUCAGAGA-3’; Si-p53: 5’-GACUCCAGUGGUAAUCUAC-3’. The cells were transfected for 48-72 hours according to the cell line and the experiments.

### RNA extraction and expression analysis

Total RNA from cells was extracted using the TRIzol Reagents (GIBCO) following the manufacturer’s instructions. RNA from FFPE samples was extracted using the miRneasy FFPE kit (QIAGEN) following the manufacturer’s instructions. The concentration and purity of total RNA was assessed using a Nanodrop TM1000 spectrophotometer (Nanodrop Technologies). Reverse transcription and qRT-PCR quantification were performed, respectively, by MMLV RT assay and SYBR Green or Taqman assays (Applied Biosystems) according to the manufacturer’s protocol. GAPDH and RNU48 were used as endogenous controls to standardize gene expression. Primers and Taqman assays used are indicated in supplementary table 2.

### SARS-CoV-2 detection

For the detection of SARS-CoV-2 in RNAs extracted from tissue samples we used Bosphore Novel Coronavirus (2019-nCoV) Detection Kit v2 (Anatolia GeneWork). This kit is a Real-Time PCR-based in vitro diagnostic medical device that allows to detect two regions of the virus in two separate reactions: E gene is used for screening purpose, where 2019-nCoV and also the closely related coronaviruses are detected, and the orf1ab target region is used to discriminate 2019-nCoV specifically. This kit includes also an internal control in order to check RNA extraction, PCR inhibition and application errors.

## Acknowledgements

Funding: GB acknowledges the support of AIRC IG 2017 – ID. 20613, Regione Lazio and MAECI Italy/USA bilateral grant program.

