## Supplementary material for "TMPRSS2, a SARS-CoV-2 internalization protease is downregulated in head and neck cancer patients": figures and tables

**Supplementary figure 1**

A

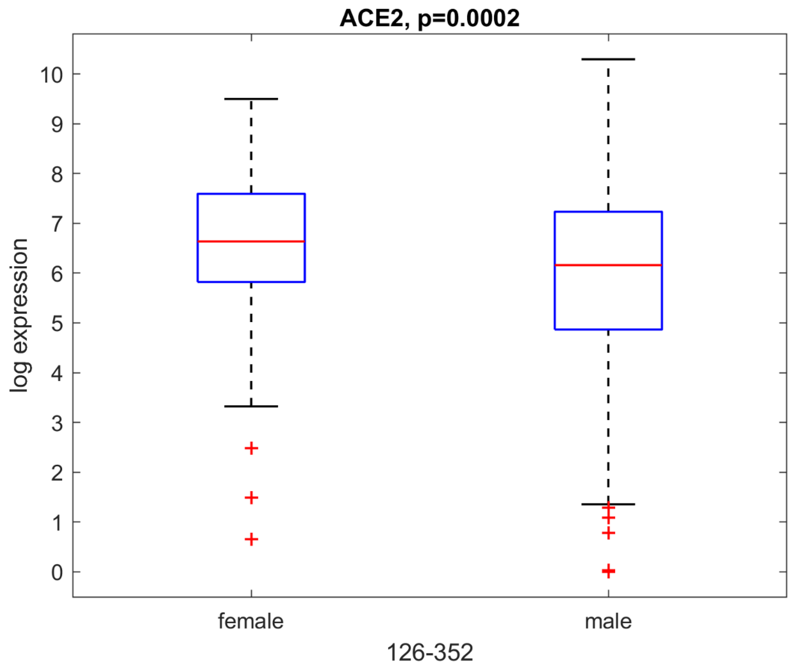


B

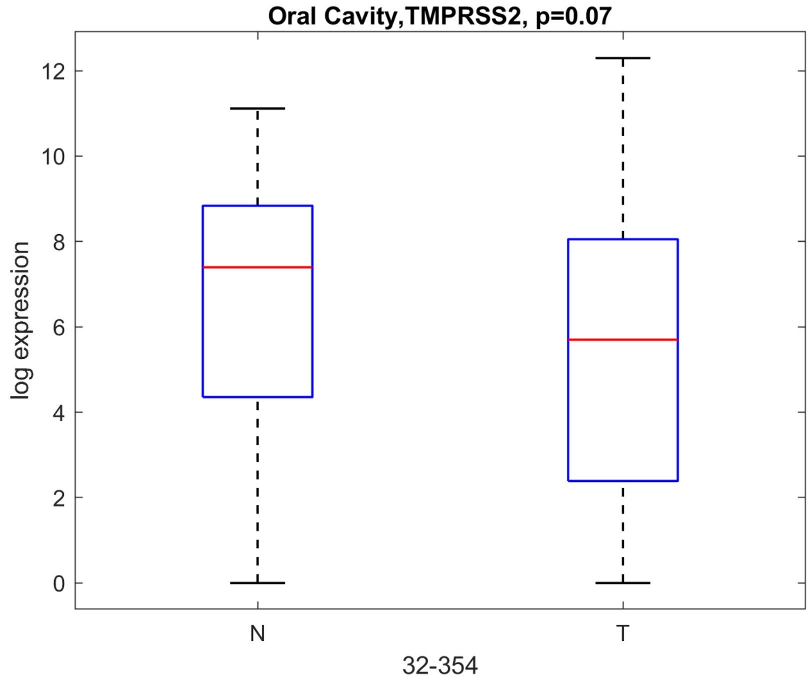

**Supplementary figure 2**

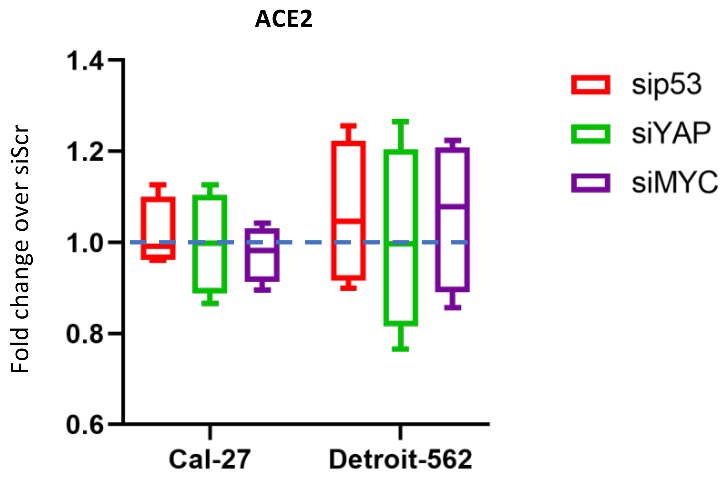

### Supplementary figure 3

A

**MYC signature**

| | | | | |
|---|---|---|---|---|
| MYC | SF3B3 | CCT2 | CCT7 | EIF3B |
| PA2G4 | PSMD1 | CCT3 | G3BP1 | PWP1 |
| POLE3 | PSMA2 | NME1 | CSTF2 | ODC1 |
| NOLC1 | RRP9 | HPRT1 | EIF3D | TCP1 |
| XRCC6 | HSP90AB1 | | | |

B

**immune signature**

| | | | | |
|---|---|---|---|---|
| CTLA4 | TNFRSF9 | KIR2DL3 | IL10 | IFNG |
| PDCD1 | TNFRSF4 | TNFRSF18 | CD40LG | LAG3 |
| CD274 | CD40 | ICOS | IDO1 | CD4 |
| KIR2DL1 | HAVCR2 | | | |

**Supplementary Table 1**

**MiRNA\TMPRSS2 predicted interactions (miRWalk)**

| mirnaid | refseqid | genesymbol | bindingp | position |
|---|---|---|---|---|
| hsa-miR-372-3p | NM_005656 | TMPRSS2 | 0.807692307692 | 3UTR |
| hsa-miR-520a-3p | NM_005656 | TMPRSS2 | 0.807692307692 | 3UTR |
| hsa-miR-587 | NM_005656 | TMPRSS2 | 0.807692307692 | 3UTR |
| hsa-miR-33b-3p | NM_005656 | TMPRSS2 | 0.807692307692 | 3UTR |
| hsa-miR-1224-3p | NM_005656 | TMPRSS2 | 0.807692307692 | 3UTR |
| hsa-miR-1227-3p | NM_005656 | TMPRSS2 | 0.807692307692 | 3UTR |
| hsa-miR-4284 | NM_005656 | TMPRSS2 | 0.807692307692 | 3UTR |
| hsa-miR-3616-3p | NM_005656 | TMPRSS2 | 0.807692307692 | 3UTR |
| hsa-miR-3620-3p | NM_005656 | TMPRSS2 | 0.807692307692 | 3UTR |
| hsa-miR-6762-3p | NM_005656 | TMPRSS2 | 0.807692307692 | 3UTR |
| hsa-miR-10527-5p | NM_005656 | TMPRSS2 | 0.807692307692 | 3UTR |
| hsa-miR-9851-3p | NM_005656 | TMPRSS2 | 0.807692307692 | 3UTR |
| hsa-miR-372-3p | NM_001135099 | TMPRSS2 | 0.807692307692 | 3UTR |
| hsa-miR-520a-3p | NM_001135099 | TMPRSS2 | 0.807692307692 | 3UTR |
| hsa-miR-513a-3p | NM_001135099 | TMPRSS2 | 0.807692307692 | 3UTR |
| hsa-miR-659-3p | NM_001135099 | TMPRSS2 | 0.807692307692 | 3UTR |
| hsa-miR-513c-3p | NM_001135099 | TMPRSS2 | 0.807692307692 | 3UTR |
| hsa-miR-4270 | NM_001135099 | TMPRSS2 | 0.807692307692 | 3UTR |
| hsa-miR-3616-3p | NM_001135099 | TMPRSS2 | 0.807692307692 | 3UTR |
| hsa-miR-3619-3p | NM_001135099 | TMPRSS2 | 0.807692307692 | 3UTR |
| hsa-miR-3620-3p | NM_001135099 | TMPRSS2 | 0.807692307692 | 3UTR |
| hsa-miR-4462 | NM_001135099 | TMPRSS2 | 0.807692307692 | 3UTR |
| hsa-miR-4494 | NM_001135099 | TMPRSS2 | 0.807692307692 | 3UTR |
| hsa-miR-3976 | NM_001135099 | TMPRSS2 | 0.807692307692 | 3UTR |
| hsa-miR-6827-5p | NM_001135099 | TMPRSS2 | 0.807692307692 | 3UTR |
| hsa-miR-6870-5p | NM_001135099 | TMPRSS2 | 0.807692307692 | 3UTR |
| hsa-miR-6881-5p | NM_001135099 | TMPRSS2 | 0.807692307692 | 3UTR |
| hsa-miR-7109-5p | NM_001135099 | TMPRSS2 | 0.807692307692 | 3UTR |
| hsa-miR-10527-5p | NM_001135099 | TMPRSS2 | 0.807692307692 | 3UTR |
| hsa-miR-3652 | NM_005656 | TMPRSS2 | 0.815384615385 | 3UTR |
| hsa-miR-6870-3p | NM_005656 | TMPRSS2 | 0.815384615385 | 3UTR |
| hsa-miR-491-3p | NM_001135099 | TMPRSS2 | 0.815384615385 | 3UTR |
| hsa-miR-6870-3p | NM_001135099 | TMPRSS2 | 0.815384615385 | 3UTR |
| hsa-miR-628-3p | NM_005656 | TMPRSS2 | 0.820512820513 | 3UTR |
| hsa-miR-4308 | NM_005656 | TMPRSS2 | 0.820512820513 | 3UTR |
| hsa-miR-6754-5p | NM_005656 | TMPRSS2 | 0.820512820513 | 3UTR |
| hsa-miR-423-5p | NM_001135099 | TMPRSS2 | 0.820512820513 | 3UTR |
| hsa-miR-3175 | NM_001135099 | TMPRSS2 | 0.820512820513 | 3UTR |
| hsa-miR-6809-5p | NM_001135099 | TMPRSS2 | 0.820512820513 | 3UTR |
| hsa-miR-12120 | NM_001135099 | TMPRSS2 | 0.826923076923 | 3UTR |
| hsa-miR-500b-3p | NM_005656 | TMPRSS2 | 0.835164835165 | 3UTR |

| | | | | |
|---|---|---|---|---|
| hsa-miR-19b-3p | NM_005656 | TMPRSS2 | 0.846153846154 | 3UTR |
| hsa-miR-20a-5p | NM_005656 | TMPRSS2 | 0.846153846154 | 3UTR |
| hsa-miR-24-1-5p | NM_005656 | TMPRSS2 | 0.846153846154 | 3UTR |
| hsa-miR-26a-5p | NM_005656 | TMPRSS2 | 0.846153846154 | 3UTR |
| hsa-miR-28-5p | NM_005656 | TMPRSS2 | 0.846153846154 | 3UTR |
| hsa-miR-31-3p | NM_005656 | TMPRSS2 | 0.846153846154 | 3UTR |
| hsa-miR-92a-1-5p | NM_005656 | TMPRSS2 | 0.846153846154 | 3UTR |
| hsa-miR-103a-3p | NM_005656 | TMPRSS2 | 0.846153846154 | 3UTR |
| hsa-miR-105-3p | NM_005656 | TMPRSS2 | 0.846153846154 | 3UTR |
| hsa-miR-107 | NM_005656 | TMPRSS2 | 0.846153846154 | 3UTR |
| hsa-miR-192-3p | NM_005656 | TMPRSS2 | 0.846153846154 | 3UTR |
| hsa-miR-197-5p | NM_005656 | TMPRSS2 | 0.846153846154 | 3UTR |
| hsa-miR-129-5p | NM_005656 | TMPRSS2 | 0.846153846154 | 3UTR |
| hsa-miR-148a-3p | NM_005656 | TMPRSS2 | 0.846153846154 | 3UTR |
| hsa-miR-30c-2-3p | NM_005656 | TMPRSS2 | 0.846153846154 | 3UTR |
| hsa-miR-30c-2-3p | NM_005656 | TMPRSS2 | 0.846153846154 | 3UTR |
| hsa-miR-139-5p | NM_005656 | TMPRSS2 | 0.846153846154 | 3UTR |
| hsa-miR-181b-5p | NM_005656 | TMPRSS2 | 0.846153846154 | 3UTR |
| hsa-miR-181c-5p | NM_005656 | TMPRSS2 | 0.846153846154 | 3UTR |
| hsa-miR-181c-3p | NM_005656 | TMPRSS2 | 0.846153846154 | 3UTR |
| hsa-miR-183-3p | NM_005656 | TMPRSS2 | 0.846153846154 | 3UTR |
| hsa-miR-204-3p | NM_005656 | TMPRSS2 | 0.846153846154 | 3UTR |
| hsa-miR-205-5p | NM_005656 | TMPRSS2 | 0.846153846154 | 3UTR |
| hsa-miR-212-3p | NM_005656 | TMPRSS2 | 0.846153846154 | 3UTR |
| hsa-miR-214-3p | NM_005656 | TMPRSS2 | 0.846153846154 | 3UTR |
| hsa-miR-217-5p | NM_005656 | TMPRSS2 | 0.846153846154 | 3UTR |
| hsa-miR-219a-1-3p | NM_005656 | TMPRSS2 | 0.846153846154 | 3UTR |
| hsa-miR-222-3p | NM_005656 | TMPRSS2 | 0.846153846154 | 3UTR |
| hsa-let-7g-3p | NM_005656 | TMPRSS2 | 0.846153846154 | 3UTR |
| hsa-miR-15b-5p | NM_005656 | TMPRSS2 | 0.846153846154 | 3UTR |
| hsa-miR-23b-3p | NM_005656 | TMPRSS2 | 0.846153846154 | 3UTR |
| hsa-miR-122-3p | NM_005656 | TMPRSS2 | 0.846153846154 | 3UTR |
| hsa-miR-138-2-3p | NM_005656 | TMPRSS2 | 0.846153846154 | 3UTR |
| hsa-miR-141-5p | NM_005656 | TMPRSS2 | 0.846153846154 | 3UTR |
| hsa-miR-143-5p | NM_005656 | TMPRSS2 | 0.846153846154 | 3UTR |
| hsa-miR-145-5p | NM_005656 | TMPRSS2 | 0.846153846154 | 3UTR |
| hsa-miR-191-5p | NM_005656 | TMPRSS2 | 0.846153846154 | 3UTR |
| hsa-miR-9-3p | NM_005656 | TMPRSS2 | 0.846153846154 | 3UTR |
| hsa-miR-138-1-3p | NM_005656 | TMPRSS2 | 0.846153846154 | 3UTR |
| hsa-miR-150-5p | NM_005656 | TMPRSS2 | 0.846153846154 | 3UTR |
| hsa-miR-185-5p | NM_005656 | TMPRSS2 | 0.846153846154 | 3UTR |
| hsa-miR-185-3p | NM_005656 | TMPRSS2 | 0.846153846154 | 3UTR |
| hsa-miR-188-5p | NM_005656 | TMPRSS2 | 0.846153846154 | 3UTR |
| hsa-miR-195-3p | NM_005656 | TMPRSS2 | 0.846153846154 | 3UTR |
| hsa-miR-200c-5p | NM_005656 | TMPRSS2 | 0.846153846154 | 3UTR |
| hsa-miR-155-5p | NM_005656 | TMPRSS2 | 0.846153846154 | 3UTR |
| hsa-miR-301a-3p | NM_005656 | TMPRSS2 | 0.846153846154 | 3UTR |

| | | | | |
|---|---|---|---|---|
| hsa-miR-130b-5p | NM_005656 | TMPRSS2 | 0.846153846154 | 3UTR |
| hsa-miR-130b-5p | NM_005656 | TMPRSS2 | 0.846153846154 | 3UTR |
| hsa-miR-365a-5p | NM_005656 | TMPRSS2 | 0.846153846154 | 3UTR |
| hsa-miR-365b-5p | NM_005656 | TMPRSS2 | 0.846153846154 | 3UTR |
| hsa-miR-302d-5p | NM_005656 | TMPRSS2 | 0.846153846154 | 3UTR |
| hsa-miR-370-3p | NM_005656 | TMPRSS2 | 0.846153846154 | 3UTR |
| hsa-miR-371a-3p | NM_005656 | TMPRSS2 | 0.846153846154 | 3UTR |
| hsa-miR-378a-3p | NM_005656 | TMPRSS2 | 0.846153846154 | 3UTR |
| hsa-miR-378a-3p | NM_005656 | TMPRSS2 | 0.846153846154 | 3UTR |
| hsa-miR-383-3p | NM_005656 | TMPRSS2 | 0.846153846154 | 3UTR |
| hsa-miR-328-5p | NM_005656 | TMPRSS2 | 0.846153846154 | 3UTR |
| hsa-miR-342-5p | NM_005656 | TMPRSS2 | 0.846153846154 | 3UTR |
| hsa-miR-342-5p | NM_005656 | TMPRSS2 | 0.846153846154 | 3UTR |
| hsa-miR-326 | NM_005656 | TMPRSS2 | 0.846153846154 | 3UTR |
| hsa-miR-135b-5p | NM_005656 | TMPRSS2 | 0.846153846154 | 3UTR |
| hsa-miR-331-3p | NM_005656 | TMPRSS2 | 0.846153846154 | 3UTR |
| hsa-miR-345-5p | NM_005656 | TMPRSS2 | 0.846153846154 | 3UTR |
| hsa-miR-345-3p | NM_005656 | TMPRSS2 | 0.846153846154 | 3UTR |
| hsa-miR-425-5p | NM_005656 | TMPRSS2 | 0.846153846154 | 3UTR |
| hsa-miR-425-3p | NM_005656 | TMPRSS2 | 0.846153846154 | 3UTR |
| hsa-miR-20b-5p | NM_005656 | TMPRSS2 | 0.846153846154 | 3UTR |
| hsa-miR-433-5p | NM_005656 | TMPRSS2 | 0.846153846154 | 3UTR |
| hsa-miR-329-3p | NM_005656 | TMPRSS2 | 0.846153846154 | 3UTR |
| hsa-miR-409-5p | NM_005656 | TMPRSS2 | 0.846153846154 | 3UTR |
| hsa-miR-484 | NM_005656 | TMPRSS2 | 0.846153846154 | 3UTR |
| hsa-miR-490-5p | NM_005656 | TMPRSS2 | 0.846153846154 | 3UTR |
| hsa-miR-490-3p | NM_005656 | TMPRSS2 | 0.846153846154 | 3UTR |
| hsa-miR-490-3p | NM_005656 | TMPRSS2 | 0.846153846154 | 3UTR |
| hsa-miR-491-3p | NM_005656 | TMPRSS2 | 0.846153846154 | 3UTR |
| hsa-miR-432-5p | NM_005656 | TMPRSS2 | 0.846153846154 | 3UTR |
| hsa-miR-494-3p | NM_005656 | TMPRSS2 | 0.846153846154 | 3UTR |
| hsa-miR-193b-3p | NM_005656 | TMPRSS2 | 0.846153846154 | 3UTR |
| hsa-miR-498-5p | NM_005656 | TMPRSS2 | 0.846153846154 | 3UTR |
| hsa-miR-520e-3p | NM_005656 | TMPRSS2 | 0.846153846154 | 3UTR |
| hsa-miR-520f-5p | NM_005656 | TMPRSS2 | 0.846153846154 | 3UTR |
| hsa-miR-519c-3p | NM_005656 | TMPRSS2 | 0.846153846154 | 3UTR |
| hsa-miR-525-3p | NM_005656 | TMPRSS2 | 0.846153846154 | 3UTR |
| hsa-miR-525-3p | NM_005656 | TMPRSS2 | 0.846153846154 | 3UTR |
| hsa-miR-518f-3p | NM_005656 | TMPRSS2 | 0.846153846154 | 3UTR |
| hsa-miR-520b-5p | NM_005656 | TMPRSS2 | 0.846153846154 | 3UTR |
| hsa-miR-518b | NM_005656 | TMPRSS2 | 0.846153846154 | 3UTR |
| hsa-miR-526a-5p | NM_005656 | TMPRSS2 | 0.846153846154 | 3UTR |
| hsa-miR-520c-5p | NM_005656 | TMPRSS2 | 0.846153846154 | 3UTR |
| hsa-miR-518c-3p | NM_005656 | TMPRSS2 | 0.846153846154 | 3UTR |
| hsa-miR-518d-5p | NM_005656 | TMPRSS2 | 0.846153846154 | 3UTR |
| hsa-miR-522-3p | NM_005656 | TMPRSS2 | 0.846153846154 | 3UTR |
| hsa-miR-519a-2-5p | NM_005656 | TMPRSS2 | 0.846153846154 | 3UTR |

| | | | | |
|---|---|---|---|---|
| hsa-miR-499a-3p | NM_005656 | TMPRSS2 | 0.846153846154 | 3UTR |
| hsa-miR-500a-3p | NM_005656 | TMPRSS2 | 0.846153846154 | 3UTR |
| hsa-miR-501-3p | NM_005656 | TMPRSS2 | 0.846153846154 | 3UTR |
| hsa-miR-502-5p | NM_005656 | TMPRSS2 | 0.846153846154 | 3UTR |
| hsa-miR-502-5p | NM_005656 | TMPRSS2 | 0.846153846154 | 3UTR |
| hsa-miR-450a-2-3p | NM_005656 | TMPRSS2 | 0.846153846154 | 3UTR |
| hsa-miR-503-5p | NM_005656 | TMPRSS2 | 0.846153846154 | 3UTR |
| hsa-miR-503-5p | NM_005656 | TMPRSS2 | 0.846153846154 | 3UTR |
| hsa-miR-513a-3p | NM_005656 | TMPRSS2 | 0.846153846154 | 3UTR |
| hsa-miR-508-5p | NM_005656 | TMPRSS2 | 0.846153846154 | 3UTR |
| hsa-miR-510-3p | NM_005656 | TMPRSS2 | 0.846153846154 | 3UTR |
| hsa-miR-551a | NM_005656 | TMPRSS2 | 0.846153846154 | 3UTR |
| hsa-miR-552-3p | NM_005656 | TMPRSS2 | 0.846153846154 | 3UTR |
| hsa-miR-554 | NM_005656 | TMPRSS2 | 0.846153846154 | 3UTR |
| hsa-miR-571 | NM_005656 | TMPRSS2 | 0.846153846154 | 3UTR |
| hsa-miR-584-5p | NM_005656 | TMPRSS2 | 0.846153846154 | 3UTR |
| hsa-miR-595 | NM_005656 | TMPRSS2 | 0.846153846154 | 3UTR |
| hsa-miR-602 | NM_005656 | TMPRSS2 | 0.846153846154 | 3UTR |
| hsa-miR-615-3p | NM_005656 | TMPRSS2 | 0.846153846154 | 3UTR |
| hsa-miR-616-3p | NM_005656 | TMPRSS2 | 0.846153846154 | 3UTR |
| hsa-miR-622 | NM_005656 | TMPRSS2 | 0.846153846154 | 3UTR |
| hsa-miR-625-5p | NM_005656 | TMPRSS2 | 0.846153846154 | 3UTR |
| hsa-miR-630 | NM_005656 | TMPRSS2 | 0.846153846154 | 3UTR |
| hsa-miR-630 | NM_005656 | TMPRSS2 | 0.846153846154 | 3UTR |
| hsa-miR-635 | NM_005656 | TMPRSS2 | 0.846153846154 | 3UTR |
| hsa-miR-642a-3p | NM_005656 | TMPRSS2 | 0.846153846154 | 3UTR |
| hsa-miR-644a | NM_005656 | TMPRSS2 | 0.846153846154 | 3UTR |
| hsa-miR-647 | NM_005656 | TMPRSS2 | 0.846153846154 | 3UTR |
| hsa-miR-650 | NM_005656 | TMPRSS2 | 0.846153846154 | 3UTR |
| hsa-miR-650 | NM_005656 | TMPRSS2 | 0.846153846154 | 3UTR |
| hsa-miR-449b-5p | NM_005656 | TMPRSS2 | 0.846153846154 | 3UTR |
| hsa-miR-654-5p | NM_005656 | TMPRSS2 | 0.846153846154 | 3UTR |
| hsa-miR-542-5p | NM_005656 | TMPRSS2 | 0.846153846154 | 3UTR |
| hsa-miR-542-5p | NM_005656 | TMPRSS2 | 0.846153846154 | 3UTR |
| hsa-miR-767-5p | NM_005656 | TMPRSS2 | 0.846153846154 | 3UTR |
| hsa-miR-767-3p | NM_005656 | TMPRSS2 | 0.846153846154 | 3UTR |
| hsa-miR-320c | NM_005656 | TMPRSS2 | 0.846153846154 | 3UTR |
| hsa-miR-1323 | NM_005656 | TMPRSS2 | 0.846153846154 | 3UTR |
| hsa-miR-1271-5p | NM_005656 | TMPRSS2 | 0.846153846154 | 3UTR |
| hsa-miR-764 | NM_005656 | TMPRSS2 | 0.846153846154 | 3UTR |
| hsa-miR-765 | NM_005656 | TMPRSS2 | 0.846153846154 | 3UTR |
| hsa-miR-770-5p | NM_005656 | TMPRSS2 | 0.846153846154 | 3UTR |
| hsa-miR-874-5p | NM_005656 | TMPRSS2 | 0.846153846154 | 3UTR |
| hsa-miR-541-5p | NM_005656 | TMPRSS2 | 0.846153846154 | 3UTR |
| hsa-miR-875-3p | NM_005656 | TMPRSS2 | 0.846153846154 | 3UTR |
| hsa-miR-877-5p | NM_005656 | TMPRSS2 | 0.846153846154 | 3UTR |
| hsa-miR-877-3p | NM_005656 | TMPRSS2 | 0.846153846154 | 3UTR |

| | | | | |
|---|---|---|---|---|
| hsa-miR-665 | NM_005656 | TMPRSS2 | 0.846153846154 | 3UTR |
| hsa-miR-301b-5p | NM_005656 | TMPRSS2 | 0.846153846154 | 3UTR |
| hsa-miR-920 | NM_005656 | TMPRSS2 | 0.846153846154 | 3UTR |
| hsa-miR-922 | NM_005656 | TMPRSS2 | 0.846153846154 | 3UTR |
| hsa-miR-934 | NM_005656 | TMPRSS2 | 0.846153846154 | 3UTR |
| hsa-miR-1229-3p | NM_005656 | TMPRSS2 | 0.846153846154 | 3UTR |
| hsa-miR-1229-3p | NM_005656 | TMPRSS2 | 0.846153846154 | 3UTR |
| hsa-miR-663b | NM_005656 | TMPRSS2 | 0.846153846154 | 3UTR |
| hsa-miR-1207-5p | NM_005656 | TMPRSS2 | 0.846153846154 | 3UTR |
| hsa-miR-1286 | NM_005656 | TMPRSS2 | 0.846153846154 | 3UTR |
| hsa-miR-1248 | NM_005656 | TMPRSS2 | 0.846153846154 | 3UTR |
| hsa-miR-1249-5p | NM_005656 | TMPRSS2 | 0.846153846154 | 3UTR |
| hsa-miR-1307-5p | NM_005656 | TMPRSS2 | 0.846153846154 | 3UTR |
| hsa-miR-1307-5p | NM_005656 | TMPRSS2 | 0.846153846154 | 3UTR |
| hsa-miR-513c-3p | NM_005656 | TMPRSS2 | 0.846153846154 | 3UTR |
| hsa-miR-320d | NM_005656 | TMPRSS2 | 0.846153846154 | 3UTR |
| hsa-miR-1912-3p | NM_005656 | TMPRSS2 | 0.846153846154 | 3UTR |
| hsa-miR-1913 | NM_005656 | TMPRSS2 | 0.846153846154 | 3UTR |
| hsa-miR-1914-3p | NM_005656 | TMPRSS2 | 0.846153846154 | 3UTR |
| hsa-miR-1976 | NM_005656 | TMPRSS2 | 0.846153846154 | 3UTR |
| hsa-miR-2110 | NM_005656 | TMPRSS2 | 0.846153846154 | 3UTR |
| hsa-miR-2115-5p | NM_005656 | TMPRSS2 | 0.846153846154 | 3UTR |
| hsa-miR-2116-5p | NM_005656 | TMPRSS2 | 0.846153846154 | 3UTR |
| hsa-miR-2116-3p | NM_005656 | TMPRSS2 | 0.846153846154 | 3UTR |
| hsa-miR-2278 | NM_005656 | TMPRSS2 | 0.846153846154 | 3UTR |
| hsa-miR-2278 | NM_005656 | TMPRSS2 | 0.846153846154 | 3UTR |
| hsa-miR-2681-5p | NM_005656 | TMPRSS2 | 0.846153846154 | 3UTR |
| hsa-miR-3117-3p | NM_005656 | TMPRSS2 | 0.846153846154 | 3UTR |
| hsa-miR-3125 | NM_005656 | TMPRSS2 | 0.846153846154 | 3UTR |
| hsa-miR-3126-5p | NM_005656 | TMPRSS2 | 0.846153846154 | 3UTR |
| hsa-miR-3127-5p | NM_005656 | TMPRSS2 | 0.846153846154 | 3UTR |
| hsa-miR-3129-3p | NM_005656 | TMPRSS2 | 0.846153846154 | 3UTR |
| hsa-miR-3130-3p | NM_005656 | TMPRSS2 | 0.846153846154 | 3UTR |
| hsa-miR-3131 | NM_005656 | TMPRSS2 | 0.846153846154 | 3UTR |
| hsa-miR-378b | NM_005656 | TMPRSS2 | 0.846153846154 | 3UTR |
| hsa-miR-3141 | NM_005656 | TMPRSS2 | 0.846153846154 | 3UTR |
| hsa-miR-3142 | NM_005656 | TMPRSS2 | 0.846153846154 | 3UTR |
| hsa-miR-3147 | NM_005656 | TMPRSS2 | 0.846153846154 | 3UTR |
| hsa-miR-3074-3p | NM_005656 | TMPRSS2 | 0.846153846154 | 3UTR |
| hsa-miR-3155a | NM_005656 | TMPRSS2 | 0.846153846154 | 3UTR |
| hsa-miR-3158-3p | NM_005656 | TMPRSS2 | 0.846153846154 | 3UTR |
| hsa-miR-3160-3p | NM_005656 | TMPRSS2 | 0.846153846154 | 3UTR |
| hsa-miR-3162-5p | NM_005656 | TMPRSS2 | 0.846153846154 | 3UTR |
| hsa-miR-3163 | NM_005656 | TMPRSS2 | 0.846153846154 | 3UTR |
| hsa-miR-3163 | NM_005656 | TMPRSS2 | 0.846153846154 | 3UTR |
| hsa-miR-3169 | NM_005656 | TMPRSS2 | 0.846153846154 | 3UTR |
| hsa-miR-3183 | NM_005656 | TMPRSS2 | 0.846153846154 | 3UTR |

| | | | | |
|---|---|---|---|---|
| hsa-miR-3065-3p | NM_005656 | TMPRSS2 | 0.846153846154 | 3UTR |
| hsa-miR-3189-3p | NM_005656 | TMPRSS2 | 0.846153846154 | 3UTR |
| hsa-miR-3192-5p | NM_005656 | TMPRSS2 | 0.846153846154 | 3UTR |
| hsa-miR-514b-3p | NM_005656 | TMPRSS2 | 0.846153846154 | 3UTR |
| hsa-miR-4296 | NM_005656 | TMPRSS2 | 0.846153846154 | 3UTR |
| hsa-miR-378c | NM_005656 | TMPRSS2 | 0.846153846154 | 3UTR |
| hsa-miR-4293 | NM_005656 | TMPRSS2 | 0.846153846154 | 3UTR |
| hsa-miR-4298 | NM_005656 | TMPRSS2 | 0.846153846154 | 3UTR |
| hsa-miR-4300 | NM_005656 | TMPRSS2 | 0.846153846154 | 3UTR |
| hsa-miR-4320 | NM_005656 | TMPRSS2 | 0.846153846154 | 3UTR |
| hsa-miR-4317 | NM_005656 | TMPRSS2 | 0.846153846154 | 3UTR |
| hsa-miR-4323 | NM_005656 | TMPRSS2 | 0.846153846154 | 3UTR |
| hsa-miR-4257 | NM_005656 | TMPRSS2 | 0.846153846154 | 3UTR |
| hsa-miR-4254 | NM_005656 | TMPRSS2 | 0.846153846154 | 3UTR |
| hsa-miR-4252 | NM_005656 | TMPRSS2 | 0.846153846154 | 3UTR |
| hsa-miR-4266 | NM_005656 | TMPRSS2 | 0.846153846154 | 3UTR |
| hsa-miR-2355-5p | NM_005656 | TMPRSS2 | 0.846153846154 | 3UTR |
| hsa-miR-4269 | NM_005656 | TMPRSS2 | 0.846153846154 | 3UTR |
| hsa-miR-4271 | NM_005656 | TMPRSS2 | 0.846153846154 | 3UTR |
| hsa-miR-4273 | NM_005656 | TMPRSS2 | 0.846153846154 | 3UTR |
| hsa-miR-4279 | NM_005656 | TMPRSS2 | 0.846153846154 | 3UTR |
| hsa-miR-4278 | NM_005656 | TMPRSS2 | 0.846153846154 | 3UTR |
| hsa-miR-4282 | NM_005656 | TMPRSS2 | 0.846153846154 | 3UTR |
| hsa-miR-4285 | NM_005656 | TMPRSS2 | 0.846153846154 | 3UTR |
| hsa-miR-4291 | NM_005656 | TMPRSS2 | 0.846153846154 | 3UTR |
| hsa-miR-3610 | NM_005656 | TMPRSS2 | 0.846153846154 | 3UTR |
| hsa-miR-3610 | NM_005656 | TMPRSS2 | 0.846153846154 | 3UTR |
| hsa-miR-3619-3p | NM_005656 | TMPRSS2 | 0.846153846154 | 3UTR |
| hsa-miR-23c | NM_005656 | TMPRSS2 | 0.846153846154 | 3UTR |
| hsa-miR-3621 | NM_005656 | TMPRSS2 | 0.846153846154 | 3UTR |
| hsa-miR-3622b-5p | NM_005656 | TMPRSS2 | 0.846153846154 | 3UTR |
| hsa-miR-3650 | NM_005656 | TMPRSS2 | 0.846153846154 | 3UTR |
| hsa-miR-3654 | NM_005656 | TMPRSS2 | 0.846153846154 | 3UTR |
| hsa-miR-3654 | NM_005656 | TMPRSS2 | 0.846153846154 | 3UTR |
| hsa-miR-3659 | NM_005656 | TMPRSS2 | 0.846153846154 | 3UTR |
| hsa-miR-3664-5p | NM_005656 | TMPRSS2 | 0.846153846154 | 3UTR |
| hsa-miR-3665 | NM_005656 | TMPRSS2 | 0.846153846154 | 3UTR |
| hsa-miR-3665 | NM_005656 | TMPRSS2 | 0.846153846154 | 3UTR |
| hsa-miR-3675-3p | NM_005656 | TMPRSS2 | 0.846153846154 | 3UTR |
| hsa-miR-3679-3p | NM_005656 | TMPRSS2 | 0.846153846154 | 3UTR |
| hsa-miR-3680-5p | NM_005656 | TMPRSS2 | 0.846153846154 | 3UTR |
| hsa-miR-3685 | NM_005656 | TMPRSS2 | 0.846153846154 | 3UTR |
| hsa-miR-3689a-5p | NM_005656 | TMPRSS2 | 0.846153846154 | 3UTR |
| hsa-miR-3692-3p | NM_005656 | TMPRSS2 | 0.846153846154 | 3UTR |
| hsa-miR-3714 | NM_005656 | TMPRSS2 | 0.846153846154 | 3UTR |
| hsa-miR-3689b-5p | NM_005656 | TMPRSS2 | 0.846153846154 | 3UTR |
| hsa-miR-3908 | NM_005656 | TMPRSS2 | 0.846153846154 | 3UTR |

| | | | | |
|---|---|---|---|---|
| hsa-miR-3909 | NM_005656 | TMPRSS2 | 0.846153846154 | 3UTR |
| hsa-miR-3911 | NM_005656 | TMPRSS2 | 0.846153846154 | 3UTR |
| hsa-miR-3916 | NM_005656 | TMPRSS2 | 0.846153846154 | 3UTR |
| hsa-miR-3917 | NM_005656 | TMPRSS2 | 0.846153846154 | 3UTR |
| hsa-miR-3924 | NM_005656 | TMPRSS2 | 0.846153846154 | 3UTR |
| hsa-miR-3934-5p | NM_005656 | TMPRSS2 | 0.846153846154 | 3UTR |
| hsa-miR-3934-5p | NM_005656 | TMPRSS2 | 0.846153846154 | 3UTR |
| hsa-miR-3934-5p | NM_005656 | TMPRSS2 | 0.846153846154 | 3UTR |
| hsa-miR-3934-3p | NM_005656 | TMPRSS2 | 0.846153846154 | 3UTR |
| hsa-miR-3936 | NM_005656 | TMPRSS2 | 0.846153846154 | 3UTR |
| hsa-miR-3943 | NM_005656 | TMPRSS2 | 0.846153846154 | 3UTR |
| hsa-miR-374c-3p | NM_005656 | TMPRSS2 | 0.846153846154 | 3UTR |
| hsa-miR-4420 | NM_005656 | TMPRSS2 | 0.846153846154 | 3UTR |
| hsa-miR-4420 | NM_005656 | TMPRSS2 | 0.846153846154 | 3UTR |
| hsa-miR-4421 | NM_005656 | TMPRSS2 | 0.846153846154 | 3UTR |
| hsa-miR-4421 | NM_005656 | TMPRSS2 | 0.846153846154 | 3UTR |
| hsa-miR-4428 | NM_005656 | TMPRSS2 | 0.846153846154 | 3UTR |
| hsa-miR-4433a-5p | NM_005656 | TMPRSS2 | 0.846153846154 | 3UTR |
| hsa-miR-4433a-3p | NM_005656 | TMPRSS2 | 0.846153846154 | 3UTR |
| hsa-miR-4433a-3p | NM_005656 | TMPRSS2 | 0.846153846154 | 3UTR |
| hsa-miR-4433a-3p | NM_005656 | TMPRSS2 | 0.846153846154 | 3UTR |
| hsa-miR-4437 | NM_005656 | TMPRSS2 | 0.846153846154 | 3UTR |
| hsa-miR-4448 | NM_005656 | TMPRSS2 | 0.846153846154 | 3UTR |
| hsa-miR-4458 | NM_005656 | TMPRSS2 | 0.846153846154 | 3UTR |
| hsa-miR-4472 | NM_005656 | TMPRSS2 | 0.846153846154 | 3UTR |
| hsa-miR-4473 | NM_005656 | TMPRSS2 | 0.846153846154 | 3UTR |
| hsa-miR-4476 | NM_005656 | TMPRSS2 | 0.846153846154 | 3UTR |
| hsa-miR-4478 | NM_005656 | TMPRSS2 | 0.846153846154 | 3UTR |
| hsa-miR-3689e | NM_005656 | TMPRSS2 | 0.846153846154 | 3UTR |
| hsa-miR-4479 | NM_005656 | TMPRSS2 | 0.846153846154 | 3UTR |
| hsa-miR-3155b | NM_005656 | TMPRSS2 | 0.846153846154 | 3UTR |
| hsa-miR-4485-5p | NM_005656 | TMPRSS2 | 0.846153846154 | 3UTR |
| hsa-miR-4487 | NM_005656 | TMPRSS2 | 0.846153846154 | 3UTR |
| hsa-miR-4496 | NM_005656 | TMPRSS2 | 0.846153846154 | 3UTR |
| hsa-miR-4498 | NM_005656 | TMPRSS2 | 0.846153846154 | 3UTR |
| hsa-miR-4500 | NM_005656 | TMPRSS2 | 0.846153846154 | 3UTR |
| hsa-miR-4502 | NM_005656 | TMPRSS2 | 0.846153846154 | 3UTR |
| hsa-miR-2392 | NM_005656 | TMPRSS2 | 0.846153846154 | 3UTR |
| hsa-miR-4514 | NM_005656 | TMPRSS2 | 0.846153846154 | 3UTR |
| hsa-miR-4518 | NM_005656 | TMPRSS2 | 0.846153846154 | 3UTR |
| hsa-miR-4524a-5p | NM_005656 | TMPRSS2 | 0.846153846154 | 3UTR |
| hsa-miR-4535 | NM_005656 | TMPRSS2 | 0.846153846154 | 3UTR |
| hsa-miR-1587 | NM_005656 | TMPRSS2 | 0.846153846154 | 3UTR |
| hsa-miR-4536-5p | NM_005656 | TMPRSS2 | 0.846153846154 | 3UTR |
| hsa-miR-3960 | NM_005656 | TMPRSS2 | 0.846153846154 | 3UTR |
| hsa-miR-3975 | NM_005656 | TMPRSS2 | 0.846153846154 | 3UTR |
| hsa-miR-4632-5p | NM_005656 | TMPRSS2 | 0.846153846154 | 3UTR |

| | | | | |
|---|---|---|---|---|
| hsa-miR-4638-3p | NM_005656 | TMPRSS2 | 0.846153846154 | 3UTR |
| hsa-miR-4640-3p | NM_005656 | TMPRSS2 | 0.846153846154 | 3UTR |
| hsa-miR-4642 | NM_005656 | TMPRSS2 | 0.846153846154 | 3UTR |
| hsa-miR-4645-5p | NM_005656 | TMPRSS2 | 0.846153846154 | 3UTR |
| hsa-miR-4648 | NM_005656 | TMPRSS2 | 0.846153846154 | 3UTR |
| hsa-miR-4655-5p | NM_005656 | TMPRSS2 | 0.846153846154 | 3UTR |
| hsa-miR-4661-3p | NM_005656 | TMPRSS2 | 0.846153846154 | 3UTR |
| hsa-miR-4664-5p | NM_005656 | TMPRSS2 | 0.846153846154 | 3UTR |
| hsa-miR-4669 | NM_005656 | TMPRSS2 | 0.846153846154 | 3UTR |
| hsa-miR-4671-3p | NM_005656 | TMPRSS2 | 0.846153846154 | 3UTR |
| hsa-miR-4681 | NM_005656 | TMPRSS2 | 0.846153846154 | 3UTR |
| hsa-miR-4685-5p | NM_005656 | TMPRSS2 | 0.846153846154 | 3UTR |
| hsa-miR-4685-5p | NM_005656 | TMPRSS2 | 0.846153846154 | 3UTR |
| hsa-miR-1343-3p | NM_005656 | TMPRSS2 | 0.846153846154 | 3UTR |
| hsa-miR-4690-5p | NM_005656 | TMPRSS2 | 0.846153846154 | 3UTR |
| hsa-miR-4695-5p | NM_005656 | TMPRSS2 | 0.846153846154 | 3UTR |
| hsa-miR-4696 | NM_005656 | TMPRSS2 | 0.846153846154 | 3UTR |
| hsa-miR-4700-5p | NM_005656 | TMPRSS2 | 0.846153846154 | 3UTR |
| hsa-miR-4701-5p | NM_005656 | TMPRSS2 | 0.846153846154 | 3UTR |
| hsa-miR-203b-5p | NM_005656 | TMPRSS2 | 0.846153846154 | 3UTR |
| hsa-miR-4710 | NM_005656 | TMPRSS2 | 0.846153846154 | 3UTR |
| hsa-miR-4712-3p | NM_005656 | TMPRSS2 | 0.846153846154 | 3UTR |
| hsa-miR-4716-3p | NM_005656 | TMPRSS2 | 0.846153846154 | 3UTR |
| hsa-miR-4723-3p | NM_005656 | TMPRSS2 | 0.846153846154 | 3UTR |
| hsa-miR-451b | NM_005656 | TMPRSS2 | 0.846153846154 | 3UTR |
| hsa-miR-4724-5p | NM_005656 | TMPRSS2 | 0.846153846154 | 3UTR |
| hsa-miR-4731-5p | NM_005656 | TMPRSS2 | 0.846153846154 | 3UTR |
| hsa-miR-4736 | NM_005656 | TMPRSS2 | 0.846153846154 | 3UTR |
| hsa-miR-3064-3p | NM_005656 | TMPRSS2 | 0.846153846154 | 3UTR |
| hsa-miR-4739 | NM_005656 | TMPRSS2 | 0.846153846154 | 3UTR |
| hsa-miR-4742-5p | NM_005656 | TMPRSS2 | 0.846153846154 | 3UTR |
| hsa-miR-4747-5p | NM_005656 | TMPRSS2 | 0.846153846154 | 3UTR |
| hsa-miR-4747-3p | NM_005656 | TMPRSS2 | 0.846153846154 | 3UTR |
| hsa-miR-4748 | NM_005656 | TMPRSS2 | 0.846153846154 | 3UTR |
| hsa-miR-4753-3p | NM_005656 | TMPRSS2 | 0.846153846154 | 3UTR |
| hsa-miR-4764-3p | NM_005656 | TMPRSS2 | 0.846153846154 | 3UTR |
| hsa-miR-4768-5p | NM_005656 | TMPRSS2 | 0.846153846154 | 3UTR |
| hsa-miR-4769-5p | NM_005656 | TMPRSS2 | 0.846153846154 | 3UTR |
| hsa-miR-4774-5p | NM_005656 | TMPRSS2 | 0.846153846154 | 3UTR |
| hsa-miR-4774-3p | NM_005656 | TMPRSS2 | 0.846153846154 | 3UTR |
| hsa-miR-4777-3p | NM_005656 | TMPRSS2 | 0.846153846154 | 3UTR |
| hsa-miR-4777-3p | NM_005656 | TMPRSS2 | 0.846153846154 | 3UTR |
| hsa-miR-4780 | NM_005656 | TMPRSS2 | 0.846153846154 | 3UTR |
| hsa-miR-4436b-3p | NM_005656 | TMPRSS2 | 0.846153846154 | 3UTR |
| hsa-miR-4781-5p | NM_005656 | TMPRSS2 | 0.846153846154 | 3UTR |
| hsa-miR-4786-5p | NM_005656 | TMPRSS2 | 0.846153846154 | 3UTR |
| hsa-miR-4793-5p | NM_005656 | TMPRSS2 | 0.846153846154 | 3UTR |

| | | | | |
|---|---|---|---|---|
| hsa-miR-4794 | NM_005656 | TMPRSS2 | 0.846153846154 | 3UTR |
| hsa-miR-4796-5p | NM_005656 | TMPRSS2 | 0.846153846154 | 3UTR |
| hsa-miR-4796-3p | NM_005656 | TMPRSS2 | 0.846153846154 | 3UTR |
| hsa-miR-4797-5p | NM_005656 | TMPRSS2 | 0.846153846154 | 3UTR |
| hsa-miR-4802-5p | NM_005656 | TMPRSS2 | 0.846153846154 | 3UTR |
| hsa-miR-4802-3p | NM_005656 | TMPRSS2 | 0.846153846154 | 3UTR |
| hsa-miR-5006-3p | NM_005656 | TMPRSS2 | 0.846153846154 | 3UTR |
| hsa-miR-5010-3p | NM_005656 | TMPRSS2 | 0.846153846154 | 3UTR |
| hsa-miR-5087 | NM_005656 | TMPRSS2 | 0.846153846154 | 3UTR |
| hsa-miR-5093 | NM_005656 | TMPRSS2 | 0.846153846154 | 3UTR |
| hsa-miR-5194 | NM_005656 | TMPRSS2 | 0.846153846154 | 3UTR |
| hsa-miR-5196-5p | NM_005656 | TMPRSS2 | 0.846153846154 | 3UTR |
| hsa-miR-5196-3p | NM_005656 | TMPRSS2 | 0.846153846154 | 3UTR |
| hsa-miR-5571-5p | NM_005656 | TMPRSS2 | 0.846153846154 | 3UTR |
| hsa-miR-5571-5p | NM_005656 | TMPRSS2 | 0.846153846154 | 3UTR |
| hsa-miR-5580-3p | NM_005656 | TMPRSS2 | 0.846153846154 | 3UTR |
| hsa-miR-5581-3p | NM_005656 | TMPRSS2 | 0.846153846154 | 3UTR |
| hsa-miR-548au-3p | NM_005656 | TMPRSS2 | 0.846153846154 | 3UTR |
| hsa-miR-1295b-5p | NM_005656 | TMPRSS2 | 0.846153846154 | 3UTR |
| hsa-miR-5591-5p | NM_005656 | TMPRSS2 | 0.846153846154 | 3UTR |
| hsa-miR-5682 | NM_005656 | TMPRSS2 | 0.846153846154 | 3UTR |
| hsa-miR-5685 | NM_005656 | TMPRSS2 | 0.846153846154 | 3UTR |
| hsa-miR-5691 | NM_005656 | TMPRSS2 | 0.846153846154 | 3UTR |
| hsa-miR-5739 | NM_005656 | TMPRSS2 | 0.846153846154 | 3UTR |
| hsa-miR-6070 | NM_005656 | TMPRSS2 | 0.846153846154 | 3UTR |
| hsa-miR-6072 | NM_005656 | TMPRSS2 | 0.846153846154 | 3UTR |
| hsa-miR-6076 | NM_005656 | TMPRSS2 | 0.846153846154 | 3UTR |
| hsa-miR-6085 | NM_005656 | TMPRSS2 | 0.846153846154 | 3UTR |
| hsa-miR-6086 | NM_005656 | TMPRSS2 | 0.846153846154 | 3UTR |
| hsa-miR-6124 | NM_005656 | TMPRSS2 | 0.846153846154 | 3UTR |
| hsa-miR-6125 | NM_005656 | TMPRSS2 | 0.846153846154 | 3UTR |
| hsa-miR-6131 | NM_005656 | TMPRSS2 | 0.846153846154 | 3UTR |
| hsa-miR-6131 | NM_005656 | TMPRSS2 | 0.846153846154 | 3UTR |
| hsa-miR-6132 | NM_005656 | TMPRSS2 | 0.846153846154 | 3UTR |
| hsa-miR-6133 | NM_005656 | TMPRSS2 | 0.846153846154 | 3UTR |
| hsa-miR-6504-5p | NM_005656 | TMPRSS2 | 0.846153846154 | 3UTR |
| hsa-miR-6508-3p | NM_005656 | TMPRSS2 | 0.846153846154 | 3UTR |
| hsa-miR-6509-3p | NM_005656 | TMPRSS2 | 0.846153846154 | 3UTR |
| hsa-miR-6511a-5p | NM_005656 | TMPRSS2 | 0.846153846154 | 3UTR |
| hsa-miR-6515-5p | NM_005656 | TMPRSS2 | 0.846153846154 | 3UTR |
| hsa-miR-6515-3p | NM_005656 | TMPRSS2 | 0.846153846154 | 3UTR |
| hsa-miR-6715a-3p | NM_005656 | TMPRSS2 | 0.846153846154 | 3UTR |
| hsa-miR-6717-5p | NM_005656 | TMPRSS2 | 0.846153846154 | 3UTR |
| hsa-miR-6719-3p | NM_005656 | TMPRSS2 | 0.846153846154 | 3UTR |
| hsa-miR-6720-5p | NM_005656 | TMPRSS2 | 0.846153846154 | 3UTR |
| hsa-miR-892c-5p | NM_005656 | TMPRSS2 | 0.846153846154 | 3UTR |
| hsa-miR-6727-3p | NM_005656 | TMPRSS2 | 0.846153846154 | 3UTR |

| | | | | |
|---|---|---|---|---|
| hsa-miR-6728-5p | NM_005656 | TMPRSS2 | 0.846153846154 | 3UTR |
| hsa-miR-6729-5p | NM_005656 | TMPRSS2 | 0.846153846154 | 3UTR |
| hsa-miR-6730-5p | NM_005656 | TMPRSS2 | 0.846153846154 | 3UTR |
| hsa-miR-6732-5p | NM_005656 | TMPRSS2 | 0.846153846154 | 3UTR |
| hsa-miR-6734-5p | NM_005656 | TMPRSS2 | 0.846153846154 | 3UTR |
| hsa-miR-6735-5p | NM_005656 | TMPRSS2 | 0.846153846154 | 3UTR |
| hsa-miR-6737-5p | NM_005656 | TMPRSS2 | 0.846153846154 | 3UTR |
| hsa-miR-6739-3p | NM_005656 | TMPRSS2 | 0.846153846154 | 3UTR |
| hsa-miR-6742-3p | NM_005656 | TMPRSS2 | 0.846153846154 | 3UTR |
| hsa-miR-6743-3p | NM_005656 | TMPRSS2 | 0.846153846154 | 3UTR |
| hsa-miR-6746-5p | NM_005656 | TMPRSS2 | 0.846153846154 | 3UTR |
| hsa-miR-6746-3p | NM_005656 | TMPRSS2 | 0.846153846154 | 3UTR |
| hsa-miR-6747-5p | NM_005656 | TMPRSS2 | 0.846153846154 | 3UTR |
| hsa-miR-6747-5p | NM_005656 | TMPRSS2 | 0.846153846154 | 3UTR |
| hsa-miR-6756-3p | NM_005656 | TMPRSS2 | 0.846153846154 | 3UTR |
| hsa-miR-6757-5p | NM_005656 | TMPRSS2 | 0.846153846154 | 3UTR |
| hsa-miR-6758-3p | NM_005656 | TMPRSS2 | 0.846153846154 | 3UTR |
| hsa-miR-6759-5p | NM_005656 | TMPRSS2 | 0.846153846154 | 3UTR |
| hsa-miR-6760-3p | NM_005656 | TMPRSS2 | 0.846153846154 | 3UTR |
| hsa-miR-6765-3p | NM_005656 | TMPRSS2 | 0.846153846154 | 3UTR |
| hsa-miR-6771-3p | NM_005656 | TMPRSS2 | 0.846153846154 | 3UTR |
| hsa-miR-6772-5p | NM_005656 | TMPRSS2 | 0.846153846154 | 3UTR |
| hsa-miR-6772-3p | NM_005656 | TMPRSS2 | 0.846153846154 | 3UTR |
| hsa-miR-6773-5p | NM_005656 | TMPRSS2 | 0.846153846154 | 3UTR |
| hsa-miR-6773-5p | NM_005656 | TMPRSS2 | 0.846153846154 | 3UTR |
| hsa-miR-6777-5p | NM_005656 | TMPRSS2 | 0.846153846154 | 3UTR |
| hsa-miR-6779-5p | NM_005656 | TMPRSS2 | 0.846153846154 | 3UTR |
| hsa-miR-6779-3p | NM_005656 | TMPRSS2 | 0.846153846154 | 3UTR |
| hsa-miR-6781-3p | NM_005656 | TMPRSS2 | 0.846153846154 | 3UTR |
| hsa-miR-6787-5p | NM_005656 | TMPRSS2 | 0.846153846154 | 3UTR |
| hsa-miR-6792-5p | NM_005656 | TMPRSS2 | 0.846153846154 | 3UTR |
| hsa-miR-6792-3p | NM_005656 | TMPRSS2 | 0.846153846154 | 3UTR |
| hsa-miR-6793-5p | NM_005656 | TMPRSS2 | 0.846153846154 | 3UTR |
| hsa-miR-6797-5p | NM_005656 | TMPRSS2 | 0.846153846154 | 3UTR |
| hsa-miR-6797-3p | NM_005656 | TMPRSS2 | 0.846153846154 | 3UTR |
| hsa-miR-6801-5p | NM_005656 | TMPRSS2 | 0.846153846154 | 3UTR |
| hsa-miR-6802-5p | NM_005656 | TMPRSS2 | 0.846153846154 | 3UTR |
| hsa-miR-6803-3p | NM_005656 | TMPRSS2 | 0.846153846154 | 3UTR |
| hsa-miR-6809-5p | NM_005656 | TMPRSS2 | 0.846153846154 | 3UTR |
| hsa-miR-6810-5p | NM_005656 | TMPRSS2 | 0.846153846154 | 3UTR |
| hsa-miR-6812-5p | NM_005656 | TMPRSS2 | 0.846153846154 | 3UTR |
| hsa-miR-6815-5p | NM_005656 | TMPRSS2 | 0.846153846154 | 3UTR |
| hsa-miR-6816-5p | NM_005656 | TMPRSS2 | 0.846153846154 | 3UTR |
| hsa-miR-6819-5p | NM_005656 | TMPRSS2 | 0.846153846154 | 3UTR |
| hsa-miR-6820-3p | NM_005656 | TMPRSS2 | 0.846153846154 | 3UTR |
| hsa-miR-6823-5p | NM_005656 | TMPRSS2 | 0.846153846154 | 3UTR |
| hsa-miR-6824-5p | NM_005656 | TMPRSS2 | 0.846153846154 | 3UTR |

| | | | | |
|---|---|---|---|---|
| hsa-miR-6825-5p | NM_005656 | TMPRSS2 | 0.846153846154 | 3UTR |
| hsa-miR-6831-5p | NM_005656 | TMPRSS2 | 0.846153846154 | 3UTR |
| hsa-miR-6832-5p | NM_005656 | TMPRSS2 | 0.846153846154 | 3UTR |
| hsa-miR-6833-3p | NM_005656 | TMPRSS2 | 0.846153846154 | 3UTR |
| hsa-miR-6834-5p | NM_005656 | TMPRSS2 | 0.846153846154 | 3UTR |
| hsa-miR-6835-5p | NM_005656 | TMPRSS2 | 0.846153846154 | 3UTR |
| hsa-miR-6835-3p | NM_005656 | TMPRSS2 | 0.846153846154 | 3UTR |
| hsa-miR-6780b-5p | NM_005656 | TMPRSS2 | 0.846153846154 | 3UTR |
| hsa-miR-6780b-5p | NM_005656 | TMPRSS2 | 0.846153846154 | 3UTR |
| hsa-miR-6836-3p | NM_005656 | TMPRSS2 | 0.846153846154 | 3UTR |
| hsa-miR-6845-5p | NM_005656 | TMPRSS2 | 0.846153846154 | 3UTR |
| hsa-miR-6845-3p | NM_005656 | TMPRSS2 | 0.846153846154 | 3UTR |
| hsa-miR-6851-5p | NM_005656 | TMPRSS2 | 0.846153846154 | 3UTR |
| hsa-miR-6853-5p | NM_005656 | TMPRSS2 | 0.846153846154 | 3UTR |
| hsa-miR-6853-3p | NM_005656 | TMPRSS2 | 0.846153846154 | 3UTR |
| hsa-miR-6856-3p | NM_005656 | TMPRSS2 | 0.846153846154 | 3UTR |
| hsa-miR-6858-5p | NM_005656 | TMPRSS2 | 0.846153846154 | 3UTR |
| hsa-miR-6859-5p | NM_005656 | TMPRSS2 | 0.846153846154 | 3UTR |
| hsa-miR-6769b-3p | NM_005656 | TMPRSS2 | 0.846153846154 | 3UTR |
| hsa-miR-6865-5p | NM_005656 | TMPRSS2 | 0.846153846154 | 3UTR |
| hsa-miR-6871-5p | NM_005656 | TMPRSS2 | 0.846153846154 | 3UTR |
| hsa-miR-6872-5p | NM_005656 | TMPRSS2 | 0.846153846154 | 3UTR |
| hsa-miR-6877-5p | NM_005656 | TMPRSS2 | 0.846153846154 | 3UTR |
| hsa-miR-6886-5p | NM_005656 | TMPRSS2 | 0.846153846154 | 3UTR |
| hsa-miR-6886-3p | NM_005656 | TMPRSS2 | 0.846153846154 | 3UTR |
| hsa-miR-6888-3p | NM_005656 | TMPRSS2 | 0.846153846154 | 3UTR |
| hsa-miR-7107-5p | NM_005656 | TMPRSS2 | 0.846153846154 | 3UTR |
| hsa-miR-7109-5p | NM_005656 | TMPRSS2 | 0.846153846154 | 3UTR |
| hsa-miR-7150 | NM_005656 | TMPRSS2 | 0.846153846154 | 3UTR |
| hsa-miR-7152-3p | NM_005656 | TMPRSS2 | 0.846153846154 | 3UTR |
| hsa-miR-7153-3p | NM_005656 | TMPRSS2 | 0.846153846154 | 3UTR |
| hsa-miR-7154-5p | NM_005656 | TMPRSS2 | 0.846153846154 | 3UTR |
| hsa-miR-7157-5p | NM_005656 | TMPRSS2 | 0.846153846154 | 3UTR |
| hsa-miR-7160-5p | NM_005656 | TMPRSS2 | 0.846153846154 | 3UTR |
| hsa-miR-7160-3p | NM_005656 | TMPRSS2 | 0.846153846154 | 3UTR |
| hsa-miR-7702 | NM_005656 | TMPRSS2 | 0.846153846154 | 3UTR |
| hsa-miR-7703 | NM_005656 | TMPRSS2 | 0.846153846154 | 3UTR |
| hsa-miR-7845-5p | NM_005656 | TMPRSS2 | 0.846153846154 | 3UTR |
| hsa-miR-7845-5p | NM_005656 | TMPRSS2 | 0.846153846154 | 3UTR |
| hsa-miR-7846-3p | NM_005656 | TMPRSS2 | 0.846153846154 | 3UTR |
| hsa-miR-7856-5p | NM_005656 | TMPRSS2 | 0.846153846154 | 3UTR |
| hsa-miR-8063 | NM_005656 | TMPRSS2 | 0.846153846154 | 3UTR |
| hsa-miR-8065 | NM_005656 | TMPRSS2 | 0.846153846154 | 3UTR |
| hsa-miR-8078 | NM_005656 | TMPRSS2 | 0.846153846154 | 3UTR |
| hsa-miR-8085 | NM_005656 | TMPRSS2 | 0.846153846154 | 3UTR |
| hsa-miR-8087 | NM_005656 | TMPRSS2 | 0.846153846154 | 3UTR |
| hsa-miR-9500 | NM_005656 | TMPRSS2 | 0.846153846154 | 3UTR |

| | | | | |
|---|---|---|---|---|
| hsa-miR-9903 | NM_005656 | TMPRSS2 | 0.846153846154 | 3UTR |
| hsa-miR-10226 | NM_005656 | TMPRSS2 | 0.846153846154 | 3UTR |
| hsa-miR-10394-5p | NM_005656 | TMPRSS2 | 0.846153846154 | 3UTR |
| hsa-miR-10395-5p | NM_005656 | TMPRSS2 | 0.846153846154 | 3UTR |
| hsa-miR-10399-3p | NM_005656 | TMPRSS2 | 0.846153846154 | 3UTR |
| hsa-miR-11399 | NM_005656 | TMPRSS2 | 0.846153846154 | 3UTR |
| hsa-miR-3085-5p | NM_005656 | TMPRSS2 | 0.846153846154 | 3UTR |
| hsa-miR-12114 | NM_005656 | TMPRSS2 | 0.846153846154 | 3UTR |
| hsa-miR-12122 | NM_005656 | TMPRSS2 | 0.846153846154 | 3UTR |
| hsa-miR-12122 | NM_005656 | TMPRSS2 | 0.846153846154 | 3UTR |
| hsa-miR-12124 | NM_005656 | TMPRSS2 | 0.846153846154 | 3UTR |
| hsa-miR-12127 | NM_005656 | TMPRSS2 | 0.846153846154 | 3UTR |
| hsa-miR-12128 | NM_005656 | TMPRSS2 | 0.846153846154 | 3UTR |
| hsa-miR-16-1-3p | NM_001135099 | TMPRSS2 | 0.846153846154 | 3UTR |
| hsa-miR-17-5p | NM_001135099 | TMPRSS2 | 0.846153846154 | 3UTR |
| hsa-miR-24-3p | NM_001135099 | TMPRSS2 | 0.846153846154 | 3UTR |
| hsa-miR-25-5p | NM_001135099 | TMPRSS2 | 0.846153846154 | 3UTR |
| hsa-miR-28-5p | NM_001135099 | TMPRSS2 | 0.846153846154 | 3UTR |
| hsa-miR-28-3p | NM_001135099 | TMPRSS2 | 0.846153846154 | 3UTR |
| hsa-miR-99a-3p | NM_001135099 | TMPRSS2 | 0.846153846154 | 3UTR |
| hsa-miR-103a-1-5p | NM_001135099 | TMPRSS2 | 0.846153846154 | 3UTR |
| hsa-miR-105-3p | NM_001135099 | TMPRSS2 | 0.846153846154 | 3UTR |
| hsa-miR-106a-5p | NM_001135099 | TMPRSS2 | 0.846153846154 | 3UTR |
| hsa-miR-148a-3p | NM_001135099 | TMPRSS2 | 0.846153846154 | 3UTR |
| hsa-miR-148a-3p | NM_001135099 | TMPRSS2 | 0.846153846154 | 3UTR |
| hsa-miR-30c-2-3p | NM_001135099 | TMPRSS2 | 0.846153846154 | 3UTR |
| hsa-miR-30c-2-3p | NM_001135099 | TMPRSS2 | 0.846153846154 | 3UTR |
| hsa-miR-139-5p | NM_001135099 | TMPRSS2 | 0.846153846154 | 3UTR |
| hsa-miR-139-5p | NM_001135099 | TMPRSS2 | 0.846153846154 | 3UTR |
| hsa-miR-7-1-3p | NM_001135099 | TMPRSS2 | 0.846153846154 | 3UTR |
| hsa-miR-34a-5p | NM_001135099 | TMPRSS2 | 0.846153846154 | 3UTR |
| hsa-miR-34a-5p | NM_001135099 | TMPRSS2 | 0.846153846154 | 3UTR |
| hsa-miR-181a-5p | NM_001135099 | TMPRSS2 | 0.846153846154 | 3UTR |
| hsa-miR-181c-3p | NM_001135099 | TMPRSS2 | 0.846153846154 | 3UTR |
| hsa-miR-183-3p | NM_001135099 | TMPRSS2 | 0.846153846154 | 3UTR |
| hsa-miR-210-5p | NM_001135099 | TMPRSS2 | 0.846153846154 | 3UTR |
| hsa-miR-212-5p | NM_001135099 | TMPRSS2 | 0.846153846154 | 3UTR |
| hsa-miR-214-3p | NM_001135099 | TMPRSS2 | 0.846153846154 | 3UTR |
| hsa-miR-217-5p | NM_001135099 | TMPRSS2 | 0.846153846154 | 3UTR |
| hsa-miR-219a-1-3p | NM_001135099 | TMPRSS2 | 0.846153846154 | 3UTR |
| hsa-miR-222-3p | NM_001135099 | TMPRSS2 | 0.846153846154 | 3UTR |
| hsa-miR-224-3p | NM_001135099 | TMPRSS2 | 0.846153846154 | 3UTR |
| hsa-miR-224-3p | NM_001135099 | TMPRSS2 | 0.846153846154 | 3UTR |
| hsa-let-7g-3p | NM_001135099 | TMPRSS2 | 0.846153846154 | 3UTR |
| hsa-miR-23b-5p | NM_001135099 | TMPRSS2 | 0.846153846154 | 3UTR |
| hsa-miR-27b-3p | NM_001135099 | TMPRSS2 | 0.846153846154 | 3UTR |
| hsa-miR-122-3p | NM_001135099 | TMPRSS2 | 0.846153846154 | 3UTR |

| | | | | |
|---|---|---|---|---|
| hsa-miR-122-3p | NM_001135099 | TMPRSS2 | 0.846153846154 | 3UTR |
| hsa-miR-128-3p | NM_001135099 | TMPRSS2 | 0.846153846154 | 3UTR |
| hsa-miR-130a-5p | NM_001135099 | TMPRSS2 | 0.846153846154 | 3UTR |
| hsa-miR-135a-2-3p | NM_001135099 | TMPRSS2 | 0.846153846154 | 3UTR |
| hsa-miR-145-5p | NM_001135099 | TMPRSS2 | 0.846153846154 | 3UTR |
| hsa-miR-9-3p | NM_001135099 | TMPRSS2 | 0.846153846154 | 3UTR |
| hsa-miR-125a-5p | NM_001135099 | TMPRSS2 | 0.846153846154 | 3UTR |
| hsa-miR-138-1-3p | NM_001135099 | TMPRSS2 | 0.846153846154 | 3UTR |
| hsa-miR-150-5p | NM_001135099 | TMPRSS2 | 0.846153846154 | 3UTR |
| hsa-miR-185-5p | NM_001135099 | TMPRSS2 | 0.846153846154 | 3UTR |
| hsa-miR-185-3p | NM_001135099 | TMPRSS2 | 0.846153846154 | 3UTR |
| hsa-miR-188-5p | NM_001135099 | TMPRSS2 | 0.846153846154 | 3UTR |
| hsa-miR-320a-5p | NM_001135099 | TMPRSS2 | 0.846153846154 | 3UTR |
| hsa-miR-155-5p | NM_001135099 | TMPRSS2 | 0.846153846154 | 3UTR |
| hsa-miR-128-2-5p | NM_001135099 | TMPRSS2 | 0.846153846154 | 3UTR |
| hsa-miR-29c-5p | NM_001135099 | TMPRSS2 | 0.846153846154 | 3UTR |
| hsa-miR-299-3p | NM_001135099 | TMPRSS2 | 0.846153846154 | 3UTR |
| hsa-miR-130b-5p | NM_001135099 | TMPRSS2 | 0.846153846154 | 3UTR |
| hsa-miR-130b-5p | NM_001135099 | TMPRSS2 | 0.846153846154 | 3UTR |
| hsa-miR-365a-5p | NM_001135099 | TMPRSS2 | 0.846153846154 | 3UTR |
| hsa-miR-365b-5p | NM_001135099 | TMPRSS2 | 0.846153846154 | 3UTR |
| hsa-miR-370-3p | NM_001135099 | TMPRSS2 | 0.846153846154 | 3UTR |
| hsa-miR-371a-3p | NM_001135099 | TMPRSS2 | 0.846153846154 | 3UTR |
| hsa-miR-373-5p | NM_001135099 | TMPRSS2 | 0.846153846154 | 3UTR |
| hsa-miR-373-5p | NM_001135099 | TMPRSS2 | 0.846153846154 | 3UTR |
| hsa-miR-373-3p | NM_001135099 | TMPRSS2 | 0.846153846154 | 3UTR |
| hsa-miR-376a-5p | NM_001135099 | TMPRSS2 | 0.846153846154 | 3UTR |
| hsa-miR-378a-3p | NM_001135099 | TMPRSS2 | 0.846153846154 | 3UTR |
| hsa-miR-330-3p | NM_001135099 | TMPRSS2 | 0.846153846154 | 3UTR |
| hsa-miR-326 | NM_001135099 | TMPRSS2 | 0.846153846154 | 3UTR |
| hsa-miR-324-3p | NM_001135099 | TMPRSS2 | 0.846153846154 | 3UTR |
| hsa-miR-345-5p | NM_001135099 | TMPRSS2 | 0.846153846154 | 3UTR |
| hsa-miR-196b-3p | NM_001135099 | TMPRSS2 | 0.846153846154 | 3UTR |
| hsa-miR-425-5p | NM_001135099 | TMPRSS2 | 0.846153846154 | 3UTR |
| hsa-miR-20b-5p | NM_001135099 | TMPRSS2 | 0.846153846154 | 3UTR |
| hsa-miR-20b-5p | NM_001135099 | TMPRSS2 | 0.846153846154 | 3UTR |
| hsa-miR-20b-5p | NM_001135099 | TMPRSS2 | 0.846153846154 | 3UTR |
| hsa-miR-329-5p | NM_001135099 | TMPRSS2 | 0.846153846154 | 3UTR |
| hsa-miR-409-5p | NM_001135099 | TMPRSS2 | 0.846153846154 | 3UTR |
| hsa-miR-412-3p | NM_001135099 | TMPRSS2 | 0.846153846154 | 3UTR |
| hsa-miR-483-5p | NM_001135099 | TMPRSS2 | 0.846153846154 | 3UTR |
| hsa-miR-485-3p | NM_001135099 | TMPRSS2 | 0.846153846154 | 3UTR |
| hsa-miR-487a-3p | NM_001135099 | TMPRSS2 | 0.846153846154 | 3UTR |
| hsa-miR-490-3p | NM_001135099 | TMPRSS2 | 0.846153846154 | 3UTR |
| hsa-miR-491-3p | NM_001135099 | TMPRSS2 | 0.846153846154 | 3UTR |
| hsa-miR-432-5p | NM_001135099 | TMPRSS2 | 0.846153846154 | 3UTR |
| hsa-miR-432-3p | NM_001135099 | TMPRSS2 | 0.846153846154 | 3UTR |

| | | | | |
|---|---|---|---|---|
| hsa-miR-494-3p | NM_001135099 | TMPRSS2 | 0.846153846154 | 3UTR |
| hsa-miR-497-5p | NM_001135099 | TMPRSS2 | 0.846153846154 | 3UTR |
| hsa-miR-498-5p | NM_001135099 | TMPRSS2 | 0.846153846154 | 3UTR |
| hsa-miR-498-5p | NM_001135099 | TMPRSS2 | 0.846153846154 | 3UTR |
| hsa-miR-520e-3p | NM_001135099 | TMPRSS2 | 0.846153846154 | 3UTR |
| hsa-miR-515-5p | NM_001135099 | TMPRSS2 | 0.846153846154 | 3UTR |
| hsa-miR-520f-5p | NM_001135099 | TMPRSS2 | 0.846153846154 | 3UTR |
| hsa-miR-525-3p | NM_001135099 | TMPRSS2 | 0.846153846154 | 3UTR |
| hsa-miR-523-3p | NM_001135099 | TMPRSS2 | 0.846153846154 | 3UTR |
| hsa-miR-523-3p | NM_001135099 | TMPRSS2 | 0.846153846154 | 3UTR |
| hsa-miR-520b-3p | NM_001135099 | TMPRSS2 | 0.846153846154 | 3UTR |
| hsa-miR-526a-5p | NM_001135099 | TMPRSS2 | 0.846153846154 | 3UTR |
| hsa-miR-520c-5p | NM_001135099 | TMPRSS2 | 0.846153846154 | 3UTR |
| hsa-miR-518c-3p | NM_001135099 | TMPRSS2 | 0.846153846154 | 3UTR |
| hsa-miR-516b-3p | NM_001135099 | TMPRSS2 | 0.846153846154 | 3UTR |
| hsa-miR-518a-5p | NM_001135099 | TMPRSS2 | 0.846153846154 | 3UTR |
| hsa-miR-518d-5p | NM_001135099 | TMPRSS2 | 0.846153846154 | 3UTR |
| hsa-miR-520h | NM_001135099 | TMPRSS2 | 0.846153846154 | 3UTR |
| hsa-miR-522-3p | NM_001135099 | TMPRSS2 | 0.846153846154 | 3UTR |
| hsa-miR-527 | NM_001135099 | TMPRSS2 | 0.846153846154 | 3UTR |
| hsa-miR-516a-3p | NM_001135099 | TMPRSS2 | 0.846153846154 | 3UTR |
| hsa-miR-499a-3p | NM_001135099 | TMPRSS2 | 0.846153846154 | 3UTR |
| hsa-miR-503-5p | NM_001135099 | TMPRSS2 | 0.846153846154 | 3UTR |
| hsa-miR-503-5p | NM_001135099 | TMPRSS2 | 0.846153846154 | 3UTR |
| hsa-miR-513a-5p | NM_001135099 | TMPRSS2 | 0.846153846154 | 3UTR |
| hsa-miR-532-5p | NM_001135099 | TMPRSS2 | 0.846153846154 | 3UTR |
| hsa-miR-551a | NM_001135099 | TMPRSS2 | 0.846153846154 | 3UTR |
| hsa-miR-552-5p | NM_001135099 | TMPRSS2 | 0.846153846154 | 3UTR |
| hsa-miR-554 | NM_001135099 | TMPRSS2 | 0.846153846154 | 3UTR |
| hsa-miR-558 | NM_001135099 | TMPRSS2 | 0.846153846154 | 3UTR |
| hsa-miR-564 | NM_001135099 | TMPRSS2 | 0.846153846154 | 3UTR |
| hsa-miR-571 | NM_001135099 | TMPRSS2 | 0.846153846154 | 3UTR |
| hsa-miR-584-5p | NM_001135099 | TMPRSS2 | 0.846153846154 | 3UTR |
| hsa-miR-587 | NM_001135099 | TMPRSS2 | 0.846153846154 | 3UTR |
| hsa-miR-550a-3p | NM_001135099 | TMPRSS2 | 0.846153846154 | 3UTR |
| hsa-miR-601 | NM_001135099 | TMPRSS2 | 0.846153846154 | 3UTR |
| hsa-miR-608 | NM_001135099 | TMPRSS2 | 0.846153846154 | 3UTR |
| hsa-miR-609 | NM_001135099 | TMPRSS2 | 0.846153846154 | 3UTR |
| hsa-miR-610 | NM_001135099 | TMPRSS2 | 0.846153846154 | 3UTR |
| hsa-miR-612 | NM_001135099 | TMPRSS2 | 0.846153846154 | 3UTR |
| hsa-miR-615-3p | NM_001135099 | TMPRSS2 | 0.846153846154 | 3UTR |
| hsa-miR-616-3p | NM_001135099 | TMPRSS2 | 0.846153846154 | 3UTR |
| hsa-miR-627-5p | NM_001135099 | TMPRSS2 | 0.846153846154 | 3UTR |
| hsa-miR-635 | NM_001135099 | TMPRSS2 | 0.846153846154 | 3UTR |
| hsa-miR-643 | NM_001135099 | TMPRSS2 | 0.846153846154 | 3UTR |
| hsa-miR-644a | NM_001135099 | TMPRSS2 | 0.846153846154 | 3UTR |
| hsa-miR-645 | NM_001135099 | TMPRSS2 | 0.846153846154 | 3UTR |

| | | | | |
|---|---|---|---|---|
| hsa-miR-646 | NM_001135099 | TMPRSS2 | 0.846153846154 | 3UTR |
| hsa-miR-646 | NM_001135099 | TMPRSS2 | 0.846153846154 | 3UTR |
| hsa-miR-651-5p | NM_001135099 | TMPRSS2 | 0.846153846154 | 3UTR |
| hsa-miR-449b-3p | NM_001135099 | TMPRSS2 | 0.846153846154 | 3UTR |
| hsa-miR-449b-3p | NM_001135099 | TMPRSS2 | 0.846153846154 | 3UTR |
| hsa-miR-657 | NM_001135099 | TMPRSS2 | 0.846153846154 | 3UTR |
| hsa-miR-658 | NM_001135099 | TMPRSS2 | 0.846153846154 | 3UTR |
| hsa-miR-542-5p | NM_001135099 | TMPRSS2 | 0.846153846154 | 3UTR |
| hsa-miR-766-5p | NM_001135099 | TMPRSS2 | 0.846153846154 | 3UTR |
| hsa-miR-670-5p | NM_001135099 | TMPRSS2 | 0.846153846154 | 3UTR |
| hsa-miR-764 | NM_001135099 | TMPRSS2 | 0.846153846154 | 3UTR |
| hsa-miR-765 | NM_001135099 | TMPRSS2 | 0.846153846154 | 3UTR |
| hsa-miR-675-5p | NM_001135099 | TMPRSS2 | 0.846153846154 | 3UTR |
| hsa-miR-450b-3p | NM_001135099 | TMPRSS2 | 0.846153846154 | 3UTR |
| hsa-miR-874-3p | NM_001135099 | TMPRSS2 | 0.846153846154 | 3UTR |
| hsa-miR-890 | NM_001135099 | TMPRSS2 | 0.846153846154 | 3UTR |
| hsa-miR-541-5p | NM_001135099 | TMPRSS2 | 0.846153846154 | 3UTR |
| hsa-miR-875-5p | NM_001135099 | TMPRSS2 | 0.846153846154 | 3UTR |
| hsa-miR-147b-3p | NM_001135099 | TMPRSS2 | 0.846153846154 | 3UTR |
| hsa-miR-877-5p | NM_001135099 | TMPRSS2 | 0.846153846154 | 3UTR |
| hsa-miR-887-5p | NM_001135099 | TMPRSS2 | 0.846153846154 | 3UTR |
| hsa-miR-665 | NM_001135099 | TMPRSS2 | 0.846153846154 | 3UTR |
| hsa-miR-301b-5p | NM_001135099 | TMPRSS2 | 0.846153846154 | 3UTR |
| hsa-miR-216b-5p | NM_001135099 | TMPRSS2 | 0.846153846154 | 3UTR |
| hsa-miR-920 | NM_001135099 | TMPRSS2 | 0.846153846154 | 3UTR |
| hsa-miR-922 | NM_001135099 | TMPRSS2 | 0.846153846154 | 3UTR |
| hsa-miR-922 | NM_001135099 | TMPRSS2 | 0.846153846154 | 3UTR |
| hsa-miR-934 | NM_001135099 | TMPRSS2 | 0.846153846154 | 3UTR |
| hsa-miR-937-3p | NM_001135099 | TMPRSS2 | 0.846153846154 | 3UTR |
| hsa-miR-943 | NM_001135099 | TMPRSS2 | 0.846153846154 | 3UTR |
| hsa-miR-1178-5p | NM_001135099 | TMPRSS2 | 0.846153846154 | 3UTR |
| hsa-miR-1182 | NM_001135099 | TMPRSS2 | 0.846153846154 | 3UTR |
| hsa-miR-1231 | NM_001135099 | TMPRSS2 | 0.846153846154 | 3UTR |
| hsa-miR-1203 | NM_001135099 | TMPRSS2 | 0.846153846154 | 3UTR |
| hsa-miR-1286 | NM_001135099 | TMPRSS2 | 0.846153846154 | 3UTR |
| hsa-miR-548k | NM_001135099 | TMPRSS2 | 0.846153846154 | 3UTR |
| hsa-miR-1249-5p | NM_001135099 | TMPRSS2 | 0.846153846154 | 3UTR |
| hsa-miR-1249-5p | NM_001135099 | TMPRSS2 | 0.846153846154 | 3UTR |
| hsa-miR-1251-5p | NM_001135099 | TMPRSS2 | 0.846153846154 | 3UTR |
| hsa-miR-1253 | NM_001135099 | TMPRSS2 | 0.846153846154 | 3UTR |
| hsa-miR-1263 | NM_001135099 | TMPRSS2 | 0.846153846154 | 3UTR |
| hsa-miR-1270 | NM_001135099 | TMPRSS2 | 0.846153846154 | 3UTR |
| hsa-miR-1275 | NM_001135099 | TMPRSS2 | 0.846153846154 | 3UTR |
| hsa-miR-1281 | NM_001135099 | TMPRSS2 | 0.846153846154 | 3UTR |
| hsa-miR-513c-5p | NM_001135099 | TMPRSS2 | 0.846153846154 | 3UTR |
| hsa-miR-1321 | NM_001135099 | TMPRSS2 | 0.846153846154 | 3UTR |
| hsa-miR-320d | NM_001135099 | TMPRSS2 | 0.846153846154 | 3UTR |

| | | | | |
|---|---|---|---|---|
| hsa-miR-1910-5p | NM_001135099 | TMPRSS2 | 0.846153846154 | 3UTR |
| hsa-miR-1911-3p | NM_001135099 | TMPRSS2 | 0.846153846154 | 3UTR |
| hsa-miR-1913 | NM_001135099 | TMPRSS2 | 0.846153846154 | 3UTR |
| hsa-miR-2114-5p | NM_001135099 | TMPRSS2 | 0.846153846154 | 3UTR |
| hsa-miR-2114-5p | NM_001135099 | TMPRSS2 | 0.846153846154 | 3UTR |
| hsa-miR-2117 | NM_001135099 | TMPRSS2 | 0.846153846154 | 3UTR |
| hsa-miR-2276-3p | NM_001135099 | TMPRSS2 | 0.846153846154 | 3UTR |
| hsa-miR-2278 | NM_001135099 | TMPRSS2 | 0.846153846154 | 3UTR |
| hsa-miR-2909 | NM_001135099 | TMPRSS2 | 0.846153846154 | 3UTR |
| hsa-miR-3116 | NM_001135099 | TMPRSS2 | 0.846153846154 | 3UTR |
| hsa-miR-548s | NM_001135099 | TMPRSS2 | 0.846153846154 | 3UTR |
| hsa-miR-3125 | NM_001135099 | TMPRSS2 | 0.846153846154 | 3UTR |
| hsa-miR-3130-3p | NM_001135099 | TMPRSS2 | 0.846153846154 | 3UTR |
| hsa-miR-3132 | NM_001135099 | TMPRSS2 | 0.846153846154 | 3UTR |
| hsa-miR-378b | NM_001135099 | TMPRSS2 | 0.846153846154 | 3UTR |
| hsa-miR-3139 | NM_001135099 | TMPRSS2 | 0.846153846154 | 3UTR |
| hsa-miR-3141 | NM_001135099 | TMPRSS2 | 0.846153846154 | 3UTR |
| hsa-miR-3141 | NM_001135099 | TMPRSS2 | 0.846153846154 | 3UTR |
| hsa-miR-1273c | NM_001135099 | TMPRSS2 | 0.846153846154 | 3UTR |
| hsa-miR-3150a-5p | NM_001135099 | TMPRSS2 | 0.846153846154 | 3UTR |
| hsa-miR-3157-3p | NM_001135099 | TMPRSS2 | 0.846153846154 | 3UTR |
| hsa-miR-3162-5p | NM_001135099 | TMPRSS2 | 0.846153846154 | 3UTR |
| hsa-miR-3162-5p | NM_001135099 | TMPRSS2 | 0.846153846154 | 3UTR |
| hsa-miR-3163 | NM_001135099 | TMPRSS2 | 0.846153846154 | 3UTR |
| hsa-miR-3164 | NM_001135099 | TMPRSS2 | 0.846153846154 | 3UTR |
| hsa-miR-3170 | NM_001135099 | TMPRSS2 | 0.846153846154 | 3UTR |
| hsa-miR-1193 | NM_001135099 | TMPRSS2 | 0.846153846154 | 3UTR |
| hsa-miR-3174 | NM_001135099 | TMPRSS2 | 0.846153846154 | 3UTR |
| hsa-miR-3184-3p | NM_001135099 | TMPRSS2 | 0.846153846154 | 3UTR |
| hsa-miR-3189-5p | NM_001135099 | TMPRSS2 | 0.846153846154 | 3UTR |
| hsa-miR-3189-3p | NM_001135099 | TMPRSS2 | 0.846153846154 | 3UTR |
| hsa-miR-3192-3p | NM_001135099 | TMPRSS2 | 0.846153846154 | 3UTR |
| hsa-miR-3197 | NM_001135099 | TMPRSS2 | 0.846153846154 | 3UTR |
| hsa-miR-378c | NM_001135099 | TMPRSS2 | 0.846153846154 | 3UTR |
| hsa-miR-4294 | NM_001135099 | TMPRSS2 | 0.846153846154 | 3UTR |
| hsa-miR-4306 | NM_001135099 | TMPRSS2 | 0.846153846154 | 3UTR |
| hsa-miR-4313 | NM_001135099 | TMPRSS2 | 0.846153846154 | 3UTR |
| hsa-miR-4320 | NM_001135099 | TMPRSS2 | 0.846153846154 | 3UTR |
| hsa-miR-4320 | NM_001135099 | TMPRSS2 | 0.846153846154 | 3UTR |
| hsa-miR-4317 | NM_001135099 | TMPRSS2 | 0.846153846154 | 3UTR |
| hsa-miR-4259 | NM_001135099 | TMPRSS2 | 0.846153846154 | 3UTR |
| hsa-miR-4266 | NM_001135099 | TMPRSS2 | 0.846153846154 | 3UTR |
| hsa-miR-4269 | NM_001135099 | TMPRSS2 | 0.846153846154 | 3UTR |
| hsa-miR-4271 | NM_001135099 | TMPRSS2 | 0.846153846154 | 3UTR |
| hsa-miR-4271 | NM_001135099 | TMPRSS2 | 0.846153846154 | 3UTR |
| hsa-miR-4273 | NM_001135099 | TMPRSS2 | 0.846153846154 | 3UTR |
| hsa-miR-4279 | NM_001135099 | TMPRSS2 | 0.846153846154 | 3UTR |

| | | | | |
|---|---|---|---|---|
| hsa-miR-4278 | NM_001135099 | TMPRSS2 | 0.846153846154 | 3UTR |
| hsa-miR-4285 | NM_001135099 | TMPRSS2 | 0.846153846154 | 3UTR |
| hsa-miR-4289 | NM_001135099 | TMPRSS2 | 0.846153846154 | 3UTR |
| hsa-miR-4291 | NM_001135099 | TMPRSS2 | 0.846153846154 | 3UTR |
| hsa-miR-500b-5p | NM_001135099 | TMPRSS2 | 0.846153846154 | 3UTR |
| hsa-miR-500b-5p | NM_001135099 | TMPRSS2 | 0.846153846154 | 3UTR |
| hsa-miR-3610 | NM_001135099 | TMPRSS2 | 0.846153846154 | 3UTR |
| hsa-miR-3610 | NM_001135099 | TMPRSS2 | 0.846153846154 | 3UTR |
| hsa-miR-3614-3p | NM_001135099 | TMPRSS2 | 0.846153846154 | 3UTR |
| hsa-miR-3617-3p | NM_001135099 | TMPRSS2 | 0.846153846154 | 3UTR |
| hsa-miR-23c | NM_001135099 | TMPRSS2 | 0.846153846154 | 3UTR |
| hsa-miR-3650 | NM_001135099 | TMPRSS2 | 0.846153846154 | 3UTR |
| hsa-miR-3654 | NM_001135099 | TMPRSS2 | 0.846153846154 | 3UTR |
| hsa-miR-3664-5p | NM_001135099 | TMPRSS2 | 0.846153846154 | 3UTR |
| hsa-miR-3665 | NM_001135099 | TMPRSS2 | 0.846153846154 | 3UTR |
| hsa-miR-3675-3p | NM_001135099 | TMPRSS2 | 0.846153846154 | 3UTR |
| hsa-miR-3678-3p | NM_001135099 | TMPRSS2 | 0.846153846154 | 3UTR |
| hsa-miR-3680-5p | NM_001135099 | TMPRSS2 | 0.846153846154 | 3UTR |
| hsa-miR-3680-3p | NM_001135099 | TMPRSS2 | 0.846153846154 | 3UTR |
| hsa-miR-3689a-5p | NM_001135099 | TMPRSS2 | 0.846153846154 | 3UTR |
| hsa-miR-3689b-5p | NM_001135099 | TMPRSS2 | 0.846153846154 | 3UTR |
| hsa-miR-3689b-3p | NM_001135099 | TMPRSS2 | 0.846153846154 | 3UTR |
| hsa-miR-3916 | NM_001135099 | TMPRSS2 | 0.846153846154 | 3UTR |
| hsa-miR-3924 | NM_001135099 | TMPRSS2 | 0.846153846154 | 3UTR |
| hsa-miR-3925-5p | NM_001135099 | TMPRSS2 | 0.846153846154 | 3UTR |
| hsa-miR-3934-5p | NM_001135099 | TMPRSS2 | 0.846153846154 | 3UTR |
| hsa-miR-374c-3p | NM_001135099 | TMPRSS2 | 0.846153846154 | 3UTR |
| hsa-miR-550b-2-5p | NM_001135099 | TMPRSS2 | 0.846153846154 | 3UTR |
| hsa-miR-378e | NM_001135099 | TMPRSS2 | 0.846153846154 | 3UTR |
| hsa-miR-4421 | NM_001135099 | TMPRSS2 | 0.846153846154 | 3UTR |
| hsa-miR-4421 | NM_001135099 | TMPRSS2 | 0.846153846154 | 3UTR |
| hsa-miR-378g | NM_001135099 | TMPRSS2 | 0.846153846154 | 3UTR |
| hsa-miR-4424 | NM_001135099 | TMPRSS2 | 0.846153846154 | 3UTR |
| hsa-miR-4433a-5p | NM_001135099 | TMPRSS2 | 0.846153846154 | 3UTR |
| hsa-miR-4433a-5p | NM_001135099 | TMPRSS2 | 0.846153846154 | 3UTR |
| hsa-miR-4433a-3p | NM_001135099 | TMPRSS2 | 0.846153846154 | 3UTR |
| hsa-miR-4439 | NM_001135099 | TMPRSS2 | 0.846153846154 | 3UTR |
| hsa-miR-4448 | NM_001135099 | TMPRSS2 | 0.846153846154 | 3UTR |
| hsa-miR-378h | NM_001135099 | TMPRSS2 | 0.846153846154 | 3UTR |
| hsa-miR-4465 | NM_001135099 | TMPRSS2 | 0.846153846154 | 3UTR |
| hsa-miR-4472 | NM_001135099 | TMPRSS2 | 0.846153846154 | 3UTR |
| hsa-miR-4473 | NM_001135099 | TMPRSS2 | 0.846153846154 | 3UTR |
| hsa-miR-4474-3p | NM_001135099 | TMPRSS2 | 0.846153846154 | 3UTR |
| hsa-miR-4478 | NM_001135099 | TMPRSS2 | 0.846153846154 | 3UTR |
| hsa-miR-3689c | NM_001135099 | TMPRSS2 | 0.846153846154 | 3UTR |
| hsa-miR-3689e | NM_001135099 | TMPRSS2 | 0.846153846154 | 3UTR |
| hsa-miR-3155b | NM_001135099 | TMPRSS2 | 0.846153846154 | 3UTR |

| | | | | |
|---|---|---|---|---|
| hsa-miR-4485-5p | NM_001135099 | TMPRSS2 | 0.846153846154 | 3UTR |
| hsa-miR-4487 | NM_001135099 | TMPRSS2 | 0.846153846154 | 3UTR |
| hsa-miR-4487 | NM_001135099 | TMPRSS2 | 0.846153846154 | 3UTR |
| hsa-miR-4488 | NM_001135099 | TMPRSS2 | 0.846153846154 | 3UTR |
| hsa-miR-2392 | NM_001135099 | TMPRSS2 | 0.846153846154 | 3UTR |
| hsa-miR-4514 | NM_001135099 | TMPRSS2 | 0.846153846154 | 3UTR |
| hsa-miR-4520-5p | NM_001135099 | TMPRSS2 | 0.846153846154 | 3UTR |
| hsa-miR-4525 | NM_001135099 | TMPRSS2 | 0.846153846154 | 3UTR |
| hsa-miR-4533 | NM_001135099 | TMPRSS2 | 0.846153846154 | 3UTR |
| hsa-miR-1587 | NM_001135099 | TMPRSS2 | 0.846153846154 | 3UTR |
| hsa-miR-4540 | NM_001135099 | TMPRSS2 | 0.846153846154 | 3UTR |
| hsa-miR-3975 | NM_001135099 | TMPRSS2 | 0.846153846154 | 3UTR |
| hsa-miR-3976 | NM_001135099 | TMPRSS2 | 0.846153846154 | 3UTR |
| hsa-miR-4632-5p | NM_001135099 | TMPRSS2 | 0.846153846154 | 3UTR |
| hsa-miR-4633-5p | NM_001135099 | TMPRSS2 | 0.846153846154 | 3UTR |
| hsa-miR-4635 | NM_001135099 | TMPRSS2 | 0.846153846154 | 3UTR |
| hsa-miR-4640-5p | NM_001135099 | TMPRSS2 | 0.846153846154 | 3UTR |
| hsa-miR-4642 | NM_001135099 | TMPRSS2 | 0.846153846154 | 3UTR |
| hsa-miR-4644 | NM_001135099 | TMPRSS2 | 0.846153846154 | 3UTR |
| hsa-miR-4645-5p | NM_001135099 | TMPRSS2 | 0.846153846154 | 3UTR |
| hsa-miR-4647 | NM_001135099 | TMPRSS2 | 0.846153846154 | 3UTR |
| hsa-miR-4648 | NM_001135099 | TMPRSS2 | 0.846153846154 | 3UTR |
| hsa-miR-4669 | NM_001135099 | TMPRSS2 | 0.846153846154 | 3UTR |
| hsa-miR-4671-3p | NM_001135099 | TMPRSS2 | 0.846153846154 | 3UTR |
| hsa-miR-4681 | NM_001135099 | TMPRSS2 | 0.846153846154 | 3UTR |
| hsa-miR-4682 | NM_001135099 | TMPRSS2 | 0.846153846154 | 3UTR |
| hsa-miR-4685-5p | NM_001135099 | TMPRSS2 | 0.846153846154 | 3UTR |
| hsa-miR-4685-5p | NM_001135099 | TMPRSS2 | 0.846153846154 | 3UTR |
| hsa-miR-1343-3p | NM_001135099 | TMPRSS2 | 0.846153846154 | 3UTR |
| hsa-miR-4690-3p | NM_001135099 | TMPRSS2 | 0.846153846154 | 3UTR |
| hsa-miR-4691-5p | NM_001135099 | TMPRSS2 | 0.846153846154 | 3UTR |
| hsa-miR-4695-3p | NM_001135099 | TMPRSS2 | 0.846153846154 | 3UTR |
| hsa-miR-4696 | NM_001135099 | TMPRSS2 | 0.846153846154 | 3UTR |
| hsa-miR-4704-5p | NM_001135099 | TMPRSS2 | 0.846153846154 | 3UTR |
| hsa-miR-4704-5p | NM_001135099 | TMPRSS2 | 0.846153846154 | 3UTR |
| hsa-miR-4705 | NM_001135099 | TMPRSS2 | 0.846153846154 | 3UTR |
| hsa-miR-4706 | NM_001135099 | TMPRSS2 | 0.846153846154 | 3UTR |
| hsa-miR-4708-3p | NM_001135099 | TMPRSS2 | 0.846153846154 | 3UTR |
| hsa-miR-203b-5p | NM_001135099 | TMPRSS2 | 0.846153846154 | 3UTR |
| hsa-miR-4712-3p | NM_001135099 | TMPRSS2 | 0.846153846154 | 3UTR |
| hsa-miR-4713-5p | NM_001135099 | TMPRSS2 | 0.846153846154 | 3UTR |
| hsa-miR-4714-5p | NM_001135099 | TMPRSS2 | 0.846153846154 | 3UTR |
| hsa-miR-4716-3p | NM_001135099 | TMPRSS2 | 0.846153846154 | 3UTR |
| hsa-miR-3529-5p | NM_001135099 | TMPRSS2 | 0.846153846154 | 3UTR |
| hsa-miR-4721 | NM_001135099 | TMPRSS2 | 0.846153846154 | 3UTR |
| hsa-miR-4724-5p | NM_001135099 | TMPRSS2 | 0.846153846154 | 3UTR |
| hsa-miR-4731-5p | NM_001135099 | TMPRSS2 | 0.846153846154 | 3UTR |

| | | | | |
|---|---|---|---|---|
| hsa-miR-4732-5p | NM_001135099 | TMPRSS2 | 0.846153846154 | 3UTR |
| hsa-miR-4737 | NM_001135099 | TMPRSS2 | 0.846153846154 | 3UTR |
| hsa-miR-3064-3p | NM_001135099 | TMPRSS2 | 0.846153846154 | 3UTR |
| hsa-miR-4743-5p | NM_001135099 | TMPRSS2 | 0.846153846154 | 3UTR |
| hsa-miR-4747-5p | NM_001135099 | TMPRSS2 | 0.846153846154 | 3UTR |
| hsa-miR-4747-3p | NM_001135099 | TMPRSS2 | 0.846153846154 | 3UTR |
| hsa-miR-4748 | NM_001135099 | TMPRSS2 | 0.846153846154 | 3UTR |
| hsa-miR-4748 | NM_001135099 | TMPRSS2 | 0.846153846154 | 3UTR |
| hsa-miR-4750-3p | NM_001135099 | TMPRSS2 | 0.846153846154 | 3UTR |
| hsa-miR-4752 | NM_001135099 | TMPRSS2 | 0.846153846154 | 3UTR |
| hsa-miR-4753-3p | NM_001135099 | TMPRSS2 | 0.846153846154 | 3UTR |
| hsa-miR-4761-5p | NM_001135099 | TMPRSS2 | 0.846153846154 | 3UTR |
| hsa-miR-4768-5p | NM_001135099 | TMPRSS2 | 0.846153846154 | 3UTR |
| hsa-miR-4771 | NM_001135099 | TMPRSS2 | 0.846153846154 | 3UTR |
| hsa-miR-4775 | NM_001135099 | TMPRSS2 | 0.846153846154 | 3UTR |
| hsa-miR-4779 | NM_001135099 | TMPRSS2 | 0.846153846154 | 3UTR |
| hsa-miR-4780 | NM_001135099 | TMPRSS2 | 0.846153846154 | 3UTR |
| hsa-miR-4436b-3p | NM_001135099 | TMPRSS2 | 0.846153846154 | 3UTR |
| hsa-miR-4785 | NM_001135099 | TMPRSS2 | 0.846153846154 | 3UTR |
| hsa-miR-4785 | NM_001135099 | TMPRSS2 | 0.846153846154 | 3UTR |
| hsa-miR-1245b-5p | NM_001135099 | TMPRSS2 | 0.846153846154 | 3UTR |
| hsa-miR-2467-3p | NM_001135099 | TMPRSS2 | 0.846153846154 | 3UTR |
| hsa-miR-4786-5p | NM_001135099 | TMPRSS2 | 0.846153846154 | 3UTR |
| hsa-miR-4790-5p | NM_001135099 | TMPRSS2 | 0.846153846154 | 3UTR |
| hsa-miR-4793-5p | NM_001135099 | TMPRSS2 | 0.846153846154 | 3UTR |
| hsa-miR-4794 | NM_001135099 | TMPRSS2 | 0.846153846154 | 3UTR |
| hsa-miR-4796-3p | NM_001135099 | TMPRSS2 | 0.846153846154 | 3UTR |
| hsa-miR-4797-3p | NM_001135099 | TMPRSS2 | 0.846153846154 | 3UTR |
| hsa-miR-4802-5p | NM_001135099 | TMPRSS2 | 0.846153846154 | 3UTR |
| hsa-miR-4802-3p | NM_001135099 | TMPRSS2 | 0.846153846154 | 3UTR |
| hsa-miR-5001-3p | NM_001135099 | TMPRSS2 | 0.846153846154 | 3UTR |
| hsa-miR-5003-3p | NM_001135099 | TMPRSS2 | 0.846153846154 | 3UTR |
| hsa-miR-5006-3p | NM_001135099 | TMPRSS2 | 0.846153846154 | 3UTR |
| hsa-miR-5010-5p | NM_001135099 | TMPRSS2 | 0.846153846154 | 3UTR |
| hsa-miR-5092 | NM_001135099 | TMPRSS2 | 0.846153846154 | 3UTR |
| hsa-miR-5187-3p | NM_001135099 | TMPRSS2 | 0.846153846154 | 3UTR |
| hsa-miR-5193 | NM_001135099 | TMPRSS2 | 0.846153846154 | 3UTR |
| hsa-miR-5193 | NM_001135099 | TMPRSS2 | 0.846153846154 | 3UTR |
| hsa-miR-5194 | NM_001135099 | TMPRSS2 | 0.846153846154 | 3UTR |
| hsa-miR-5581-3p | NM_001135099 | TMPRSS2 | 0.846153846154 | 3UTR |
| hsa-miR-548at-5p | NM_001135099 | TMPRSS2 | 0.846153846154 | 3UTR |
| hsa-miR-5584-3p | NM_001135099 | TMPRSS2 | 0.846153846154 | 3UTR |
| hsa-miR-5587-5p | NM_001135099 | TMPRSS2 | 0.846153846154 | 3UTR |
| hsa-miR-1295b-5p | NM_001135099 | TMPRSS2 | 0.846153846154 | 3UTR |
| hsa-miR-5588-3p | NM_001135099 | TMPRSS2 | 0.846153846154 | 3UTR |
| hsa-miR-5591-5p | NM_001135099 | TMPRSS2 | 0.846153846154 | 3UTR |
| hsa-miR-5690 | NM_001135099 | TMPRSS2 | 0.846153846154 | 3UTR |

| | | | | |
|---|---|---|---|---|
| hsa-miR-5691 | NM_001135099 | TMPRSS2 | 0.846153846154 | 3UTR |
| hsa-miR-5699-5p | NM_001135099 | TMPRSS2 | 0.846153846154 | 3UTR |
| hsa-miR-5703 | NM_001135099 | TMPRSS2 | 0.846153846154 | 3UTR |
| hsa-miR-6072 | NM_001135099 | TMPRSS2 | 0.846153846154 | 3UTR |
| hsa-miR-6085 | NM_001135099 | TMPRSS2 | 0.846153846154 | 3UTR |
| hsa-miR-6086 | NM_001135099 | TMPRSS2 | 0.846153846154 | 3UTR |
| hsa-miR-6124 | NM_001135099 | TMPRSS2 | 0.846153846154 | 3UTR |
| hsa-miR-6124 | NM_001135099 | TMPRSS2 | 0.846153846154 | 3UTR |
| hsa-miR-378j | NM_001135099 | TMPRSS2 | 0.846153846154 | 3UTR |
| hsa-miR-6132 | NM_001135099 | TMPRSS2 | 0.846153846154 | 3UTR |
| hsa-miR-6132 | NM_001135099 | TMPRSS2 | 0.846153846154 | 3UTR |
| hsa-miR-6165 | NM_001135099 | TMPRSS2 | 0.846153846154 | 3UTR |
| hsa-miR-6499-3p | NM_001135099 | TMPRSS2 | 0.846153846154 | 3UTR |
| hsa-miR-6501-3p | NM_001135099 | TMPRSS2 | 0.846153846154 | 3UTR |
| hsa-miR-6504-5p | NM_001135099 | TMPRSS2 | 0.846153846154 | 3UTR |
| hsa-miR-6505-3p | NM_001135099 | TMPRSS2 | 0.846153846154 | 3UTR |
| hsa-miR-6509-5p | NM_001135099 | TMPRSS2 | 0.846153846154 | 3UTR |
| hsa-miR-6509-3p | NM_001135099 | TMPRSS2 | 0.846153846154 | 3UTR |
| hsa-miR-6510-3p | NM_001135099 | TMPRSS2 | 0.846153846154 | 3UTR |
| hsa-miR-6511a-5p | NM_001135099 | TMPRSS2 | 0.846153846154 | 3UTR |
| hsa-miR-6513-3p | NM_001135099 | TMPRSS2 | 0.846153846154 | 3UTR |
| hsa-miR-6721-5p | NM_001135099 | TMPRSS2 | 0.846153846154 | 3UTR |
| hsa-miR-892c-5p | NM_001135099 | TMPRSS2 | 0.846153846154 | 3UTR |
| hsa-miR-6726-3p | NM_001135099 | TMPRSS2 | 0.846153846154 | 3UTR |
| hsa-miR-6727-3p | NM_001135099 | TMPRSS2 | 0.846153846154 | 3UTR |
| hsa-miR-6729-5p | NM_001135099 | TMPRSS2 | 0.846153846154 | 3UTR |
| hsa-miR-6734-5p | NM_001135099 | TMPRSS2 | 0.846153846154 | 3UTR |
| hsa-miR-6734-3p | NM_001135099 | TMPRSS2 | 0.846153846154 | 3UTR |
| hsa-miR-6738-5p | NM_001135099 | TMPRSS2 | 0.846153846154 | 3UTR |
| hsa-miR-6739-3p | NM_001135099 | TMPRSS2 | 0.846153846154 | 3UTR |
| hsa-miR-6746-5p | NM_001135099 | TMPRSS2 | 0.846153846154 | 3UTR |
| hsa-miR-6750-3p | NM_001135099 | TMPRSS2 | 0.846153846154 | 3UTR |
| hsa-miR-6752-5p | NM_001135099 | TMPRSS2 | 0.846153846154 | 3UTR |
| hsa-miR-6753-5p | NM_001135099 | TMPRSS2 | 0.846153846154 | 3UTR |
| hsa-miR-6753-5p | NM_001135099 | TMPRSS2 | 0.846153846154 | 3UTR |
| hsa-miR-6754-5p | NM_001135099 | TMPRSS2 | 0.846153846154 | 3UTR |
| hsa-miR-6757-5p | NM_001135099 | TMPRSS2 | 0.846153846154 | 3UTR |
| hsa-miR-6759-5p | NM_001135099 | TMPRSS2 | 0.846153846154 | 3UTR |
| hsa-miR-6760-5p | NM_001135099 | TMPRSS2 | 0.846153846154 | 3UTR |
| hsa-miR-6762-5p | NM_001135099 | TMPRSS2 | 0.846153846154 | 3UTR |
| hsa-miR-6767-3p | NM_001135099 | TMPRSS2 | 0.846153846154 | 3UTR |
| hsa-miR-6768-3p | NM_001135099 | TMPRSS2 | 0.846153846154 | 3UTR |
| hsa-miR-6769a-5p | NM_001135099 | TMPRSS2 | 0.846153846154 | 3UTR |
| hsa-miR-6771-3p | NM_001135099 | TMPRSS2 | 0.846153846154 | 3UTR |
| hsa-miR-6772-5p | NM_001135099 | TMPRSS2 | 0.846153846154 | 3UTR |
| hsa-miR-6778-3p | NM_001135099 | TMPRSS2 | 0.846153846154 | 3UTR |
| hsa-miR-6779-5p | NM_001135099 | TMPRSS2 | 0.846153846154 | 3UTR |

| | | | | |
|---|---|---|---|---|
| hsa-miR-6781-5p | NM_001135099 | TMPRSS2 | 0.846153846154 | 3UTR |
| hsa-miR-6784-3p | NM_001135099 | TMPRSS2 | 0.846153846154 | 3UTR |
| hsa-miR-6787-3p | NM_001135099 | TMPRSS2 | 0.846153846154 | 3UTR |
| hsa-miR-6788-3p | NM_001135099 | TMPRSS2 | 0.846153846154 | 3UTR |
| hsa-miR-6792-5p | NM_001135099 | TMPRSS2 | 0.846153846154 | 3UTR |
| hsa-miR-6792-3p | NM_001135099 | TMPRSS2 | 0.846153846154 | 3UTR |
| hsa-miR-6793-3p | NM_001135099 | TMPRSS2 | 0.846153846154 | 3UTR |
| hsa-miR-6794-5p | NM_001135099 | TMPRSS2 | 0.846153846154 | 3UTR |
| hsa-miR-6796-5p | NM_001135099 | TMPRSS2 | 0.846153846154 | 3UTR |
| hsa-miR-6797-5p | NM_001135099 | TMPRSS2 | 0.846153846154 | 3UTR |
| hsa-miR-6797-3p | NM_001135099 | TMPRSS2 | 0.846153846154 | 3UTR |
| hsa-miR-6801-5p | NM_001135099 | TMPRSS2 | 0.846153846154 | 3UTR |
| hsa-miR-6803-5p | NM_001135099 | TMPRSS2 | 0.846153846154 | 3UTR |
| hsa-miR-6804-5p | NM_001135099 | TMPRSS2 | 0.846153846154 | 3UTR |
| hsa-miR-6811-5p | NM_001135099 | TMPRSS2 | 0.846153846154 | 3UTR |
| hsa-miR-6814-5p | NM_001135099 | TMPRSS2 | 0.846153846154 | 3UTR |
| hsa-miR-6815-5p | NM_001135099 | TMPRSS2 | 0.846153846154 | 3UTR |
| hsa-miR-6826-5p | NM_001135099 | TMPRSS2 | 0.846153846154 | 3UTR |
| hsa-miR-6828-5p | NM_001135099 | TMPRSS2 | 0.846153846154 | 3UTR |
| hsa-miR-6829-5p | NM_001135099 | TMPRSS2 | 0.846153846154 | 3UTR |
| hsa-miR-6830-5p | NM_001135099 | TMPRSS2 | 0.846153846154 | 3UTR |
| hsa-miR-6830-3p | NM_001135099 | TMPRSS2 | 0.846153846154 | 3UTR |
| hsa-miR-6831-5p | NM_001135099 | TMPRSS2 | 0.846153846154 | 3UTR |
| hsa-miR-6831-5p | NM_001135099 | TMPRSS2 | 0.846153846154 | 3UTR |
| hsa-miR-6832-3p | NM_001135099 | TMPRSS2 | 0.846153846154 | 3UTR |
| hsa-miR-6833-3p | NM_001135099 | TMPRSS2 | 0.846153846154 | 3UTR |
| hsa-miR-6834-3p | NM_001135099 | TMPRSS2 | 0.846153846154 | 3UTR |
| hsa-miR-6835-5p | NM_001135099 | TMPRSS2 | 0.846153846154 | 3UTR |
| hsa-miR-6835-3p | NM_001135099 | TMPRSS2 | 0.846153846154 | 3UTR |
| hsa-miR-6780b-5p | NM_001135099 | TMPRSS2 | 0.846153846154 | 3UTR |
| hsa-miR-6780b-5p | NM_001135099 | TMPRSS2 | 0.846153846154 | 3UTR |
| hsa-miR-6845-5p | NM_001135099 | TMPRSS2 | 0.846153846154 | 3UTR |
| hsa-miR-6845-3p | NM_001135099 | TMPRSS2 | 0.846153846154 | 3UTR |
| hsa-miR-6847-5p | NM_001135099 | TMPRSS2 | 0.846153846154 | 3UTR |
| hsa-miR-6847-5p | NM_001135099 | TMPRSS2 | 0.846153846154 | 3UTR |
| hsa-miR-6848-3p | NM_001135099 | TMPRSS2 | 0.846153846154 | 3UTR |
| hsa-miR-6849-3p | NM_001135099 | TMPRSS2 | 0.846153846154 | 3UTR |
| hsa-miR-6853-5p | NM_001135099 | TMPRSS2 | 0.846153846154 | 3UTR |
| hsa-miR-6854-3p | NM_001135099 | TMPRSS2 | 0.846153846154 | 3UTR |
| hsa-miR-6857-3p | NM_001135099 | TMPRSS2 | 0.846153846154 | 3UTR |
| hsa-miR-6858-5p | NM_001135099 | TMPRSS2 | 0.846153846154 | 3UTR |
| hsa-miR-6769b-5p | NM_001135099 | TMPRSS2 | 0.846153846154 | 3UTR |
| hsa-miR-6865-5p | NM_001135099 | TMPRSS2 | 0.846153846154 | 3UTR |
| hsa-miR-6866-3p | NM_001135099 | TMPRSS2 | 0.846153846154 | 3UTR |
| hsa-miR-6867-3p | NM_001135099 | TMPRSS2 | 0.846153846154 | 3UTR |
| hsa-miR-6871-5p | NM_001135099 | TMPRSS2 | 0.846153846154 | 3UTR |
| hsa-miR-6872-5p | NM_001135099 | TMPRSS2 | 0.846153846154 | 3UTR |

| | | | | |
|---|---|---|---|---|
| hsa-miR-6876-5p | NM_001135099 | TMPRSS2 | 0.846153846154 | 3UTR |
| hsa-miR-6877-5p | NM_001135099 | TMPRSS2 | 0.846153846154 | 3UTR |
| hsa-miR-6877-5p | NM_001135099 | TMPRSS2 | 0.846153846154 | 3UTR |
| hsa-miR-6878-5p | NM_001135099 | TMPRSS2 | 0.846153846154 | 3UTR |
| hsa-miR-6882-5p | NM_001135099 | TMPRSS2 | 0.846153846154 | 3UTR |
| hsa-miR-6883-5p | NM_001135099 | TMPRSS2 | 0.846153846154 | 3UTR |
| hsa-miR-6885-5p | NM_001135099 | TMPRSS2 | 0.846153846154 | 3UTR |
| hsa-miR-6886-5p | NM_001135099 | TMPRSS2 | 0.846153846154 | 3UTR |
| hsa-miR-6890-5p | NM_001135099 | TMPRSS2 | 0.846153846154 | 3UTR |
| hsa-miR-6893-5p | NM_001135099 | TMPRSS2 | 0.846153846154 | 3UTR |
| hsa-miR-6893-3p | NM_001135099 | TMPRSS2 | 0.846153846154 | 3UTR |
| hsa-miR-6894-5p | NM_001135099 | TMPRSS2 | 0.846153846154 | 3UTR |
| hsa-miR-7107-5p | NM_001135099 | TMPRSS2 | 0.846153846154 | 3UTR |
| hsa-miR-7111-5p | NM_001135099 | TMPRSS2 | 0.846153846154 | 3UTR |
| hsa-miR-7113-3p | NM_001135099 | TMPRSS2 | 0.846153846154 | 3UTR |
| hsa-miR-7150 | NM_001135099 | TMPRSS2 | 0.846153846154 | 3UTR |
| hsa-miR-7151-3p | NM_001135099 | TMPRSS2 | 0.846153846154 | 3UTR |
| hsa-miR-7154-5p | NM_001135099 | TMPRSS2 | 0.846153846154 | 3UTR |
| hsa-miR-7156-5p | NM_001135099 | TMPRSS2 | 0.846153846154 | 3UTR |
| hsa-miR-7157-5p | NM_001135099 | TMPRSS2 | 0.846153846154 | 3UTR |
| hsa-miR-7158-5p | NM_001135099 | TMPRSS2 | 0.846153846154 | 3UTR |
| hsa-miR-7162-5p | NM_001135099 | TMPRSS2 | 0.846153846154 | 3UTR |
| hsa-miR-7702 | NM_001135099 | TMPRSS2 | 0.846153846154 | 3UTR |
| hsa-miR-7706 | NM_001135099 | TMPRSS2 | 0.846153846154 | 3UTR |
| hsa-miR-7843-3p | NM_001135099 | TMPRSS2 | 0.846153846154 | 3UTR |
| hsa-miR-1273h-3p | NM_001135099 | TMPRSS2 | 0.846153846154 | 3UTR |
| hsa-miR-7845-5p | NM_001135099 | TMPRSS2 | 0.846153846154 | 3UTR |
| hsa-miR-7845-5p | NM_001135099 | TMPRSS2 | 0.846153846154 | 3UTR |
| hsa-miR-7851-3p | NM_001135099 | TMPRSS2 | 0.846153846154 | 3UTR |
| hsa-miR-7854-3p | NM_001135099 | TMPRSS2 | 0.846153846154 | 3UTR |
| hsa-miR-7856-5p | NM_001135099 | TMPRSS2 | 0.846153846154 | 3UTR |
| hsa-miR-7974 | NM_001135099 | TMPRSS2 | 0.846153846154 | 3UTR |
| hsa-miR-8059 | NM_001135099 | TMPRSS2 | 0.846153846154 | 3UTR |
| hsa-miR-8060 | NM_001135099 | TMPRSS2 | 0.846153846154 | 3UTR |
| hsa-miR-8063 | NM_001135099 | TMPRSS2 | 0.846153846154 | 3UTR |
| hsa-miR-8065 | NM_001135099 | TMPRSS2 | 0.846153846154 | 3UTR |
| hsa-miR-8078 | NM_001135099 | TMPRSS2 | 0.846153846154 | 3UTR |
| hsa-miR-8082 | NM_001135099 | TMPRSS2 | 0.846153846154 | 3UTR |
| hsa-miR-8086 | NM_001135099 | TMPRSS2 | 0.846153846154 | 3UTR |
| hsa-miR-9901 | NM_001135099 | TMPRSS2 | 0.846153846154 | 3UTR |
| hsa-miR-10522-5p | NM_001135099 | TMPRSS2 | 0.846153846154 | 3UTR |
| hsa-miR-11399 | NM_001135099 | TMPRSS2 | 0.846153846154 | 3UTR |
| hsa-miR-3085-5p | NM_001135099 | TMPRSS2 | 0.846153846154 | 3UTR |
| hsa-miR-12114 | NM_001135099 | TMPRSS2 | 0.846153846154 | 3UTR |
| hsa-miR-12127 | NM_001135099 | TMPRSS2 | 0.846153846154 | 3UTR |
| hsa-miR-3674 | NM_005656 | TMPRSS2 | 0.858974358974 | 3UTR |
| hsa-miR-7153-3p | NM_005656 | TMPRSS2 | 0.861538461538 | 3UTR |

| | | | | |
|---|---|---|---|---|
| hsa-miR-7153-3p | NM_001135099 | TMPRSS2 | 0.861538461538 | 3UTR |
| hsa-miR-4720-3p | NM_001135099 | TMPRSS2 | 0.862637362637 | 3UTR |
| hsa-miR-133b | NM_005656 | TMPRSS2 | 0.865384615385 | 3UTR |
| hsa-miR-302a-3p | NM_001135099 | TMPRSS2 | 0.865384615385 | 3UTR |
| hsa-miR-4293 | NM_001135099 | TMPRSS2 | 0.865384615385 | 3UTR |
| hsa-miR-523-3p | NM_005656 | TMPRSS2 | 0.871794871795 | 3UTR |
| hsa-miR-598-3p | NM_005656 | TMPRSS2 | 0.871794871795 | 3UTR |
| hsa-miR-1207-5p | NM_005656 | TMPRSS2 | 0.871794871795 | 3UTR |
| hsa-miR-1908-3p | NM_005656 | TMPRSS2 | 0.871794871795 | 3UTR |
| hsa-miR-6749-5p | NM_005656 | TMPRSS2 | 0.871794871795 | 3UTR |
| hsa-miR-6803-5p | NM_005656 | TMPRSS2 | 0.871794871795 | 3UTR |
| hsa-miR-195-5p | NM_001135099 | TMPRSS2 | 0.871794871795 | 3UTR |
| hsa-miR-381-3p | NM_001135099 | TMPRSS2 | 0.871794871795 | 3UTR |
| hsa-miR-3945 | NM_001135099 | TMPRSS2 | 0.871794871795 | 3UTR |
| hsa-miR-4740-5p | NM_001135099 | TMPRSS2 | 0.871794871795 | 3UTR |
| hsa-miR-6849-5p | NM_001135099 | TMPRSS2 | 0.871794871795 | 3UTR |
| hsa-miR-7156-5p | NM_001135099 | TMPRSS2 | 0.871794871795 | 3UTR |
| hsa-miR-10394-5p | NM_001135099 | TMPRSS2 | 0.876923076923 | 3UTR |
| hsa-miR-608 | NM_005656 | TMPRSS2 | 0.884615384615 | 3UTR |
| hsa-miR-659-5p | NM_005656 | TMPRSS2 | 0.884615384615 | 3UTR |
| hsa-miR-4314 | NM_005656 | TMPRSS2 | 0.884615384615 | 3UTR |
| hsa-miR-4270 | NM_005656 | TMPRSS2 | 0.884615384615 | 3UTR |
| hsa-miR-4290 | NM_005656 | TMPRSS2 | 0.884615384615 | 3UTR |
| hsa-miR-4430 | NM_005656 | TMPRSS2 | 0.884615384615 | 3UTR |
| hsa-miR-4689 | NM_005656 | TMPRSS2 | 0.884615384615 | 3UTR |
| hsa-miR-4720-3p | NM_005656 | TMPRSS2 | 0.884615384615 | 3UTR |
| hsa-miR-4769-5p | NM_005656 | TMPRSS2 | 0.884615384615 | 3UTR |
| hsa-miR-5094 | NM_005656 | TMPRSS2 | 0.884615384615 | 3UTR |
| hsa-miR-5588-5p | NM_005656 | TMPRSS2 | 0.884615384615 | 3UTR |
| hsa-miR-5699-3p | NM_005656 | TMPRSS2 | 0.884615384615 | 3UTR |
| hsa-miR-6132 | NM_005656 | TMPRSS2 | 0.884615384615 | 3UTR |
| hsa-miR-6501-3p | NM_005656 | TMPRSS2 | 0.884615384615 | 3UTR |
| hsa-miR-6862-5p | NM_005656 | TMPRSS2 | 0.884615384615 | 3UTR |
| hsa-miR-6870-5p | NM_005656 | TMPRSS2 | 0.884615384615 | 3UTR |
| hsa-miR-6876-3p | NM_005656 | TMPRSS2 | 0.884615384615 | 3UTR |
| hsa-miR-608 | NM_001135099 | TMPRSS2 | 0.884615384615 | 3UTR |
| hsa-miR-616-3p | NM_001135099 | TMPRSS2 | 0.884615384615 | 3UTR |
| hsa-miR-4283 | NM_001135099 | TMPRSS2 | 0.884615384615 | 3UTR |
| hsa-miR-3663-5p | NM_001135099 | TMPRSS2 | 0.884615384615 | 3UTR |
| hsa-miR-4430 | NM_001135099 | TMPRSS2 | 0.884615384615 | 3UTR |
| hsa-miR-5699-3p | NM_001135099 | TMPRSS2 | 0.884615384615 | 3UTR |
| hsa-miR-6737-5p | NM_001135099 | TMPRSS2 | 0.884615384615 | 3UTR |
| hsa-miR-6878-5p | NM_001135099 | TMPRSS2 | 0.884615384615 | 3UTR |
| hsa-miR-3174 | NM_005656 | TMPRSS2 | 0.892307692308 | 3UTR |
| hsa-miR-3689a-3p | NM_001135099 | TMPRSS2 | 0.892307692308 | 3UTR |
| hsa-miR-3929 | NM_001135099 | TMPRSS2 | 0.892307692308 | 3UTR |
| hsa-miR-520c-3p | NM_005656 | TMPRSS2 | 0.897435897436 | 3UTR |

| | | | | |
|---|---|---|---|---|
| hsa-miR-33b-3p | NM_005656 | TMPRSS2 | 0.897435897436 | 3UTR |
| hsa-miR-6746-5p | NM_005656 | TMPRSS2 | 0.897435897436 | 3UTR |
| hsa-miR-7114-5p | NM_005656 | TMPRSS2 | 0.897435897436 | 3UTR |
| hsa-miR-141-3p | NM_001135099 | TMPRSS2 | 0.897435897436 | 3UTR |
| hsa-miR-520c-3p | NM_001135099 | TMPRSS2 | 0.897435897436 | 3UTR |
| hsa-miR-642b-3p | NM_001135099 | TMPRSS2 | 0.897435897436 | 3UTR |
| hsa-miR-4802-5p | NM_001135099 | TMPRSS2 | 0.897435897436 | 3UTR |
| hsa-miR-27b-5p | NM_005656 | TMPRSS2 | 0.903846153846 | 3UTR |
| hsa-miR-6726-5p | NM_005656 | TMPRSS2 | 0.903846153846 | 3UTR |
| hsa-miR-6726-5p | NM_001135099 | TMPRSS2 | 0.903846153846 | 3UTR |
| hsa-miR-4769-5p | NM_001135099 | TMPRSS2 | 0.910256410256 | 3UTR |
| hsa-miR-1251-5p | NM_005656 | TMPRSS2 | 0.912087912088 | 3UTR |
| hsa-let-7a-5p | NM_005656 | TMPRSS2 | 0.923076923077 | 3UTR |
| hsa-let-7a-2-3p | NM_005656 | TMPRSS2 | 0.923076923077 | 3UTR |
| hsa-let-7c-5p | NM_005656 | TMPRSS2 | 0.923076923077 | 3UTR |
| hsa-let-7e-5p | NM_005656 | TMPRSS2 | 0.923076923077 | 3UTR |
| hsa-let-7f-5p | NM_005656 | TMPRSS2 | 0.923076923077 | 3UTR |
| hsa-miR-17-5p | NM_005656 | TMPRSS2 | 0.923076923077 | 3UTR |
| hsa-miR-17-5p | NM_005656 | TMPRSS2 | 0.923076923077 | 3UTR |
| hsa-miR-17-5p | NM_005656 | TMPRSS2 | 0.923076923077 | 3UTR |
| hsa-miR-21-3p | NM_005656 | TMPRSS2 | 0.923076923077 | 3UTR |
| hsa-miR-24-3p | NM_005656 | TMPRSS2 | 0.923076923077 | 3UTR |
| hsa-miR-30a-3p | NM_005656 | TMPRSS2 | 0.923076923077 | 3UTR |
| hsa-miR-92a-1-5p | NM_005656 | TMPRSS2 | 0.923076923077 | 3UTR |
| hsa-miR-93-5p | NM_005656 | TMPRSS2 | 0.923076923077 | 3UTR |
| hsa-miR-95-3p | NM_005656 | TMPRSS2 | 0.923076923077 | 3UTR |
| hsa-miR-103a-2-5p | NM_005656 | TMPRSS2 | 0.923076923077 | 3UTR |
| hsa-miR-106a-5p | NM_005656 | TMPRSS2 | 0.923076923077 | 3UTR |
| hsa-miR-106a-5p | NM_005656 | TMPRSS2 | 0.923076923077 | 3UTR |
| hsa-miR-106a-5p | NM_005656 | TMPRSS2 | 0.923076923077 | 3UTR |
| hsa-miR-16-2-3p | NM_005656 | TMPRSS2 | 0.923076923077 | 3UTR |
| hsa-miR-196a-5p | NM_005656 | TMPRSS2 | 0.923076923077 | 3UTR |
| hsa-miR-197-5p | NM_005656 | TMPRSS2 | 0.923076923077 | 3UTR |
| hsa-miR-197-3p | NM_005656 | TMPRSS2 | 0.923076923077 | 3UTR |
| hsa-miR-198 | NM_005656 | TMPRSS2 | 0.923076923077 | 3UTR |
| hsa-miR-199a-3p | NM_005656 | TMPRSS2 | 0.923076923077 | 3UTR |
| hsa-miR-129-1-3p | NM_005656 | TMPRSS2 | 0.923076923077 | 3UTR |
| hsa-miR-30d-3p | NM_005656 | TMPRSS2 | 0.923076923077 | 3UTR |
| hsa-miR-7-5p | NM_005656 | TMPRSS2 | 0.923076923077 | 3UTR |
| hsa-miR-182-5p | NM_005656 | TMPRSS2 | 0.923076923077 | 3UTR |
| hsa-miR-187-5p | NM_005656 | TMPRSS2 | 0.923076923077 | 3UTR |
| hsa-miR-199b-3p | NM_005656 | TMPRSS2 | 0.923076923077 | 3UTR |
| hsa-miR-216a-3p | NM_005656 | TMPRSS2 | 0.923076923077 | 3UTR |
| hsa-miR-218-1-3p | NM_005656 | TMPRSS2 | 0.923076923077 | 3UTR |
| hsa-miR-224-5p | NM_005656 | TMPRSS2 | 0.923076923077 | 3UTR |
| hsa-miR-224-3p | NM_005656 | TMPRSS2 | 0.923076923077 | 3UTR |
| hsa-miR-224-3p | NM_005656 | TMPRSS2 | 0.923076923077 | 3UTR |

| | | | | |
|---|---|---|---|---|
| hsa-miR-23b-5p | NM_005656 | TMPRSS2 | 0.923076923077 | 3UTR |
| hsa-miR-27b-3p | NM_005656 | TMPRSS2 | 0.923076923077 | 3UTR |
| hsa-miR-128-1-5p | NM_005656 | TMPRSS2 | 0.923076923077 | 3UTR |
| hsa-miR-128-1-5p | NM_005656 | TMPRSS2 | 0.923076923077 | 3UTR |
| hsa-miR-132-3p | NM_005656 | TMPRSS2 | 0.923076923077 | 3UTR |
| hsa-miR-135a-2-3p | NM_005656 | TMPRSS2 | 0.923076923077 | 3UTR |
| hsa-miR-138-2-3p | NM_005656 | TMPRSS2 | 0.923076923077 | 3UTR |
| hsa-miR-140-5p | NM_005656 | TMPRSS2 | 0.923076923077 | 3UTR |
| hsa-miR-140-3p | NM_005656 | TMPRSS2 | 0.923076923077 | 3UTR |
| hsa-miR-145-3p | NM_005656 | TMPRSS2 | 0.923076923077 | 3UTR |
| hsa-miR-152-3p | NM_005656 | TMPRSS2 | 0.923076923077 | 3UTR |
| hsa-miR-153-5p | NM_005656 | TMPRSS2 | 0.923076923077 | 3UTR |
| hsa-miR-9-5p | NM_005656 | TMPRSS2 | 0.923076923077 | 3UTR |
| hsa-miR-9-5p | NM_005656 | TMPRSS2 | 0.923076923077 | 3UTR |
| hsa-miR-193a-5p | NM_005656 | TMPRSS2 | 0.923076923077 | 3UTR |
| hsa-miR-195-5p | NM_005656 | TMPRSS2 | 0.923076923077 | 3UTR |
| hsa-miR-29c-5p | NM_005656 | TMPRSS2 | 0.923076923077 | 3UTR |
| hsa-miR-29c-5p | NM_005656 | TMPRSS2 | 0.923076923077 | 3UTR |
| hsa-miR-34c-3p | NM_005656 | TMPRSS2 | 0.923076923077 | 3UTR |
| hsa-miR-296-3p | NM_005656 | TMPRSS2 | 0.923076923077 | 3UTR |
| hsa-miR-296-3p | NM_005656 | TMPRSS2 | 0.923076923077 | 3UTR |
| hsa-miR-361-5p | NM_005656 | TMPRSS2 | 0.923076923077 | 3UTR |
| hsa-miR-365b-5p | NM_005656 | TMPRSS2 | 0.923076923077 | 3UTR |
| hsa-miR-370-3p | NM_005656 | TMPRSS2 | 0.923076923077 | 3UTR |
| hsa-miR-331-5p | NM_005656 | TMPRSS2 | 0.923076923077 | 3UTR |
| hsa-miR-324-5p | NM_005656 | TMPRSS2 | 0.923076923077 | 3UTR |
| hsa-miR-339-5p | NM_005656 | TMPRSS2 | 0.923076923077 | 3UTR |
| hsa-miR-339-3p | NM_005656 | TMPRSS2 | 0.923076923077 | 3UTR |
| hsa-miR-423-5p | NM_005656 | TMPRSS2 | 0.923076923077 | 3UTR |
| hsa-miR-20b-5p | NM_005656 | TMPRSS2 | 0.923076923077 | 3UTR |
| hsa-miR-20b-5p | NM_005656 | TMPRSS2 | 0.923076923077 | 3UTR |
| hsa-miR-20b-5p | NM_005656 | TMPRSS2 | 0.923076923077 | 3UTR |
| hsa-miR-448 | NM_005656 | TMPRSS2 | 0.923076923077 | 3UTR |
| hsa-miR-449a | NM_005656 | TMPRSS2 | 0.923076923077 | 3UTR |
| hsa-miR-449a | NM_005656 | TMPRSS2 | 0.923076923077 | 3UTR |
| hsa-miR-329-5p | NM_005656 | TMPRSS2 | 0.923076923077 | 3UTR |
| hsa-miR-329-5p | NM_005656 | TMPRSS2 | 0.923076923077 | 3UTR |
| hsa-miR-483-3p | NM_005656 | TMPRSS2 | 0.923076923077 | 3UTR |
| hsa-miR-485-3p | NM_005656 | TMPRSS2 | 0.923076923077 | 3UTR |
| hsa-miR-486-3p | NM_005656 | TMPRSS2 | 0.923076923077 | 3UTR |
| hsa-miR-491-5p | NM_005656 | TMPRSS2 | 0.923076923077 | 3UTR |
| hsa-miR-491-3p | NM_005656 | TMPRSS2 | 0.923076923077 | 3UTR |
| hsa-miR-146b-3p | NM_005656 | TMPRSS2 | 0.923076923077 | 3UTR |
| hsa-miR-492 | NM_005656 | TMPRSS2 | 0.923076923077 | 3UTR |
| hsa-miR-432-5p | NM_005656 | TMPRSS2 | 0.923076923077 | 3UTR |
| hsa-miR-512-5p | NM_005656 | TMPRSS2 | 0.923076923077 | 3UTR |
| hsa-miR-512-5p | NM_005656 | TMPRSS2 | 0.923076923077 | 3UTR |

| | | | | |
|---|---|---|---|---|
| hsa-miR-498-5p | NM_005656 | TMPRSS2 | 0.923076923077 | 3UTR |
| hsa-miR-520e-5p | NM_005656 | TMPRSS2 | 0.923076923077 | 3UTR |
| hsa-miR-515-5p | NM_005656 | TMPRSS2 | 0.923076923077 | 3UTR |
| hsa-miR-520f-3p | NM_005656 | TMPRSS2 | 0.923076923077 | 3UTR |
| hsa-miR-519c-5p | NM_005656 | TMPRSS2 | 0.923076923077 | 3UTR |
| hsa-miR-520a-5p | NM_005656 | TMPRSS2 | 0.923076923077 | 3UTR |
| hsa-miR-526b-5p | NM_005656 | TMPRSS2 | 0.923076923077 | 3UTR |
| hsa-miR-519b-5p | NM_005656 | TMPRSS2 | 0.923076923077 | 3UTR |
| hsa-miR-523-5p | NM_005656 | TMPRSS2 | 0.923076923077 | 3UTR |
| hsa-miR-523-3p | NM_005656 | TMPRSS2 | 0.923076923077 | 3UTR |
| hsa-miR-518f-5p | NM_005656 | TMPRSS2 | 0.923076923077 | 3UTR |
| hsa-miR-524-3p | NM_005656 | TMPRSS2 | 0.923076923077 | 3UTR |
| hsa-miR-520d-5p | NM_005656 | TMPRSS2 | 0.923076923077 | 3UTR |
| hsa-miR-516b-3p | NM_005656 | TMPRSS2 | 0.923076923077 | 3UTR |
| hsa-miR-516b-3p | NM_005656 | TMPRSS2 | 0.923076923077 | 3UTR |
| hsa-miR-518e-5p | NM_005656 | TMPRSS2 | 0.923076923077 | 3UTR |
| hsa-miR-520h | NM_005656 | TMPRSS2 | 0.923076923077 | 3UTR |
| hsa-miR-522-5p | NM_005656 | TMPRSS2 | 0.923076923077 | 3UTR |
| hsa-miR-519a-5p | NM_005656 | TMPRSS2 | 0.923076923077 | 3UTR |
| hsa-miR-516a-3p | NM_005656 | TMPRSS2 | 0.923076923077 | 3UTR |
| hsa-miR-516a-3p | NM_005656 | TMPRSS2 | 0.923076923077 | 3UTR |
| hsa-miR-500a-3p | NM_005656 | TMPRSS2 | 0.923076923077 | 3UTR |
| hsa-miR-501-5p | NM_005656 | TMPRSS2 | 0.923076923077 | 3UTR |
| hsa-miR-504-5p | NM_005656 | TMPRSS2 | 0.923076923077 | 3UTR |
| hsa-miR-513a-5p | NM_005656 | TMPRSS2 | 0.923076923077 | 3UTR |
| hsa-miR-507 | NM_005656 | TMPRSS2 | 0.923076923077 | 3UTR |
| hsa-miR-509-3p | NM_005656 | TMPRSS2 | 0.923076923077 | 3UTR |
| hsa-miR-532-3p | NM_005656 | TMPRSS2 | 0.923076923077 | 3UTR |
| hsa-miR-376a-2-5p | NM_005656 | TMPRSS2 | 0.923076923077 | 3UTR |
| hsa-miR-562 | NM_005656 | TMPRSS2 | 0.923076923077 | 3UTR |
| hsa-miR-584-5p | NM_005656 | TMPRSS2 | 0.923076923077 | 3UTR |
| hsa-miR-548a-3p | NM_005656 | TMPRSS2 | 0.923076923077 | 3UTR |
| hsa-miR-548b-3p | NM_005656 | TMPRSS2 | 0.923076923077 | 3UTR |
| hsa-miR-589-5p | NM_005656 | TMPRSS2 | 0.923076923077 | 3UTR |
| hsa-miR-591 | NM_005656 | TMPRSS2 | 0.923076923077 | 3UTR |
| hsa-miR-601 | NM_005656 | TMPRSS2 | 0.923076923077 | 3UTR |
| hsa-miR-604 | NM_005656 | TMPRSS2 | 0.923076923077 | 3UTR |
| hsa-miR-609 | NM_005656 | TMPRSS2 | 0.923076923077 | 3UTR |
| hsa-miR-616-5p | NM_005656 | TMPRSS2 | 0.923076923077 | 3UTR |
| hsa-miR-616-3p | NM_005656 | TMPRSS2 | 0.923076923077 | 3UTR |
| hsa-miR-617 | NM_005656 | TMPRSS2 | 0.923076923077 | 3UTR |
| hsa-miR-618 | NM_005656 | TMPRSS2 | 0.923076923077 | 3UTR |
| hsa-miR-619-5p | NM_005656 | TMPRSS2 | 0.923076923077 | 3UTR |
| hsa-miR-619-5p | NM_005656 | TMPRSS2 | 0.923076923077 | 3UTR |
| hsa-miR-619-3p | NM_005656 | TMPRSS2 | 0.923076923077 | 3UTR |
| hsa-miR-625-5p | NM_005656 | TMPRSS2 | 0.923076923077 | 3UTR |
| hsa-miR-627-5p | NM_005656 | TMPRSS2 | 0.923076923077 | 3UTR |

| | | | | |
|---|---|---|---|---|
| hsa-miR-632 | NM_005656 | TMPRSS2 | 0.923076923077 | 3UTR |
| hsa-miR-635 | NM_005656 | TMPRSS2 | 0.923076923077 | 3UTR |
| hsa-miR-643 | NM_005656 | TMPRSS2 | 0.923076923077 | 3UTR |
| hsa-miR-646 | NM_005656 | TMPRSS2 | 0.923076923077 | 3UTR |
| hsa-miR-646 | NM_005656 | TMPRSS2 | 0.923076923077 | 3UTR |
| hsa-miR-648 | NM_005656 | TMPRSS2 | 0.923076923077 | 3UTR |
| hsa-miR-652-3p | NM_005656 | TMPRSS2 | 0.923076923077 | 3UTR |
| hsa-miR-449b-3p | NM_005656 | TMPRSS2 | 0.923076923077 | 3UTR |
| hsa-miR-449b-3p | NM_005656 | TMPRSS2 | 0.923076923077 | 3UTR |
| hsa-miR-550a-3-5p | NM_005656 | TMPRSS2 | 0.923076923077 | 3UTR |
| hsa-miR-151b | NM_005656 | TMPRSS2 | 0.923076923077 | 3UTR |
| hsa-miR-1323 | NM_005656 | TMPRSS2 | 0.923076923077 | 3UTR |
| hsa-miR-1271-5p | NM_005656 | TMPRSS2 | 0.923076923077 | 3UTR |
| hsa-miR-1271-3p | NM_005656 | TMPRSS2 | 0.923076923077 | 3UTR |
| hsa-miR-1301-3p | NM_005656 | TMPRSS2 | 0.923076923077 | 3UTR |
| hsa-miR-769-3p | NM_005656 | TMPRSS2 | 0.923076923077 | 3UTR |
| hsa-miR-378d | NM_005656 | TMPRSS2 | 0.923076923077 | 3UTR |
| hsa-miR-675-3p | NM_005656 | TMPRSS2 | 0.923076923077 | 3UTR |
| hsa-miR-874-3p | NM_005656 | TMPRSS2 | 0.923076923077 | 3UTR |
| hsa-miR-890 | NM_005656 | TMPRSS2 | 0.923076923077 | 3UTR |
| hsa-miR-892b | NM_005656 | TMPRSS2 | 0.923076923077 | 3UTR |
| hsa-miR-875-5p | NM_005656 | TMPRSS2 | 0.923076923077 | 3UTR |
| hsa-miR-885-5p | NM_005656 | TMPRSS2 | 0.923076923077 | 3UTR |
| hsa-miR-877-5p | NM_005656 | TMPRSS2 | 0.923076923077 | 3UTR |
| hsa-miR-887-5p | NM_005656 | TMPRSS2 | 0.923076923077 | 3UTR |
| hsa-miR-665 | NM_005656 | TMPRSS2 | 0.923076923077 | 3UTR |
| hsa-miR-216b-5p | NM_005656 | TMPRSS2 | 0.923076923077 | 3UTR |
| hsa-miR-936 | NM_005656 | TMPRSS2 | 0.923076923077 | 3UTR |
| hsa-miR-1181 | NM_005656 | TMPRSS2 | 0.923076923077 | 3UTR |
| hsa-miR-1182 | NM_005656 | TMPRSS2 | 0.923076923077 | 3UTR |
| hsa-miR-1184 | NM_005656 | TMPRSS2 | 0.923076923077 | 3UTR |
| hsa-miR-1236-5p | NM_005656 | TMPRSS2 | 0.923076923077 | 3UTR |
| hsa-miR-1236-3p | NM_005656 | TMPRSS2 | 0.923076923077 | 3UTR |
| hsa-miR-1202 | NM_005656 | TMPRSS2 | 0.923076923077 | 3UTR |
| hsa-miR-1289 | NM_005656 | TMPRSS2 | 0.923076923077 | 3UTR |
| hsa-miR-1289 | NM_005656 | TMPRSS2 | 0.923076923077 | 3UTR |
| hsa-miR-1291 | NM_005656 | TMPRSS2 | 0.923076923077 | 3UTR |
| hsa-miR-548k | NM_005656 | TMPRSS2 | 0.923076923077 | 3UTR |
| hsa-miR-1243 | NM_005656 | TMPRSS2 | 0.923076923077 | 3UTR |
| hsa-miR-1249-5p | NM_005656 | TMPRSS2 | 0.923076923077 | 3UTR |
| hsa-miR-1250-5p | NM_005656 | TMPRSS2 | 0.923076923077 | 3UTR |
| hsa-miR-1251-5p | NM_005656 | TMPRSS2 | 0.923076923077 | 3UTR |
| hsa-miR-1258 | NM_005656 | TMPRSS2 | 0.923076923077 | 3UTR |
| hsa-miR-1265 | NM_005656 | TMPRSS2 | 0.923076923077 | 3UTR |
| hsa-miR-1270 | NM_005656 | TMPRSS2 | 0.923076923077 | 3UTR |
| hsa-miR-1275 | NM_005656 | TMPRSS2 | 0.923076923077 | 3UTR |
| hsa-miR-1276 | NM_005656 | TMPRSS2 | 0.923076923077 | 3UTR |

| | | | | |
|---|---|---|---|---|
| hsa-miR-1321 | NM_005656 | TMPRSS2 | 0.923076923077 | 3UTR |
| hsa-miR-1321 | NM_005656 | TMPRSS2 | 0.923076923077 | 3UTR |
| hsa-miR-1324 | NM_005656 | TMPRSS2 | 0.923076923077 | 3UTR |
| hsa-miR-320d | NM_005656 | TMPRSS2 | 0.923076923077 | 3UTR |
| hsa-miR-1908-5p | NM_005656 | TMPRSS2 | 0.923076923077 | 3UTR |
| hsa-miR-1909-3p | NM_005656 | TMPRSS2 | 0.923076923077 | 3UTR |
| hsa-miR-1911-3p | NM_005656 | TMPRSS2 | 0.923076923077 | 3UTR |
| hsa-miR-1912-5p | NM_005656 | TMPRSS2 | 0.923076923077 | 3UTR |
| hsa-miR-2114-5p | NM_005656 | TMPRSS2 | 0.923076923077 | 3UTR |
| hsa-miR-2117 | NM_005656 | TMPRSS2 | 0.923076923077 | 3UTR |
| hsa-miR-2276-3p | NM_005656 | TMPRSS2 | 0.923076923077 | 3UTR |
| hsa-miR-2277-3p | NM_005656 | TMPRSS2 | 0.923076923077 | 3UTR |
| hsa-miR-2682-5p | NM_005656 | TMPRSS2 | 0.923076923077 | 3UTR |
| hsa-miR-711 | NM_005656 | TMPRSS2 | 0.923076923077 | 3UTR |
| hsa-miR-548s | NM_005656 | TMPRSS2 | 0.923076923077 | 3UTR |
| hsa-miR-3132 | NM_005656 | TMPRSS2 | 0.923076923077 | 3UTR |
| hsa-miR-378b | NM_005656 | TMPRSS2 | 0.923076923077 | 3UTR |
| hsa-miR-3138 | NM_005656 | TMPRSS2 | 0.923076923077 | 3UTR |
| hsa-miR-3141 | NM_005656 | TMPRSS2 | 0.923076923077 | 3UTR |
| hsa-miR-1273c | NM_005656 | TMPRSS2 | 0.923076923077 | 3UTR |
| hsa-miR-3150a-3p | NM_005656 | TMPRSS2 | 0.923076923077 | 3UTR |
| hsa-miR-3152-3p | NM_005656 | TMPRSS2 | 0.923076923077 | 3UTR |
| hsa-miR-3153 | NM_005656 | TMPRSS2 | 0.923076923077 | 3UTR |
| hsa-miR-3154 | NM_005656 | TMPRSS2 | 0.923076923077 | 3UTR |
| hsa-miR-3158-5p | NM_005656 | TMPRSS2 | 0.923076923077 | 3UTR |
| hsa-miR-3162-3p | NM_005656 | TMPRSS2 | 0.923076923077 | 3UTR |
| hsa-miR-1260b | NM_005656 | TMPRSS2 | 0.923076923077 | 3UTR |
| hsa-miR-3168 | NM_005656 | TMPRSS2 | 0.923076923077 | 3UTR |
| hsa-miR-3173-5p | NM_005656 | TMPRSS2 | 0.923076923077 | 3UTR |
| hsa-miR-3173-5p | NM_005656 | TMPRSS2 | 0.923076923077 | 3UTR |
| hsa-miR-3173-3p | NM_005656 | TMPRSS2 | 0.923076923077 | 3UTR |
| hsa-miR-3179 | NM_005656 | TMPRSS2 | 0.923076923077 | 3UTR |
| hsa-miR-3180-5p | NM_005656 | TMPRSS2 | 0.923076923077 | 3UTR |
| hsa-miR-3184-5p | NM_005656 | TMPRSS2 | 0.923076923077 | 3UTR |
| hsa-miR-3184-5p | NM_005656 | TMPRSS2 | 0.923076923077 | 3UTR |
| hsa-miR-3185 | NM_005656 | TMPRSS2 | 0.923076923077 | 3UTR |
| hsa-miR-3189-5p | NM_005656 | TMPRSS2 | 0.923076923077 | 3UTR |
| hsa-miR-3189-3p | NM_005656 | TMPRSS2 | 0.923076923077 | 3UTR |
| hsa-miR-3190-3p | NM_005656 | TMPRSS2 | 0.923076923077 | 3UTR |
| hsa-miR-3194-5p | NM_005656 | TMPRSS2 | 0.923076923077 | 3UTR |
| hsa-miR-3197 | NM_005656 | TMPRSS2 | 0.923076923077 | 3UTR |
| hsa-miR-3198 | NM_005656 | TMPRSS2 | 0.923076923077 | 3UTR |
| hsa-miR-3202 | NM_005656 | TMPRSS2 | 0.923076923077 | 3UTR |
| hsa-miR-4301 | NM_005656 | TMPRSS2 | 0.923076923077 | 3UTR |
| hsa-miR-4301 | NM_005656 | TMPRSS2 | 0.923076923077 | 3UTR |
| hsa-miR-4299 | NM_005656 | TMPRSS2 | 0.923076923077 | 3UTR |
| hsa-miR-4300 | NM_005656 | TMPRSS2 | 0.923076923077 | 3UTR |

| | | | | |
|---|---|---|---|---|
| hsa-miR-4304 | NM_005656 | TMPRSS2 | 0.923076923077 | 3UTR |
| hsa-miR-4306 | NM_005656 | TMPRSS2 | 0.923076923077 | 3UTR |
| hsa-miR-4316 | NM_005656 | TMPRSS2 | 0.923076923077 | 3UTR |
| hsa-miR-4318 | NM_005656 | TMPRSS2 | 0.923076923077 | 3UTR |
| hsa-miR-4261 | NM_005656 | TMPRSS2 | 0.923076923077 | 3UTR |
| hsa-miR-4266 | NM_005656 | TMPRSS2 | 0.923076923077 | 3UTR |
| hsa-miR-4270 | NM_005656 | TMPRSS2 | 0.923076923077 | 3UTR |
| hsa-miR-4278 | NM_005656 | TMPRSS2 | 0.923076923077 | 3UTR |
| hsa-miR-4280 | NM_005656 | TMPRSS2 | 0.923076923077 | 3UTR |
| hsa-miR-4283 | NM_005656 | TMPRSS2 | 0.923076923077 | 3UTR |
| hsa-miR-4283 | NM_005656 | TMPRSS2 | 0.923076923077 | 3UTR |
| hsa-miR-500b-5p | NM_005656 | TMPRSS2 | 0.923076923077 | 3UTR |
| hsa-miR-3612 | NM_005656 | TMPRSS2 | 0.923076923077 | 3UTR |
| hsa-miR-3617-3p | NM_005656 | TMPRSS2 | 0.923076923077 | 3UTR |
| hsa-miR-3617-3p | NM_005656 | TMPRSS2 | 0.923076923077 | 3UTR |
| hsa-miR-3619-3p | NM_005656 | TMPRSS2 | 0.923076923077 | 3UTR |
| hsa-miR-3649 | NM_005656 | TMPRSS2 | 0.923076923077 | 3UTR |
| hsa-miR-3661 | NM_005656 | TMPRSS2 | 0.923076923077 | 3UTR |
| hsa-miR-3662 | NM_005656 | TMPRSS2 | 0.923076923077 | 3UTR |
| hsa-miR-3665 | NM_005656 | TMPRSS2 | 0.923076923077 | 3UTR |
| hsa-miR-3667-3p | NM_005656 | TMPRSS2 | 0.923076923077 | 3UTR |
| hsa-miR-3677-3p | NM_005656 | TMPRSS2 | 0.923076923077 | 3UTR |
| hsa-miR-3678-3p | NM_005656 | TMPRSS2 | 0.923076923077 | 3UTR |
| hsa-miR-3680-3p | NM_005656 | TMPRSS2 | 0.923076923077 | 3UTR |
| hsa-miR-3680-3p | NM_005656 | TMPRSS2 | 0.923076923077 | 3UTR |
| hsa-miR-3682-3p | NM_005656 | TMPRSS2 | 0.923076923077 | 3UTR |
| hsa-miR-3689a-5p | NM_005656 | TMPRSS2 | 0.923076923077 | 3UTR |
| hsa-miR-3692-5p | NM_005656 | TMPRSS2 | 0.923076923077 | 3UTR |
| hsa-miR-3689b-5p | NM_005656 | TMPRSS2 | 0.923076923077 | 3UTR |
| hsa-miR-3689b-3p | NM_005656 | TMPRSS2 | 0.923076923077 | 3UTR |
| hsa-miR-3913-3p | NM_005656 | TMPRSS2 | 0.923076923077 | 3UTR |
| hsa-miR-3919 | NM_005656 | TMPRSS2 | 0.923076923077 | 3UTR |
| hsa-miR-3923 | NM_005656 | TMPRSS2 | 0.923076923077 | 3UTR |
| hsa-miR-3934-3p | NM_005656 | TMPRSS2 | 0.923076923077 | 3UTR |
| hsa-miR-3939 | NM_005656 | TMPRSS2 | 0.923076923077 | 3UTR |
| hsa-miR-3940-5p | NM_005656 | TMPRSS2 | 0.923076923077 | 3UTR |
| hsa-miR-3945 | NM_005656 | TMPRSS2 | 0.923076923077 | 3UTR |
| hsa-miR-550b-2-5p | NM_005656 | TMPRSS2 | 0.923076923077 | 3UTR |
| hsa-miR-378g | NM_005656 | TMPRSS2 | 0.923076923077 | 3UTR |
| hsa-miR-4429 | NM_005656 | TMPRSS2 | 0.923076923077 | 3UTR |
| hsa-miR-4433a-5p | NM_005656 | TMPRSS2 | 0.923076923077 | 3UTR |
| hsa-miR-4442 | NM_005656 | TMPRSS2 | 0.923076923077 | 3UTR |
| hsa-miR-4448 | NM_005656 | TMPRSS2 | 0.923076923077 | 3UTR |
| hsa-miR-4451 | NM_005656 | TMPRSS2 | 0.923076923077 | 3UTR |
| hsa-miR-4458 | NM_005656 | TMPRSS2 | 0.923076923077 | 3UTR |
| hsa-miR-4465 | NM_005656 | TMPRSS2 | 0.923076923077 | 3UTR |
| hsa-miR-4466 | NM_005656 | TMPRSS2 | 0.923076923077 | 3UTR |

| | | | | |
|---|---|---|---|---|
| hsa-miR-4470 | NM_005656 | TMPRSS2 | 0.923076923077 | 3UTR |
| hsa-miR-4471 | NM_005656 | TMPRSS2 | 0.923076923077 | 3UTR |
| hsa-miR-4474-3p | NM_005656 | TMPRSS2 | 0.923076923077 | 3UTR |
| hsa-miR-3689c | NM_005656 | TMPRSS2 | 0.923076923077 | 3UTR |
| hsa-miR-3689e | NM_005656 | TMPRSS2 | 0.923076923077 | 3UTR |
| hsa-miR-4480 | NM_005656 | TMPRSS2 | 0.923076923077 | 3UTR |
| hsa-miR-4482-3p | NM_005656 | TMPRSS2 | 0.923076923077 | 3UTR |
| hsa-miR-4487 | NM_005656 | TMPRSS2 | 0.923076923077 | 3UTR |
| hsa-miR-4494 | NM_005656 | TMPRSS2 | 0.923076923077 | 3UTR |
| hsa-miR-4515 | NM_005656 | TMPRSS2 | 0.923076923077 | 3UTR |
| hsa-miR-4519 | NM_005656 | TMPRSS2 | 0.923076923077 | 3UTR |
| hsa-miR-4530 | NM_005656 | TMPRSS2 | 0.923076923077 | 3UTR |
| hsa-miR-4533 | NM_005656 | TMPRSS2 | 0.923076923077 | 3UTR |
| hsa-miR-4534 | NM_005656 | TMPRSS2 | 0.923076923077 | 3UTR |
| hsa-miR-378i | NM_005656 | TMPRSS2 | 0.923076923077 | 3UTR |
| hsa-miR-4540 | NM_005656 | TMPRSS2 | 0.923076923077 | 3UTR |
| hsa-miR-3972 | NM_005656 | TMPRSS2 | 0.923076923077 | 3UTR |
| hsa-miR-3976 | NM_005656 | TMPRSS2 | 0.923076923077 | 3UTR |
| hsa-miR-3978 | NM_005656 | TMPRSS2 | 0.923076923077 | 3UTR |
| hsa-miR-4635 | NM_005656 | TMPRSS2 | 0.923076923077 | 3UTR |
| hsa-miR-4640-3p | NM_005656 | TMPRSS2 | 0.923076923077 | 3UTR |
| hsa-miR-4644 | NM_005656 | TMPRSS2 | 0.923076923077 | 3UTR |
| hsa-miR-4647 | NM_005656 | TMPRSS2 | 0.923076923077 | 3UTR |
| hsa-miR-4649-3p | NM_005656 | TMPRSS2 | 0.923076923077 | 3UTR |
| hsa-miR-4653-3p | NM_005656 | TMPRSS2 | 0.923076923077 | 3UTR |
| hsa-miR-4654 | NM_005656 | TMPRSS2 | 0.923076923077 | 3UTR |
| hsa-miR-4655-5p | NM_005656 | TMPRSS2 | 0.923076923077 | 3UTR |
| hsa-miR-4656 | NM_005656 | TMPRSS2 | 0.923076923077 | 3UTR |
| hsa-miR-4659a-5p | NM_005656 | TMPRSS2 | 0.923076923077 | 3UTR |
| hsa-miR-4663 | NM_005656 | TMPRSS2 | 0.923076923077 | 3UTR |
| hsa-miR-4667-5p | NM_005656 | TMPRSS2 | 0.923076923077 | 3UTR |
| hsa-miR-4667-5p | NM_005656 | TMPRSS2 | 0.923076923077 | 3UTR |
| hsa-miR-4682 | NM_005656 | TMPRSS2 | 0.923076923077 | 3UTR |
| hsa-miR-4687-5p | NM_005656 | TMPRSS2 | 0.923076923077 | 3UTR |
| hsa-miR-4687-3p | NM_005656 | TMPRSS2 | 0.923076923077 | 3UTR |
| hsa-miR-4689 | NM_005656 | TMPRSS2 | 0.923076923077 | 3UTR |
| hsa-miR-4690-5p | NM_005656 | TMPRSS2 | 0.923076923077 | 3UTR |
| hsa-miR-4695-3p | NM_005656 | TMPRSS2 | 0.923076923077 | 3UTR |
| hsa-miR-4704-5p | NM_005656 | TMPRSS2 | 0.923076923077 | 3UTR |
| hsa-miR-4704-5p | NM_005656 | TMPRSS2 | 0.923076923077 | 3UTR |
| hsa-miR-4706 | NM_005656 | TMPRSS2 | 0.923076923077 | 3UTR |
| hsa-miR-4708-3p | NM_005656 | TMPRSS2 | 0.923076923077 | 3UTR |
| hsa-miR-4709-3p | NM_005656 | TMPRSS2 | 0.923076923077 | 3UTR |
| hsa-miR-4710 | NM_005656 | TMPRSS2 | 0.923076923077 | 3UTR |
| hsa-miR-4713-5p | NM_005656 | TMPRSS2 | 0.923076923077 | 3UTR |
| hsa-miR-4713-3p | NM_005656 | TMPRSS2 | 0.923076923077 | 3UTR |
| hsa-miR-4715-3p | NM_005656 | TMPRSS2 | 0.923076923077 | 3UTR |

| | | | | |
|---|---|---|---|---|
| hsa-miR-4725-5p | NM_005656 | TMPRSS2 | 0.923076923077 | 3UTR |
| hsa-miR-4731-3p | NM_005656 | TMPRSS2 | 0.923076923077 | 3UTR |
| hsa-miR-4732-5p | NM_005656 | TMPRSS2 | 0.923076923077 | 3UTR |
| hsa-miR-4738-3p | NM_005656 | TMPRSS2 | 0.923076923077 | 3UTR |
| hsa-miR-4742-5p | NM_005656 | TMPRSS2 | 0.923076923077 | 3UTR |
| hsa-miR-4743-5p | NM_005656 | TMPRSS2 | 0.923076923077 | 3UTR |
| hsa-miR-4744 | NM_005656 | TMPRSS2 | 0.923076923077 | 3UTR |
| hsa-miR-4748 | NM_005656 | TMPRSS2 | 0.923076923077 | 3UTR |
| hsa-miR-4749-3p | NM_005656 | TMPRSS2 | 0.923076923077 | 3UTR |
| hsa-miR-4755-5p | NM_005656 | TMPRSS2 | 0.923076923077 | 3UTR |
| hsa-miR-4756-3p | NM_005656 | TMPRSS2 | 0.923076923077 | 3UTR |
| hsa-miR-4760-3p | NM_005656 | TMPRSS2 | 0.923076923077 | 3UTR |
| hsa-miR-4764-5p | NM_005656 | TMPRSS2 | 0.923076923077 | 3UTR |
| hsa-miR-4775 | NM_005656 | TMPRSS2 | 0.923076923077 | 3UTR |
| hsa-miR-4776-5p | NM_005656 | TMPRSS2 | 0.923076923077 | 3UTR |
| hsa-miR-4436b-3p | NM_005656 | TMPRSS2 | 0.923076923077 | 3UTR |
| hsa-miR-4785 | NM_005656 | TMPRSS2 | 0.923076923077 | 3UTR |
| hsa-miR-2467-3p | NM_005656 | TMPRSS2 | 0.923076923077 | 3UTR |
| hsa-miR-4786-3p | NM_005656 | TMPRSS2 | 0.923076923077 | 3UTR |
| hsa-miR-4790-5p | NM_005656 | TMPRSS2 | 0.923076923077 | 3UTR |
| hsa-miR-4793-5p | NM_005656 | TMPRSS2 | 0.923076923077 | 3UTR |
| hsa-miR-4796-3p | NM_005656 | TMPRSS2 | 0.923076923077 | 3UTR |
| hsa-miR-4797-3p | NM_005656 | TMPRSS2 | 0.923076923077 | 3UTR |
| hsa-miR-4799-3p | NM_005656 | TMPRSS2 | 0.923076923077 | 3UTR |
| hsa-miR-4799-3p | NM_005656 | TMPRSS2 | 0.923076923077 | 3UTR |
| hsa-miR-4800-5p | NM_005656 | TMPRSS2 | 0.923076923077 | 3UTR |
| hsa-miR-5004-5p | NM_005656 | TMPRSS2 | 0.923076923077 | 3UTR |
| hsa-miR-5004-3p | NM_005656 | TMPRSS2 | 0.923076923077 | 3UTR |
| hsa-miR-548ao-3p | NM_005656 | TMPRSS2 | 0.923076923077 | 3UTR |
| hsa-miR-5006-5p | NM_005656 | TMPRSS2 | 0.923076923077 | 3UTR |
| hsa-miR-5006-5p | NM_005656 | TMPRSS2 | 0.923076923077 | 3UTR |
| hsa-miR-5008-3p | NM_005656 | TMPRSS2 | 0.923076923077 | 3UTR |
| hsa-miR-5010-5p | NM_005656 | TMPRSS2 | 0.923076923077 | 3UTR |
| hsa-miR-5047 | NM_005656 | TMPRSS2 | 0.923076923077 | 3UTR |
| hsa-miR-5088-5p | NM_005656 | TMPRSS2 | 0.923076923077 | 3UTR |
| hsa-miR-5088-3p | NM_005656 | TMPRSS2 | 0.923076923077 | 3UTR |
| hsa-miR-5090 | NM_005656 | TMPRSS2 | 0.923076923077 | 3UTR |
| hsa-miR-5092 | NM_005656 | TMPRSS2 | 0.923076923077 | 3UTR |
| hsa-miR-5187-3p | NM_005656 | TMPRSS2 | 0.923076923077 | 3UTR |
| hsa-miR-5187-3p | NM_005656 | TMPRSS2 | 0.923076923077 | 3UTR |
| hsa-miR-5190 | NM_005656 | TMPRSS2 | 0.923076923077 | 3UTR |
| hsa-miR-5192 | NM_005656 | TMPRSS2 | 0.923076923077 | 3UTR |
| hsa-miR-5192 | NM_005656 | TMPRSS2 | 0.923076923077 | 3UTR |
| hsa-miR-5193 | NM_005656 | TMPRSS2 | 0.923076923077 | 3UTR |
| hsa-miR-5193 | NM_005656 | TMPRSS2 | 0.923076923077 | 3UTR |
| hsa-miR-5193 | NM_005656 | TMPRSS2 | 0.923076923077 | 3UTR |
| hsa-miR-5196-5p | NM_005656 | TMPRSS2 | 0.923076923077 | 3UTR |

| | | | | |
|---|---|---|---|---|
| hsa-miR-5197-3p | NM_005656 | TMPRSS2 | 0.923076923077 | 3UTR |
| hsa-miR-5571-3p | NM_005656 | TMPRSS2 | 0.923076923077 | 3UTR |
| hsa-miR-5580-3p | NM_005656 | TMPRSS2 | 0.923076923077 | 3UTR |
| hsa-miR-5584-3p | NM_005656 | TMPRSS2 | 0.923076923077 | 3UTR |
| hsa-miR-5587-5p | NM_005656 | TMPRSS2 | 0.923076923077 | 3UTR |
| hsa-miR-1295b-5p | NM_005656 | TMPRSS2 | 0.923076923077 | 3UTR |
| hsa-miR-1295b-3p | NM_005656 | TMPRSS2 | 0.923076923077 | 3UTR |
| hsa-miR-5589-5p | NM_005656 | TMPRSS2 | 0.923076923077 | 3UTR |
| hsa-miR-5591-5p | NM_005656 | TMPRSS2 | 0.923076923077 | 3UTR |
| hsa-miR-5591-5p | NM_005656 | TMPRSS2 | 0.923076923077 | 3UTR |
| hsa-miR-5591-3p | NM_005656 | TMPRSS2 | 0.923076923077 | 3UTR |
| hsa-miR-5682 | NM_005656 | TMPRSS2 | 0.923076923077 | 3UTR |
| hsa-miR-5690 | NM_005656 | TMPRSS2 | 0.923076923077 | 3UTR |
| hsa-miR-5693 | NM_005656 | TMPRSS2 | 0.923076923077 | 3UTR |
| hsa-miR-5704 | NM_005656 | TMPRSS2 | 0.923076923077 | 3UTR |
| hsa-miR-6068 | NM_005656 | TMPRSS2 | 0.923076923077 | 3UTR |
| hsa-miR-6077 | NM_005656 | TMPRSS2 | 0.923076923077 | 3UTR |
| hsa-miR-6085 | NM_005656 | TMPRSS2 | 0.923076923077 | 3UTR |
| hsa-miR-6127 | NM_005656 | TMPRSS2 | 0.923076923077 | 3UTR |
| hsa-miR-6127 | NM_005656 | TMPRSS2 | 0.923076923077 | 3UTR |
| hsa-miR-378j | NM_005656 | TMPRSS2 | 0.923076923077 | 3UTR |
| hsa-miR-6134 | NM_005656 | TMPRSS2 | 0.923076923077 | 3UTR |
| hsa-miR-548ay-3p | NM_005656 | TMPRSS2 | 0.923076923077 | 3UTR |
| hsa-miR-6500-5p | NM_005656 | TMPRSS2 | 0.923076923077 | 3UTR |
| hsa-miR-6503-3p | NM_005656 | TMPRSS2 | 0.923076923077 | 3UTR |
| hsa-miR-6503-3p | NM_005656 | TMPRSS2 | 0.923076923077 | 3UTR |
| hsa-miR-6509-5p | NM_005656 | TMPRSS2 | 0.923076923077 | 3UTR |
| hsa-miR-6509-3p | NM_005656 | TMPRSS2 | 0.923076923077 | 3UTR |
| hsa-miR-6511a-5p | NM_005656 | TMPRSS2 | 0.923076923077 | 3UTR |
| hsa-miR-6515-5p | NM_005656 | TMPRSS2 | 0.923076923077 | 3UTR |
| hsa-miR-6716-3p | NM_005656 | TMPRSS2 | 0.923076923077 | 3UTR |
| hsa-miR-6511b-5p | NM_005656 | TMPRSS2 | 0.923076923077 | 3UTR |
| hsa-miR-6720-3p | NM_005656 | TMPRSS2 | 0.923076923077 | 3UTR |
| hsa-miR-6728-5p | NM_005656 | TMPRSS2 | 0.923076923077 | 3UTR |
| hsa-miR-6729-5p | NM_005656 | TMPRSS2 | 0.923076923077 | 3UTR |
| hsa-miR-6731-5p | NM_005656 | TMPRSS2 | 0.923076923077 | 3UTR |
| hsa-miR-6734-5p | NM_005656 | TMPRSS2 | 0.923076923077 | 3UTR |
| hsa-miR-6734-5p | NM_005656 | TMPRSS2 | 0.923076923077 | 3UTR |
| hsa-miR-6735-5p | NM_005656 | TMPRSS2 | 0.923076923077 | 3UTR |
| hsa-miR-6735-3p | NM_005656 | TMPRSS2 | 0.923076923077 | 3UTR |
| hsa-miR-6736-5p | NM_005656 | TMPRSS2 | 0.923076923077 | 3UTR |
| hsa-miR-6736-3p | NM_005656 | TMPRSS2 | 0.923076923077 | 3UTR |
| hsa-miR-6738-5p | NM_005656 | TMPRSS2 | 0.923076923077 | 3UTR |
| hsa-miR-6740-5p | NM_005656 | TMPRSS2 | 0.923076923077 | 3UTR |
| hsa-miR-6742-5p | NM_005656 | TMPRSS2 | 0.923076923077 | 3UTR |
| hsa-miR-6747-5p | NM_005656 | TMPRSS2 | 0.923076923077 | 3UTR |
| hsa-miR-6748-5p | NM_005656 | TMPRSS2 | 0.923076923077 | 3UTR |

| | | | | |
|---|---|---|---|---|
| hsa-miR-6751-3p | NM_005656 | TMPRSS2 | 0.923076923077 | 3UTR |
| hsa-miR-6753-5p | NM_005656 | TMPRSS2 | 0.923076923077 | 3UTR |
| hsa-miR-6754-5p | NM_005656 | TMPRSS2 | 0.923076923077 | 3UTR |
| hsa-miR-6758-5p | NM_005656 | TMPRSS2 | 0.923076923077 | 3UTR |
| hsa-miR-6763-5p | NM_005656 | TMPRSS2 | 0.923076923077 | 3UTR |
| hsa-miR-6763-5p | NM_005656 | TMPRSS2 | 0.923076923077 | 3UTR |
| hsa-miR-6765-5p | NM_005656 | TMPRSS2 | 0.923076923077 | 3UTR |
| hsa-miR-6765-3p | NM_005656 | TMPRSS2 | 0.923076923077 | 3UTR |
| hsa-miR-6766-3p | NM_005656 | TMPRSS2 | 0.923076923077 | 3UTR |
| hsa-miR-6767-3p | NM_005656 | TMPRSS2 | 0.923076923077 | 3UTR |
| hsa-miR-6768-3p | NM_005656 | TMPRSS2 | 0.923076923077 | 3UTR |
| hsa-miR-6769a-5p | NM_005656 | TMPRSS2 | 0.923076923077 | 3UTR |
| hsa-miR-6773-3p | NM_005656 | TMPRSS2 | 0.923076923077 | 3UTR |
| hsa-miR-6774-5p | NM_005656 | TMPRSS2 | 0.923076923077 | 3UTR |
| hsa-miR-6774-3p | NM_005656 | TMPRSS2 | 0.923076923077 | 3UTR |
| hsa-miR-6780a-3p | NM_005656 | TMPRSS2 | 0.923076923077 | 3UTR |
| hsa-miR-6781-5p | NM_005656 | TMPRSS2 | 0.923076923077 | 3UTR |
| hsa-miR-6787-3p | NM_005656 | TMPRSS2 | 0.923076923077 | 3UTR |
| hsa-miR-6791-5p | NM_005656 | TMPRSS2 | 0.923076923077 | 3UTR |
| hsa-miR-6792-5p | NM_005656 | TMPRSS2 | 0.923076923077 | 3UTR |
| hsa-miR-6793-3p | NM_005656 | TMPRSS2 | 0.923076923077 | 3UTR |
| hsa-miR-6799-5p | NM_005656 | TMPRSS2 | 0.923076923077 | 3UTR |
| hsa-miR-6799-3p | NM_005656 | TMPRSS2 | 0.923076923077 | 3UTR |
| hsa-miR-6800-3p | NM_005656 | TMPRSS2 | 0.923076923077 | 3UTR |
| hsa-miR-6801-5p | NM_005656 | TMPRSS2 | 0.923076923077 | 3UTR |
| hsa-miR-6801-3p | NM_005656 | TMPRSS2 | 0.923076923077 | 3UTR |
| hsa-miR-6802-5p | NM_005656 | TMPRSS2 | 0.923076923077 | 3UTR |
| hsa-miR-6804-5p | NM_005656 | TMPRSS2 | 0.923076923077 | 3UTR |
| hsa-miR-6804-5p | NM_005656 | TMPRSS2 | 0.923076923077 | 3UTR |
| hsa-miR-6806-5p | NM_005656 | TMPRSS2 | 0.923076923077 | 3UTR |
| hsa-miR-6811-5p | NM_005656 | TMPRSS2 | 0.923076923077 | 3UTR |
| hsa-miR-6812-3p | NM_005656 | TMPRSS2 | 0.923076923077 | 3UTR |
| hsa-miR-6812-3p | NM_005656 | TMPRSS2 | 0.923076923077 | 3UTR |
| hsa-miR-6813-5p | NM_005656 | TMPRSS2 | 0.923076923077 | 3UTR |
| hsa-miR-6815-5p | NM_005656 | TMPRSS2 | 0.923076923077 | 3UTR |
| hsa-miR-6815-3p | NM_005656 | TMPRSS2 | 0.923076923077 | 3UTR |
| hsa-miR-6815-3p | NM_005656 | TMPRSS2 | 0.923076923077 | 3UTR |
| hsa-miR-6816-5p | NM_005656 | TMPRSS2 | 0.923076923077 | 3UTR |
| hsa-miR-6821-3p | NM_005656 | TMPRSS2 | 0.923076923077 | 3UTR |
| hsa-miR-6823-3p | NM_005656 | TMPRSS2 | 0.923076923077 | 3UTR |
| hsa-miR-6826-5p | NM_005656 | TMPRSS2 | 0.923076923077 | 3UTR |
| hsa-miR-6828-5p | NM_005656 | TMPRSS2 | 0.923076923077 | 3UTR |
| hsa-miR-6829-5p | NM_005656 | TMPRSS2 | 0.923076923077 | 3UTR |
| hsa-miR-6829-3p | NM_005656 | TMPRSS2 | 0.923076923077 | 3UTR |
| hsa-miR-6833-3p | NM_005656 | TMPRSS2 | 0.923076923077 | 3UTR |
| hsa-miR-6834-3p | NM_005656 | TMPRSS2 | 0.923076923077 | 3UTR |
| hsa-miR-6838-3p | NM_005656 | TMPRSS2 | 0.923076923077 | 3UTR |

| | | | | |
|---|---|---|---|---|
| hsa-miR-6846-5p | NM_005656 | TMPRSS2 | 0.923076923077 | 3UTR |
| hsa-miR-6848-5p | NM_005656 | TMPRSS2 | 0.923076923077 | 3UTR |
| hsa-miR-6848-3p | NM_005656 | TMPRSS2 | 0.923076923077 | 3UTR |
| hsa-miR-6849-3p | NM_005656 | TMPRSS2 | 0.923076923077 | 3UTR |
| hsa-miR-6854-5p | NM_005656 | TMPRSS2 | 0.923076923077 | 3UTR |
| hsa-miR-6856-5p | NM_005656 | TMPRSS2 | 0.923076923077 | 3UTR |
| hsa-miR-6857-3p | NM_005656 | TMPRSS2 | 0.923076923077 | 3UTR |
| hsa-miR-6859-5p | NM_005656 | TMPRSS2 | 0.923076923077 | 3UTR |
| hsa-miR-6859-3p | NM_005656 | TMPRSS2 | 0.923076923077 | 3UTR |
| hsa-miR-6861-5p | NM_005656 | TMPRSS2 | 0.923076923077 | 3UTR |
| hsa-miR-6864-3p | NM_005656 | TMPRSS2 | 0.923076923077 | 3UTR |
| hsa-miR-6865-5p | NM_005656 | TMPRSS2 | 0.923076923077 | 3UTR |
| hsa-miR-6865-3p | NM_005656 | TMPRSS2 | 0.923076923077 | 3UTR |
| hsa-miR-6866-3p | NM_005656 | TMPRSS2 | 0.923076923077 | 3UTR |
| hsa-miR-6867-3p | NM_005656 | TMPRSS2 | 0.923076923077 | 3UTR |
| hsa-miR-6867-3p | NM_005656 | TMPRSS2 | 0.923076923077 | 3UTR |
| hsa-miR-6873-5p | NM_005656 | TMPRSS2 | 0.923076923077 | 3UTR |
| hsa-miR-6876-5p | NM_005656 | TMPRSS2 | 0.923076923077 | 3UTR |
| hsa-miR-6877-5p | NM_005656 | TMPRSS2 | 0.923076923077 | 3UTR |
| hsa-miR-6877-3p | NM_005656 | TMPRSS2 | 0.923076923077 | 3UTR |
| hsa-miR-6878-5p | NM_005656 | TMPRSS2 | 0.923076923077 | 3UTR |
| hsa-miR-6878-3p | NM_005656 | TMPRSS2 | 0.923076923077 | 3UTR |
| hsa-miR-6880-5p | NM_005656 | TMPRSS2 | 0.923076923077 | 3UTR |
| hsa-miR-6882-5p | NM_005656 | TMPRSS2 | 0.923076923077 | 3UTR |
| hsa-miR-6884-5p | NM_005656 | TMPRSS2 | 0.923076923077 | 3UTR |
| hsa-miR-6887-5p | NM_005656 | TMPRSS2 | 0.923076923077 | 3UTR |
| hsa-miR-6887-5p | NM_005656 | TMPRSS2 | 0.923076923077 | 3UTR |
| hsa-miR-6889-3p | NM_005656 | TMPRSS2 | 0.923076923077 | 3UTR |
| hsa-miR-6891-5p | NM_005656 | TMPRSS2 | 0.923076923077 | 3UTR |
| hsa-miR-6892-5p | NM_005656 | TMPRSS2 | 0.923076923077 | 3UTR |
| hsa-miR-6892-5p | NM_005656 | TMPRSS2 | 0.923076923077 | 3UTR |
| hsa-miR-6892-3p | NM_005656 | TMPRSS2 | 0.923076923077 | 3UTR |
| hsa-miR-6894-3p | NM_005656 | TMPRSS2 | 0.923076923077 | 3UTR |
| hsa-miR-7106-5p | NM_005656 | TMPRSS2 | 0.923076923077 | 3UTR |
| hsa-miR-7108-5p | NM_005656 | TMPRSS2 | 0.923076923077 | 3UTR |
| hsa-miR-7108-3p | NM_005656 | TMPRSS2 | 0.923076923077 | 3UTR |
| hsa-miR-7110-5p | NM_005656 | TMPRSS2 | 0.923076923077 | 3UTR |
| hsa-miR-7111-5p | NM_005656 | TMPRSS2 | 0.923076923077 | 3UTR |
| hsa-miR-7111-5p | NM_005656 | TMPRSS2 | 0.923076923077 | 3UTR |
| hsa-miR-7111-3p | NM_005656 | TMPRSS2 | 0.923076923077 | 3UTR |
| hsa-miR-7111-3p | NM_005656 | TMPRSS2 | 0.923076923077 | 3UTR |
| hsa-miR-7112-3p | NM_005656 | TMPRSS2 | 0.923076923077 | 3UTR |
| hsa-miR-7113-5p | NM_005656 | TMPRSS2 | 0.923076923077 | 3UTR |
| hsa-miR-7152-5p | NM_005656 | TMPRSS2 | 0.923076923077 | 3UTR |
| hsa-miR-7152-3p | NM_005656 | TMPRSS2 | 0.923076923077 | 3UTR |
| hsa-miR-7152-3p | NM_005656 | TMPRSS2 | 0.923076923077 | 3UTR |
| hsa-miR-7161-3p | NM_005656 | TMPRSS2 | 0.923076923077 | 3UTR |

| | | | | |
|---|---|---|---|---|
| hsa-miR-7162-3p | NM_005656 | TMPRSS2 | 0.923076923077 | 3UTR |
| hsa-miR-7706 | NM_005656 | TMPRSS2 | 0.923076923077 | 3UTR |
| hsa-miR-7843-5p | NM_005656 | TMPRSS2 | 0.923076923077 | 3UTR |
| hsa-miR-4433b-5p | NM_005656 | TMPRSS2 | 0.923076923077 | 3UTR |
| hsa-miR-4433b-3p | NM_005656 | TMPRSS2 | 0.923076923077 | 3UTR |
| hsa-miR-4433b-3p | NM_005656 | TMPRSS2 | 0.923076923077 | 3UTR |
| hsa-miR-1273h-3p | NM_005656 | TMPRSS2 | 0.923076923077 | 3UTR |
| hsa-miR-7851-3p | NM_005656 | TMPRSS2 | 0.923076923077 | 3UTR |
| hsa-miR-7851-3p | NM_005656 | TMPRSS2 | 0.923076923077 | 3UTR |
| hsa-miR-7854-3p | NM_005656 | TMPRSS2 | 0.923076923077 | 3UTR |
| hsa-miR-7974 | NM_005656 | TMPRSS2 | 0.923076923077 | 3UTR |
| hsa-miR-7978 | NM_005656 | TMPRSS2 | 0.923076923077 | 3UTR |
| hsa-miR-8072 | NM_005656 | TMPRSS2 | 0.923076923077 | 3UTR |
| hsa-miR-8077 | NM_005656 | TMPRSS2 | 0.923076923077 | 3UTR |
| hsa-miR-8080 | NM_005656 | TMPRSS2 | 0.923076923077 | 3UTR |
| hsa-miR-8085 | NM_005656 | TMPRSS2 | 0.923076923077 | 3UTR |
| hsa-miR-8086 | NM_005656 | TMPRSS2 | 0.923076923077 | 3UTR |
| hsa-miR-10392-3p | NM_005656 | TMPRSS2 | 0.923076923077 | 3UTR |
| hsa-miR-10397-3p | NM_005656 | TMPRSS2 | 0.923076923077 | 3UTR |
| hsa-miR-10401-5p | NM_005656 | TMPRSS2 | 0.923076923077 | 3UTR |
| hsa-miR-10522-5p | NM_005656 | TMPRSS2 | 0.923076923077 | 3UTR |
| hsa-miR-10524-5p | NM_005656 | TMPRSS2 | 0.923076923077 | 3UTR |
| hsa-miR-10526-3p | NM_005656 | TMPRSS2 | 0.923076923077 | 3UTR |
| hsa-miR-11181-5p | NM_005656 | TMPRSS2 | 0.923076923077 | 3UTR |
| hsa-miR-3059-5p | NM_005656 | TMPRSS2 | 0.923076923077 | 3UTR |
| hsa-miR-12126 | NM_005656 | TMPRSS2 | 0.923076923077 | 3UTR |
| hsa-miR-12127 | NM_005656 | TMPRSS2 | 0.923076923077 | 3UTR |
| hsa-miR-17-5p | NM_001135099 | TMPRSS2 | 0.923076923077 | 3UTR |
| hsa-miR-17-5p | NM_001135099 | TMPRSS2 | 0.923076923077 | 3UTR |
| hsa-miR-19b-3p | NM_001135099 | TMPRSS2 | 0.923076923077 | 3UTR |
| hsa-miR-20a-5p | NM_001135099 | TMPRSS2 | 0.923076923077 | 3UTR |
| hsa-miR-21-3p | NM_001135099 | TMPRSS2 | 0.923076923077 | 3UTR |
| hsa-miR-30a-3p | NM_001135099 | TMPRSS2 | 0.923076923077 | 3UTR |
| hsa-miR-96-3p | NM_001135099 | TMPRSS2 | 0.923076923077 | 3UTR |
| hsa-miR-103a-2-5p | NM_001135099 | TMPRSS2 | 0.923076923077 | 3UTR |
| hsa-miR-106a-5p | NM_001135099 | TMPRSS2 | 0.923076923077 | 3UTR |
| hsa-miR-106a-5p | NM_001135099 | TMPRSS2 | 0.923076923077 | 3UTR |
| hsa-miR-107 | NM_001135099 | TMPRSS2 | 0.923076923077 | 3UTR |
| hsa-miR-192-3p | NM_001135099 | TMPRSS2 | 0.923076923077 | 3UTR |
| hsa-miR-199a-3p | NM_001135099 | TMPRSS2 | 0.923076923077 | 3UTR |
| hsa-miR-129-5p | NM_001135099 | TMPRSS2 | 0.923076923077 | 3UTR |
| hsa-miR-30d-3p | NM_001135099 | TMPRSS2 | 0.923076923077 | 3UTR |
| hsa-miR-7-5p | NM_001135099 | TMPRSS2 | 0.923076923077 | 3UTR |
| hsa-miR-181b-5p | NM_001135099 | TMPRSS2 | 0.923076923077 | 3UTR |
| hsa-miR-181c-5p | NM_001135099 | TMPRSS2 | 0.923076923077 | 3UTR |
| hsa-miR-199b-3p | NM_001135099 | TMPRSS2 | 0.923076923077 | 3UTR |
| hsa-miR-216a-3p | NM_001135099 | TMPRSS2 | 0.923076923077 | 3UTR |

| | | | | |
|---|---|---|---|---|
| hsa-miR-218-1-3p | NM_001135099 | TMPRSS2 | 0.923076923077 | 3UTR |
| hsa-miR-222-3p | NM_001135099 | TMPRSS2 | 0.923076923077 | 3UTR |
| hsa-miR-224-5p | NM_001135099 | TMPRSS2 | 0.923076923077 | 3UTR |
| hsa-let-7g-3p | NM_001135099 | TMPRSS2 | 0.923076923077 | 3UTR |
| hsa-let-7i-3p | NM_001135099 | TMPRSS2 | 0.923076923077 | 3UTR |
| hsa-miR-15b-5p | NM_001135099 | TMPRSS2 | 0.923076923077 | 3UTR |
| hsa-miR-128-1-5p | NM_001135099 | TMPRSS2 | 0.923076923077 | 3UTR |
| hsa-miR-128-1-5p | NM_001135099 | TMPRSS2 | 0.923076923077 | 3UTR |
| hsa-miR-140-5p | NM_001135099 | TMPRSS2 | 0.923076923077 | 3UTR |
| hsa-miR-140-3p | NM_001135099 | TMPRSS2 | 0.923076923077 | 3UTR |
| hsa-miR-9-5p | NM_001135099 | TMPRSS2 | 0.923076923077 | 3UTR |
| hsa-miR-185-3p | NM_001135099 | TMPRSS2 | 0.923076923077 | 3UTR |
| hsa-miR-195-3p | NM_001135099 | TMPRSS2 | 0.923076923077 | 3UTR |
| hsa-miR-320a-3p | NM_001135099 | TMPRSS2 | 0.923076923077 | 3UTR |
| hsa-miR-194-3p | NM_001135099 | TMPRSS2 | 0.923076923077 | 3UTR |
| hsa-miR-106b-5p | NM_001135099 | TMPRSS2 | 0.923076923077 | 3UTR |
| hsa-miR-29c-5p | NM_001135099 | TMPRSS2 | 0.923076923077 | 3UTR |
| hsa-miR-34c-3p | NM_001135099 | TMPRSS2 | 0.923076923077 | 3UTR |
| hsa-miR-296-3p | NM_001135099 | TMPRSS2 | 0.923076923077 | 3UTR |
| hsa-miR-296-3p | NM_001135099 | TMPRSS2 | 0.923076923077 | 3UTR |
| hsa-miR-365b-5p | NM_001135099 | TMPRSS2 | 0.923076923077 | 3UTR |
| hsa-miR-302d-5p | NM_001135099 | TMPRSS2 | 0.923076923077 | 3UTR |
| hsa-miR-302d-5p | NM_001135099 | TMPRSS2 | 0.923076923077 | 3UTR |
| hsa-miR-367-3p | NM_001135099 | TMPRSS2 | 0.923076923077 | 3UTR |
| hsa-miR-370-5p | NM_001135099 | TMPRSS2 | 0.923076923077 | 3UTR |
| hsa-miR-370-3p | NM_001135099 | TMPRSS2 | 0.923076923077 | 3UTR |
| hsa-miR-371a-5p | NM_001135099 | TMPRSS2 | 0.923076923077 | 3UTR |
| hsa-miR-373-5p | NM_001135099 | TMPRSS2 | 0.923076923077 | 3UTR |
| hsa-miR-342-5p | NM_001135099 | TMPRSS2 | 0.923076923077 | 3UTR |
| hsa-miR-339-5p | NM_001135099 | TMPRSS2 | 0.923076923077 | 3UTR |
| hsa-miR-339-5p | NM_001135099 | TMPRSS2 | 0.923076923077 | 3UTR |
| hsa-miR-345-3p | NM_001135099 | TMPRSS2 | 0.923076923077 | 3UTR |
| hsa-miR-20b-5p | NM_001135099 | TMPRSS2 | 0.923076923077 | 3UTR |
| hsa-miR-448 | NM_001135099 | TMPRSS2 | 0.923076923077 | 3UTR |
| hsa-miR-449a | NM_001135099 | TMPRSS2 | 0.923076923077 | 3UTR |
| hsa-miR-449a | NM_001135099 | TMPRSS2 | 0.923076923077 | 3UTR |
| hsa-miR-431-3p | NM_001135099 | TMPRSS2 | 0.923076923077 | 3UTR |
| hsa-miR-490-5p | NM_001135099 | TMPRSS2 | 0.923076923077 | 3UTR |
| hsa-miR-490-3p | NM_001135099 | TMPRSS2 | 0.923076923077 | 3UTR |
| hsa-miR-146b-3p | NM_001135099 | TMPRSS2 | 0.923076923077 | 3UTR |
| hsa-miR-202-3p | NM_001135099 | TMPRSS2 | 0.923076923077 | 3UTR |
| hsa-miR-493-3p | NM_001135099 | TMPRSS2 | 0.923076923077 | 3UTR |
| hsa-miR-193b-3p | NM_001135099 | TMPRSS2 | 0.923076923077 | 3UTR |
| hsa-miR-512-5p | NM_001135099 | TMPRSS2 | 0.923076923077 | 3UTR |
| hsa-miR-520e-5p | NM_001135099 | TMPRSS2 | 0.923076923077 | 3UTR |
| hsa-miR-520f-3p | NM_001135099 | TMPRSS2 | 0.923076923077 | 3UTR |
| hsa-miR-519c-5p | NM_001135099 | TMPRSS2 | 0.923076923077 | 3UTR |

| | | | | |
|---|---|---|---|---|
| hsa-miR-520a-5p | NM_001135099 | TMPRSS2 | 0.923076923077 | 3UTR |
| hsa-miR-519b-5p | NM_001135099 | TMPRSS2 | 0.923076923077 | 3UTR |
| hsa-miR-523-5p | NM_001135099 | TMPRSS2 | 0.923076923077 | 3UTR |
| hsa-miR-518f-5p | NM_001135099 | TMPRSS2 | 0.923076923077 | 3UTR |
| hsa-miR-519d-5p | NM_001135099 | TMPRSS2 | 0.923076923077 | 3UTR |
| hsa-miR-516b-3p | NM_001135099 | TMPRSS2 | 0.923076923077 | 3UTR |
| hsa-miR-518e-5p | NM_001135099 | TMPRSS2 | 0.923076923077 | 3UTR |
| hsa-miR-518a-5p | NM_001135099 | TMPRSS2 | 0.923076923077 | 3UTR |
| hsa-miR-522-5p | NM_001135099 | TMPRSS2 | 0.923076923077 | 3UTR |
| hsa-miR-519a-5p | NM_001135099 | TMPRSS2 | 0.923076923077 | 3UTR |
| hsa-miR-527 | NM_001135099 | TMPRSS2 | 0.923076923077 | 3UTR |
| hsa-miR-516a-3p | NM_001135099 | TMPRSS2 | 0.923076923077 | 3UTR |
| hsa-miR-500a-3p | NM_001135099 | TMPRSS2 | 0.923076923077 | 3UTR |
| hsa-miR-450a-2-3p | NM_001135099 | TMPRSS2 | 0.923076923077 | 3UTR |
| hsa-miR-510-3p | NM_001135099 | TMPRSS2 | 0.923076923077 | 3UTR |
| hsa-miR-514a-5p | NM_001135099 | TMPRSS2 | 0.923076923077 | 3UTR |
| hsa-miR-532-3p | NM_001135099 | TMPRSS2 | 0.923076923077 | 3UTR |
| hsa-miR-557 | NM_001135099 | TMPRSS2 | 0.923076923077 | 3UTR |
| hsa-miR-548a-3p | NM_001135099 | TMPRSS2 | 0.923076923077 | 3UTR |
| hsa-miR-591 | NM_001135099 | TMPRSS2 | 0.923076923077 | 3UTR |
| hsa-miR-595 | NM_001135099 | TMPRSS2 | 0.923076923077 | 3UTR |
| hsa-miR-598-3p | NM_001135099 | TMPRSS2 | 0.923076923077 | 3UTR |
| hsa-miR-604 | NM_001135099 | TMPRSS2 | 0.923076923077 | 3UTR |
| hsa-miR-616-5p | NM_001135099 | TMPRSS2 | 0.923076923077 | 3UTR |
| hsa-miR-617 | NM_001135099 | TMPRSS2 | 0.923076923077 | 3UTR |
| hsa-miR-618 | NM_001135099 | TMPRSS2 | 0.923076923077 | 3UTR |
| hsa-miR-619-5p | NM_001135099 | TMPRSS2 | 0.923076923077 | 3UTR |
| hsa-miR-619-5p | NM_001135099 | TMPRSS2 | 0.923076923077 | 3UTR |
| hsa-miR-623 | NM_001135099 | TMPRSS2 | 0.923076923077 | 3UTR |
| hsa-miR-625-5p | NM_001135099 | TMPRSS2 | 0.923076923077 | 3UTR |
| hsa-miR-628-3p | NM_001135099 | TMPRSS2 | 0.923076923077 | 3UTR |
| hsa-miR-630 | NM_001135099 | TMPRSS2 | 0.923076923077 | 3UTR |
| hsa-miR-33b-3p | NM_001135099 | TMPRSS2 | 0.923076923077 | 3UTR |
| hsa-miR-639 | NM_001135099 | TMPRSS2 | 0.923076923077 | 3UTR |
| hsa-miR-647 | NM_001135099 | TMPRSS2 | 0.923076923077 | 3UTR |
| hsa-miR-648 | NM_001135099 | TMPRSS2 | 0.923076923077 | 3UTR |
| hsa-miR-650 | NM_001135099 | TMPRSS2 | 0.923076923077 | 3UTR |
| hsa-miR-650 | NM_001135099 | TMPRSS2 | 0.923076923077 | 3UTR |
| hsa-miR-652-3p | NM_001135099 | TMPRSS2 | 0.923076923077 | 3UTR |
| hsa-miR-652-3p | NM_001135099 | TMPRSS2 | 0.923076923077 | 3UTR |
| hsa-miR-449b-5p | NM_001135099 | TMPRSS2 | 0.923076923077 | 3UTR |
| hsa-miR-654-3p | NM_001135099 | TMPRSS2 | 0.923076923077 | 3UTR |
| hsa-miR-655-5p | NM_001135099 | TMPRSS2 | 0.923076923077 | 3UTR |
| hsa-miR-549a-5p | NM_001135099 | TMPRSS2 | 0.923076923077 | 3UTR |
| hsa-miR-659-3p | NM_001135099 | TMPRSS2 | 0.923076923077 | 3UTR |
| hsa-miR-542-5p | NM_001135099 | TMPRSS2 | 0.923076923077 | 3UTR |
| hsa-miR-671-5p | NM_001135099 | TMPRSS2 | 0.923076923077 | 3UTR |

| | | | | |
|---|---|---|---|---|
| hsa-miR-767-3p | NM_001135099 | TMPRSS2 | 0.923076923077 | 3UTR |
| hsa-miR-151b | NM_001135099 | TMPRSS2 | 0.923076923077 | 3UTR |
| hsa-miR-320b | NM_001135099 | TMPRSS2 | 0.923076923077 | 3UTR |
| hsa-miR-320c | NM_001135099 | TMPRSS2 | 0.923076923077 | 3UTR |
| hsa-miR-1323 | NM_001135099 | TMPRSS2 | 0.923076923077 | 3UTR |
| hsa-miR-1271-3p | NM_001135099 | TMPRSS2 | 0.923076923077 | 3UTR |
| hsa-miR-1301-5p | NM_001135099 | TMPRSS2 | 0.923076923077 | 3UTR |
| hsa-miR-378d | NM_001135099 | TMPRSS2 | 0.923076923077 | 3UTR |
| hsa-miR-770-5p | NM_001135099 | TMPRSS2 | 0.923076923077 | 3UTR |
| hsa-miR-675-3p | NM_001135099 | TMPRSS2 | 0.923076923077 | 3UTR |
| hsa-miR-885-5p | NM_001135099 | TMPRSS2 | 0.923076923077 | 3UTR |
| hsa-miR-873-5p | NM_001135099 | TMPRSS2 | 0.923076923077 | 3UTR |
| hsa-miR-921 | NM_001135099 | TMPRSS2 | 0.923076923077 | 3UTR |
| hsa-miR-921 | NM_001135099 | TMPRSS2 | 0.923076923077 | 3UTR |
| hsa-miR-933 | NM_001135099 | TMPRSS2 | 0.923076923077 | 3UTR |
| hsa-miR-939-5p | NM_001135099 | TMPRSS2 | 0.923076923077 | 3UTR |
| hsa-miR-1180-5p | NM_001135099 | TMPRSS2 | 0.923076923077 | 3UTR |
| hsa-miR-1183 | NM_001135099 | TMPRSS2 | 0.923076923077 | 3UTR |
| hsa-miR-1184 | NM_001135099 | TMPRSS2 | 0.923076923077 | 3UTR |
| hsa-miR-1229-3p | NM_001135099 | TMPRSS2 | 0.923076923077 | 3UTR |
| hsa-miR-1233-3p | NM_001135099 | TMPRSS2 | 0.923076923077 | 3UTR |
| hsa-miR-1236-5p | NM_001135099 | TMPRSS2 | 0.923076923077 | 3UTR |
| hsa-miR-1236-3p | NM_001135099 | TMPRSS2 | 0.923076923077 | 3UTR |
| hsa-miR-1236-3p | NM_001135099 | TMPRSS2 | 0.923076923077 | 3UTR |
| hsa-miR-1205 | NM_001135099 | TMPRSS2 | 0.923076923077 | 3UTR |
| hsa-miR-1207-5p | NM_001135099 | TMPRSS2 | 0.923076923077 | 3UTR |
| hsa-miR-1285-5p | NM_001135099 | TMPRSS2 | 0.923076923077 | 3UTR |
| hsa-miR-1286 | NM_001135099 | TMPRSS2 | 0.923076923077 | 3UTR |
| hsa-miR-1289 | NM_001135099 | TMPRSS2 | 0.923076923077 | 3UTR |
| hsa-miR-1291 | NM_001135099 | TMPRSS2 | 0.923076923077 | 3UTR |
| hsa-miR-1295a | NM_001135099 | TMPRSS2 | 0.923076923077 | 3UTR |
| hsa-miR-1245a | NM_001135099 | TMPRSS2 | 0.923076923077 | 3UTR |
| hsa-miR-1249-5p | NM_001135099 | TMPRSS2 | 0.923076923077 | 3UTR |
| hsa-miR-1249-3p | NM_001135099 | TMPRSS2 | 0.923076923077 | 3UTR |
| hsa-miR-1256 | NM_001135099 | TMPRSS2 | 0.923076923077 | 3UTR |
| hsa-miR-1258 | NM_001135099 | TMPRSS2 | 0.923076923077 | 3UTR |
| hsa-miR-1276 | NM_001135099 | TMPRSS2 | 0.923076923077 | 3UTR |
| hsa-miR-1306-5p | NM_001135099 | TMPRSS2 | 0.923076923077 | 3UTR |
| hsa-miR-1539 | NM_001135099 | TMPRSS2 | 0.923076923077 | 3UTR |
| hsa-miR-320d | NM_001135099 | TMPRSS2 | 0.923076923077 | 3UTR |
| hsa-miR-1912-5p | NM_001135099 | TMPRSS2 | 0.923076923077 | 3UTR |
| hsa-miR-1915-3p | NM_001135099 | TMPRSS2 | 0.923076923077 | 3UTR |
| hsa-miR-2277-3p | NM_001135099 | TMPRSS2 | 0.923076923077 | 3UTR |
| hsa-miR-2682-5p | NM_001135099 | TMPRSS2 | 0.923076923077 | 3UTR |
| hsa-miR-711 | NM_001135099 | TMPRSS2 | 0.923076923077 | 3UTR |
| hsa-miR-3117-3p | NM_001135099 | TMPRSS2 | 0.923076923077 | 3UTR |
| hsa-miR-3124-5p | NM_001135099 | TMPRSS2 | 0.923076923077 | 3UTR |

| | | | | |
|---|---|---|---|---|
| hsa-miR-3127-3p | NM_001135099 | TMPRSS2 | 0.923076923077 | 3UTR |
| hsa-miR-3129-3p | NM_001135099 | TMPRSS2 | 0.923076923077 | 3UTR |
| hsa-miR-378b | NM_001135099 | TMPRSS2 | 0.923076923077 | 3UTR |
| hsa-miR-3141 | NM_001135099 | TMPRSS2 | 0.923076923077 | 3UTR |
| hsa-miR-1273c | NM_001135099 | TMPRSS2 | 0.923076923077 | 3UTR |
| hsa-miR-3147 | NM_001135099 | TMPRSS2 | 0.923076923077 | 3UTR |
| hsa-miR-3152-3p | NM_001135099 | TMPRSS2 | 0.923076923077 | 3UTR |
| hsa-miR-3153 | NM_001135099 | TMPRSS2 | 0.923076923077 | 3UTR |
| hsa-miR-3154 | NM_001135099 | TMPRSS2 | 0.923076923077 | 3UTR |
| hsa-miR-3154 | NM_001135099 | TMPRSS2 | 0.923076923077 | 3UTR |
| hsa-miR-3154 | NM_001135099 | TMPRSS2 | 0.923076923077 | 3UTR |
| hsa-miR-3155a | NM_001135099 | TMPRSS2 | 0.923076923077 | 3UTR |
| hsa-miR-3156-5p | NM_001135099 | TMPRSS2 | 0.923076923077 | 3UTR |
| hsa-miR-3159 | NM_001135099 | TMPRSS2 | 0.923076923077 | 3UTR |
| hsa-miR-3162-3p | NM_001135099 | TMPRSS2 | 0.923076923077 | 3UTR |
| hsa-miR-3163 | NM_001135099 | TMPRSS2 | 0.923076923077 | 3UTR |
| hsa-miR-3168 | NM_001135099 | TMPRSS2 | 0.923076923077 | 3UTR |
| hsa-miR-3169 | NM_001135099 | TMPRSS2 | 0.923076923077 | 3UTR |
| hsa-miR-3173-5p | NM_001135099 | TMPRSS2 | 0.923076923077 | 3UTR |
| hsa-miR-3173-5p | NM_001135099 | TMPRSS2 | 0.923076923077 | 3UTR |
| hsa-miR-3173-3p | NM_001135099 | TMPRSS2 | 0.923076923077 | 3UTR |
| hsa-miR-3180-5p | NM_001135099 | TMPRSS2 | 0.923076923077 | 3UTR |
| hsa-miR-3184-5p | NM_001135099 | TMPRSS2 | 0.923076923077 | 3UTR |
| hsa-miR-3185 | NM_001135099 | TMPRSS2 | 0.923076923077 | 3UTR |
| hsa-miR-3189-3p | NM_001135099 | TMPRSS2 | 0.923076923077 | 3UTR |
| hsa-miR-3191-3p | NM_001135099 | TMPRSS2 | 0.923076923077 | 3UTR |
| hsa-miR-3191-3p | NM_001135099 | TMPRSS2 | 0.923076923077 | 3UTR |
| hsa-miR-3192-5p | NM_001135099 | TMPRSS2 | 0.923076923077 | 3UTR |
| hsa-miR-3194-5p | NM_001135099 | TMPRSS2 | 0.923076923077 | 3UTR |
| hsa-miR-3198 | NM_001135099 | TMPRSS2 | 0.923076923077 | 3UTR |
| hsa-miR-514b-5p | NM_001135099 | TMPRSS2 | 0.923076923077 | 3UTR |
| hsa-miR-3202 | NM_001135099 | TMPRSS2 | 0.923076923077 | 3UTR |
| hsa-miR-4299 | NM_001135099 | TMPRSS2 | 0.923076923077 | 3UTR |
| hsa-miR-4298 | NM_001135099 | TMPRSS2 | 0.923076923077 | 3UTR |
| hsa-miR-4298 | NM_001135099 | TMPRSS2 | 0.923076923077 | 3UTR |
| hsa-miR-4304 | NM_001135099 | TMPRSS2 | 0.923076923077 | 3UTR |
| hsa-miR-4316 | NM_001135099 | TMPRSS2 | 0.923076923077 | 3UTR |
| hsa-miR-4321 | NM_001135099 | TMPRSS2 | 0.923076923077 | 3UTR |
| hsa-miR-4257 | NM_001135099 | TMPRSS2 | 0.923076923077 | 3UTR |
| hsa-miR-4254 | NM_001135099 | TMPRSS2 | 0.923076923077 | 3UTR |
| hsa-miR-4252 | NM_001135099 | TMPRSS2 | 0.923076923077 | 3UTR |
| hsa-miR-4261 | NM_001135099 | TMPRSS2 | 0.923076923077 | 3UTR |
| hsa-miR-4270 | NM_001135099 | TMPRSS2 | 0.923076923077 | 3UTR |
| hsa-miR-4278 | NM_001135099 | TMPRSS2 | 0.923076923077 | 3UTR |
| hsa-miR-4280 | NM_001135099 | TMPRSS2 | 0.923076923077 | 3UTR |
| hsa-miR-4284 | NM_001135099 | TMPRSS2 | 0.923076923077 | 3UTR |
| hsa-miR-4290 | NM_001135099 | TMPRSS2 | 0.923076923077 | 3UTR |

| | | | | |
|---|---|---|---|---|
| hsa-miR-3612 | NM_001135099 | TMPRSS2 | 0.923076923077 | 3UTR |
| hsa-miR-3619-3p | NM_001135099 | TMPRSS2 | 0.923076923077 | 3UTR |
| hsa-miR-3621 | NM_001135099 | TMPRSS2 | 0.923076923077 | 3UTR |
| hsa-miR-3649 | NM_001135099 | TMPRSS2 | 0.923076923077 | 3UTR |
| hsa-miR-3652 | NM_001135099 | TMPRSS2 | 0.923076923077 | 3UTR |
| hsa-miR-3659 | NM_001135099 | TMPRSS2 | 0.923076923077 | 3UTR |
| hsa-miR-3661 | NM_001135099 | TMPRSS2 | 0.923076923077 | 3UTR |
| hsa-miR-3663-5p | NM_001135099 | TMPRSS2 | 0.923076923077 | 3UTR |
| hsa-miR-3665 | NM_001135099 | TMPRSS2 | 0.923076923077 | 3UTR |
| hsa-miR-3667-3p | NM_001135099 | TMPRSS2 | 0.923076923077 | 3UTR |
| hsa-miR-3677-3p | NM_001135099 | TMPRSS2 | 0.923076923077 | 3UTR |
| hsa-miR-3678-3p | NM_001135099 | TMPRSS2 | 0.923076923077 | 3UTR |
| hsa-miR-3679-5p | NM_001135099 | TMPRSS2 | 0.923076923077 | 3UTR |
| hsa-miR-3680-3p | NM_001135099 | TMPRSS2 | 0.923076923077 | 3UTR |
| hsa-miR-3689a-5p | NM_001135099 | TMPRSS2 | 0.923076923077 | 3UTR |
| hsa-miR-3692-5p | NM_001135099 | TMPRSS2 | 0.923076923077 | 3UTR |
| hsa-miR-3692-3p | NM_001135099 | TMPRSS2 | 0.923076923077 | 3UTR |
| hsa-miR-3689b-5p | NM_001135099 | TMPRSS2 | 0.923076923077 | 3UTR |
| hsa-miR-3908 | NM_001135099 | TMPRSS2 | 0.923076923077 | 3UTR |
| hsa-miR-3917 | NM_001135099 | TMPRSS2 | 0.923076923077 | 3UTR |
| hsa-miR-3919 | NM_001135099 | TMPRSS2 | 0.923076923077 | 3UTR |
| hsa-miR-3934-3p | NM_001135099 | TMPRSS2 | 0.923076923077 | 3UTR |
| hsa-miR-3935 | NM_001135099 | TMPRSS2 | 0.923076923077 | 3UTR |
| hsa-miR-3943 | NM_001135099 | TMPRSS2 | 0.923076923077 | 3UTR |
| hsa-miR-3945 | NM_001135099 | TMPRSS2 | 0.923076923077 | 3UTR |
| hsa-miR-4425 | NM_001135099 | TMPRSS2 | 0.923076923077 | 3UTR |
| hsa-miR-4428 | NM_001135099 | TMPRSS2 | 0.923076923077 | 3UTR |
| hsa-miR-4429 | NM_001135099 | TMPRSS2 | 0.923076923077 | 3UTR |
| hsa-miR-4442 | NM_001135099 | TMPRSS2 | 0.923076923077 | 3UTR |
| hsa-miR-4448 | NM_001135099 | TMPRSS2 | 0.923076923077 | 3UTR |
| hsa-miR-548ah-5p | NM_001135099 | TMPRSS2 | 0.923076923077 | 3UTR |
| hsa-miR-4451 | NM_001135099 | TMPRSS2 | 0.923076923077 | 3UTR |
| hsa-miR-4458 | NM_001135099 | TMPRSS2 | 0.923076923077 | 3UTR |
| hsa-miR-4466 | NM_001135099 | TMPRSS2 | 0.923076923077 | 3UTR |
| hsa-miR-4471 | NM_001135099 | TMPRSS2 | 0.923076923077 | 3UTR |
| hsa-miR-4476 | NM_001135099 | TMPRSS2 | 0.923076923077 | 3UTR |
| hsa-miR-3689e | NM_001135099 | TMPRSS2 | 0.923076923077 | 3UTR |
| hsa-miR-4480 | NM_001135099 | TMPRSS2 | 0.923076923077 | 3UTR |
| hsa-miR-4482-3p | NM_001135099 | TMPRSS2 | 0.923076923077 | 3UTR |
| hsa-miR-4494 | NM_001135099 | TMPRSS2 | 0.923076923077 | 3UTR |
| hsa-miR-4500 | NM_001135099 | TMPRSS2 | 0.923076923077 | 3UTR |
| hsa-miR-4515 | NM_001135099 | TMPRSS2 | 0.923076923077 | 3UTR |
| hsa-miR-4517 | NM_001135099 | TMPRSS2 | 0.923076923077 | 3UTR |
| hsa-miR-4524a-5p | NM_001135099 | TMPRSS2 | 0.923076923077 | 3UTR |
| hsa-miR-4530 | NM_001135099 | TMPRSS2 | 0.923076923077 | 3UTR |
| hsa-miR-4534 | NM_001135099 | TMPRSS2 | 0.923076923077 | 3UTR |
| hsa-miR-378i | NM_001135099 | TMPRSS2 | 0.923076923077 | 3UTR |

| | | | | |
|---|---|---|---|---|
| hsa-miR-378i | NM_001135099 | TMPRSS2 | 0.923076923077 | 3UTR |
| hsa-miR-4537 | NM_001135099 | TMPRSS2 | 0.923076923077 | 3UTR |
| hsa-miR-3978 | NM_001135099 | TMPRSS2 | 0.923076923077 | 3UTR |
| hsa-miR-4632-5p | NM_001135099 | TMPRSS2 | 0.923076923077 | 3UTR |
| hsa-miR-4640-3p | NM_001135099 | TMPRSS2 | 0.923076923077 | 3UTR |
| hsa-miR-4650-3p | NM_001135099 | TMPRSS2 | 0.923076923077 | 3UTR |
| hsa-miR-4651 | NM_001135099 | TMPRSS2 | 0.923076923077 | 3UTR |
| hsa-miR-4655-5p | NM_001135099 | TMPRSS2 | 0.923076923077 | 3UTR |
| hsa-miR-4656 | NM_001135099 | TMPRSS2 | 0.923076923077 | 3UTR |
| hsa-miR-4657 | NM_001135099 | TMPRSS2 | 0.923076923077 | 3UTR |
| hsa-miR-4661-3p | NM_001135099 | TMPRSS2 | 0.923076923077 | 3UTR |
| hsa-miR-4663 | NM_001135099 | TMPRSS2 | 0.923076923077 | 3UTR |
| hsa-miR-4663 | NM_001135099 | TMPRSS2 | 0.923076923077 | 3UTR |
| hsa-miR-4664-5p | NM_001135099 | TMPRSS2 | 0.923076923077 | 3UTR |
| hsa-miR-4667-5p | NM_001135099 | TMPRSS2 | 0.923076923077 | 3UTR |
| hsa-miR-4667-3p | NM_001135099 | TMPRSS2 | 0.923076923077 | 3UTR |
| hsa-miR-4687-5p | NM_001135099 | TMPRSS2 | 0.923076923077 | 3UTR |
| hsa-miR-4687-3p | NM_001135099 | TMPRSS2 | 0.923076923077 | 3UTR |
| hsa-miR-4689 | NM_001135099 | TMPRSS2 | 0.923076923077 | 3UTR |
| hsa-miR-4690-5p | NM_001135099 | TMPRSS2 | 0.923076923077 | 3UTR |
| hsa-miR-4691-3p | NM_001135099 | TMPRSS2 | 0.923076923077 | 3UTR |
| hsa-miR-4691-3p | NM_001135099 | TMPRSS2 | 0.923076923077 | 3UTR |
| hsa-miR-4695-5p | NM_001135099 | TMPRSS2 | 0.923076923077 | 3UTR |
| hsa-miR-4699-3p | NM_001135099 | TMPRSS2 | 0.923076923077 | 3UTR |
| hsa-miR-4700-3p | NM_001135099 | TMPRSS2 | 0.923076923077 | 3UTR |
| hsa-miR-4709-3p | NM_001135099 | TMPRSS2 | 0.923076923077 | 3UTR |
| hsa-miR-4710 | NM_001135099 | TMPRSS2 | 0.923076923077 | 3UTR |
| hsa-miR-4711-3p | NM_001135099 | TMPRSS2 | 0.923076923077 | 3UTR |
| hsa-miR-4713-3p | NM_001135099 | TMPRSS2 | 0.923076923077 | 3UTR |
| hsa-miR-4715-3p | NM_001135099 | TMPRSS2 | 0.923076923077 | 3UTR |
| hsa-miR-4723-3p | NM_001135099 | TMPRSS2 | 0.923076923077 | 3UTR |
| hsa-miR-4726-5p | NM_001135099 | TMPRSS2 | 0.923076923077 | 3UTR |
| hsa-miR-4730 | NM_001135099 | TMPRSS2 | 0.923076923077 | 3UTR |
| hsa-miR-4731-3p | NM_001135099 | TMPRSS2 | 0.923076923077 | 3UTR |
| hsa-miR-4732-5p | NM_001135099 | TMPRSS2 | 0.923076923077 | 3UTR |
| hsa-miR-3064-5p | NM_001135099 | TMPRSS2 | 0.923076923077 | 3UTR |
| hsa-miR-4738-3p | NM_001135099 | TMPRSS2 | 0.923076923077 | 3UTR |
| hsa-miR-4738-3p | NM_001135099 | TMPRSS2 | 0.923076923077 | 3UTR |
| hsa-miR-4744 | NM_001135099 | TMPRSS2 | 0.923076923077 | 3UTR |
| hsa-miR-4746-5p | NM_001135099 | TMPRSS2 | 0.923076923077 | 3UTR |
| hsa-miR-4747-5p | NM_001135099 | TMPRSS2 | 0.923076923077 | 3UTR |
| hsa-miR-4748 | NM_001135099 | TMPRSS2 | 0.923076923077 | 3UTR |
| hsa-miR-4750-3p | NM_001135099 | TMPRSS2 | 0.923076923077 | 3UTR |
| hsa-miR-4755-5p | NM_001135099 | TMPRSS2 | 0.923076923077 | 3UTR |
| hsa-miR-4756-3p | NM_001135099 | TMPRSS2 | 0.923076923077 | 3UTR |
| hsa-miR-4772-3p | NM_001135099 | TMPRSS2 | 0.923076923077 | 3UTR |
| hsa-miR-4773 | NM_001135099 | TMPRSS2 | 0.923076923077 | 3UTR |

| | | | | |
|---|---|---|---|---|
| hsa-miR-4775 | NM_001135099 | TMPRSS2 | 0.923076923077 | 3UTR |
| hsa-miR-4776-5p | NM_001135099 | TMPRSS2 | 0.923076923077 | 3UTR |
| hsa-miR-1245b-3p | NM_001135099 | TMPRSS2 | 0.923076923077 | 3UTR |
| hsa-miR-4786-3p | NM_001135099 | TMPRSS2 | 0.923076923077 | 3UTR |
| hsa-miR-4788 | NM_001135099 | TMPRSS2 | 0.923076923077 | 3UTR |
| hsa-miR-4790-3p | NM_001135099 | TMPRSS2 | 0.923076923077 | 3UTR |
| hsa-miR-4793-5p | NM_001135099 | TMPRSS2 | 0.923076923077 | 3UTR |
| hsa-miR-4796-5p | NM_001135099 | TMPRSS2 | 0.923076923077 | 3UTR |
| hsa-miR-4797-5p | NM_001135099 | TMPRSS2 | 0.923076923077 | 3UTR |
| hsa-miR-4799-3p | NM_001135099 | TMPRSS2 | 0.923076923077 | 3UTR |
| hsa-miR-4800-5p | NM_001135099 | TMPRSS2 | 0.923076923077 | 3UTR |
| hsa-miR-5001-3p | NM_001135099 | TMPRSS2 | 0.923076923077 | 3UTR |
| hsa-miR-5002-5p | NM_001135099 | TMPRSS2 | 0.923076923077 | 3UTR |
| hsa-miR-5004-5p | NM_001135099 | TMPRSS2 | 0.923076923077 | 3UTR |
| hsa-miR-5008-3p | NM_001135099 | TMPRSS2 | 0.923076923077 | 3UTR |
| hsa-miR-5047 | NM_001135099 | TMPRSS2 | 0.923076923077 | 3UTR |
| hsa-miR-5087 | NM_001135099 | TMPRSS2 | 0.923076923077 | 3UTR |
| hsa-miR-5089-5p | NM_001135099 | TMPRSS2 | 0.923076923077 | 3UTR |
| hsa-miR-5090 | NM_001135099 | TMPRSS2 | 0.923076923077 | 3UTR |
| hsa-miR-5187-3p | NM_001135099 | TMPRSS2 | 0.923076923077 | 3UTR |
| hsa-miR-5193 | NM_001135099 | TMPRSS2 | 0.923076923077 | 3UTR |
| hsa-miR-5195-3p | NM_001135099 | TMPRSS2 | 0.923076923077 | 3UTR |
| hsa-miR-5196-5p | NM_001135099 | TMPRSS2 | 0.923076923077 | 3UTR |
| hsa-miR-5197-3p | NM_001135099 | TMPRSS2 | 0.923076923077 | 3UTR |
| hsa-miR-5580-3p | NM_001135099 | TMPRSS2 | 0.923076923077 | 3UTR |
| hsa-miR-5585-3p | NM_001135099 | TMPRSS2 | 0.923076923077 | 3UTR |
| hsa-miR-5589-5p | NM_001135099 | TMPRSS2 | 0.923076923077 | 3UTR |
| hsa-miR-5591-5p | NM_001135099 | TMPRSS2 | 0.923076923077 | 3UTR |
| hsa-miR-5591-5p | NM_001135099 | TMPRSS2 | 0.923076923077 | 3UTR |
| hsa-miR-5591-3p | NM_001135099 | TMPRSS2 | 0.923076923077 | 3UTR |
| hsa-miR-5682 | NM_001135099 | TMPRSS2 | 0.923076923077 | 3UTR |
| hsa-miR-5682 | NM_001135099 | TMPRSS2 | 0.923076923077 | 3UTR |
| hsa-miR-5687 | NM_001135099 | TMPRSS2 | 0.923076923077 | 3UTR |
| hsa-miR-5693 | NM_001135099 | TMPRSS2 | 0.923076923077 | 3UTR |
| hsa-miR-5704 | NM_001135099 | TMPRSS2 | 0.923076923077 | 3UTR |
| hsa-miR-5708 | NM_001135099 | TMPRSS2 | 0.923076923077 | 3UTR |
| hsa-miR-5739 | NM_001135099 | TMPRSS2 | 0.923076923077 | 3UTR |
| hsa-miR-5739 | NM_001135099 | TMPRSS2 | 0.923076923077 | 3UTR |
| hsa-miR-1199-3p | NM_001135099 | TMPRSS2 | 0.923076923077 | 3UTR |
| hsa-miR-6081 | NM_001135099 | TMPRSS2 | 0.923076923077 | 3UTR |
| hsa-miR-6085 | NM_001135099 | TMPRSS2 | 0.923076923077 | 3UTR |
| hsa-miR-6088 | NM_001135099 | TMPRSS2 | 0.923076923077 | 3UTR |
| hsa-miR-6127 | NM_001135099 | TMPRSS2 | 0.923076923077 | 3UTR |
| hsa-miR-6127 | NM_001135099 | TMPRSS2 | 0.923076923077 | 3UTR |
| hsa-miR-378j | NM_001135099 | TMPRSS2 | 0.923076923077 | 3UTR |
| hsa-miR-6131 | NM_001135099 | TMPRSS2 | 0.923076923077 | 3UTR |
| hsa-miR-6134 | NM_001135099 | TMPRSS2 | 0.923076923077 | 3UTR |

| | | | | |
|---|---|---|---|---|
| hsa-miR-6165 | NM_001135099 | TMPRSS2 | 0.923076923077 | 3UTR |
| hsa-miR-6165 | NM_001135099 | TMPRSS2 | 0.923076923077 | 3UTR |
| hsa-miR-6499-5p | NM_001135099 | TMPRSS2 | 0.923076923077 | 3UTR |
| hsa-miR-6503-3p | NM_001135099 | TMPRSS2 | 0.923076923077 | 3UTR |
| hsa-miR-6509-3p | NM_001135099 | TMPRSS2 | 0.923076923077 | 3UTR |
| hsa-miR-6510-5p | NM_001135099 | TMPRSS2 | 0.923076923077 | 3UTR |
| hsa-miR-6511a-3p | NM_001135099 | TMPRSS2 | 0.923076923077 | 3UTR |
| hsa-miR-6515-5p | NM_001135099 | TMPRSS2 | 0.923076923077 | 3UTR |
| hsa-miR-6515-5p | NM_001135099 | TMPRSS2 | 0.923076923077 | 3UTR |
| hsa-miR-6715b-3p | NM_001135099 | TMPRSS2 | 0.923076923077 | 3UTR |
| hsa-miR-6716-3p | NM_001135099 | TMPRSS2 | 0.923076923077 | 3UTR |
| hsa-miR-6717-5p | NM_001135099 | TMPRSS2 | 0.923076923077 | 3UTR |
| hsa-miR-6511b-5p | NM_001135099 | TMPRSS2 | 0.923076923077 | 3UTR |
| hsa-miR-6511b-5p | NM_001135099 | TMPRSS2 | 0.923076923077 | 3UTR |
| hsa-miR-6719-3p | NM_001135099 | TMPRSS2 | 0.923076923077 | 3UTR |
| hsa-miR-6720-5p | NM_001135099 | TMPRSS2 | 0.923076923077 | 3UTR |
| hsa-miR-6726-3p | NM_001135099 | TMPRSS2 | 0.923076923077 | 3UTR |
| hsa-miR-6728-5p | NM_001135099 | TMPRSS2 | 0.923076923077 | 3UTR |
| hsa-miR-6728-5p | NM_001135099 | TMPRSS2 | 0.923076923077 | 3UTR |
| hsa-miR-6731-3p | NM_001135099 | TMPRSS2 | 0.923076923077 | 3UTR |
| hsa-miR-6732-5p | NM_001135099 | TMPRSS2 | 0.923076923077 | 3UTR |
| hsa-miR-6734-5p | NM_001135099 | TMPRSS2 | 0.923076923077 | 3UTR |
| hsa-miR-6741-5p | NM_001135099 | TMPRSS2 | 0.923076923077 | 3UTR |
| hsa-miR-6746-5p | NM_001135099 | TMPRSS2 | 0.923076923077 | 3UTR |
| hsa-miR-6747-5p | NM_001135099 | TMPRSS2 | 0.923076923077 | 3UTR |
| hsa-miR-6747-5p | NM_001135099 | TMPRSS2 | 0.923076923077 | 3UTR |
| hsa-miR-6747-3p | NM_001135099 | TMPRSS2 | 0.923076923077 | 3UTR |
| hsa-miR-6747-3p | NM_001135099 | TMPRSS2 | 0.923076923077 | 3UTR |
| hsa-miR-6748-5p | NM_001135099 | TMPRSS2 | 0.923076923077 | 3UTR |
| hsa-miR-6749-5p | NM_001135099 | TMPRSS2 | 0.923076923077 | 3UTR |
| hsa-miR-6750-5p | NM_001135099 | TMPRSS2 | 0.923076923077 | 3UTR |
| hsa-miR-6751-3p | NM_001135099 | TMPRSS2 | 0.923076923077 | 3UTR |
| hsa-miR-6753-5p | NM_001135099 | TMPRSS2 | 0.923076923077 | 3UTR |
| hsa-miR-6758-5p | NM_001135099 | TMPRSS2 | 0.923076923077 | 3UTR |
| hsa-miR-6760-3p | NM_001135099 | TMPRSS2 | 0.923076923077 | 3UTR |
| hsa-miR-6762-3p | NM_001135099 | TMPRSS2 | 0.923076923077 | 3UTR |
| hsa-miR-6763-5p | NM_001135099 | TMPRSS2 | 0.923076923077 | 3UTR |
| hsa-miR-6763-5p | NM_001135099 | TMPRSS2 | 0.923076923077 | 3UTR |
| hsa-miR-6765-5p | NM_001135099 | TMPRSS2 | 0.923076923077 | 3UTR |
| hsa-miR-6765-3p | NM_001135099 | TMPRSS2 | 0.923076923077 | 3UTR |
| hsa-miR-6767-3p | NM_001135099 | TMPRSS2 | 0.923076923077 | 3UTR |
| hsa-miR-6769a-3p | NM_001135099 | TMPRSS2 | 0.923076923077 | 3UTR |
| hsa-miR-6771-5p | NM_001135099 | TMPRSS2 | 0.923076923077 | 3UTR |
| hsa-miR-6773-3p | NM_001135099 | TMPRSS2 | 0.923076923077 | 3UTR |
| hsa-miR-6774-3p | NM_001135099 | TMPRSS2 | 0.923076923077 | 3UTR |
| hsa-miR-6776-3p | NM_001135099 | TMPRSS2 | 0.923076923077 | 3UTR |
| hsa-miR-6776-3p | NM_001135099 | TMPRSS2 | 0.923076923077 | 3UTR |

| | | | | |
|---|---|---|---|---|
| hsa-miR-6777-5p | NM_001135099 | TMPRSS2 | 0.923076923077 | 3UTR |
| hsa-miR-6779-3p | NM_001135099 | TMPRSS2 | 0.923076923077 | 3UTR |
| hsa-miR-6780a-3p | NM_001135099 | TMPRSS2 | 0.923076923077 | 3UTR |
| hsa-miR-6781-3p | NM_001135099 | TMPRSS2 | 0.923076923077 | 3UTR |
| hsa-miR-6785-5p | NM_001135099 | TMPRSS2 | 0.923076923077 | 3UTR |
| hsa-miR-6791-3p | NM_001135099 | TMPRSS2 | 0.923076923077 | 3UTR |
| hsa-miR-6792-3p | NM_001135099 | TMPRSS2 | 0.923076923077 | 3UTR |
| hsa-miR-6793-3p | NM_001135099 | TMPRSS2 | 0.923076923077 | 3UTR |
| hsa-miR-6795-5p | NM_001135099 | TMPRSS2 | 0.923076923077 | 3UTR |
| hsa-miR-6797-5p | NM_001135099 | TMPRSS2 | 0.923076923077 | 3UTR |
| hsa-miR-6797-3p | NM_001135099 | TMPRSS2 | 0.923076923077 | 3UTR |
| hsa-miR-6798-5p | NM_001135099 | TMPRSS2 | 0.923076923077 | 3UTR |
| hsa-miR-6799-5p | NM_001135099 | TMPRSS2 | 0.923076923077 | 3UTR |
| hsa-miR-6799-3p | NM_001135099 | TMPRSS2 | 0.923076923077 | 3UTR |
| hsa-miR-6801-3p | NM_001135099 | TMPRSS2 | 0.923076923077 | 3UTR |
| hsa-miR-6802-5p | NM_001135099 | TMPRSS2 | 0.923076923077 | 3UTR |
| hsa-miR-6803-3p | NM_001135099 | TMPRSS2 | 0.923076923077 | 3UTR |
| hsa-miR-6804-5p | NM_001135099 | TMPRSS2 | 0.923076923077 | 3UTR |
| hsa-miR-6805-5p | NM_001135099 | TMPRSS2 | 0.923076923077 | 3UTR |
| hsa-miR-6807-3p | NM_001135099 | TMPRSS2 | 0.923076923077 | 3UTR |
| hsa-miR-6808-5p | NM_001135099 | TMPRSS2 | 0.923076923077 | 3UTR |
| hsa-miR-6809-5p | NM_001135099 | TMPRSS2 | 0.923076923077 | 3UTR |
| hsa-miR-6810-5p | NM_001135099 | TMPRSS2 | 0.923076923077 | 3UTR |
| hsa-miR-6811-3p | NM_001135099 | TMPRSS2 | 0.923076923077 | 3UTR |
| hsa-miR-6812-5p | NM_001135099 | TMPRSS2 | 0.923076923077 | 3UTR |
| hsa-miR-6812-3p | NM_001135099 | TMPRSS2 | 0.923076923077 | 3UTR |
| hsa-miR-6814-3p | NM_001135099 | TMPRSS2 | 0.923076923077 | 3UTR |
| hsa-miR-6815-5p | NM_001135099 | TMPRSS2 | 0.923076923077 | 3UTR |
| hsa-miR-6815-3p | NM_001135099 | TMPRSS2 | 0.923076923077 | 3UTR |
| hsa-miR-6817-5p | NM_001135099 | TMPRSS2 | 0.923076923077 | 3UTR |
| hsa-miR-6819-5p | NM_001135099 | TMPRSS2 | 0.923076923077 | 3UTR |
| hsa-miR-6821-3p | NM_001135099 | TMPRSS2 | 0.923076923077 | 3UTR |
| hsa-miR-6823-5p | NM_001135099 | TMPRSS2 | 0.923076923077 | 3UTR |
| hsa-miR-6825-5p | NM_001135099 | TMPRSS2 | 0.923076923077 | 3UTR |
| hsa-miR-6829-3p | NM_001135099 | TMPRSS2 | 0.923076923077 | 3UTR |
| hsa-miR-6832-5p | NM_001135099 | TMPRSS2 | 0.923076923077 | 3UTR |
| hsa-miR-6832-5p | NM_001135099 | TMPRSS2 | 0.923076923077 | 3UTR |
| hsa-miR-6836-3p | NM_001135099 | TMPRSS2 | 0.923076923077 | 3UTR |
| hsa-miR-6846-5p | NM_001135099 | TMPRSS2 | 0.923076923077 | 3UTR |
| hsa-miR-6846-3p | NM_001135099 | TMPRSS2 | 0.923076923077 | 3UTR |
| hsa-miR-6851-5p | NM_001135099 | TMPRSS2 | 0.923076923077 | 3UTR |
| hsa-miR-6851-3p | NM_001135099 | TMPRSS2 | 0.923076923077 | 3UTR |
| hsa-miR-6854-5p | NM_001135099 | TMPRSS2 | 0.923076923077 | 3UTR |
| hsa-miR-6856-5p | NM_001135099 | TMPRSS2 | 0.923076923077 | 3UTR |
| hsa-miR-6857-3p | NM_001135099 | TMPRSS2 | 0.923076923077 | 3UTR |
| hsa-miR-6859-5p | NM_001135099 | TMPRSS2 | 0.923076923077 | 3UTR |
| hsa-miR-6862-5p | NM_001135099 | TMPRSS2 | 0.923076923077 | 3UTR |

| | | | | |
|---|---|---|---|---|
| hsa-miR-6863 | NM_001135099 | TMPRSS2 | 0.923076923077 | 3UTR |
| hsa-miR-6864-3p | NM_001135099 | TMPRSS2 | 0.923076923077 | 3UTR |
| hsa-miR-6865-5p | NM_001135099 | TMPRSS2 | 0.923076923077 | 3UTR |
| hsa-miR-6865-3p | NM_001135099 | TMPRSS2 | 0.923076923077 | 3UTR |
| hsa-miR-6867-3p | NM_001135099 | TMPRSS2 | 0.923076923077 | 3UTR |
| hsa-miR-6868-5p | NM_001135099 | TMPRSS2 | 0.923076923077 | 3UTR |
| hsa-miR-6873-5p | NM_001135099 | TMPRSS2 | 0.923076923077 | 3UTR |
| hsa-miR-6873-3p | NM_001135099 | TMPRSS2 | 0.923076923077 | 3UTR |
| hsa-miR-6878-3p | NM_001135099 | TMPRSS2 | 0.923076923077 | 3UTR |
| hsa-miR-6880-5p | NM_001135099 | TMPRSS2 | 0.923076923077 | 3UTR |
| hsa-miR-6883-3p | NM_001135099 | TMPRSS2 | 0.923076923077 | 3UTR |
| hsa-miR-6884-5p | NM_001135099 | TMPRSS2 | 0.923076923077 | 3UTR |
| hsa-miR-6884-5p | NM_001135099 | TMPRSS2 | 0.923076923077 | 3UTR |
| hsa-miR-6886-3p | NM_001135099 | TMPRSS2 | 0.923076923077 | 3UTR |
| hsa-miR-6887-5p | NM_001135099 | TMPRSS2 | 0.923076923077 | 3UTR |
| hsa-miR-6887-3p | NM_001135099 | TMPRSS2 | 0.923076923077 | 3UTR |
| hsa-miR-6890-3p | NM_001135099 | TMPRSS2 | 0.923076923077 | 3UTR |
| hsa-miR-6890-3p | NM_001135099 | TMPRSS2 | 0.923076923077 | 3UTR |
| hsa-miR-6892-5p | NM_001135099 | TMPRSS2 | 0.923076923077 | 3UTR |
| hsa-miR-6892-3p | NM_001135099 | TMPRSS2 | 0.923076923077 | 3UTR |
| hsa-miR-6895-3p | NM_001135099 | TMPRSS2 | 0.923076923077 | 3UTR |
| hsa-miR-7110-5p | NM_001135099 | TMPRSS2 | 0.923076923077 | 3UTR |
| hsa-miR-7111-5p | NM_001135099 | TMPRSS2 | 0.923076923077 | 3UTR |
| hsa-miR-7111-3p | NM_001135099 | TMPRSS2 | 0.923076923077 | 3UTR |
| hsa-miR-7113-5p | NM_001135099 | TMPRSS2 | 0.923076923077 | 3UTR |
| hsa-miR-7114-3p | NM_001135099 | TMPRSS2 | 0.923076923077 | 3UTR |
| hsa-miR-7152-3p | NM_001135099 | TMPRSS2 | 0.923076923077 | 3UTR |
| hsa-miR-7152-3p | NM_001135099 | TMPRSS2 | 0.923076923077 | 3UTR |
| hsa-miR-7152-3p | NM_001135099 | TMPRSS2 | 0.923076923077 | 3UTR |
| hsa-miR-7153-3p | NM_001135099 | TMPRSS2 | 0.923076923077 | 3UTR |
| hsa-miR-7161-5p | NM_001135099 | TMPRSS2 | 0.923076923077 | 3UTR |
| hsa-miR-7161-3p | NM_001135099 | TMPRSS2 | 0.923076923077 | 3UTR |
| hsa-miR-7160-3p | NM_001135099 | TMPRSS2 | 0.923076923077 | 3UTR |
| hsa-miR-7162-3p | NM_001135099 | TMPRSS2 | 0.923076923077 | 3UTR |
| hsa-miR-7703 | NM_001135099 | TMPRSS2 | 0.923076923077 | 3UTR |
| hsa-miR-7706 | NM_001135099 | TMPRSS2 | 0.923076923077 | 3UTR |
| hsa-miR-7843-5p | NM_001135099 | TMPRSS2 | 0.923076923077 | 3UTR |
| hsa-miR-4433b-5p | NM_001135099 | TMPRSS2 | 0.923076923077 | 3UTR |
| hsa-miR-4433b-3p | NM_001135099 | TMPRSS2 | 0.923076923077 | 3UTR |
| hsa-miR-4433b-3p | NM_001135099 | TMPRSS2 | 0.923076923077 | 3UTR |
| hsa-miR-7848-3p | NM_001135099 | TMPRSS2 | 0.923076923077 | 3UTR |
| hsa-miR-7854-3p | NM_001135099 | TMPRSS2 | 0.923076923077 | 3UTR |
| hsa-miR-7978 | NM_001135099 | TMPRSS2 | 0.923076923077 | 3UTR |
| hsa-miR-8077 | NM_001135099 | TMPRSS2 | 0.923076923077 | 3UTR |
| hsa-miR-8085 | NM_001135099 | TMPRSS2 | 0.923076923077 | 3UTR |
| hsa-miR-10226 | NM_001135099 | TMPRSS2 | 0.923076923077 | 3UTR |
| hsa-miR-10392-3p | NM_001135099 | TMPRSS2 | 0.923076923077 | 3UTR |

| | | | | |
|---|---|---|---|---|
| hsa-miR-10394-5p | NM_001135099 | TMPRSS2 | 0.923076923077 | 3UTR |
| hsa-miR-10397-5p | NM_001135099 | TMPRSS2 | 0.923076923077 | 3UTR |
| hsa-miR-10399-3p | NM_001135099 | TMPRSS2 | 0.923076923077 | 3UTR |
| hsa-miR-10401-5p | NM_001135099 | TMPRSS2 | 0.923076923077 | 3UTR |
| hsa-miR-10524-5p | NM_001135099 | TMPRSS2 | 0.923076923077 | 3UTR |
| hsa-miR-10526-3p | NM_001135099 | TMPRSS2 | 0.923076923077 | 3UTR |
| hsa-miR-11181-5p | NM_001135099 | TMPRSS2 | 0.923076923077 | 3UTR |
| hsa-miR-3059-5p | NM_001135099 | TMPRSS2 | 0.923076923077 | 3UTR |
| hsa-miR-3059-3p | NM_001135099 | TMPRSS2 | 0.923076923077 | 3UTR |
| hsa-miR-3059-3p | NM_001135099 | TMPRSS2 | 0.923076923077 | 3UTR |
| hsa-miR-3085-3p | NM_001135099 | TMPRSS2 | 0.923076923077 | 3UTR |
| hsa-miR-12115 | NM_001135099 | TMPRSS2 | 0.923076923077 | 3UTR |
| hsa-miR-12118 | NM_001135099 | TMPRSS2 | 0.923076923077 | 3UTR |
| hsa-miR-12119 | NM_001135099 | TMPRSS2 | 0.923076923077 | 3UTR |
| hsa-miR-12122 | NM_001135099 | TMPRSS2 | 0.923076923077 | 3UTR |
| hsa-miR-12124 | NM_001135099 | TMPRSS2 | 0.923076923077 | 3UTR |
| hsa-miR-12126 | NM_001135099 | TMPRSS2 | 0.923076923077 | 3UTR |
| hsa-miR-12128 | NM_001135099 | TMPRSS2 | 0.923076923077 | 3UTR |
| hsa-miR-12131 | NM_001135099 | TMPRSS2 | 0.923076923077 | 3UTR |
| hsa-miR-6861-3p | NM_001135099 | TMPRSS2 | 0.935897435897 | 3UTR |
| hsa-miR-149-5p | NM_005656 | TMPRSS2 | 0.948717948718 | 3UTR |
| hsa-miR-381-3p | NM_005656 | TMPRSS2 | 0.948717948718 | 3UTR |
| hsa-miR-3620-5p | NM_005656 | TMPRSS2 | 0.948717948718 | 3UTR |
| hsa-miR-6727-5p | NM_005656 | TMPRSS2 | 0.948717948718 | 3UTR |
| hsa-miR-6809-3p | NM_005656 | TMPRSS2 | 0.948717948718 | 3UTR |
| hsa-miR-149-5p | NM_001135099 | TMPRSS2 | 0.948717948718 | 3UTR |
| hsa-miR-6809-3p | NM_001135099 | TMPRSS2 | 0.948717948718 | 3UTR |
| hsa-miR-6866-3p | NM_001135099 | TMPRSS2 | 0.948717948718 | 3UTR |
| hsa-miR-605-5p | NM_005656 | TMPRSS2 | 0.953846153846 | 3UTR |
| hsa-miR-6133 | NM_001135099 | TMPRSS2 | 0.953846153846 | 3UTR |
| hsa-miR-619-5p | NM_005656 | TMPRSS2 | 0.961538461538 | 3UTR |
| hsa-miR-4429 | NM_005656 | TMPRSS2 | 0.961538461538 | 3UTR |
| hsa-miR-12115 | NM_005656 | TMPRSS2 | 0.961538461538 | 3UTR |
| hsa-miR-12131 | NM_005656 | TMPRSS2 | 0.961538461538 | 3UTR |
| hsa-let-7a-5p | NM_001135099 | TMPRSS2 | 0.961538461538 | 3UTR |
| hsa-let-7f-5p | NM_001135099 | TMPRSS2 | 0.961538461538 | 3UTR |
| hsa-miR-98-5p | NM_001135099 | TMPRSS2 | 0.961538461538 | 3UTR |
| hsa-miR-140-5p | NM_001135099 | TMPRSS2 | 0.961538461538 | 3UTR |
| hsa-miR-3177-3p | NM_001135099 | TMPRSS2 | 0.961538461538 | 3UTR |
| hsa-miR-4448 | NM_001135099 | TMPRSS2 | 0.961538461538 | 3UTR |
| hsa-miR-4689 | NM_001135099 | TMPRSS2 | 0.961538461538 | 3UTR |
| hsa-miR-5094 | NM_001135099 | TMPRSS2 | 0.961538461538 | 3UTR |
| hsa-miR-6812-3p | NM_001135099 | TMPRSS2 | 0.961538461538 | 3UTR |
| hsa-miR-6887-5p | NM_001135099 | TMPRSS2 | 0.961538461538 | 3UTR |
| hsa-miR-10394-3p | NM_001135099 | TMPRSS2 | 0.961538461538 | 3UTR |
| hsa-miR-3929 | NM_005656 | TMPRSS2 | 0.969230769231 | 3UTR |
| hsa-miR-1304-3p | NM_001135099 | TMPRSS2 | 0.969230769231 | 3UTR |

| | | | | |
|---|---|---|---|---|
| hsa-miR-3174 | NM_001135099 | TMPRSS2 | 0.969230769231 | 3UTR |
| hsa-miR-141-3p | NM_005656 | TMPRSS2 | 0.974358974359 | 3UTR |
| hsa-miR-452-5p | NM_005656 | TMPRSS2 | 0.974358974359 | 3UTR |
| hsa-miR-504-3p | NM_005656 | TMPRSS2 | 0.974358974359 | 3UTR |
| hsa-miR-652-3p | NM_005656 | TMPRSS2 | 0.974358974359 | 3UTR |
| hsa-miR-4519 | NM_005656 | TMPRSS2 | 0.974358974359 | 3UTR |
| hsa-miR-6781-3p | NM_005656 | TMPRSS2 | 0.974358974359 | 3UTR |
| hsa-miR-452-5p | NM_001135099 | TMPRSS2 | 0.974358974359 | 3UTR |
| hsa-miR-504-3p | NM_001135099 | TMPRSS2 | 0.974358974359 | 3UTR |
| hsa-miR-652-3p | NM_001135099 | TMPRSS2 | 0.974358974359 | 3UTR |
| hsa-miR-1296-3p | NM_001135099 | TMPRSS2 | 0.974358974359 | 3UTR |
| hsa-miR-4519 | NM_001135099 | TMPRSS2 | 0.974358974359 | 3UTR |
| hsa-miR-6781-3p | NM_001135099 | TMPRSS2 | 0.974358974359 | 3UTR |
| hsa-let-7b-5p | NM_005656 | TMPRSS2 | 1.0 | 3UTR |
| hsa-miR-21-3p | NM_005656 | TMPRSS2 | 1.0 | 3UTR |
| hsa-miR-23a-5p | NM_005656 | TMPRSS2 | 1.0 | 3UTR |
| hsa-miR-25-5p | NM_005656 | TMPRSS2 | 1.0 | 3UTR |
| hsa-miR-26b-3p | NM_005656 | TMPRSS2 | 1.0 | 3UTR |
| hsa-miR-28-5p | NM_005656 | TMPRSS2 | 1.0 | 3UTR |
| hsa-miR-96-5p | NM_005656 | TMPRSS2 | 1.0 | 3UTR |
| hsa-miR-98-5p | NM_005656 | TMPRSS2 | 1.0 | 3UTR |
| hsa-miR-101-5p | NM_005656 | TMPRSS2 | 1.0 | 3UTR |
| hsa-miR-103a-3p | NM_005656 | TMPRSS2 | 1.0 | 3UTR |
| hsa-miR-107 | NM_005656 | TMPRSS2 | 1.0 | 3UTR |
| hsa-miR-129-1-3p | NM_005656 | TMPRSS2 | 1.0 | 3UTR |
| hsa-miR-30d-5p | NM_005656 | TMPRSS2 | 1.0 | 3UTR |
| hsa-miR-139-5p | NM_005656 | TMPRSS2 | 1.0 | 3UTR |
| hsa-miR-34a-5p | NM_005656 | TMPRSS2 | 1.0 | 3UTR |
| hsa-miR-34a-5p | NM_005656 | TMPRSS2 | 1.0 | 3UTR |
| hsa-miR-183-5p | NM_005656 | TMPRSS2 | 1.0 | 3UTR |
| hsa-miR-200b-5p | NM_005656 | TMPRSS2 | 1.0 | 3UTR |
| hsa-let-7g-5p | NM_005656 | TMPRSS2 | 1.0 | 3UTR |
| hsa-let-7i-5p | NM_005656 | TMPRSS2 | 1.0 | 3UTR |
| hsa-miR-30b-3p | NM_005656 | TMPRSS2 | 1.0 | 3UTR |
| hsa-miR-124-5p | NM_005656 | TMPRSS2 | 1.0 | 3UTR |
| hsa-miR-140-5p | NM_005656 | TMPRSS2 | 1.0 | 3UTR |
| hsa-miR-129-2-3p | NM_005656 | TMPRSS2 | 1.0 | 3UTR |
| hsa-miR-134-5p | NM_005656 | TMPRSS2 | 1.0 | 3UTR |
| hsa-miR-185-5p | NM_005656 | TMPRSS2 | 1.0 | 3UTR |
| hsa-miR-185-3p | NM_005656 | TMPRSS2 | 1.0 | 3UTR |
| hsa-miR-188-5p | NM_005656 | TMPRSS2 | 1.0 | 3UTR |
| hsa-miR-320a-3p | NM_005656 | TMPRSS2 | 1.0 | 3UTR |
| hsa-miR-194-3p | NM_005656 | TMPRSS2 | 1.0 | 3UTR |
| hsa-miR-106b-3p | NM_005656 | TMPRSS2 | 1.0 | 3UTR |
| hsa-miR-302c-5p | NM_005656 | TMPRSS2 | 1.0 | 3UTR |
| hsa-miR-367-3p | NM_005656 | TMPRSS2 | 1.0 | 3UTR |
| hsa-miR-371a-5p | NM_005656 | TMPRSS2 | 1.0 | 3UTR |

| | | | | |
|---|---|---|---|---|
| hsa-miR-373-5p | NM_005656 | TMPRSS2 | 1.0 | 3UTR |
| hsa-miR-379-5p | NM_005656 | TMPRSS2 | 1.0 | 3UTR |
| hsa-miR-151a-5p | NM_005656 | TMPRSS2 | 1.0 | 3UTR |
| hsa-miR-151a-3p | NM_005656 | TMPRSS2 | 1.0 | 3UTR |
| hsa-miR-324-5p | NM_005656 | TMPRSS2 | 1.0 | 3UTR |
| hsa-miR-324-3p | NM_005656 | TMPRSS2 | 1.0 | 3UTR |
| hsa-miR-339-5p | NM_005656 | TMPRSS2 | 1.0 | 3UTR |
| hsa-miR-423-5p | NM_005656 | TMPRSS2 | 1.0 | 3UTR |
| hsa-miR-423-3p | NM_005656 | TMPRSS2 | 1.0 | 3UTR |
| hsa-miR-20b-3p | NM_005656 | TMPRSS2 | 1.0 | 3UTR |
| hsa-miR-431-3p | NM_005656 | TMPRSS2 | 1.0 | 3UTR |
| hsa-miR-412-3p | NM_005656 | TMPRSS2 | 1.0 | 3UTR |
| hsa-miR-491-5p | NM_005656 | TMPRSS2 | 1.0 | 3UTR |
| hsa-miR-146b-3p | NM_005656 | TMPRSS2 | 1.0 | 3UTR |
| hsa-miR-202-3p | NM_005656 | TMPRSS2 | 1.0 | 3UTR |
| hsa-miR-493-3p | NM_005656 | TMPRSS2 | 1.0 | 3UTR |
| hsa-miR-432-5p | NM_005656 | TMPRSS2 | 1.0 | 3UTR |
| hsa-miR-193b-5p | NM_005656 | TMPRSS2 | 1.0 | 3UTR |
| hsa-miR-181d-5p | NM_005656 | TMPRSS2 | 1.0 | 3UTR |
| hsa-miR-520c-3p | NM_005656 | TMPRSS2 | 1.0 | 3UTR |
| hsa-miR-518a-5p | NM_005656 | TMPRSS2 | 1.0 | 3UTR |
| hsa-miR-527 | NM_005656 | TMPRSS2 | 1.0 | 3UTR |
| hsa-miR-500a-5p | NM_005656 | TMPRSS2 | 1.0 | 3UTR |
| hsa-miR-502-3p | NM_005656 | TMPRSS2 | 1.0 | 3UTR |
| hsa-miR-503-3p | NM_005656 | TMPRSS2 | 1.0 | 3UTR |
| hsa-miR-504-3p | NM_005656 | TMPRSS2 | 1.0 | 3UTR |
| hsa-miR-505-5p | NM_005656 | TMPRSS2 | 1.0 | 3UTR |
| hsa-miR-505-5p | NM_005656 | TMPRSS2 | 1.0 | 3UTR |
| hsa-miR-505-3p | NM_005656 | TMPRSS2 | 1.0 | 3UTR |
| hsa-miR-514a-5p | NM_005656 | TMPRSS2 | 1.0 | 3UTR |
| hsa-miR-455-5p | NM_005656 | TMPRSS2 | 1.0 | 3UTR |
| hsa-miR-455-3p | NM_005656 | TMPRSS2 | 1.0 | 3UTR |
| hsa-miR-554 | NM_005656 | TMPRSS2 | 1.0 | 3UTR |
| hsa-miR-557 | NM_005656 | TMPRSS2 | 1.0 | 3UTR |
| hsa-miR-575 | NM_005656 | TMPRSS2 | 1.0 | 3UTR |
| hsa-miR-584-3p | NM_005656 | TMPRSS2 | 1.0 | 3UTR |
| hsa-miR-550a-5p | NM_005656 | TMPRSS2 | 1.0 | 3UTR |
| hsa-miR-550a-5p | NM_005656 | TMPRSS2 | 1.0 | 3UTR |
| hsa-miR-600 | NM_005656 | TMPRSS2 | 1.0 | 3UTR |
| hsa-miR-608 | NM_005656 | TMPRSS2 | 1.0 | 3UTR |
| hsa-miR-608 | NM_005656 | TMPRSS2 | 1.0 | 3UTR |
| hsa-miR-614 | NM_005656 | TMPRSS2 | 1.0 | 3UTR |
| hsa-miR-622 | NM_005656 | TMPRSS2 | 1.0 | 3UTR |
| hsa-miR-645 | NM_005656 | TMPRSS2 | 1.0 | 3UTR |
| hsa-miR-648 | NM_005656 | TMPRSS2 | 1.0 | 3UTR |
| hsa-miR-650 | NM_005656 | TMPRSS2 | 1.0 | 3UTR |
| hsa-miR-449b-5p | NM_005656 | TMPRSS2 | 1.0 | 3UTR |

| | | | | |
|---|---|---|---|---|
| hsa-miR-549a-5p | NM_005656 | TMPRSS2 | 1.0 | 3UTR |
| hsa-miR-671-5p | NM_005656 | TMPRSS2 | 1.0 | 3UTR |
| hsa-miR-671-5p | NM_005656 | TMPRSS2 | 1.0 | 3UTR |
| hsa-miR-671-3p | NM_005656 | TMPRSS2 | 1.0 | 3UTR |
| hsa-miR-767-5p | NM_005656 | TMPRSS2 | 1.0 | 3UTR |
| hsa-miR-320b | NM_005656 | TMPRSS2 | 1.0 | 3UTR |
| hsa-miR-1296-3p | NM_005656 | TMPRSS2 | 1.0 | 3UTR |
| hsa-miR-1271-3p | NM_005656 | TMPRSS2 | 1.0 | 3UTR |
| hsa-miR-449c-3p | NM_005656 | TMPRSS2 | 1.0 | 3UTR |
| hsa-miR-769-5p | NM_005656 | TMPRSS2 | 1.0 | 3UTR |
| hsa-miR-766-5p | NM_005656 | TMPRSS2 | 1.0 | 3UTR |
| hsa-miR-675-5p | NM_005656 | TMPRSS2 | 1.0 | 3UTR |
| hsa-miR-890 | NM_005656 | TMPRSS2 | 1.0 | 3UTR |
| hsa-miR-708-5p | NM_005656 | TMPRSS2 | 1.0 | 3UTR |
| hsa-miR-147b-3p | NM_005656 | TMPRSS2 | 1.0 | 3UTR |
| hsa-miR-921 | NM_005656 | TMPRSS2 | 1.0 | 3UTR |
| hsa-miR-922 | NM_005656 | TMPRSS2 | 1.0 | 3UTR |
| hsa-miR-933 | NM_005656 | TMPRSS2 | 1.0 | 3UTR |
| hsa-miR-933 | NM_005656 | TMPRSS2 | 1.0 | 3UTR |
| hsa-miR-939-5p | NM_005656 | TMPRSS2 | 1.0 | 3UTR |
| hsa-miR-939-5p | NM_005656 | TMPRSS2 | 1.0 | 3UTR |
| hsa-miR-943 | NM_005656 | TMPRSS2 | 1.0 | 3UTR |
| hsa-miR-1180-5p | NM_005656 | TMPRSS2 | 1.0 | 3UTR |
| hsa-miR-1181 | NM_005656 | TMPRSS2 | 1.0 | 3UTR |
| hsa-miR-1227-5p | NM_005656 | TMPRSS2 | 1.0 | 3UTR |
| hsa-miR-1227-3p | NM_005656 | TMPRSS2 | 1.0 | 3UTR |
| hsa-miR-1229-5p | NM_005656 | TMPRSS2 | 1.0 | 3UTR |
| hsa-miR-1234-3p | NM_005656 | TMPRSS2 | 1.0 | 3UTR |
| hsa-miR-1234-3p | NM_005656 | TMPRSS2 | 1.0 | 3UTR |
| hsa-miR-1207-3p | NM_005656 | TMPRSS2 | 1.0 | 3UTR |
| hsa-miR-1286 | NM_005656 | TMPRSS2 | 1.0 | 3UTR |
| hsa-miR-1304-3p | NM_005656 | TMPRSS2 | 1.0 | 3UTR |
| hsa-miR-1245a | NM_005656 | TMPRSS2 | 1.0 | 3UTR |
| hsa-miR-1249-3p | NM_005656 | TMPRSS2 | 1.0 | 3UTR |
| hsa-miR-1253 | NM_005656 | TMPRSS2 | 1.0 | 3UTR |
| hsa-miR-1256 | NM_005656 | TMPRSS2 | 1.0 | 3UTR |
| hsa-miR-1263 | NM_005656 | TMPRSS2 | 1.0 | 3UTR |
| hsa-miR-1266-5p | NM_005656 | TMPRSS2 | 1.0 | 3UTR |
| hsa-miR-1275 | NM_005656 | TMPRSS2 | 1.0 | 3UTR |
| hsa-miR-1255b-2-3p | NM_005656 | TMPRSS2 | 1.0 | 3UTR |
| hsa-miR-103b | NM_005656 | TMPRSS2 | 1.0 | 3UTR |
| hsa-miR-1825 | NM_005656 | TMPRSS2 | 1.0 | 3UTR |
| hsa-miR-1909-5p | NM_005656 | TMPRSS2 | 1.0 | 3UTR |
| hsa-miR-1910-3p | NM_005656 | TMPRSS2 | 1.0 | 3UTR |
| hsa-miR-2117 | NM_005656 | TMPRSS2 | 1.0 | 3UTR |
| hsa-miR-2277-5p | NM_005656 | TMPRSS2 | 1.0 | 3UTR |
| hsa-miR-2681-5p | NM_005656 | TMPRSS2 | 1.0 | 3UTR |

| | | | | |
|---|---|---|---|---|
| hsa-miR-2682-3p | NM_005656 | TMPRSS2 | 1.0 | 3UTR |
| hsa-miR-3125 | NM_005656 | TMPRSS2 | 1.0 | 3UTR |
| hsa-miR-3129-5p | NM_005656 | TMPRSS2 | 1.0 | 3UTR |
| hsa-miR-3131 | NM_005656 | TMPRSS2 | 1.0 | 3UTR |
| hsa-miR-3137 | NM_005656 | TMPRSS2 | 1.0 | 3UTR |
| hsa-miR-3144-5p | NM_005656 | TMPRSS2 | 1.0 | 3UTR |
| hsa-miR-3150a-5p | NM_005656 | TMPRSS2 | 1.0 | 3UTR |
| hsa-miR-3153 | NM_005656 | TMPRSS2 | 1.0 | 3UTR |
| hsa-miR-3156-5p | NM_005656 | TMPRSS2 | 1.0 | 3UTR |
| hsa-miR-3162-5p | NM_005656 | TMPRSS2 | 1.0 | 3UTR |
| hsa-miR-3166 | NM_005656 | TMPRSS2 | 1.0 | 3UTR |
| hsa-miR-3170 | NM_005656 | TMPRSS2 | 1.0 | 3UTR |
| hsa-miR-3175 | NM_005656 | TMPRSS2 | 1.0 | 3UTR |
| hsa-miR-3177-3p | NM_005656 | TMPRSS2 | 1.0 | 3UTR |
| hsa-miR-3187-3p | NM_005656 | TMPRSS2 | 1.0 | 3UTR |
| hsa-miR-3191-3p | NM_005656 | TMPRSS2 | 1.0 | 3UTR |
| hsa-miR-3192-5p | NM_005656 | TMPRSS2 | 1.0 | 3UTR |
| hsa-miR-3193 | NM_005656 | TMPRSS2 | 1.0 | 3UTR |
| hsa-miR-3194-5p | NM_005656 | TMPRSS2 | 1.0 | 3UTR |
| hsa-miR-3202 | NM_005656 | TMPRSS2 | 1.0 | 3UTR |
| hsa-miR-4298 | NM_005656 | TMPRSS2 | 1.0 | 3UTR |
| hsa-miR-4313 | NM_005656 | TMPRSS2 | 1.0 | 3UTR |
| hsa-miR-4321 | NM_005656 | TMPRSS2 | 1.0 | 3UTR |
| hsa-miR-4259 | NM_005656 | TMPRSS2 | 1.0 | 3UTR |
| hsa-miR-4260 | NM_005656 | TMPRSS2 | 1.0 | 3UTR |
| hsa-miR-4252 | NM_005656 | TMPRSS2 | 1.0 | 3UTR |
| hsa-miR-4327 | NM_005656 | TMPRSS2 | 1.0 | 3UTR |
| hsa-miR-4327 | NM_005656 | TMPRSS2 | 1.0 | 3UTR |
| hsa-miR-4269 | NM_005656 | TMPRSS2 | 1.0 | 3UTR |
| hsa-miR-3619-5p | NM_005656 | TMPRSS2 | 1.0 | 3UTR |
| hsa-miR-3621 | NM_005656 | TMPRSS2 | 1.0 | 3UTR |
| hsa-miR-3646 | NM_005656 | TMPRSS2 | 1.0 | 3UTR |
| hsa-miR-3652 | NM_005656 | TMPRSS2 | 1.0 | 3UTR |
| hsa-miR-3652 | NM_005656 | TMPRSS2 | 1.0 | 3UTR |
| hsa-miR-3655 | NM_005656 | TMPRSS2 | 1.0 | 3UTR |
| hsa-miR-3663-5p | NM_005656 | TMPRSS2 | 1.0 | 3UTR |
| hsa-miR-3663-5p | NM_005656 | TMPRSS2 | 1.0 | 3UTR |
| hsa-miR-3664-3p | NM_005656 | TMPRSS2 | 1.0 | 3UTR |
| hsa-miR-3667-3p | NM_005656 | TMPRSS2 | 1.0 | 3UTR |
| hsa-miR-3678-3p | NM_005656 | TMPRSS2 | 1.0 | 3UTR |
| hsa-miR-3679-5p | NM_005656 | TMPRSS2 | 1.0 | 3UTR |
| hsa-miR-3679-5p | NM_005656 | TMPRSS2 | 1.0 | 3UTR |
| hsa-miR-3689a-3p | NM_005656 | TMPRSS2 | 1.0 | 3UTR |
| hsa-miR-3907 | NM_005656 | TMPRSS2 | 1.0 | 3UTR |
| hsa-miR-3917 | NM_005656 | TMPRSS2 | 1.0 | 3UTR |
| hsa-miR-3150b-5p | NM_005656 | TMPRSS2 | 1.0 | 3UTR |
| hsa-miR-3926 | NM_005656 | TMPRSS2 | 1.0 | 3UTR |

| | | | | |
|---|---|---|---|---|
| hsa-miR-3928-3p | NM_005656 | TMPRSS2 | 1.0 | 3UTR |
| hsa-miR-3934-3p | NM_005656 | TMPRSS2 | 1.0 | 3UTR |
| hsa-miR-3944-5p | NM_005656 | TMPRSS2 | 1.0 | 3UTR |
| hsa-miR-642b-3p | NM_005656 | TMPRSS2 | 1.0 | 3UTR |
| hsa-miR-642b-3p | NM_005656 | TMPRSS2 | 1.0 | 3UTR |
| hsa-miR-550b-2-5p | NM_005656 | TMPRSS2 | 1.0 | 3UTR |
| hsa-miR-4418 | NM_005656 | TMPRSS2 | 1.0 | 3UTR |
| hsa-miR-378f | NM_005656 | TMPRSS2 | 1.0 | 3UTR |
| hsa-miR-4425 | NM_005656 | TMPRSS2 | 1.0 | 3UTR |
| hsa-miR-4428 | NM_005656 | TMPRSS2 | 1.0 | 3UTR |
| hsa-miR-4433a-3p | NM_005656 | TMPRSS2 | 1.0 | 3UTR |
| hsa-miR-4436a | NM_005656 | TMPRSS2 | 1.0 | 3UTR |
| hsa-miR-4439 | NM_005656 | TMPRSS2 | 1.0 | 3UTR |
| hsa-miR-4441 | NM_005656 | TMPRSS2 | 1.0 | 3UTR |
| hsa-miR-4446-3p | NM_005656 | TMPRSS2 | 1.0 | 3UTR |
| hsa-miR-4448 | NM_005656 | TMPRSS2 | 1.0 | 3UTR |
| hsa-miR-4449 | NM_005656 | TMPRSS2 | 1.0 | 3UTR |
| hsa-miR-548ah-5p | NM_005656 | TMPRSS2 | 1.0 | 3UTR |
| hsa-miR-378h | NM_005656 | TMPRSS2 | 1.0 | 3UTR |
| hsa-miR-4469 | NM_005656 | TMPRSS2 | 1.0 | 3UTR |
| hsa-miR-4472 | NM_005656 | TMPRSS2 | 1.0 | 3UTR |
| hsa-miR-4481 | NM_005656 | TMPRSS2 | 1.0 | 3UTR |
| hsa-miR-4484 | NM_005656 | TMPRSS2 | 1.0 | 3UTR |
| hsa-miR-4488 | NM_005656 | TMPRSS2 | 1.0 | 3UTR |
| hsa-miR-4489 | NM_005656 | TMPRSS2 | 1.0 | 3UTR |
| hsa-miR-4501 | NM_005656 | TMPRSS2 | 1.0 | 3UTR |
| hsa-miR-4505 | NM_005656 | TMPRSS2 | 1.0 | 3UTR |
| hsa-miR-4516 | NM_005656 | TMPRSS2 | 1.0 | 3UTR |
| hsa-miR-4517 | NM_005656 | TMPRSS2 | 1.0 | 3UTR |
| hsa-miR-4525 | NM_005656 | TMPRSS2 | 1.0 | 3UTR |
| hsa-miR-4533 | NM_005656 | TMPRSS2 | 1.0 | 3UTR |
| hsa-miR-378i | NM_005656 | TMPRSS2 | 1.0 | 3UTR |
| hsa-miR-378i | NM_005656 | TMPRSS2 | 1.0 | 3UTR |
| hsa-miR-3978 | NM_005656 | TMPRSS2 | 1.0 | 3UTR |
| hsa-miR-4632-5p | NM_005656 | TMPRSS2 | 1.0 | 3UTR |
| hsa-miR-4646-5p | NM_005656 | TMPRSS2 | 1.0 | 3UTR |
| hsa-miR-4651 | NM_005656 | TMPRSS2 | 1.0 | 3UTR |
| hsa-miR-4653-3p | NM_005656 | TMPRSS2 | 1.0 | 3UTR |
| hsa-miR-4657 | NM_005656 | TMPRSS2 | 1.0 | 3UTR |
| hsa-miR-4669 | NM_005656 | TMPRSS2 | 1.0 | 3UTR |
| hsa-miR-4685-3p | NM_005656 | TMPRSS2 | 1.0 | 3UTR |
| hsa-miR-4688 | NM_005656 | TMPRSS2 | 1.0 | 3UTR |
| hsa-miR-4691-3p | NM_005656 | TMPRSS2 | 1.0 | 3UTR |
| hsa-miR-4695-5p | NM_005656 | TMPRSS2 | 1.0 | 3UTR |
| hsa-miR-4698 | NM_005656 | TMPRSS2 | 1.0 | 3UTR |
| hsa-miR-4700-5p | NM_005656 | TMPRSS2 | 1.0 | 3UTR |
| hsa-miR-4705 | NM_005656 | TMPRSS2 | 1.0 | 3UTR |

| | | | | |
|---|---|---|---|---|
| hsa-miR-4716-3p | NM_005656 | TMPRSS2 | 1.0 | 3UTR |
| hsa-miR-4717-5p | NM_005656 | TMPRSS2 | 1.0 | 3UTR |
| hsa-miR-4723-5p | NM_005656 | TMPRSS2 | 1.0 | 3UTR |
| hsa-miR-4726-5p | NM_005656 | TMPRSS2 | 1.0 | 3UTR |
| hsa-miR-4741 | NM_005656 | TMPRSS2 | 1.0 | 3UTR |
| hsa-miR-4746-5p | NM_005656 | TMPRSS2 | 1.0 | 3UTR |
| hsa-miR-4747-5p | NM_005656 | TMPRSS2 | 1.0 | 3UTR |
| hsa-miR-4747-3p | NM_005656 | TMPRSS2 | 1.0 | 3UTR |
| hsa-miR-4750-5p | NM_005656 | TMPRSS2 | 1.0 | 3UTR |
| hsa-miR-4761-3p | NM_005656 | TMPRSS2 | 1.0 | 3UTR |
| hsa-miR-4763-3p | NM_005656 | TMPRSS2 | 1.0 | 3UTR |
| hsa-miR-4779 | NM_005656 | TMPRSS2 | 1.0 | 3UTR |
| hsa-miR-4436b-3p | NM_005656 | TMPRSS2 | 1.0 | 3UTR |
| hsa-miR-1245b-5p | NM_005656 | TMPRSS2 | 1.0 | 3UTR |
| hsa-miR-4787-5p | NM_005656 | TMPRSS2 | 1.0 | 3UTR |
| hsa-miR-4787-5p | NM_005656 | TMPRSS2 | 1.0 | 3UTR |
| hsa-miR-4788 | NM_005656 | TMPRSS2 | 1.0 | 3UTR |
| hsa-miR-4793-5p | NM_005656 | TMPRSS2 | 1.0 | 3UTR |
| hsa-miR-5002-5p | NM_005656 | TMPRSS2 | 1.0 | 3UTR |
| hsa-miR-5002-3p | NM_005656 | TMPRSS2 | 1.0 | 3UTR |
| hsa-miR-5003-3p | NM_005656 | TMPRSS2 | 1.0 | 3UTR |
| hsa-miR-5004-3p | NM_005656 | TMPRSS2 | 1.0 | 3UTR |
| hsa-miR-5010-5p | NM_005656 | TMPRSS2 | 1.0 | 3UTR |
| hsa-miR-5088-3p | NM_005656 | TMPRSS2 | 1.0 | 3UTR |
| hsa-miR-5093 | NM_005656 | TMPRSS2 | 1.0 | 3UTR |
| hsa-miR-5195-3p | NM_005656 | TMPRSS2 | 1.0 | 3UTR |
| hsa-miR-5195-3p | NM_005656 | TMPRSS2 | 1.0 | 3UTR |
| hsa-miR-4524b-3p | NM_005656 | TMPRSS2 | 1.0 | 3UTR |
| hsa-miR-5581-5p | NM_005656 | TMPRSS2 | 1.0 | 3UTR |
| hsa-miR-5585-3p | NM_005656 | TMPRSS2 | 1.0 | 3UTR |
| hsa-miR-1295b-3p | NM_005656 | TMPRSS2 | 1.0 | 3UTR |
| hsa-miR-5589-3p | NM_005656 | TMPRSS2 | 1.0 | 3UTR |
| hsa-miR-5682 | NM_005656 | TMPRSS2 | 1.0 | 3UTR |
| hsa-miR-5687 | NM_005656 | TMPRSS2 | 1.0 | 3UTR |
| hsa-miR-5691 | NM_005656 | TMPRSS2 | 1.0 | 3UTR |
| hsa-miR-5698 | NM_005656 | TMPRSS2 | 1.0 | 3UTR |
| hsa-miR-1199-3p | NM_005656 | TMPRSS2 | 1.0 | 3UTR |
| hsa-miR-6074 | NM_005656 | TMPRSS2 | 1.0 | 3UTR |
| hsa-miR-6081 | NM_005656 | TMPRSS2 | 1.0 | 3UTR |
| hsa-miR-6086 | NM_005656 | TMPRSS2 | 1.0 | 3UTR |
| hsa-miR-6088 | NM_005656 | TMPRSS2 | 1.0 | 3UTR |
| hsa-miR-6090 | NM_005656 | TMPRSS2 | 1.0 | 3UTR |
| hsa-miR-6133 | NM_005656 | TMPRSS2 | 1.0 | 3UTR |
| hsa-miR-6133 | NM_005656 | TMPRSS2 | 1.0 | 3UTR |
| hsa-miR-6134 | NM_005656 | TMPRSS2 | 1.0 | 3UTR |
| hsa-miR-6165 | NM_005656 | TMPRSS2 | 1.0 | 3UTR |
| hsa-miR-6505-5p | NM_005656 | TMPRSS2 | 1.0 | 3UTR |

| | | | | |
|---|---|---|---|---|
| hsa-miR-6510-5p | NM_005656 | TMPRSS2 | 1.0 | 3UTR |
| hsa-miR-6510-5p | NM_005656 | TMPRSS2 | 1.0 | 3UTR |
| hsa-miR-6514-5p | NM_005656 | TMPRSS2 | 1.0 | 3UTR |
| hsa-miR-6515-5p | NM_005656 | TMPRSS2 | 1.0 | 3UTR |
| hsa-miR-6715b-3p | NM_005656 | TMPRSS2 | 1.0 | 3UTR |
| hsa-miR-6717-5p | NM_005656 | TMPRSS2 | 1.0 | 3UTR |
| hsa-miR-6511b-5p | NM_005656 | TMPRSS2 | 1.0 | 3UTR |
| hsa-miR-6721-5p | NM_005656 | TMPRSS2 | 1.0 | 3UTR |
| hsa-miR-6722-3p | NM_005656 | TMPRSS2 | 1.0 | 3UTR |
| hsa-miR-892c-3p | NM_005656 | TMPRSS2 | 1.0 | 3UTR |
| hsa-miR-6731-3p | NM_005656 | TMPRSS2 | 1.0 | 3UTR |
| hsa-miR-6735-5p | NM_005656 | TMPRSS2 | 1.0 | 3UTR |
| hsa-miR-6740-3p | NM_005656 | TMPRSS2 | 1.0 | 3UTR |
| hsa-miR-6743-5p | NM_005656 | TMPRSS2 | 1.0 | 3UTR |
| hsa-miR-6747-5p | NM_005656 | TMPRSS2 | 1.0 | 3UTR |
| hsa-miR-6747-3p | NM_005656 | TMPRSS2 | 1.0 | 3UTR |
| hsa-miR-6748-5p | NM_005656 | TMPRSS2 | 1.0 | 3UTR |
| hsa-miR-6749-3p | NM_005656 | TMPRSS2 | 1.0 | 3UTR |
| hsa-miR-6749-3p | NM_005656 | TMPRSS2 | 1.0 | 3UTR |
| hsa-miR-6751-3p | NM_005656 | TMPRSS2 | 1.0 | 3UTR |
| hsa-miR-6753-5p | NM_005656 | TMPRSS2 | 1.0 | 3UTR |
| hsa-miR-6753-5p | NM_005656 | TMPRSS2 | 1.0 | 3UTR |
| hsa-miR-6755-3p | NM_005656 | TMPRSS2 | 1.0 | 3UTR |
| hsa-miR-6759-3p | NM_005656 | TMPRSS2 | 1.0 | 3UTR |
| hsa-miR-6765-5p | NM_005656 | TMPRSS2 | 1.0 | 3UTR |
| hsa-miR-6767-5p | NM_005656 | TMPRSS2 | 1.0 | 3UTR |
| hsa-miR-6769a-5p | NM_005656 | TMPRSS2 | 1.0 | 3UTR |
| hsa-miR-6771-5p | NM_005656 | TMPRSS2 | 1.0 | 3UTR |
| hsa-miR-6771-5p | NM_005656 | TMPRSS2 | 1.0 | 3UTR |
| hsa-miR-6772-5p | NM_005656 | TMPRSS2 | 1.0 | 3UTR |
| hsa-miR-6772-5p | NM_005656 | TMPRSS2 | 1.0 | 3UTR |
| hsa-miR-6773-3p | NM_005656 | TMPRSS2 | 1.0 | 3UTR |
| hsa-miR-6774-5p | NM_005656 | TMPRSS2 | 1.0 | 3UTR |
| hsa-miR-6776-3p | NM_005656 | TMPRSS2 | 1.0 | 3UTR |
| hsa-miR-6777-5p | NM_005656 | TMPRSS2 | 1.0 | 3UTR |
| hsa-miR-6778-5p | NM_005656 | TMPRSS2 | 1.0 | 3UTR |
| hsa-miR-6779-5p | NM_005656 | TMPRSS2 | 1.0 | 3UTR |
| hsa-miR-6779-5p | NM_005656 | TMPRSS2 | 1.0 | 3UTR |
| hsa-miR-6780a-5p | NM_005656 | TMPRSS2 | 1.0 | 3UTR |
| hsa-miR-6780a-5p | NM_005656 | TMPRSS2 | 1.0 | 3UTR |
| hsa-miR-6782-5p | NM_005656 | TMPRSS2 | 1.0 | 3UTR |
| hsa-miR-6792-3p | NM_005656 | TMPRSS2 | 1.0 | 3UTR |
| hsa-miR-6793-5p | NM_005656 | TMPRSS2 | 1.0 | 3UTR |
| hsa-miR-6794-5p | NM_005656 | TMPRSS2 | 1.0 | 3UTR |
| hsa-miR-6795-5p | NM_005656 | TMPRSS2 | 1.0 | 3UTR |
| hsa-miR-6795-3p | NM_005656 | TMPRSS2 | 1.0 | 3UTR |
| hsa-miR-6797-3p | NM_005656 | TMPRSS2 | 1.0 | 3UTR |

| | | | | |
|---|---|---|---|---|
| hsa-miR-6798-5p | NM_005656 | TMPRSS2 | 1.0 | 3UTR |
| hsa-miR-6799-5p | NM_005656 | TMPRSS2 | 1.0 | 3UTR |
| hsa-miR-6802-3p | NM_005656 | TMPRSS2 | 1.0 | 3UTR |
| hsa-miR-6804-3p | NM_005656 | TMPRSS2 | 1.0 | 3UTR |
| hsa-miR-6805-5p | NM_005656 | TMPRSS2 | 1.0 | 3UTR |
| hsa-miR-6807-5p | NM_005656 | TMPRSS2 | 1.0 | 3UTR |
| hsa-miR-6807-3p | NM_005656 | TMPRSS2 | 1.0 | 3UTR |
| hsa-miR-6808-5p | NM_005656 | TMPRSS2 | 1.0 | 3UTR |
| hsa-miR-6810-5p | NM_005656 | TMPRSS2 | 1.0 | 3UTR |
| hsa-miR-6813-3p | NM_005656 | TMPRSS2 | 1.0 | 3UTR |
| hsa-miR-6814-3p | NM_005656 | TMPRSS2 | 1.0 | 3UTR |
| hsa-miR-6816-3p | NM_005656 | TMPRSS2 | 1.0 | 3UTR |
| hsa-miR-6818-5p | NM_005656 | TMPRSS2 | 1.0 | 3UTR |
| hsa-miR-6819-5p | NM_005656 | TMPRSS2 | 1.0 | 3UTR |
| hsa-miR-6819-3p | NM_005656 | TMPRSS2 | 1.0 | 3UTR |
| hsa-miR-6822-5p | NM_005656 | TMPRSS2 | 1.0 | 3UTR |
| hsa-miR-6823-5p | NM_005656 | TMPRSS2 | 1.0 | 3UTR |
| hsa-miR-6824-5p | NM_005656 | TMPRSS2 | 1.0 | 3UTR |
| hsa-miR-6824-3p | NM_005656 | TMPRSS2 | 1.0 | 3UTR |
| hsa-miR-6828-5p | NM_005656 | TMPRSS2 | 1.0 | 3UTR |
| hsa-miR-6829-5p | NM_005656 | TMPRSS2 | 1.0 | 3UTR |
| hsa-miR-6832-5p | NM_005656 | TMPRSS2 | 1.0 | 3UTR |
| hsa-miR-6835-5p | NM_005656 | TMPRSS2 | 1.0 | 3UTR |
| hsa-miR-6837-5p | NM_005656 | TMPRSS2 | 1.0 | 3UTR |
| hsa-miR-6841-5p | NM_005656 | TMPRSS2 | 1.0 | 3UTR |
| hsa-miR-6842-5p | NM_005656 | TMPRSS2 | 1.0 | 3UTR |
| hsa-miR-6842-3p | NM_005656 | TMPRSS2 | 1.0 | 3UTR |
| hsa-miR-6843-3p | NM_005656 | TMPRSS2 | 1.0 | 3UTR |
| hsa-miR-6845-3p | NM_005656 | TMPRSS2 | 1.0 | 3UTR |
| hsa-miR-6846-5p | NM_005656 | TMPRSS2 | 1.0 | 3UTR |
| hsa-miR-6849-5p | NM_005656 | TMPRSS2 | 1.0 | 3UTR |
| hsa-miR-6852-5p | NM_005656 | TMPRSS2 | 1.0 | 3UTR |
| hsa-miR-6854-3p | NM_005656 | TMPRSS2 | 1.0 | 3UTR |
| hsa-miR-6769b-5p | NM_005656 | TMPRSS2 | 1.0 | 3UTR |
| hsa-miR-6860 | NM_005656 | TMPRSS2 | 1.0 | 3UTR |
| hsa-miR-6864-5p | NM_005656 | TMPRSS2 | 1.0 | 3UTR |
| hsa-miR-6868-5p | NM_005656 | TMPRSS2 | 1.0 | 3UTR |
| hsa-miR-6871-5p | NM_005656 | TMPRSS2 | 1.0 | 3UTR |
| hsa-miR-6872-5p | NM_005656 | TMPRSS2 | 1.0 | 3UTR |
| hsa-miR-6875-3p | NM_005656 | TMPRSS2 | 1.0 | 3UTR |
| hsa-miR-6876-3p | NM_005656 | TMPRSS2 | 1.0 | 3UTR |
| hsa-miR-6878-3p | NM_005656 | TMPRSS2 | 1.0 | 3UTR |
| hsa-miR-6879-5p | NM_005656 | TMPRSS2 | 1.0 | 3UTR |
| hsa-miR-6879-5p | NM_005656 | TMPRSS2 | 1.0 | 3UTR |
| hsa-miR-6880-3p | NM_005656 | TMPRSS2 | 1.0 | 3UTR |
| hsa-miR-6883-5p | NM_005656 | TMPRSS2 | 1.0 | 3UTR |
| hsa-miR-6883-3p | NM_005656 | TMPRSS2 | 1.0 | 3UTR |

| | | | | |
|---|---|---|---|---|
| hsa-miR-6884-5p | NM_005656 | TMPRSS2 | 1.0 | 3UTR |
| hsa-miR-6888-5p | NM_005656 | TMPRSS2 | 1.0 | 3UTR |
| hsa-miR-6888-5p | NM_005656 | TMPRSS2 | 1.0 | 3UTR |
| hsa-miR-6890-5p | NM_005656 | TMPRSS2 | 1.0 | 3UTR |
| hsa-miR-6893-5p | NM_005656 | TMPRSS2 | 1.0 | 3UTR |
| hsa-miR-6894-5p | NM_005656 | TMPRSS2 | 1.0 | 3UTR |
| hsa-miR-7107-5p | NM_005656 | TMPRSS2 | 1.0 | 3UTR |
| hsa-miR-7110-3p | NM_005656 | TMPRSS2 | 1.0 | 3UTR |
| hsa-miR-7112-5p | NM_005656 | TMPRSS2 | 1.0 | 3UTR |
| hsa-miR-7112-3p | NM_005656 | TMPRSS2 | 1.0 | 3UTR |
| hsa-miR-7113-5p | NM_005656 | TMPRSS2 | 1.0 | 3UTR |
| hsa-miR-7114-3p | NM_005656 | TMPRSS2 | 1.0 | 3UTR |
| hsa-miR-7151-3p | NM_005656 | TMPRSS2 | 1.0 | 3UTR |
| hsa-miR-7151-3p | NM_005656 | TMPRSS2 | 1.0 | 3UTR |
| hsa-miR-7158-5p | NM_005656 | TMPRSS2 | 1.0 | 3UTR |
| hsa-miR-7702 | NM_005656 | TMPRSS2 | 1.0 | 3UTR |
| hsa-miR-7843-5p | NM_005656 | TMPRSS2 | 1.0 | 3UTR |
| hsa-miR-7846-3p | NM_005656 | TMPRSS2 | 1.0 | 3UTR |
| hsa-miR-7846-3p | NM_005656 | TMPRSS2 | 1.0 | 3UTR |
| hsa-miR-7847-3p | NM_005656 | TMPRSS2 | 1.0 | 3UTR |
| hsa-miR-7849-3p | NM_005656 | TMPRSS2 | 1.0 | 3UTR |
| hsa-miR-7854-3p | NM_005656 | TMPRSS2 | 1.0 | 3UTR |
| hsa-miR-8052 | NM_005656 | TMPRSS2 | 1.0 | 3UTR |
| hsa-miR-8063 | NM_005656 | TMPRSS2 | 1.0 | 3UTR |
| hsa-miR-8069 | NM_005656 | TMPRSS2 | 1.0 | 3UTR |
| hsa-miR-8072 | NM_005656 | TMPRSS2 | 1.0 | 3UTR |
| hsa-miR-8078 | NM_005656 | TMPRSS2 | 1.0 | 3UTR |
| hsa-miR-8088 | NM_005656 | TMPRSS2 | 1.0 | 3UTR |
| hsa-miR-9985 | NM_005656 | TMPRSS2 | 1.0 | 3UTR |
| hsa-miR-10394-3p | NM_005656 | TMPRSS2 | 1.0 | 3UTR |
| hsa-miR-10397-5p | NM_005656 | TMPRSS2 | 1.0 | 3UTR |
| hsa-miR-10398-5p | NM_005656 | TMPRSS2 | 1.0 | 3UTR |
| hsa-miR-10526-3p | NM_005656 | TMPRSS2 | 1.0 | 3UTR |
| hsa-miR-11181-3p | NM_005656 | TMPRSS2 | 1.0 | 3UTR |
| hsa-miR-3059-3p | NM_005656 | TMPRSS2 | 1.0 | 3UTR |
| hsa-miR-3059-3p | NM_005656 | TMPRSS2 | 1.0 | 3UTR |
| hsa-miR-3085-5p | NM_005656 | TMPRSS2 | 1.0 | 3UTR |
| hsa-miR-3085-3p | NM_005656 | TMPRSS2 | 1.0 | 3UTR |
| hsa-miR-6529-5p | NM_005656 | TMPRSS2 | 1.0 | 3UTR |
| hsa-miR-9851-5p | NM_005656 | TMPRSS2 | 1.0 | 3UTR |
| hsa-miR-9851-3p | NM_005656 | TMPRSS2 | 1.0 | 3UTR |
| hsa-miR-12119 | NM_005656 | TMPRSS2 | 1.0 | 3UTR |
| hsa-miR-12119 | NM_005656 | TMPRSS2 | 1.0 | 3UTR |
| hsa-miR-12120 | NM_005656 | TMPRSS2 | 1.0 | 3UTR |
| hsa-miR-12128 | NM_005656 | TMPRSS2 | 1.0 | 3UTR |
| hsa-miR-12131 | NM_005656 | TMPRSS2 | 1.0 | 3UTR |
| hsa-let-7b-5p | NM_001135099 | TMPRSS2 | 1.0 | 3UTR |

| | | | | |
|---|---|---|---|---|
| hsa-let-7c-5p | NM_001135099 | TMPRSS2 | 1.0 | 3UTR |
| hsa-miR-23a-5p | NM_001135099 | TMPRSS2 | 1.0 | 3UTR |
| hsa-miR-26a-5p | NM_001135099 | TMPRSS2 | 1.0 | 3UTR |
| hsa-miR-26b-3p | NM_001135099 | TMPRSS2 | 1.0 | 3UTR |
| hsa-miR-28-5p | NM_001135099 | TMPRSS2 | 1.0 | 3UTR |
| hsa-miR-92a-1-5p | NM_001135099 | TMPRSS2 | 1.0 | 3UTR |
| hsa-miR-93-5p | NM_001135099 | TMPRSS2 | 1.0 | 3UTR |
| hsa-miR-95-3p | NM_001135099 | TMPRSS2 | 1.0 | 3UTR |
| hsa-miR-96-5p | NM_001135099 | TMPRSS2 | 1.0 | 3UTR |
| hsa-miR-101-5p | NM_001135099 | TMPRSS2 | 1.0 | 3UTR |
| hsa-miR-103a-2-5p | NM_001135099 | TMPRSS2 | 1.0 | 3UTR |
| hsa-miR-103a-3p | NM_001135099 | TMPRSS2 | 1.0 | 3UTR |
| hsa-miR-103a-1-5p | NM_001135099 | TMPRSS2 | 1.0 | 3UTR |
| hsa-miR-107 | NM_001135099 | TMPRSS2 | 1.0 | 3UTR |
| hsa-miR-197-3p | NM_001135099 | TMPRSS2 | 1.0 | 3UTR |
| hsa-miR-198 | NM_001135099 | TMPRSS2 | 1.0 | 3UTR |
| hsa-miR-129-1-3p | NM_001135099 | TMPRSS2 | 1.0 | 3UTR |
| hsa-miR-30d-5p | NM_001135099 | TMPRSS2 | 1.0 | 3UTR |
| hsa-miR-34a-5p | NM_001135099 | TMPRSS2 | 1.0 | 3UTR |
| hsa-miR-182-5p | NM_001135099 | TMPRSS2 | 1.0 | 3UTR |
| hsa-miR-183-5p | NM_001135099 | TMPRSS2 | 1.0 | 3UTR |
| hsa-miR-204-3p | NM_001135099 | TMPRSS2 | 1.0 | 3UTR |
| hsa-miR-218-1-3p | NM_001135099 | TMPRSS2 | 1.0 | 3UTR |
| hsa-miR-200b-5p | NM_001135099 | TMPRSS2 | 1.0 | 3UTR |
| hsa-let-7g-5p | NM_001135099 | TMPRSS2 | 1.0 | 3UTR |
| hsa-let-7i-5p | NM_001135099 | TMPRSS2 | 1.0 | 3UTR |
| hsa-miR-27b-5p | NM_001135099 | TMPRSS2 | 1.0 | 3UTR |
| hsa-miR-30b-3p | NM_001135099 | TMPRSS2 | 1.0 | 3UTR |
| hsa-miR-124-5p | NM_001135099 | TMPRSS2 | 1.0 | 3UTR |
| hsa-miR-135a-2-3p | NM_001135099 | TMPRSS2 | 1.0 | 3UTR |
| hsa-miR-152-3p | NM_001135099 | TMPRSS2 | 1.0 | 3UTR |
| hsa-miR-191-5p | NM_001135099 | TMPRSS2 | 1.0 | 3UTR |
| hsa-miR-9-5p | NM_001135099 | TMPRSS2 | 1.0 | 3UTR |
| hsa-miR-134-5p | NM_001135099 | TMPRSS2 | 1.0 | 3UTR |
| hsa-miR-149-5p | NM_001135099 | TMPRSS2 | 1.0 | 3UTR |
| hsa-miR-185-5p | NM_001135099 | TMPRSS2 | 1.0 | 3UTR |
| hsa-miR-188-5p | NM_001135099 | TMPRSS2 | 1.0 | 3UTR |
| hsa-miR-193a-5p | NM_001135099 | TMPRSS2 | 1.0 | 3UTR |
| hsa-miR-320a-3p | NM_001135099 | TMPRSS2 | 1.0 | 3UTR |
| hsa-miR-200c-5p | NM_001135099 | TMPRSS2 | 1.0 | 3UTR |
| hsa-miR-106b-3p | NM_001135099 | TMPRSS2 | 1.0 | 3UTR |
| hsa-miR-361-5p | NM_001135099 | TMPRSS2 | 1.0 | 3UTR |
| hsa-miR-302c-5p | NM_001135099 | TMPRSS2 | 1.0 | 3UTR |
| hsa-miR-379-5p | NM_001135099 | TMPRSS2 | 1.0 | 3UTR |
| hsa-miR-328-5p | NM_001135099 | TMPRSS2 | 1.0 | 3UTR |
| hsa-miR-151a-5p | NM_001135099 | TMPRSS2 | 1.0 | 3UTR |
| hsa-miR-151a-3p | NM_001135099 | TMPRSS2 | 1.0 | 3UTR |

| | | | | |
|---|---|---|---|---|
| hsa-miR-331-5p | NM_001135099 | TMPRSS2 | 1.0 | 3UTR |
| hsa-miR-331-3p | NM_001135099 | TMPRSS2 | 1.0 | 3UTR |
| hsa-miR-324-5p | NM_001135099 | TMPRSS2 | 1.0 | 3UTR |
| hsa-miR-324-3p | NM_001135099 | TMPRSS2 | 1.0 | 3UTR |
| hsa-miR-339-5p | NM_001135099 | TMPRSS2 | 1.0 | 3UTR |
| hsa-miR-422a | NM_001135099 | TMPRSS2 | 1.0 | 3UTR |
| hsa-miR-423-5p | NM_001135099 | TMPRSS2 | 1.0 | 3UTR |
| hsa-miR-423-5p | NM_001135099 | TMPRSS2 | 1.0 | 3UTR |
| hsa-miR-423-3p | NM_001135099 | TMPRSS2 | 1.0 | 3UTR |
| hsa-miR-20b-3p | NM_001135099 | TMPRSS2 | 1.0 | 3UTR |
| hsa-miR-329-5p | NM_001135099 | TMPRSS2 | 1.0 | 3UTR |
| hsa-miR-483-3p | NM_001135099 | TMPRSS2 | 1.0 | 3UTR |
| hsa-miR-491-5p | NM_001135099 | TMPRSS2 | 1.0 | 3UTR |
| hsa-miR-146b-3p | NM_001135099 | TMPRSS2 | 1.0 | 3UTR |
| hsa-miR-432-5p | NM_001135099 | TMPRSS2 | 1.0 | 3UTR |
| hsa-miR-432-5p | NM_001135099 | TMPRSS2 | 1.0 | 3UTR |
| hsa-miR-193b-5p | NM_001135099 | TMPRSS2 | 1.0 | 3UTR |
| hsa-miR-181d-5p | NM_001135099 | TMPRSS2 | 1.0 | 3UTR |
| hsa-miR-526b-5p | NM_001135099 | TMPRSS2 | 1.0 | 3UTR |
| hsa-miR-520b-5p | NM_001135099 | TMPRSS2 | 1.0 | 3UTR |
| hsa-miR-520c-3p | NM_001135099 | TMPRSS2 | 1.0 | 3UTR |
| hsa-miR-519a-2-5p | NM_001135099 | TMPRSS2 | 1.0 | 3UTR |
| hsa-miR-501-5p | NM_001135099 | TMPRSS2 | 1.0 | 3UTR |
| hsa-miR-502-5p | NM_001135099 | TMPRSS2 | 1.0 | 3UTR |
| hsa-miR-502-3p | NM_001135099 | TMPRSS2 | 1.0 | 3UTR |
| hsa-miR-503-3p | NM_001135099 | TMPRSS2 | 1.0 | 3UTR |
| hsa-miR-504-3p | NM_001135099 | TMPRSS2 | 1.0 | 3UTR |
| hsa-miR-505-5p | NM_001135099 | TMPRSS2 | 1.0 | 3UTR |
| hsa-miR-505-5p | NM_001135099 | TMPRSS2 | 1.0 | 3UTR |
| hsa-miR-505-3p | NM_001135099 | TMPRSS2 | 1.0 | 3UTR |
| hsa-miR-507 | NM_001135099 | TMPRSS2 | 1.0 | 3UTR |
| hsa-miR-508-5p | NM_001135099 | TMPRSS2 | 1.0 | 3UTR |
| hsa-miR-509-3p | NM_001135099 | TMPRSS2 | 1.0 | 3UTR |
| hsa-miR-514a-5p | NM_001135099 | TMPRSS2 | 1.0 | 3UTR |
| hsa-miR-455-5p | NM_001135099 | TMPRSS2 | 1.0 | 3UTR |
| hsa-miR-455-3p | NM_001135099 | TMPRSS2 | 1.0 | 3UTR |
| hsa-miR-562 | NM_001135099 | TMPRSS2 | 1.0 | 3UTR |
| hsa-miR-575 | NM_001135099 | TMPRSS2 | 1.0 | 3UTR |
| hsa-miR-584-3p | NM_001135099 | TMPRSS2 | 1.0 | 3UTR |
| hsa-miR-548b-3p | NM_001135099 | TMPRSS2 | 1.0 | 3UTR |
| hsa-miR-589-5p | NM_001135099 | TMPRSS2 | 1.0 | 3UTR |
| hsa-miR-550a-5p | NM_001135099 | TMPRSS2 | 1.0 | 3UTR |
| hsa-miR-550a-5p | NM_001135099 | TMPRSS2 | 1.0 | 3UTR |
| hsa-miR-602 | NM_001135099 | TMPRSS2 | 1.0 | 3UTR |
| hsa-miR-608 | NM_001135099 | TMPRSS2 | 1.0 | 3UTR |
| hsa-miR-613 | NM_001135099 | TMPRSS2 | 1.0 | 3UTR |
| hsa-miR-614 | NM_001135099 | TMPRSS2 | 1.0 | 3UTR |

| | | | | |
|---|---|---|---|---|
| hsa-miR-619-5p | NM_001135099 | TMPRSS2 | 1.0 | 3UTR |
| hsa-miR-622 | NM_001135099 | TMPRSS2 | 1.0 | 3UTR |
| hsa-miR-622 | NM_001135099 | TMPRSS2 | 1.0 | 3UTR |
| hsa-miR-33b-3p | NM_001135099 | TMPRSS2 | 1.0 | 3UTR |
| hsa-miR-642a-5p | NM_001135099 | TMPRSS2 | 1.0 | 3UTR |
| hsa-miR-647 | NM_001135099 | TMPRSS2 | 1.0 | 3UTR |
| hsa-miR-648 | NM_001135099 | TMPRSS2 | 1.0 | 3UTR |
| hsa-miR-650 | NM_001135099 | TMPRSS2 | 1.0 | 3UTR |
| hsa-miR-449b-5p | NM_001135099 | TMPRSS2 | 1.0 | 3UTR |
| hsa-miR-671-5p | NM_001135099 | TMPRSS2 | 1.0 | 3UTR |
| hsa-miR-671-3p | NM_001135099 | TMPRSS2 | 1.0 | 3UTR |
| hsa-miR-550a-3-5p | NM_001135099 | TMPRSS2 | 1.0 | 3UTR |
| hsa-miR-767-5p | NM_001135099 | TMPRSS2 | 1.0 | 3UTR |
| hsa-miR-767-5p | NM_001135099 | TMPRSS2 | 1.0 | 3UTR |
| hsa-miR-1224-3p | NM_001135099 | TMPRSS2 | 1.0 | 3UTR |
| hsa-miR-320b | NM_001135099 | TMPRSS2 | 1.0 | 3UTR |
| hsa-miR-1271-5p | NM_001135099 | TMPRSS2 | 1.0 | 3UTR |
| hsa-miR-1271-3p | NM_001135099 | TMPRSS2 | 1.0 | 3UTR |
| hsa-miR-449c-3p | NM_001135099 | TMPRSS2 | 1.0 | 3UTR |
| hsa-miR-769-5p | NM_001135099 | TMPRSS2 | 1.0 | 3UTR |
| hsa-miR-766-5p | NM_001135099 | TMPRSS2 | 1.0 | 3UTR |
| hsa-miR-675-5p | NM_001135099 | TMPRSS2 | 1.0 | 3UTR |
| hsa-miR-890 | NM_001135099 | TMPRSS2 | 1.0 | 3UTR |
| hsa-miR-875-3p | NM_001135099 | TMPRSS2 | 1.0 | 3UTR |
| hsa-miR-708-5p | NM_001135099 | TMPRSS2 | 1.0 | 3UTR |
| hsa-miR-665 | NM_001135099 | TMPRSS2 | 1.0 | 3UTR |
| hsa-miR-921 | NM_001135099 | TMPRSS2 | 1.0 | 3UTR |
| hsa-miR-933 | NM_001135099 | TMPRSS2 | 1.0 | 3UTR |
| hsa-miR-936 | NM_001135099 | TMPRSS2 | 1.0 | 3UTR |
| hsa-miR-939-5p | NM_001135099 | TMPRSS2 | 1.0 | 3UTR |
| hsa-miR-1182 | NM_001135099 | TMPRSS2 | 1.0 | 3UTR |
| hsa-miR-1227-3p | NM_001135099 | TMPRSS2 | 1.0 | 3UTR |
| hsa-miR-1229-5p | NM_001135099 | TMPRSS2 | 1.0 | 3UTR |
| hsa-miR-1229-3p | NM_001135099 | TMPRSS2 | 1.0 | 3UTR |
| hsa-miR-1234-3p | NM_001135099 | TMPRSS2 | 1.0 | 3UTR |
| hsa-miR-1234-3p | NM_001135099 | TMPRSS2 | 1.0 | 3UTR |
| hsa-miR-1236-5p | NM_001135099 | TMPRSS2 | 1.0 | 3UTR |
| hsa-miR-1237-3p | NM_001135099 | TMPRSS2 | 1.0 | 3UTR |
| hsa-miR-1238-3p | NM_001135099 | TMPRSS2 | 1.0 | 3UTR |
| hsa-miR-1207-3p | NM_001135099 | TMPRSS2 | 1.0 | 3UTR |
| hsa-miR-1243 | NM_001135099 | TMPRSS2 | 1.0 | 3UTR |
| hsa-miR-1250-5p | NM_001135099 | TMPRSS2 | 1.0 | 3UTR |
| hsa-miR-1253 | NM_001135099 | TMPRSS2 | 1.0 | 3UTR |
| hsa-miR-1265 | NM_001135099 | TMPRSS2 | 1.0 | 3UTR |
| hsa-miR-1266-5p | NM_001135099 | TMPRSS2 | 1.0 | 3UTR |
| hsa-miR-1275 | NM_001135099 | TMPRSS2 | 1.0 | 3UTR |
| hsa-miR-1255b-2-3p | NM_001135099 | TMPRSS2 | 1.0 | 3UTR |

| | | | | |
|---|---|---|---|---|
| hsa-miR-1321 | NM_001135099 | TMPRSS2 | 1.0 | 3UTR |
| hsa-miR-1324 | NM_001135099 | TMPRSS2 | 1.0 | 3UTR |
| hsa-miR-103b | NM_001135099 | TMPRSS2 | 1.0 | 3UTR |
| hsa-miR-1825 | NM_001135099 | TMPRSS2 | 1.0 | 3UTR |
| hsa-miR-1908-5p | NM_001135099 | TMPRSS2 | 1.0 | 3UTR |
| hsa-miR-1909-5p | NM_001135099 | TMPRSS2 | 1.0 | 3UTR |
| hsa-miR-1910-3p | NM_001135099 | TMPRSS2 | 1.0 | 3UTR |
| hsa-miR-1912-3p | NM_001135099 | TMPRSS2 | 1.0 | 3UTR |
| hsa-miR-2116-5p | NM_001135099 | TMPRSS2 | 1.0 | 3UTR |
| hsa-miR-2277-5p | NM_001135099 | TMPRSS2 | 1.0 | 3UTR |
| hsa-miR-2682-3p | NM_001135099 | TMPRSS2 | 1.0 | 3UTR |
| hsa-miR-3126-5p | NM_001135099 | TMPRSS2 | 1.0 | 3UTR |
| hsa-miR-3129-5p | NM_001135099 | TMPRSS2 | 1.0 | 3UTR |
| hsa-miR-3131 | NM_001135099 | TMPRSS2 | 1.0 | 3UTR |
| hsa-miR-3137 | NM_001135099 | TMPRSS2 | 1.0 | 3UTR |
| hsa-miR-3138 | NM_001135099 | TMPRSS2 | 1.0 | 3UTR |
| hsa-miR-3144-5p | NM_001135099 | TMPRSS2 | 1.0 | 3UTR |
| hsa-miR-3153 | NM_001135099 | TMPRSS2 | 1.0 | 3UTR |
| hsa-miR-3158-5p | NM_001135099 | TMPRSS2 | 1.0 | 3UTR |
| hsa-miR-3158-3p | NM_001135099 | TMPRSS2 | 1.0 | 3UTR |
| hsa-miR-3166 | NM_001135099 | TMPRSS2 | 1.0 | 3UTR |
| hsa-miR-1260b | NM_001135099 | TMPRSS2 | 1.0 | 3UTR |
| hsa-miR-3170 | NM_001135099 | TMPRSS2 | 1.0 | 3UTR |
| hsa-miR-3179 | NM_001135099 | TMPRSS2 | 1.0 | 3UTR |
| hsa-miR-3184-3p | NM_001135099 | TMPRSS2 | 1.0 | 3UTR |
| hsa-miR-3187-3p | NM_001135099 | TMPRSS2 | 1.0 | 3UTR |
| hsa-miR-3190-3p | NM_001135099 | TMPRSS2 | 1.0 | 3UTR |
| hsa-miR-3192-5p | NM_001135099 | TMPRSS2 | 1.0 | 3UTR |
| hsa-miR-3193 | NM_001135099 | TMPRSS2 | 1.0 | 3UTR |
| hsa-miR-3194-5p | NM_001135099 | TMPRSS2 | 1.0 | 3UTR |
| hsa-miR-4300 | NM_001135099 | TMPRSS2 | 1.0 | 3UTR |
| hsa-miR-4314 | NM_001135099 | TMPRSS2 | 1.0 | 3UTR |
| hsa-miR-4318 | NM_001135099 | TMPRSS2 | 1.0 | 3UTR |
| hsa-miR-4259 | NM_001135099 | TMPRSS2 | 1.0 | 3UTR |
| hsa-miR-4260 | NM_001135099 | TMPRSS2 | 1.0 | 3UTR |
| hsa-miR-4327 | NM_001135099 | TMPRSS2 | 1.0 | 3UTR |
| hsa-miR-4327 | NM_001135099 | TMPRSS2 | 1.0 | 3UTR |
| hsa-miR-4269 | NM_001135099 | TMPRSS2 | 1.0 | 3UTR |
| hsa-miR-4283 | NM_001135099 | TMPRSS2 | 1.0 | 3UTR |
| hsa-miR-3619-5p | NM_001135099 | TMPRSS2 | 1.0 | 3UTR |
| hsa-miR-3620-5p | NM_001135099 | TMPRSS2 | 1.0 | 3UTR |
| hsa-miR-3652 | NM_001135099 | TMPRSS2 | 1.0 | 3UTR |
| hsa-miR-3655 | NM_001135099 | TMPRSS2 | 1.0 | 3UTR |
| hsa-miR-3664-3p | NM_001135099 | TMPRSS2 | 1.0 | 3UTR |
| hsa-miR-3665 | NM_001135099 | TMPRSS2 | 1.0 | 3UTR |
| hsa-miR-3667-3p | NM_001135099 | TMPRSS2 | 1.0 | 3UTR |
| hsa-miR-3679-5p | NM_001135099 | TMPRSS2 | 1.0 | 3UTR |

| | | | | |
|---|---|---|---|---|
| hsa-miR-3682-3p | NM_001135099 | TMPRSS2 | 1.0 | 3UTR |
| hsa-miR-3907 | NM_001135099 | TMPRSS2 | 1.0 | 3UTR |
| hsa-miR-3911 | NM_001135099 | TMPRSS2 | 1.0 | 3UTR |
| hsa-miR-3913-3p | NM_001135099 | TMPRSS2 | 1.0 | 3UTR |
| hsa-miR-3926 | NM_001135099 | TMPRSS2 | 1.0 | 3UTR |
| hsa-miR-3928-3p | NM_001135099 | TMPRSS2 | 1.0 | 3UTR |
| hsa-miR-3936 | NM_001135099 | TMPRSS2 | 1.0 | 3UTR |
| hsa-miR-3939 | NM_001135099 | TMPRSS2 | 1.0 | 3UTR |
| hsa-miR-3940-5p | NM_001135099 | TMPRSS2 | 1.0 | 3UTR |
| hsa-miR-3944-5p | NM_001135099 | TMPRSS2 | 1.0 | 3UTR |
| hsa-miR-642b-3p | NM_001135099 | TMPRSS2 | 1.0 | 3UTR |
| hsa-miR-550b-2-5p | NM_001135099 | TMPRSS2 | 1.0 | 3UTR |
| hsa-miR-4418 | NM_001135099 | TMPRSS2 | 1.0 | 3UTR |
| hsa-miR-378f | NM_001135099 | TMPRSS2 | 1.0 | 3UTR |
| hsa-miR-4425 | NM_001135099 | TMPRSS2 | 1.0 | 3UTR |
| hsa-miR-4428 | NM_001135099 | TMPRSS2 | 1.0 | 3UTR |
| hsa-miR-4429 | NM_001135099 | TMPRSS2 | 1.0 | 3UTR |
| hsa-miR-4433a-3p | NM_001135099 | TMPRSS2 | 1.0 | 3UTR |
| hsa-miR-4436a | NM_001135099 | TMPRSS2 | 1.0 | 3UTR |
| hsa-miR-4439 | NM_001135099 | TMPRSS2 | 1.0 | 3UTR |
| hsa-miR-4441 | NM_001135099 | TMPRSS2 | 1.0 | 3UTR |
| hsa-miR-4446-3p | NM_001135099 | TMPRSS2 | 1.0 | 3UTR |
| hsa-miR-4449 | NM_001135099 | TMPRSS2 | 1.0 | 3UTR |
| hsa-miR-4456 | NM_001135099 | TMPRSS2 | 1.0 | 3UTR |
| hsa-miR-4458 | NM_001135099 | TMPRSS2 | 1.0 | 3UTR |
| hsa-miR-4469 | NM_001135099 | TMPRSS2 | 1.0 | 3UTR |
| hsa-miR-4481 | NM_001135099 | TMPRSS2 | 1.0 | 3UTR |
| hsa-miR-4484 | NM_001135099 | TMPRSS2 | 1.0 | 3UTR |
| hsa-miR-4489 | NM_001135099 | TMPRSS2 | 1.0 | 3UTR |
| hsa-miR-4501 | NM_001135099 | TMPRSS2 | 1.0 | 3UTR |
| hsa-miR-4505 | NM_001135099 | TMPRSS2 | 1.0 | 3UTR |
| hsa-miR-4516 | NM_001135099 | TMPRSS2 | 1.0 | 3UTR |
| hsa-miR-4519 | NM_001135099 | TMPRSS2 | 1.0 | 3UTR |
| hsa-miR-4533 | NM_001135099 | TMPRSS2 | 1.0 | 3UTR |
| hsa-miR-378i | NM_001135099 | TMPRSS2 | 1.0 | 3UTR |
| hsa-miR-3978 | NM_001135099 | TMPRSS2 | 1.0 | 3UTR |
| hsa-miR-4638-3p | NM_001135099 | TMPRSS2 | 1.0 | 3UTR |
| hsa-miR-4646-5p | NM_001135099 | TMPRSS2 | 1.0 | 3UTR |
| hsa-miR-4653-3p | NM_001135099 | TMPRSS2 | 1.0 | 3UTR |
| hsa-miR-4653-3p | NM_001135099 | TMPRSS2 | 1.0 | 3UTR |
| hsa-miR-4654 | NM_001135099 | TMPRSS2 | 1.0 | 3UTR |
| hsa-miR-4655-5p | NM_001135099 | TMPRSS2 | 1.0 | 3UTR |
| hsa-miR-4667-5p | NM_001135099 | TMPRSS2 | 1.0 | 3UTR |
| hsa-miR-4685-3p | NM_001135099 | TMPRSS2 | 1.0 | 3UTR |
| hsa-miR-4688 | NM_001135099 | TMPRSS2 | 1.0 | 3UTR |
| hsa-miR-4698 | NM_001135099 | TMPRSS2 | 1.0 | 3UTR |
| hsa-miR-4700-5p | NM_001135099 | TMPRSS2 | 1.0 | 3UTR |

| | | | | |
|---|---|---|---|---|
| hsa-miR-4716-3p | NM_001135099 | TMPRSS2 | 1.0 | 3UTR |
| hsa-miR-4716-3p | NM_001135099 | TMPRSS2 | 1.0 | 3UTR |
| hsa-miR-4717-5p | NM_001135099 | TMPRSS2 | 1.0 | 3UTR |
| hsa-miR-4722-5p | NM_001135099 | TMPRSS2 | 1.0 | 3UTR |
| hsa-miR-4723-5p | NM_001135099 | TMPRSS2 | 1.0 | 3UTR |
| hsa-miR-4725-5p | NM_001135099 | TMPRSS2 | 1.0 | 3UTR |
| hsa-miR-4739 | NM_001135099 | TMPRSS2 | 1.0 | 3UTR |
| hsa-miR-4741 | NM_001135099 | TMPRSS2 | 1.0 | 3UTR |
| hsa-miR-4746-3p | NM_001135099 | TMPRSS2 | 1.0 | 3UTR |
| hsa-miR-4749-3p | NM_001135099 | TMPRSS2 | 1.0 | 3UTR |
| hsa-miR-4750-5p | NM_001135099 | TMPRSS2 | 1.0 | 3UTR |
| hsa-miR-4760-3p | NM_001135099 | TMPRSS2 | 1.0 | 3UTR |
| hsa-miR-4761-3p | NM_001135099 | TMPRSS2 | 1.0 | 3UTR |
| hsa-miR-4764-3p | NM_001135099 | TMPRSS2 | 1.0 | 3UTR |
| hsa-miR-4776-5p | NM_001135099 | TMPRSS2 | 1.0 | 3UTR |
| hsa-miR-4436b-3p | NM_001135099 | TMPRSS2 | 1.0 | 3UTR |
| hsa-miR-4436b-3p | NM_001135099 | TMPRSS2 | 1.0 | 3UTR |
| hsa-miR-2467-3p | NM_001135099 | TMPRSS2 | 1.0 | 3UTR |
| hsa-miR-4799-3p | NM_001135099 | TMPRSS2 | 1.0 | 3UTR |
| hsa-miR-5001-3p | NM_001135099 | TMPRSS2 | 1.0 | 3UTR |
| hsa-miR-5002-3p | NM_001135099 | TMPRSS2 | 1.0 | 3UTR |
| hsa-miR-5003-3p | NM_001135099 | TMPRSS2 | 1.0 | 3UTR |
| hsa-miR-5004-3p | NM_001135099 | TMPRSS2 | 1.0 | 3UTR |
| hsa-miR-548ao-3p | NM_001135099 | TMPRSS2 | 1.0 | 3UTR |
| hsa-miR-5006-5p | NM_001135099 | TMPRSS2 | 1.0 | 3UTR |
| hsa-miR-5006-5p | NM_001135099 | TMPRSS2 | 1.0 | 3UTR |
| hsa-miR-5010-5p | NM_001135099 | TMPRSS2 | 1.0 | 3UTR |
| hsa-miR-5088-5p | NM_001135099 | TMPRSS2 | 1.0 | 3UTR |
| hsa-miR-5093 | NM_001135099 | TMPRSS2 | 1.0 | 3UTR |
| hsa-miR-5190 | NM_001135099 | TMPRSS2 | 1.0 | 3UTR |
| hsa-miR-5192 | NM_001135099 | TMPRSS2 | 1.0 | 3UTR |
| hsa-miR-5195-3p | NM_001135099 | TMPRSS2 | 1.0 | 3UTR |
| hsa-miR-5196-5p | NM_001135099 | TMPRSS2 | 1.0 | 3UTR |
| hsa-miR-4524b-3p | NM_001135099 | TMPRSS2 | 1.0 | 3UTR |
| hsa-miR-5571-3p | NM_001135099 | TMPRSS2 | 1.0 | 3UTR |
| hsa-miR-5572 | NM_001135099 | TMPRSS2 | 1.0 | 3UTR |
| hsa-miR-5581-5p | NM_001135099 | TMPRSS2 | 1.0 | 3UTR |
| hsa-miR-548au-3p | NM_001135099 | TMPRSS2 | 1.0 | 3UTR |
| hsa-miR-1295b-3p | NM_001135099 | TMPRSS2 | 1.0 | 3UTR |
| hsa-miR-1295b-3p | NM_001135099 | TMPRSS2 | 1.0 | 3UTR |
| hsa-miR-5589-3p | NM_001135099 | TMPRSS2 | 1.0 | 3UTR |
| hsa-miR-5682 | NM_001135099 | TMPRSS2 | 1.0 | 3UTR |
| hsa-miR-5691 | NM_001135099 | TMPRSS2 | 1.0 | 3UTR |
| hsa-miR-5698 | NM_001135099 | TMPRSS2 | 1.0 | 3UTR |
| hsa-miR-5704 | NM_001135099 | TMPRSS2 | 1.0 | 3UTR |
| hsa-miR-1199-3p | NM_001135099 | TMPRSS2 | 1.0 | 3UTR |
| hsa-miR-6068 | NM_001135099 | TMPRSS2 | 1.0 | 3UTR |

| | | | | |
|---|---|---|---|---|
| hsa-miR-6070 | NM_001135099 | TMPRSS2 | 1.0 | 3UTR |
| hsa-miR-6074 | NM_001135099 | TMPRSS2 | 1.0 | 3UTR |
| hsa-miR-6077 | NM_001135099 | TMPRSS2 | 1.0 | 3UTR |
| hsa-miR-6086 | NM_001135099 | TMPRSS2 | 1.0 | 3UTR |
| hsa-miR-6088 | NM_001135099 | TMPRSS2 | 1.0 | 3UTR |
| hsa-miR-6090 | NM_001135099 | TMPRSS2 | 1.0 | 3UTR |
| hsa-miR-6133 | NM_001135099 | TMPRSS2 | 1.0 | 3UTR |
| hsa-miR-6134 | NM_001135099 | TMPRSS2 | 1.0 | 3UTR |
| hsa-miR-6165 | NM_001135099 | TMPRSS2 | 1.0 | 3UTR |
| hsa-miR-548ay-3p | NM_001135099 | TMPRSS2 | 1.0 | 3UTR |
| hsa-miR-6500-5p | NM_001135099 | TMPRSS2 | 1.0 | 3UTR |
| hsa-miR-6505-5p | NM_001135099 | TMPRSS2 | 1.0 | 3UTR |
| hsa-miR-6510-5p | NM_001135099 | TMPRSS2 | 1.0 | 3UTR |
| hsa-miR-6515-5p | NM_001135099 | TMPRSS2 | 1.0 | 3UTR |
| hsa-miR-6515-5p | NM_001135099 | TMPRSS2 | 1.0 | 3UTR |
| hsa-miR-6717-5p | NM_001135099 | TMPRSS2 | 1.0 | 3UTR |
| hsa-miR-6511b-3p | NM_001135099 | TMPRSS2 | 1.0 | 3UTR |
| hsa-miR-6720-3p | NM_001135099 | TMPRSS2 | 1.0 | 3UTR |
| hsa-miR-892c-3p | NM_001135099 | TMPRSS2 | 1.0 | 3UTR |
| hsa-miR-6727-5p | NM_001135099 | TMPRSS2 | 1.0 | 3UTR |
| hsa-miR-6728-3p | NM_001135099 | TMPRSS2 | 1.0 | 3UTR |
| hsa-miR-6729-5p | NM_001135099 | TMPRSS2 | 1.0 | 3UTR |
| hsa-miR-6731-5p | NM_001135099 | TMPRSS2 | 1.0 | 3UTR |
| hsa-miR-6734-5p | NM_001135099 | TMPRSS2 | 1.0 | 3UTR |
| hsa-miR-6735-5p | NM_001135099 | TMPRSS2 | 1.0 | 3UTR |
| hsa-miR-6735-3p | NM_001135099 | TMPRSS2 | 1.0 | 3UTR |
| hsa-miR-6736-3p | NM_001135099 | TMPRSS2 | 1.0 | 3UTR |
| hsa-miR-6737-5p | NM_001135099 | TMPRSS2 | 1.0 | 3UTR |
| hsa-miR-6737-3p | NM_001135099 | TMPRSS2 | 1.0 | 3UTR |
| hsa-miR-6740-5p | NM_001135099 | TMPRSS2 | 1.0 | 3UTR |
| hsa-miR-6740-3p | NM_001135099 | TMPRSS2 | 1.0 | 3UTR |
| hsa-miR-6748-5p | NM_001135099 | TMPRSS2 | 1.0 | 3UTR |
| hsa-miR-6751-3p | NM_001135099 | TMPRSS2 | 1.0 | 3UTR |
| hsa-miR-6754-3p | NM_001135099 | TMPRSS2 | 1.0 | 3UTR |
| hsa-miR-6755-3p | NM_001135099 | TMPRSS2 | 1.0 | 3UTR |
| hsa-miR-6757-3p | NM_001135099 | TMPRSS2 | 1.0 | 3UTR |
| hsa-miR-6759-3p | NM_001135099 | TMPRSS2 | 1.0 | 3UTR |
| hsa-miR-6761-3p | NM_001135099 | TMPRSS2 | 1.0 | 3UTR |
| hsa-miR-6767-5p | NM_001135099 | TMPRSS2 | 1.0 | 3UTR |
| hsa-miR-6769a-5p | NM_001135099 | TMPRSS2 | 1.0 | 3UTR |
| hsa-miR-6771-5p | NM_001135099 | TMPRSS2 | 1.0 | 3UTR |
| hsa-miR-6772-5p | NM_001135099 | TMPRSS2 | 1.0 | 3UTR |
| hsa-miR-6772-3p | NM_001135099 | TMPRSS2 | 1.0 | 3UTR |
| hsa-miR-6773-3p | NM_001135099 | TMPRSS2 | 1.0 | 3UTR |
| hsa-miR-6774-5p | NM_001135099 | TMPRSS2 | 1.0 | 3UTR |
| hsa-miR-6774-5p | NM_001135099 | TMPRSS2 | 1.0 | 3UTR |
| hsa-miR-6775-3p | NM_001135099 | TMPRSS2 | 1.0 | 3UTR |

| | | | | |
|---|---|---|---|---|
| hsa-miR-6777-3p | NM_001135099 | TMPRSS2 | 1.0 | 3UTR |
| hsa-miR-6778-5p | NM_001135099 | TMPRSS2 | 1.0 | 3UTR |
| hsa-miR-6780a-5p | NM_001135099 | TMPRSS2 | 1.0 | 3UTR |
| hsa-miR-6780a-5p | NM_001135099 | TMPRSS2 | 1.0 | 3UTR |
| hsa-miR-6787-5p | NM_001135099 | TMPRSS2 | 1.0 | 3UTR |
| hsa-miR-6793-5p | NM_001135099 | TMPRSS2 | 1.0 | 3UTR |
| hsa-miR-6794-5p | NM_001135099 | TMPRSS2 | 1.0 | 3UTR |
| hsa-miR-6795-3p | NM_001135099 | TMPRSS2 | 1.0 | 3UTR |
| hsa-miR-6799-3p | NM_001135099 | TMPRSS2 | 1.0 | 3UTR |
| hsa-miR-6800-3p | NM_001135099 | TMPRSS2 | 1.0 | 3UTR |
| hsa-miR-6802-3p | NM_001135099 | TMPRSS2 | 1.0 | 3UTR |
| hsa-miR-6804-3p | NM_001135099 | TMPRSS2 | 1.0 | 3UTR |
| hsa-miR-6805-5p | NM_001135099 | TMPRSS2 | 1.0 | 3UTR |
| hsa-miR-6810-5p | NM_001135099 | TMPRSS2 | 1.0 | 3UTR |
| hsa-miR-6813-5p | NM_001135099 | TMPRSS2 | 1.0 | 3UTR |
| hsa-miR-6813-3p | NM_001135099 | TMPRSS2 | 1.0 | 3UTR |
| hsa-miR-6815-3p | NM_001135099 | TMPRSS2 | 1.0 | 3UTR |
| hsa-miR-6816-3p | NM_001135099 | TMPRSS2 | 1.0 | 3UTR |
| hsa-miR-6818-5p | NM_001135099 | TMPRSS2 | 1.0 | 3UTR |
| hsa-miR-6819-3p | NM_001135099 | TMPRSS2 | 1.0 | 3UTR |
| hsa-miR-6823-5p | NM_001135099 | TMPRSS2 | 1.0 | 3UTR |
| hsa-miR-6823-3p | NM_001135099 | TMPRSS2 | 1.0 | 3UTR |
| hsa-miR-6824-5p | NM_001135099 | TMPRSS2 | 1.0 | 3UTR |
| hsa-miR-6824-3p | NM_001135099 | TMPRSS2 | 1.0 | 3UTR |
| hsa-miR-6825-3p | NM_001135099 | TMPRSS2 | 1.0 | 3UTR |
| hsa-miR-6828-5p | NM_001135099 | TMPRSS2 | 1.0 | 3UTR |
| hsa-miR-6829-5p | NM_001135099 | TMPRSS2 | 1.0 | 3UTR |
| hsa-miR-6833-3p | NM_001135099 | TMPRSS2 | 1.0 | 3UTR |
| hsa-miR-6834-5p | NM_001135099 | TMPRSS2 | 1.0 | 3UTR |
| hsa-miR-6835-5p | NM_001135099 | TMPRSS2 | 1.0 | 3UTR |
| hsa-miR-6780b-3p | NM_001135099 | TMPRSS2 | 1.0 | 3UTR |
| hsa-miR-6837-5p | NM_001135099 | TMPRSS2 | 1.0 | 3UTR |
| hsa-miR-6841-5p | NM_001135099 | TMPRSS2 | 1.0 | 3UTR |
| hsa-miR-6842-5p | NM_001135099 | TMPRSS2 | 1.0 | 3UTR |
| hsa-miR-6842-3p | NM_001135099 | TMPRSS2 | 1.0 | 3UTR |
| hsa-miR-6843-3p | NM_001135099 | TMPRSS2 | 1.0 | 3UTR |
| hsa-miR-6845-3p | NM_001135099 | TMPRSS2 | 1.0 | 3UTR |
| hsa-miR-6848-5p | NM_001135099 | TMPRSS2 | 1.0 | 3UTR |
| hsa-miR-6852-5p | NM_001135099 | TMPRSS2 | 1.0 | 3UTR |
| hsa-miR-6852-5p | NM_001135099 | TMPRSS2 | 1.0 | 3UTR |
| hsa-miR-6854-3p | NM_001135099 | TMPRSS2 | 1.0 | 3UTR |
| hsa-miR-6859-5p | NM_001135099 | TMPRSS2 | 1.0 | 3UTR |
| hsa-miR-6859-3p | NM_001135099 | TMPRSS2 | 1.0 | 3UTR |
| hsa-miR-6769b-5p | NM_001135099 | TMPRSS2 | 1.0 | 3UTR |
| hsa-miR-6769b-3p | NM_001135099 | TMPRSS2 | 1.0 | 3UTR |
| hsa-miR-6860 | NM_001135099 | TMPRSS2 | 1.0 | 3UTR |
| hsa-miR-6860 | NM_001135099 | TMPRSS2 | 1.0 | 3UTR |

| | | | | |
|---|---|---|---|---|
| hsa-miR-6861-5p | NM_001135099 | TMPRSS2 | 1.0 | 3UTR |
| hsa-miR-6864-5p | NM_001135099 | TMPRSS2 | 1.0 | 3UTR |
| hsa-miR-6865-5p | NM_001135099 | TMPRSS2 | 1.0 | 3UTR |
| hsa-miR-6871-5p | NM_001135099 | TMPRSS2 | 1.0 | 3UTR |
| hsa-miR-6872-5p | NM_001135099 | TMPRSS2 | 1.0 | 3UTR |
| hsa-miR-6875-3p | NM_001135099 | TMPRSS2 | 1.0 | 3UTR |
| hsa-miR-6876-3p | NM_001135099 | TMPRSS2 | 1.0 | 3UTR |
| hsa-miR-6876-3p | NM_001135099 | TMPRSS2 | 1.0 | 3UTR |
| hsa-miR-6877-3p | NM_001135099 | TMPRSS2 | 1.0 | 3UTR |
| hsa-miR-6878-3p | NM_001135099 | TMPRSS2 | 1.0 | 3UTR |
| hsa-miR-6879-5p | NM_001135099 | TMPRSS2 | 1.0 | 3UTR |
| hsa-miR-6880-3p | NM_001135099 | TMPRSS2 | 1.0 | 3UTR |
| hsa-miR-6883-5p | NM_001135099 | TMPRSS2 | 1.0 | 3UTR |
| hsa-miR-6888-5p | NM_001135099 | TMPRSS2 | 1.0 | 3UTR |
| hsa-miR-6889-3p | NM_001135099 | TMPRSS2 | 1.0 | 3UTR |
| hsa-miR-6891-5p | NM_001135099 | TMPRSS2 | 1.0 | 3UTR |
| hsa-miR-6894-3p | NM_001135099 | TMPRSS2 | 1.0 | 3UTR |
| hsa-miR-7107-5p | NM_001135099 | TMPRSS2 | 1.0 | 3UTR |
| hsa-miR-7108-5p | NM_001135099 | TMPRSS2 | 1.0 | 3UTR |
| hsa-miR-7108-3p | NM_001135099 | TMPRSS2 | 1.0 | 3UTR |
| hsa-miR-7110-3p | NM_001135099 | TMPRSS2 | 1.0 | 3UTR |
| hsa-miR-7111-3p | NM_001135099 | TMPRSS2 | 1.0 | 3UTR |
| hsa-miR-7112-5p | NM_001135099 | TMPRSS2 | 1.0 | 3UTR |
| hsa-miR-7112-3p | NM_001135099 | TMPRSS2 | 1.0 | 3UTR |
| hsa-miR-7112-3p | NM_001135099 | TMPRSS2 | 1.0 | 3UTR |
| hsa-miR-7113-5p | NM_001135099 | TMPRSS2 | 1.0 | 3UTR |
| hsa-miR-7151-3p | NM_001135099 | TMPRSS2 | 1.0 | 3UTR |
| hsa-miR-7152-5p | NM_001135099 | TMPRSS2 | 1.0 | 3UTR |
| hsa-miR-7158-5p | NM_001135099 | TMPRSS2 | 1.0 | 3UTR |
| hsa-miR-7702 | NM_001135099 | TMPRSS2 | 1.0 | 3UTR |
| hsa-miR-7703 | NM_001135099 | TMPRSS2 | 1.0 | 3UTR |
| hsa-miR-7843-5p | NM_001135099 | TMPRSS2 | 1.0 | 3UTR |
| hsa-miR-7845-5p | NM_001135099 | TMPRSS2 | 1.0 | 3UTR |
| hsa-miR-7846-3p | NM_001135099 | TMPRSS2 | 1.0 | 3UTR |
| hsa-miR-7846-3p | NM_001135099 | TMPRSS2 | 1.0 | 3UTR |
| hsa-miR-7847-3p | NM_001135099 | TMPRSS2 | 1.0 | 3UTR |
| hsa-miR-7851-3p | NM_001135099 | TMPRSS2 | 1.0 | 3UTR |
| hsa-miR-8052 | NM_001135099 | TMPRSS2 | 1.0 | 3UTR |
| hsa-miR-8063 | NM_001135099 | TMPRSS2 | 1.0 | 3UTR |
| hsa-miR-8072 | NM_001135099 | TMPRSS2 | 1.0 | 3UTR |
| hsa-miR-8085 | NM_001135099 | TMPRSS2 | 1.0 | 3UTR |
| hsa-miR-8088 | NM_001135099 | TMPRSS2 | 1.0 | 3UTR |
| hsa-miR-9985 | NM_001135099 | TMPRSS2 | 1.0 | 3UTR |
| hsa-miR-10395-5p | NM_001135099 | TMPRSS2 | 1.0 | 3UTR |
| hsa-miR-10398-5p | NM_001135099 | TMPRSS2 | 1.0 | 3UTR |
| hsa-miR-10526-3p | NM_001135099 | TMPRSS2 | 1.0 | 3UTR |
| hsa-miR-11181-3p | NM_001135099 | TMPRSS2 | 1.0 | 3UTR |

| | | | | |
|---|---|---|---|---|
| hsa-miR-3085-5p | NM_001135099 | TMPRSS2 | 1.0 | 3UTR |
| hsa-miR-6529-5p | NM_001135099 | TMPRSS2 | 1.0 | 3UTR |
| hsa-miR-9851-5p | NM_001135099 | TMPRSS2 | 1.0 | 3UTR |
| hsa-miR-9851-3p | NM_001135099 | TMPRSS2 | 1.0 | 3UTR |
| hsa-miR-12116 | NM_001135099 | TMPRSS2 | 1.0 | 3UTR |
| hsa-miR-12119 | NM_001135099 | TMPRSS2 | 1.0 | 3UTR |

**Supplementary Table 2**

**miRNA-target experimental validated interactions (miRNet)**

| ID | Target | Experiment |
|---|---|---|
| hsa-mir-193b-3p | AAMP | CLASH |
| hsa-mir-193b-3p | AARS | CLASH |
| hsa-mir-193b-3p | ACACA | CLASH |
| hsa-mir-193b-3p | ACTG1 | CLASH |
| hsa-mir-193b-3p | ACTN4 | CLASH//HITS-CLIP |
| hsa-mir-193b-3p | ACTN1 | Proteomics |
| hsa-mir-193b-3p | ADARB1 | CLASH |
| hsa-mir-193b-3p | ADCY9 | Microarray |
| hsa-mir-193b-3p | PARP1 | CLASH |
| hsa-mir-193b-3p | AKT1 | CLASH |
| hsa-mir-193b-3p | ALDH3A2 | CLASH |
| hsa-mir-193b-3p | APEH | CLASH |
| hsa-mir-193b-3p | ARCN1 | CLASH |
| hsa-mir-193b-3p | ATP1A1 | Proteomics |
| hsa-mir-193b-3p | ALDH7A1 | CLASH |
| hsa-mir-193b-3p | ATP5B | Proteomics |
| hsa-mir-193b-3p | BARD1 | Microarray |
| hsa-mir-193b-3p | BCKDHA | CLASH |
| hsa-mir-193b-3p | CCND1 | CLASH//Immunohistochemistry//Luciferase reporter assay//Microarray//qRT-PCR//Reporter assay//Western blot |
| hsa-mir-193b-3p | BLM | Microarray |
| hsa-mir-193b-3p | BRCA1 | Microarray |
| hsa-mir-193b-3p | BUB1 | PAR-CLIP |
| hsa-mir-193b-3p | BUB1B | CLASH//Microarray |
| hsa-mir-193b-3p | C1QBP | HITS-CLIP |
| hsa-mir-193b-3p | C5 | Microarray |
| hsa-mir-193b-3p | CAPNS1 | CLASH |
| hsa-mir-193b-3p | CASP9 | Microarray |
| hsa-mir-193b-3p | CBS | CLASH |
| hsa-mir-193b-3p | KYAT1 | CLASH |
| hsa-mir-193b-3p | CCNA2 | Microarray |
| hsa-mir-193b-3p | CDK1 | Microarray |
| hsa-mir-193b-3p | CDC6 | Microarray |
| hsa-mir-193b-3p | CDC20 | Microarray |
| hsa-mir-193b-3p | CDC25A | Microarray |
| hsa-mir-193b-3p | CDH1 | Proteomics |
| hsa-mir-193b-3p | CDK4 | CLASH |
| hsa-mir-193b-3p | CDK6 | Microarray |
| hsa-mir-193b-3p | CDK8 | CLASH |
| hsa-mir-193b-3p | CDK9 | CLASH |
| hsa-mir-193b-3p | CFL1 | CLASH |
| hsa-mir-193b-3p | RCC1 | CLASH//Microarray |
| hsa-mir-193b-3p | CHD4 | CLASH//Proteomics |

| | | |
|---|---|---|
| hsa-mir-193b-3p | CHEK1 | Microarray |
| hsa-mir-193b-3p | COL4A1 | CLASH |
| hsa-mir-193b-3p | COPA | CLASH |
| hsa-mir-193b-3p | COX7C | CLASH |
| hsa-mir-193b-3p | CS | CLASH |
| hsa-mir-193b-3p | SLC25A10 | CLASH |
| hsa-mir-193b-3p | DDB1 | Proteomics |
| hsa-mir-193b-3p | AKR1C2 | Luciferase reporter assay//Proteomics//Reporter assay |
| hsa-mir-193b-3p | DLX1 | Microarray |
| hsa-mir-193b-3p | DNMT1 | Microarray |
| hsa-mir-193b-3p | DNMT3A | Microarray |
| hsa-mir-193b-3p | DPH1 | CLASH |
| hsa-mir-193b-3p | DRG2 | CLASH |
| hsa-mir-193b-3p | DUT | Proteomics |
| hsa-mir-193b-3p | E2F1 | Microarray |
| hsa-mir-193b-3p | E2F2 | Microarray |
| hsa-mir-193b-3p | E2F6 | Microarray |
| hsa-mir-193b-3p | ECH1 | Proteomics |
| hsa-mir-193b-3p | ECT2 | Microarray |
| hsa-mir-193b-3p | EEF2 | CLASH |
| hsa-mir-193b-3p | MEGF8 | CLASH |
| hsa-mir-193b-3p | EPHA2 | Microarray |
| hsa-mir-193b-3p | EIF4B | CLASH//Proteomics |
| hsa-mir-193b-3p | ELK3 | Microarray |
| hsa-mir-193b-3p | MARK2 | CLASH |
| hsa-mir-193b-3p | EP300 | CLASH |
| hsa-mir-193b-3p | EPHX1 | Proteomics |
| hsa-mir-193b-3p | EPRS | CLASH |
| hsa-mir-193b-3p | ESD | CLASH |
| hsa-mir-193b-3p | ESR1 | Luciferase reporter assay//Microarray |
| hsa-mir-193b-3p | ETS1 | Luciferase reporter assay//qRT-PCR//Western blot |
| hsa-mir-193b-3p | EZH2 | Microarray |
| hsa-mir-193b-3p | FANCA | Microarray |
| hsa-mir-193b-3p | FANCD2 | Microarray |
| hsa-mir-193b-3p | FANCE | Microarray |
| hsa-mir-193b-3p | FANCG | Microarray |
| hsa-mir-193b-3p | FASN | CLASH |
| hsa-mir-193b-3p | FEN1 | Microarray//Proteomics |
| hsa-mir-193b-3p | FGF11 | CLASH |
| hsa-mir-193b-3p | FOXC1 | CLASH |
| hsa-mir-193b-3p | FLNA | CLASH |
| hsa-mir-193b-3p | KDSR | HITS-CLIP |
| hsa-mir-193b-3p | SLC37A4 | CLASH |
| hsa-mir-193b-3p | XRCC6 | CLASH |
| hsa-mir-193b-3p | GDI2 | CLASH |
| hsa-mir-193b-3p | GJB2 | Microarray |
| hsa-mir-193b-3p | GLO1 | Microarray |

| | | |
|---|---|---|
| hsa-mir-193b-3p | GLUD1 | CLASH |
| hsa-mir-193b-3p | GPI | CLASH |
| hsa-mir-193b-3p | RAPGEF1 | CLASH |
| hsa-mir-193b-3p | GSS | CLASH |
| hsa-mir-193b-3p | MSH6 | Microarray |
| hsa-mir-193b-3p | HIST1H1D | Microarray |
| hsa-mir-193b-3p | HIST1H1E | CLASH |
| hsa-mir-193b-3p | HIST1H1B | CLASH |
| hsa-mir-193b-3p | HADHB | Proteomics |
| hsa-mir-193b-3p | HCFC1 | CLASH |
| hsa-mir-193b-3p | HDGF | CLASH |
| hsa-mir-193b-3p | HELLS | Microarray |
| hsa-mir-193b-3p | HMGB1 | Microarray |
| hsa-mir-193b-3p | HMGCR | Microarray |
| hsa-mir-193b-3p | HNRNPAB | CLASH |
| hsa-mir-193b-3p | HNRNPH1 | CLASH |
| hsa-mir-193b-3p | HNRNPL | Proteomics |
| hsa-mir-193b-3p | HPRT1 | Microarray |
| hsa-mir-193b-3p | DNAJA1 | CLASH |
| hsa-mir-193b-3p | HSPA1B | CLASH |
| hsa-mir-193b-3p | HSPA1L | CLASH |
| hsa-mir-193b-3p | HSP90AB1 | Proteomics |
| hsa-mir-193b-3p | NDST1 | CLASH |
| hsa-mir-193b-3p | IDI1 | Proteomics |
| hsa-mir-193b-3p | IGFBP5 | PAR-CLIP |
| hsa-mir-193b-3p | INSIG1 | CLASH |
| hsa-mir-193b-3p | IRAK1 | CLASH |
| hsa-mir-193b-3p | IRF1 | HITS-CLIP |
| hsa-mir-193b-3p | ITPKA | Microarray |
| hsa-mir-193b-3p | KIT | Luciferase reporter assay |
| hsa-mir-193b-3p | KIF11 | Microarray |
| hsa-mir-193b-3p | KIF22 | Microarray |
| hsa-mir-193b-3p | KPNA2 | Proteomics |
| hsa-mir-193b-3p | KRAS | In situ hybridization//Luciferase reporter assay//Microarray//PAR-CLIP//RTPCR//Western blot |
| hsa-mir-193b-3p | KRT19 | Proteomics |
| hsa-mir-193b-3p | LAMB1 | CLASH |
| hsa-mir-193b-3p | LAMC1 | PAR-CLIP |
| hsa-mir-193b-3p | STMN1 | Microarray |
| hsa-mir-193b-3p | LASP1 | Proteomics |
| hsa-mir-193b-3p | LETM1 | Proteomics |
| hsa-mir-193b-3p | LMNB1 | CLASH |
| hsa-mir-193b-3p | LOXL1 | Microarray |
| hsa-mir-193b-3p | SMAD3 | CLASH//Luciferase reporter assay//qRT-PCR//Western blot |
| hsa-mir-193b-3p | MAGEB2 | CLASH |
| hsa-mir-193b-3p | MARS | CLASH |
| hsa-mir-193b-3p | MAT2A | Proteomics |

| | | |
|---|---|---|
| hsa-mir-193b-3p | MAX | In situ hybridization//Luciferase reporter assay//Microarray//RTPCR//Western blot |
| hsa-mir-193b-3p | MAZ | CLASH |
| hsa-mir-193b-3p | MCL1 | Luciferase reporter assay//Microarray//qRT-PCR//Western blot |
| hsa-mir-193b-3p | MCM3 | Microarray |
| hsa-mir-193b-3p | MCM4 | Microarray//Proteomics |
| hsa-mir-193b-3p | MCM5 | Microarray//Proteomics |
| hsa-mir-193b-3p | MCM6 | Microarray//Proteomics |
| hsa-mir-193b-3p | MCM7 | Microarray//Proteomics |
| hsa-mir-193b-3p | DNAJB9 | PAR-CLIP |
| hsa-mir-193b-3p | MDH2 | CLASH//PAR-CLIP |
| hsa-mir-193b-3p | MAP3K1 | HITS-CLIP |
| hsa-mir-193b-3p | MAP3K3 | CLASH//HITS-CLIP |
| hsa-mir-193b-3p | MPST | Microarray |
| hsa-mir-193b-3p | COX1 | CLASH |
| hsa-mir-193b-3p | MTHFD1 | Proteomics |
| hsa-mir-193b-3p | MTR | CLASH |
| hsa-mir-193b-3p | MYB | Luciferase reporter assay//Microarray |
| hsa-mir-193b-3p | MYBL1 | Microarray |
| hsa-mir-193b-3p | MYH9 | CLASH |
| hsa-mir-193b-3p | MYLK | Microarray |
| hsa-mir-193b-3p | MYO1D | CLASH |
| hsa-mir-193b-3p | MYO5A | CLASH |
| hsa-mir-193b-3p | NAGA | Microarray |
| hsa-mir-193b-3p | HNRNPM | CLASH |
| hsa-mir-193b-3p | NCL | CLASH |
| hsa-mir-193b-3p | NDUFS6 | CLASH |
| hsa-mir-193b-3p | NF1 | CLASH//HITS-CLIP//In situ hybridization//Luciferase reporter assay//qRT-PCR//Western blot |
| hsa-mir-193b-3p | NF2 | Microarray |
| hsa-mir-193b-3p | NFIA | CLASH |
| hsa-mir-193b-3p | NFKB2 | Microarray |
| hsa-mir-193b-3p | NME4 | CLASH |
| hsa-mir-193b-3p | NQO2 | HITS-CLIP |
| hsa-mir-193b-3p | NMT1 | CLASH |
| hsa-mir-193b-3p | NONO | CLASH |
| hsa-mir-193b-3p | NPM1 | CLASH |
| hsa-mir-193b-3p | NPPC | Microarray |
| hsa-mir-193b-3p | NSF | CLASH |
| hsa-mir-193b-3p | NUCB1 | Proteomics |
| hsa-mir-193b-3p | NUCB2 | Proteomics |
| hsa-mir-193b-3p | ODC1 | CLASH |
| hsa-mir-193b-3p | OPHN1 | PAR-CLIP |
| hsa-mir-193b-3p | PAK2 | CLASH |
| hsa-mir-193b-3p | PCNA | Microarray |
| hsa-mir-193b-3p | PCNT | Microarray |
| hsa-mir-193b-3p | CDK17 | CLASH |

| | | |
|---|---|---|
| hsa-mir-193b-3p | PFDN2 | Proteomics |
| hsa-mir-193b-3p | PFKP | CLASH |
| hsa-mir-193b-3p | PFN1 | CLASH |
| hsa-mir-193b-3p | PGAM1 | CLASH |
| hsa-mir-193b-3p | PGM3 | HITS-CLIP |
| hsa-mir-193b-3p | PIP4K2A | CLASH |
| hsa-mir-193b-3p | PLAU | HITS-CLIP//Luciferase reporter assay//Microarray//qRT-PCR//Western blot |
| hsa-mir-193b-3p | PLS3 | CLASH |
| hsa-mir-193b-3p | PMAIP1 | CLASH//PAR-CLIP |
| hsa-mir-193b-3p | EXOSC10 | CLASH |
| hsa-mir-193b-3p | POLA1 | Microarray |
| hsa-mir-193b-3p | POLD1 | Microarray |
| hsa-mir-193b-3p | POLE | Microarray |
| hsa-mir-193b-3p | POLE2 | Microarray |
| hsa-mir-193b-3p | POLR2C | CLASH |
| hsa-mir-193b-3p | PPP2R5C | HITS-CLIP |
| hsa-mir-193b-3p | PRIM1 | Microarray |
| hsa-mir-193b-3p | PRKCA | CLASH |
| hsa-mir-193b-3p | MAPK8 | Microarray//PAR-CLIP |
| hsa-mir-193b-3p | PRNP | Microarray |
| hsa-mir-193b-3p | PSMA7 | CLASH |
| hsa-mir-193b-3p | PSMC5 | CLASH |
| hsa-mir-193b-3p | PTEN | CLASH |
| hsa-mir-193b-3p | PTK7 | CLASH |
| hsa-mir-193b-3p | TWF1 | CLASH |
| hsa-mir-193b-3p | PTMS | Proteomics |
| hsa-mir-193b-3p | PTPN9 | HITS-CLIP |
| hsa-mir-193b-3p | PTPN11 | CLASH |
| hsa-mir-193b-3p | PTPRG | CLASH |
| hsa-mir-193b-3p | RABGGTB | CLASH |
| hsa-mir-193b-3p | RAB5C | Proteomics |
| hsa-mir-193b-3p | RAC2 | Microarray |
| hsa-mir-193b-3p | RAD51 | ChIP-seq//Luciferase reporter assay//Microarray//qRT-PCR//Western blot |
| hsa-mir-193b-3p | RBBP5 | Microarray |
| hsa-mir-193b-3p | RBL1 | PAR-CLIP |
| hsa-mir-193b-3p | RFC4 | Microarray |
| hsa-mir-193b-3p | RFC5 | Microarray |
| hsa-mir-193b-3p | RPL8 | CLASH |
| hsa-mir-193b-3p | RPL9 | CLASH |
| hsa-mir-193b-3p | RPL12 | CLASH |
| hsa-mir-193b-3p | RPL22 | CLASH |
| hsa-mir-193b-3p | RPL23A | CLASH |
| hsa-mir-193b-3p | RPL26 | CLASH |
| hsa-mir-193b-3p | RPL27A | CLASH |
| hsa-mir-193b-3p | RPS3 | CLASH |

| | | |
|---|---|---|
| hsa-mir-193b-3p | RPS6KA1 | CLASH |
| hsa-mir-193b-3p | RPS10 | CLASH |
| hsa-mir-193b-3p | RPS18 | CLASH |
| hsa-mir-193b-3p | RPS21 | PAR-CLIP |
| hsa-mir-193b-3p | RRM2 | CLASH//Microarray |
| hsa-mir-193b-3p | RTKN | PAR-CLIP |
| hsa-mir-193b-3p | SCO1 | CLASH |
| hsa-mir-193b-3p | SFPQ | CLASH |
| hsa-mir-193b-3p | SRSF1 | CLASH |
| hsa-mir-193b-3p | SH3GL1 | CLASH |
| hsa-mir-193b-3p | SHMT1 | Microarray |
| hsa-mir-193b-3p | SHMT2 | CLASH//Luciferase reporter assay//Microarray//Proteomics//Reporter assay |
| hsa-mir-193b-3p | SIX1 | Microarray |
| hsa-mir-193b-3p | SLC1A5 | Proteomics |
| hsa-mir-193b-3p | SLC3A2 | CLASH |
| hsa-mir-193b-3p | SNRNP70 | CLASH |
| hsa-mir-193b-3p | SNRPB | CLASH |
| hsa-mir-193b-3p | SNRPD3 | CLASH |
| hsa-mir-193b-3p | SOAT1 | Microarray |
| hsa-mir-193b-3p | CAPN15 | CLASH |
| hsa-mir-193b-3p | SPTBN1 | CLASH |
| hsa-mir-193b-3p | SPTBN2 | CLASH |
| hsa-mir-193b-3p | SSRP1 | CLASH |
| hsa-mir-193b-3p | SURF4 | Proteomics |
| hsa-mir-193b-3p | TACC1 | Microarray |
| hsa-mir-193b-3p | ELOA | CLASH |
| hsa-mir-193b-3p | TCF7L2 | PAR-CLIP |
| hsa-mir-193b-3p | TERT | Microarray |
| hsa-mir-193b-3p | TGFBR3 | Microarray |
| hsa-mir-193b-3p | TK1 | Microarray |
| hsa-mir-193b-3p | TLE4 | CLASH |
| hsa-mir-193b-3p | TLN1 | HITS-CLIP |
| hsa-mir-193b-3p | TM7SF2 | Proteomics |
| hsa-mir-193b-3p | TRAPPC10 | CLASH |
| hsa-mir-193b-3p | TNFRSF1B | HITS-CLIP |
| hsa-mir-193b-3p | TOP2A | Microarray |
| hsa-mir-193b-3p | TPI1 | CLASH |
| hsa-mir-193b-3p | TSC1 | CLASH |
| hsa-mir-193b-3p | PHLDA2 | Microarray//PAR-CLIP |
| hsa-mir-193b-3p | TYMS | Microarray |
| hsa-mir-193b-3p | UCHL1 | CLASH |
| hsa-mir-193b-3p | UGP2 | Proteomics |
| hsa-mir-193b-3p | VASP | Proteomics |
| hsa-mir-193b-3p | VAV2 | CLASH |
| hsa-mir-193b-3p | NSD2 | CLASH//Microarray |
| hsa-mir-193b-3p | XPO1 | Proteomics |

| | | |
|---|---|---|
| hsa-mir-193b-3p | XRCC1 | CLASH |
| hsa-mir-193b-3p | YY1 | PAR-CLIP |
| hsa-mir-193b-3p | YWHAZ | CLASH//HITS-CLIP//Luciferase reporter assay//Proteomics//Reporter assay |
| hsa-mir-193b-3p | ZNF3 | CLASH |
| hsa-mir-193b-3p | SLC30A1 | CLASH |
| hsa-mir-193b-3p | ZYX | CLASH |
| hsa-mir-193b-3p | PRRC2A | CLASH |
| hsa-mir-193b-3p | BAG6 | CLASH |
| hsa-mir-193b-3p | AIMP2 | Microarray |
| hsa-mir-193b-3p | KAT6A | CLASH |
| hsa-mir-193b-3p | NUP214 | CLASH |
| hsa-mir-193b-3p | PDHX | CLASH |
| hsa-mir-193b-3p | AAAS | CLASH |
| hsa-mir-193b-3p | SLC7A5 | Microarray |
| hsa-mir-193b-3p | SYMPK | CLASH |
| hsa-mir-193b-3p | CHAF1B | Microarray |
| hsa-mir-193b-3p | ARID1A | CLASH |
| hsa-mir-193b-3p | TRRAP | CLASH |
| hsa-mir-193b-3p | HIST1H2AJ | CLASH |
| hsa-mir-193b-3p | HIST2H2AA3 | CLASH |
| hsa-mir-193b-3p | HIST1H3D | CLASH |
| hsa-mir-193b-3p | HIST1H3H | CLASH |
| hsa-mir-193b-3p | HIST1H3B | CLASH |
| hsa-mir-193b-3p | STX7 | CLASH |
| hsa-mir-193b-3p | DYRK2 | HITS-CLIP |
| hsa-mir-193b-3p | CUL4A | CLASH |
| hsa-mir-193b-3p | TEAD2 | CLASH |
| hsa-mir-193b-3p | IRS4 | CLASH |
| hsa-mir-193b-3p | YBX3 | CLASH |
| hsa-mir-193b-3p | YARS | CLASH |
| hsa-mir-193b-3p | PDXK | Proteomics |
| hsa-mir-193b-3p | RUVBL1 | CLASH |
| hsa-mir-193b-3p | SSNA1 | CLASH |
| hsa-mir-193b-3p | EIF3I | CLASH//Proteomics |
| hsa-mir-193b-3p | VAMP8 | Microarray//Proteomics |
| hsa-mir-193b-3p | STX16 | CLASH |
| hsa-mir-193b-3p | MBTPS1 | CLASH |
| hsa-mir-193b-3p | GBF1 | Proteomics |
| hsa-mir-193b-3p | PEX11B | Microarray |
| hsa-mir-193b-3p | DLEU2 | Microarray |
| hsa-mir-193b-3p | ALDH1A2 | CLASH |
| hsa-mir-193b-3p | HERC2 | CLASH |
| hsa-mir-193b-3p | HERC1 | CLASH |
| hsa-mir-193b-3p | BTRC | Microarray |
| hsa-mir-193b-3p | BAZ1B | CLASH |
| hsa-mir-193b-3p | SPAG9 | CLASH |

| | | |
|---|---|---|
| hsa-mir-193b-3p | DOK2 | CLASH |
| hsa-mir-193b-3p | SLC7A6 | CLASH |
| hsa-mir-193b-3p | PKMYT1 | Microarray |
| hsa-mir-193b-3p | DNAJA3 | CLASH |
| hsa-mir-193b-3p | MTMR4 | CLASH |
| hsa-mir-193b-3p | SMC3 | CLASH |
| hsa-mir-193b-3p | FAM50A | CLASH |
| hsa-mir-193b-3p | CTDP1 | CLASH |
| hsa-mir-193b-3p | EXO1 | Microarray |
| hsa-mir-193b-3p | EBAG9 | PAR-CLIP |
| hsa-mir-193b-3p | DDX21 | Proteomics |
| hsa-mir-193b-3p | MAPKAPK2 | CLASH |
| hsa-mir-193b-3p | CYTH1 | Microarray |
| hsa-mir-193b-3p | TRIP13 | Microarray |
| hsa-mir-193b-3p | TRIP12 | CLASH |
| hsa-mir-193b-3p | EFTUD2 | CLASH |
| hsa-mir-193b-3p | CIAO1 | CLASH//HITS-CLIP |
| hsa-mir-193b-3p | RECQL4 | Microarray |
| hsa-mir-193b-3p | MED21 | PAR-CLIP |
| hsa-mir-193b-3p | TJP2 | Microarray |
| hsa-mir-193b-3p | SH3BP5 | CLASH |
| hsa-mir-193b-3p | TECR | Proteomics |
| hsa-mir-193b-3p | GOSR1 | CLASH |
| hsa-mir-193b-3p | APOBEC3B | Microarray |
| hsa-mir-193b-3p | GCC2 | CLASH |
| hsa-mir-193b-3p | ESPL1 | Microarray |
| hsa-mir-193b-3p | KNTC1 | Microarray |
| hsa-mir-193b-3p | CCP110 | Microarray |
| hsa-mir-193b-3p | JADE3 | Microarray |
| hsa-mir-193b-3p | PCLAF | Microarray |
| hsa-mir-193b-3p | TMEM94 | CLASH |
| hsa-mir-193b-3p | DAZAP2 | Microarray |
| hsa-mir-193b-3p | NUP58 | Microarray |
| hsa-mir-193b-3p | RB1CC1 | CLASH |
| hsa-mir-193b-3p | ARHGAP11A | Microarray |
| hsa-mir-193b-3p | MELK | Microarray |
| hsa-mir-193b-3p | AREL1 | Microarray |
| hsa-mir-193b-3p | DENND4B | CLASH |
| hsa-mir-193b-3p | FAM20B | Microarray |
| hsa-mir-193b-3p | NCAPD2 | Proteomics |
| hsa-mir-193b-3p | KBTBD11 | Microarray |
| hsa-mir-193b-3p | ZBTB5 | Microarray |
| hsa-mir-193b-3p | LPGAT1 | CLASH |
| hsa-mir-193b-3p | MFN2 | CLASH |
| hsa-mir-193b-3p | RBM8A | CLASH//Microarray |
| hsa-mir-193b-3p | FARSB | CLASH |
| hsa-mir-193b-3p | ABCF2 | CLASH |

| | | |
|---|---|---|
| hsa-mir-193b-3p | HUWE1 | CLASH |
| hsa-mir-193b-3p | HHLA1 | PAR-CLIP |
| hsa-mir-193b-3p | PPIF | HITS-CLIP |
| hsa-mir-193b-3p | CTDSP2 | Microarray |
| hsa-mir-193b-3p | ACTR1A | CLASH |
| hsa-mir-193b-3p | TRAP1 | CLASH |
| hsa-mir-193b-3p | G3BP1 | CLASH |
| hsa-mir-193b-3p | ABI2 | Microarray//PAR-CLIP |
| hsa-mir-193b-3p | PLXNC1 | Microarray |
| hsa-mir-193b-3p | TRIM28 | CLASH |
| hsa-mir-193b-3p | SLC25A15 | Microarray |
| hsa-mir-193b-3p | NUTF2 | CLASH |
| hsa-mir-193b-3p | EIF1 | CLASH |
| hsa-mir-193b-3p | GDF11 | PAR-CLIP |
| hsa-mir-193b-3p | PLIN3 | Proteomics |
| hsa-mir-193b-3p | COQ7 | HITS-CLIP |
| hsa-mir-193b-3p | DCAF7 | HITS-CLIP//Microarray//PAR-CLIP |
| hsa-mir-193b-3p | EFS | Microarray |
| hsa-mir-193b-3p | SF3A1 | CLASH |
| hsa-mir-193b-3p | TFG | CLASH |
| hsa-mir-193b-3p | TUBA1B | Proteomics |
| hsa-mir-193b-3p | TUBB3 | CLASH//Proteomics |
| hsa-mir-193b-3p | NDC80 | Microarray |
| hsa-mir-193b-3p | BASP1 | Proteomics |
| hsa-mir-193b-3p | LYPLA1 | HITS-CLIP |
| hsa-mir-193b-3p | MCRS1 | CLASH |
| hsa-mir-193b-3p | VAV3 | Microarray |
| hsa-mir-193b-3p | TACC3 | CLASH//Microarray |
| hsa-mir-193b-3p | EIF3M | CLASH |
| hsa-mir-193b-3p | SEC23A | Proteomics |
| hsa-mir-193b-3p | ENOX2 | Microarray |
| hsa-mir-193b-3p | NCOA2 | CLASH |
| hsa-mir-193b-3p | MYBBP1A | CLASH |
| hsa-mir-193b-3p | HYOU1 | CLASH//Microarray |
| hsa-mir-193b-3p | IPO8 | CLASH |
| hsa-mir-193b-3p | PITRM1 | CLASH |
| hsa-mir-193b-3p | ANP32B | CLASH |
| hsa-mir-193b-3p | AGPAT1 | Microarray |
| hsa-mir-193b-3p | CCT4 | CLASH |
| hsa-mir-193b-3p | CCT2 | CLASH |
| hsa-mir-193b-3p | PRPF8 | CLASH |
| hsa-mir-193b-3p | AHSA1 | CLASH |
| hsa-mir-193b-3p | USP39 | Microarray |
| hsa-mir-193b-3p | POLQ | Microarray |
| hsa-mir-193b-3p | TCFL5 | CLASH |
| hsa-mir-193b-3p | STAG2 | CLASH |
| hsa-mir-193b-3p | HSPH1 | CLASH |

| | | |
|---|---|---|
| hsa-mir-193b-3p | TRAFD1 | HITS-CLIP |
| hsa-mir-193b-3p | TXNL4A | CLASH |
| hsa-mir-193b-3p | PRDX3 | Proteomics |
| hsa-mir-193b-3p | AP3M2 | CLASH |
| hsa-mir-193b-3p | CBX1 | Microarray |
| hsa-mir-193b-3p | TMED2 | Proteomics |
| hsa-mir-193b-3p | EBNA1BP2 | CLASH |
| hsa-mir-193b-3p | YWHAQ | CLASH |
| hsa-mir-193b-3p | RAB32 | PAR-CLIP |
| hsa-mir-193b-3p | SF3B2 | CLASH |
| hsa-mir-193b-3p | ILVBL | CLASH |
| hsa-mir-193b-3p | RAB35 | CLASH |
| hsa-mir-193b-3p | SLC35D2 | Microarray |
| hsa-mir-193b-3p | UBE2C | Microarray |
| hsa-mir-193b-3p | CACFD1 | HITS-CLIP |
| hsa-mir-193b-3p | HNRNPUL1 | CLASH |
| hsa-mir-193b-3p | FAF1 | CLASH |
| hsa-mir-193b-3p | ZWINT | Microarray |
| hsa-mir-193b-3p | CDC37 | Proteomics |
| hsa-mir-193b-3p | PSIP1 | CLASH//Microarray |
| hsa-mir-193b-3p | BAZ2A | HITS-CLIP |
| hsa-mir-193b-3p | AKAP13 | CLASH |
| hsa-mir-193b-3p | PACSIN2 | CLASH |
| hsa-mir-193b-3p | SYNRG | HITS-CLIP |
| hsa-mir-193b-3p | PHB2 | CLASH |
| hsa-mir-193b-3p | OIP5 | Microarray |
| hsa-mir-193b-3p | CASC3 | CLASH |
| hsa-mir-193b-3p | SEC31A | Microarray |
| hsa-mir-193b-3p | CLSTN1 | Microarray |
| hsa-mir-193b-3p | ZNF365 | Microarray |
| hsa-mir-193b-3p | DHX30 | CLASH |
| hsa-mir-193b-3p | SEPHS2 | CLASH |
| hsa-mir-193b-3p | SEPHS1 | Microarray |
| hsa-mir-193b-3p | TPX2 | CLASH |
| hsa-mir-193b-3p | NT5C2 | CLASH |
| hsa-mir-193b-3p | SPEN | CLASH |
| hsa-mir-193b-3p | CNOT1 | CLASH |
| hsa-mir-193b-3p | SNRNP200 | CLASH |
| hsa-mir-193b-3p | XPO7 | CLASH |
| hsa-mir-193b-3p | PDXDC1 | Proteomics |
| hsa-mir-193b-3p | ENDOD1 | Microarray |
| hsa-mir-193b-3p | TBC1D9B | CLASH |
| hsa-mir-193b-3p | WAPL | CLASH |
| hsa-mir-193b-3p | EMC1 | CLASH |
| hsa-mir-193b-3p | CMTR1 | CLASH |
| hsa-mir-193b-3p | PEG10 | CLASH |
| hsa-mir-193b-3p | KIF1B | Microarray |

| | | |
|---|---|---|
| hsa-mir-193b-3p | ZBTB43 | CLASH |
| hsa-mir-193b-3p | PHF8 | CLASH |
| hsa-mir-193b-3p | CIC | CLASH |
| hsa-mir-193b-3p | TTLL12 | CLASH |
| hsa-mir-193b-3p | CYFIP1 | CLASH |
| hsa-mir-193b-3p | FAF2 | CLASH |
| hsa-mir-193b-3p | TBC1D1 | Microarray |
| hsa-mir-193b-3p | SYNE2 | Microarray |
| hsa-mir-193b-3p | DNAJC9 | Microarray |
| hsa-mir-193b-3p | ZC3H7B | CLASH |
| hsa-mir-193b-3p | BICD2 | CLASH |
| hsa-mir-193b-3p | ESYT1 | CLASH |
| hsa-mir-193b-3p | SUN1 | CLASH |
| hsa-mir-193b-3p | USP24 | CLASH |
| hsa-mir-193b-3p | SMG5 | CLASH |
| hsa-mir-193b-3p | PIP5K1C | CLASH |
| hsa-mir-193b-3p | NCAPH | Microarray |
| hsa-mir-193b-3p | ATP13A2 | CLASH |
| hsa-mir-193b-3p | COTL1 | Proteomics |
| hsa-mir-193b-3p | SF3B3 | CLASH |
| hsa-mir-193b-3p | SCRIB | CLASH |
| hsa-mir-193b-3p | DDAH1 | PAR-CLIP |
| hsa-mir-193b-3p | CDC42EP4 | Microarray |
| hsa-mir-193b-3p | ORC6 | Microarray |
| hsa-mir-193b-3p | ACOT9 | CLASH |
| hsa-mir-193b-3p | AMACR | CLASH |
| hsa-mir-193b-3p | RUSC1 | Microarray |
| hsa-mir-193b-3p | SH3BP4 | CLASH |
| hsa-mir-193b-3p | PITPNB | Microarray |
| hsa-mir-193b-3p | PRPF40B | Microarray |
| hsa-mir-193b-3p | GCA | CLASH |
| hsa-mir-193b-3p | YIPF3 | CLASH |
| hsa-mir-193b-3p | CCDC28A | Microarray |
| hsa-mir-193b-3p | CLIC4 | Proteomics |
| hsa-mir-193b-3p | WWTR1 | Microarray |
| hsa-mir-193b-3p | KANK2 | Microarray |
| hsa-mir-193b-3p | C2CD2 | CLASH |
| hsa-mir-193b-3p | TSKU | PAR-CLIP |
| hsa-mir-193b-3p | TENM4 | CLASH |
| hsa-mir-193b-3p | CHTOP | CLASH |
| hsa-mir-193b-3p | PRPF31 | CLASH |
| hsa-mir-193b-3p | PHF19 | Microarray |
| hsa-mir-193b-3p | RSL1D1 | CLASH |
| hsa-mir-193b-3p | GIMAP2 | Microarray |
| hsa-mir-193b-3p | FBXO5 | Microarray |
| hsa-mir-193b-3p | PLEK2 | Microarray |
| hsa-mir-193b-3p | ago-01 | CLASH |

| | | |
|---|---|---|
| hsa-mir-193b-3p | DAZAP1 | CLASH |
| hsa-mir-193b-3p | PABPC1 | CLASH |
| hsa-mir-193b-3p | TRMT2A | CLASH |
| hsa-mir-193b-3p | PELP1 | CLASH |
| hsa-mir-193b-3p | NSG1 | Microarray |
| hsa-mir-193b-3p | PPA2 | Proteomics |
| hsa-mir-193b-3p | TNFRSF21 | Microarray |
| hsa-mir-193b-3p | AHDC1 | CLASH |
| hsa-mir-193b-3p | RNF115 | CLASH |
| hsa-mir-193b-3p | PRPF19 | CLASH |
| hsa-mir-193b-3p | TMEM97 | CLASH |
| hsa-mir-193b-3p | HDHD5 | CLASH |
| hsa-mir-193b-3p | DCPS | CLASH |
| hsa-mir-193b-3p | MRPL42 | Microarray |
| hsa-mir-193b-3p | DBNL | CLASH |
| hsa-mir-193b-3p | ATAD2 | CLASH |
| hsa-mir-193b-3p | ANKRD11 | CLASH |
| hsa-mir-193b-3p | RACGAP1 | Microarray |
| hsa-mir-193b-3p | ABT1 | CLASH |
| hsa-mir-193b-3p | NCAPH2 | Microarray |
| hsa-mir-193b-3p | SENP1 | Microarray |
| hsa-mir-193b-3p | PSMC3IP | Microarray |
| hsa-mir-193b-3p | GMPPB | CLASH |
| hsa-mir-193b-3p | SEC61A1 | Microarray |
| hsa-mir-193b-3p | PKN3 | Microarray |
| hsa-mir-193b-3p | PRICKLE4 | CLASH |
| hsa-mir-193b-3p | UBQLN2 | CLASH |
| hsa-mir-193b-3p | DONSON | Microarray |
| hsa-mir-193b-3p | LMCD1 | Microarray |
| hsa-mir-193b-3p | BICRA | CLASH |
| hsa-mir-193b-3p | GEMIN4 | CLASH |
| hsa-mir-193b-3p | HP1BP3 | CLASH |
| hsa-mir-193b-3p | NSDHL | Proteomics |
| hsa-mir-193b-3p | GMNN | Microarray |
| hsa-mir-193b-3p | NMD3 | Proteomics |
| hsa-mir-193b-3p | NOSIP | Proteomics |
| hsa-mir-193b-3p | MRPL4 | CLASH//Proteomics |
| hsa-mir-193b-3p | UTP18 | HITS-CLIP |
| hsa-mir-193b-3p | ASB3 | HITS-CLIP |
| hsa-mir-193b-3p | LEF1 | Microarray |
| hsa-mir-193b-3p | GMPR2 | CLASH |
| hsa-mir-193b-3p | TAOK3 | CLASH |
| hsa-mir-193b-3p | SNX9 | Microarray |
| hsa-mir-193b-3p | ANKFY1 | HITS-CLIP |
| hsa-mir-193b-3p | HACD3 | Proteomics |
| hsa-mir-193b-3p | CTDSPL2 | CLASH |
| hsa-mir-193b-3p | SCLY | CLASH |

| | | |
|---|---|---|
| hsa-mir-193b-3p | NT5DC3 | Microarray |
| hsa-mir-193b-3p | METTL13 | CLASH |
| hsa-mir-193b-3p | TAF9B | CLASH |
| hsa-mir-193b-3p | PTRH2 | CLASH |
| hsa-mir-193b-3p | GINS2 | Microarray |
| hsa-mir-193b-3p | CDK12 | CLASH |
| hsa-mir-193b-3p | RSF1 | Microarray |
| hsa-mir-193b-3p | C11orf24 | Microarray |
| hsa-mir-193b-3p | A4GALT | Microarray |
| hsa-mir-193b-3p | ERRFI1 | Microarray |
| hsa-mir-193b-3p | SLC38A2 | CLASH |
| hsa-mir-193b-3p | TOLLIP | CLASH |
| hsa-mir-193b-3p | CHPF2 | CLASH |
| hsa-mir-193b-3p | FBXL19 | CLASH |
| hsa-mir-193b-3p | EPDR1 | Microarray |
| hsa-mir-193b-3p | IL17RD | CLASH |
| hsa-mir-193b-3p | ALKBH4 | CLASH |
| hsa-mir-193b-3p | SNRK | CLASH |
| hsa-mir-193b-3p | NSUN2 | CLASH |
| hsa-mir-193b-3p | ALKBH5 | HITS-CLIP |
| hsa-mir-193b-3p | INO80D | PAR-CLIP |
| hsa-mir-193b-3p | SEMA4C | CLASH |
| hsa-mir-193b-3p | SARS2 | CLASH |
| hsa-mir-193b-3p | PARP16 | Microarray |
| hsa-mir-193b-3p | HEATR3 | CLASH |
| hsa-mir-193b-3p | CDCA4 | Microarray |
| hsa-mir-193b-3p | C9orf40 | Microarray |
| hsa-mir-193b-3p | WDYHV1 | Microarray |
| hsa-mir-193b-3p | ELP3 | CLASH |
| hsa-mir-193b-3p | UBR7 | Microarray |
| hsa-mir-193b-3p | ARMC1 | HITS-CLIP |
| hsa-mir-193b-3p | TMEM33 | Proteomics |
| hsa-mir-193b-3p | CENPQ | Microarray |
| hsa-mir-193b-3p | RMDN3 | CLASH |
| hsa-mir-193b-3p | FANCI | Microarray |
| hsa-mir-193b-3p | ADI1 | Proteomics |
| hsa-mir-193b-3p | C5orf22 | PAR-CLIP |
| hsa-mir-193b-3p | CHDH | Microarray |
| hsa-mir-193b-3p | COPRS | Microarray |
| hsa-mir-193b-3p | MCM10 | CLASH//Microarray |
| hsa-mir-193b-3p | STRADB | Microarray |
| hsa-mir-193b-3p | SVOP | HITS-CLIP |
| hsa-mir-193b-3p | ZNF823 | Microarray |
| hsa-mir-193b-3p | LRRC40 | Microarray |
| hsa-mir-193b-3p | HIF1AN | CLASH |
| hsa-mir-193b-3p | IWS1 | CLASH |
| hsa-mir-193b-3p | ASF1B | Microarray |

| | | |
|---|---|---|
| hsa-mir-193b-3p | C1orf112 | Microarray |
| hsa-mir-193b-3p | HHAT | Microarray |
| hsa-mir-193b-3p | DNAJC11 | CLASH |
| hsa-mir-193b-3p | ENAH | CLASH |
| hsa-mir-193b-3p | AP5M1 | Microarray |
| hsa-mir-193b-3p | TMEM30A | HITS-CLIP |
| hsa-mir-193b-3p | WDR12 | CLASH |
| hsa-mir-193b-3p | TDP1 | CLASH |
| hsa-mir-193b-3p | CENPJ | Microarray |
| hsa-mir-193b-3p | ASH1L | CLASH |
| hsa-mir-193b-3p | ZNF395 | Microarray |
| hsa-mir-193b-3p | FAM212B | CLASH |
| hsa-mir-193b-3p | LIN37 | CLASH |
| hsa-mir-193b-3p | SULF2 | Microarray |
| hsa-mir-193b-3p | UBFD1 | HITS-CLIP |
| hsa-mir-193b-3p | MEPCE | CLASH |
| hsa-mir-193b-3p | LRRC8A | Microarray |
| hsa-mir-193b-3p | TM9SF3 | Proteomics |
| hsa-mir-193b-3p | STARD7 | CLASH//HITS-CLIP//Microarray |
| hsa-mir-193b-3p | SMCO4 | Microarray |
| hsa-mir-193b-3p | ARNTL2 | Microarray |
| hsa-mir-193b-3p | RGMA | PAR-CLIP |
| hsa-mir-193b-3p | ATXN7L3 | CLASH |
| hsa-mir-193b-3p | KIF15 | Microarray |
| hsa-mir-193b-3p | CIAPIN1 | CLASH |
| hsa-mir-193b-3p | TIGAR | CLASH//HITS-CLIP |
| hsa-mir-193b-3p | SELENON | Microarray |
| hsa-mir-193b-3p | LYRM2 | PAR-CLIP |
| hsa-mir-193b-3p | S100A14 | Proteomics |
| hsa-mir-193b-3p | SPC25 | Microarray |
| hsa-mir-193b-3p | GATAD2B | CLASH |
| hsa-mir-193b-3p | CNOT6 | Microarray |
| hsa-mir-193b-3p | ZNF512B | CLASH//Microarray |
| hsa-mir-193b-3p | ESYT2 | Proteomics |
| hsa-mir-193b-3p | HACE1 | Microarray |
| hsa-mir-193b-3p | NUFIP2 | HITS-CLIP |
| hsa-mir-193b-3p | MIB1 | CLASH |
| hsa-mir-193b-3p | KLHL42 | PAR-CLIP |
| hsa-mir-193b-3p | CRAMP1 | CLASH |
| hsa-mir-193b-3p | DHX37 | CLASH |
| hsa-mir-193b-3p | PHRF1 | CLASH |
| hsa-mir-193b-3p | ZNF317 | CLASH |
| hsa-mir-193b-3p | FANCM | Microarray |
| hsa-mir-193b-3p | C6orf47 | PAR-CLIP |
| hsa-mir-193b-3p | ZNF71 | CLASH |
| hsa-mir-193b-3p | SINHCAF | CLASH |
| hsa-mir-193b-3p | PLEKHA2 | HITS-CLIP |

| | | |
|---|---|---|
| hsa-mir-193b-3p | EEFSEC | CLASH |
| hsa-mir-193b-3p | HEATR6 | Proteomics |
| hsa-mir-193b-3p | FAM111A | Microarray |
| hsa-mir-193b-3p | ELMO2 | HITS-CLIP//Microarray |
| hsa-mir-193b-3p | CHTF18 | Microarray |
| hsa-mir-193b-3p | MCCC2 | Proteomics |
| hsa-mir-193b-3p | TMBIM1 | CLASH |
| hsa-mir-193b-3p | DUS1L | CLASH |
| hsa-mir-193b-3p | NCAPG | Microarray |
| hsa-mir-193b-3p | IRF2BPL | PAR-CLIP |
| hsa-mir-193b-3p | NFKBIZ | CLASH |
| hsa-mir-193b-3p | ZMAT3 | PAR-CLIP |
| hsa-mir-193b-3p | MTMR14 | CLASH |
| hsa-mir-193b-3p | GIGYF1 | CLASH |
| hsa-mir-193b-3p | ANAPC1 | CLASH |
| hsa-mir-193b-3p | NUCKS1 | CLASH |
| hsa-mir-193b-3p | C16orf58 | CLASH |
| hsa-mir-193b-3p | C6orf106 | HITS-CLIP |
| hsa-mir-193b-3p | GINS3 | Microarray |
| hsa-mir-193b-3p | TUT1 | CLASH |
| hsa-mir-193b-3p | MRPS9 | Proteomics |
| hsa-mir-193b-3p | MRPL32 | CLASH |
| hsa-mir-193b-3p | NOL6 | CLASH |
| hsa-mir-193b-3p | SOWAHC | CLASH |
| hsa-mir-193b-3p | WNK1 | CLASH |
| hsa-mir-193b-3p | UBE2Z | CLASH |
| hsa-mir-193b-3p | FUNDC2 | CLASH |
| hsa-mir-193b-3p | TRAK2 | CLASH |
| hsa-mir-193b-3p | PCYOX1L | Microarray |
| hsa-mir-193b-3p | AUNIP | Microarray |
| hsa-mir-193b-3p | NUP37 | CLASH |
| hsa-mir-193b-3p | C7orf26 | CLASH |
| hsa-mir-193b-3p | KXD1 | CLASH |
| hsa-mir-193b-3p | TSEN34 | CLASH |
| hsa-mir-193b-3p | METTL16 | CLASH |
| hsa-mir-193b-3p | DSCC1 | Microarray |
| hsa-mir-193b-3p | BRCC3 | CLASH |
| hsa-mir-193b-3p | TANGO6 | CLASH |
| hsa-mir-193b-3p | ARMC7 | CLASH |
| hsa-mir-193b-3p | MAP7D3 | Microarray |
| hsa-mir-193b-3p | TMEM204 | Microarray |
| hsa-mir-193b-3p | RTL10 | CLASH |
| hsa-mir-193b-3p | CENPU | Microarray |
| hsa-mir-193b-3p | MSANTD2 | PAR-CLIP |
| hsa-mir-193b-3p | QTRT2 | CLASH |
| hsa-mir-193b-3p | MORC4 | Microarray |
| hsa-mir-193b-3p | IPO4 | CLASH |

| | | |
|---|---|---|
| hsa-mir-193b-3p | ISOC2 | CLASH |
| hsa-mir-193b-3p | IQCA1 | CLASH |
| hsa-mir-193b-3p | MYH14 | CLASH |
| hsa-mir-193b-3p | C12orf49 | Microarray |
| hsa-mir-193b-3p | SHCBP1 | Microarray |
| hsa-mir-193b-3p | PIP4K2C | Proteomics |
| hsa-mir-193b-3p | BORA | Microarray |
| hsa-mir-193b-3p | LPCAT1 | HITS-CLIP//Microarray |
| hsa-mir-193b-3p | DSN1 | Microarray |
| hsa-mir-193b-3p | RMI1 | CLASH |
| hsa-mir-193b-3p | FBXL18 | CLASH |
| hsa-mir-193b-3p | VCPIP1 | CLASH |
| hsa-mir-193b-3p | PTGES2 | Proteomics |
| hsa-mir-193b-3p | CTC1 | HITS-CLIP//PAR-CLIP |
| hsa-mir-193b-3p | TEDC2 | Microarray |
| hsa-mir-193b-3p | EFHD1 | Microarray |
| hsa-mir-193b-3p | PUS1 | CLASH |
| hsa-mir-193b-3p | WDR82 | HITS-CLIP |
| hsa-mir-193b-3p | REEP4 | CLASH |
| hsa-mir-193b-3p | THAP7 | CLASH |
| hsa-mir-193b-3p | INTS5 | CLASH |
| hsa-mir-193b-3p | CEP44 | Microarray |
| hsa-mir-193b-3p | PTDSS2 | CLASH |
| hsa-mir-193b-3p | HM13 | CLASH |
| hsa-mir-193b-3p | YIPF5 | CLASH |
| hsa-mir-193b-3p | CLPB | PAR-CLIP |
| hsa-mir-193b-3p | APOLD1 | Microarray |
| hsa-mir-193b-3p | CDT1 | Microarray |
| hsa-mir-193b-3p | RNF146 | HITS-CLIP |
| hsa-mir-193b-3p | EMC6 | CLASH |
| hsa-mir-193b-3p | EPPK1 | Proteomics |
| hsa-mir-193b-3p | RASSF5 | CLASH |
| hsa-mir-193b-3p | SESN2 | Microarray |
| hsa-mir-193b-3p | LONP2 | CLASH |
| hsa-mir-193b-3p | RBM4B | CLASH |
| hsa-mir-193b-3p | TMTC1 | Microarray |
| hsa-mir-193b-3p | CDCA7 | Microarray |
| hsa-mir-193b-3p | KCTD10 | CLASH |
| hsa-mir-193b-3p | HASPIN | CLASH//Microarray |
| hsa-mir-193b-3p | SPRTN | PAR-CLIP |
| hsa-mir-193b-3p | KREMEN1 | CLASH |
| hsa-mir-193b-3p | MND1 | Microarray |
| hsa-mir-193b-3p | ENKD1 | CLASH |
| hsa-mir-193b-3p | TMEM164 | Microarray |
| hsa-mir-193b-3p | SLC25A33 | CLASH |
| hsa-mir-193b-3p | CCDC77 | Microarray |
| hsa-mir-193b-3p | ACBD6 | CLASH |

| | | |
|---|---|---|
| hsa-mir-193b-3p | LCOR | CLASH |
| hsa-mir-193b-3p | MCM8 | Microarray |
| hsa-mir-193b-3p | DCTN5 | PAR-CLIP |
| hsa-mir-193b-3p | KIAA1841 | HITS-CLIP |
| hsa-mir-193b-3p | PSRC1 | Microarray |
| hsa-mir-193b-3p | PRRC2B | CLASH |
| hsa-mir-193b-3p | LMNB2 | CLASH |
| hsa-mir-193b-3p | KLHL22 | CLASH |
| hsa-mir-193b-3p | NFATC2IP | CLASH |
| hsa-mir-193b-3p | ATOH8 | Microarray |
| hsa-mir-193b-3p | ARHGAP19 | CLASH//Microarray |
| hsa-mir-193b-3p | REPS1 | CLASH |
| hsa-mir-193b-3p | STRIP1 | CLASH |
| hsa-mir-193b-3p | TUBGCP6 | CLASH |
| hsa-mir-193b-3p | TICRR | Microarray |
| hsa-mir-193b-3p | CCDC32 | Microarray |
| hsa-mir-193b-3p | JPT2 | Proteomics |
| hsa-mir-193b-3p | MSANTD3 | Microarray |
| hsa-mir-193b-3p | PXYLP1 | Microarray |
| hsa-mir-193b-3p | MRRF | PAR-CLIP |
| hsa-mir-193b-3p | TIMM50 | Proteomics |
| hsa-mir-193b-3p | HAUS8 | Microarray |
| hsa-mir-193b-3p | MAPK1IP1L | CLASH |
| hsa-mir-193b-3p | CEP41 | Microarray |
| hsa-mir-193b-3p | MYL12B | CLASH |
| hsa-mir-193b-3p | SNX18 | CLASH |
| hsa-mir-193b-3p | MTFR2 | Microarray |
| hsa-mir-193b-3p | CDCA5 | Microarray |
| hsa-mir-193b-3p | SCAMP4 | CLASH |
| hsa-mir-193b-3p | CHST14 | Microarray |
| hsa-mir-193b-3p | NLRP3 | Microarray |
| hsa-mir-193b-3p | C1QTNF2 | Microarray |
| hsa-mir-193b-3p | ZNF618 | Microarray |
| hsa-mir-193b-3p | NT5C3B | CLASH |
| hsa-mir-193b-3p | ARHGAP33 | CLASH |
| hsa-mir-193b-3p | SLC18B1 | CLASH |
| hsa-mir-193b-3p | SSX2IP | Microarray |
| hsa-mir-193b-3p | PRAP1 | Luciferase reporter assay |
| hsa-mir-193b-3p | ZFYVE27 | CLASH |
| hsa-mir-193b-3p | SFXN2 | Microarray |
| hsa-mir-193b-3p | LRR1 | Microarray |
| hsa-mir-193b-3p | RAVER1 | Proteomics |
| hsa-mir-193b-3p | TYW3 | Microarray |
| hsa-mir-193b-3p | AHSA2 | CLASH |
| hsa-mir-193b-3p | PPARGC1B | CLASH |
| hsa-mir-193b-3p | PRRC1 | Proteomics |
| hsa-mir-193b-3p | WDR36 | CLASH |

| | | |
|---|---|---|
| hsa-mir-193b-3p | PM20D2 | Microarray |
| hsa-mir-193b-3p | S100A16 | Proteomics |
| hsa-mir-193b-3p | CEP128 | Microarray |
| hsa-mir-193b-3p | TMC8 | Microarray |
| hsa-mir-193b-3p | TRIM16L | CLASH |
| hsa-mir-193b-3p | APCDD1 | Microarray |
| hsa-mir-193b-3p | TICAM1 | CLASH |
| hsa-mir-193b-3p | SLC30A7 | Microarray |
| hsa-mir-193b-3p | YDJC | CLASH |
| hsa-mir-193b-3p | FAM109B | Microarray |
| hsa-mir-193b-3p | CKAP2L | Microarray |
| hsa-mir-193b-3p | GAREM2 | CLASH |
| hsa-mir-193b-3p | ERFE | Microarray |
| hsa-mir-193b-3p | SGO1 | Microarray |
| hsa-mir-193b-3p | CDCA2 | Microarray |
| hsa-mir-193b-3p | SLC35G1 | Microarray |
| hsa-mir-193b-3p | PARP15 | HITS-CLIP |
| hsa-mir-193b-3p | DACT2 | CLASH |
| hsa-mir-193b-3p | ZNF384 | HITS-CLIP |
| hsa-mir-193b-3p | ASXL1 | CLASH |
| hsa-mir-193b-3p | RHOV | CLASH |
| hsa-mir-193b-3p | UBR1 | CLASH |
| hsa-mir-193b-3p | TXLNA | CLASH |
| hsa-mir-193b-3p | TET3 | HITS-CLIP |
| hsa-mir-193b-3p | HACD2 | Proteomics |
| hsa-mir-193b-3p | SMIM14 | PAR-CLIP |
| hsa-mir-193b-3p | TUBB | CLASH |
| hsa-mir-193b-3p | SENP5 | HITS-CLIP |
| hsa-mir-193b-3p | ATAD3C | CLASH |
| hsa-mir-193b-3p | UNC5B | CLASH |
| hsa-mir-193b-3p | CCNY | CLASH |
| hsa-mir-193b-3p | RTKN2 | Microarray |
| hsa-mir-193b-3p | SKA1 | Microarray |
| hsa-mir-193b-3p | SKA3 | CLASH |
| hsa-mir-193b-3p | DAGLB | Proteomics |
| hsa-mir-193b-3p | NAPEPLD | PAR-CLIP |
| hsa-mir-193b-3p | RNASEH1 | PAR-CLIP |
| hsa-mir-193b-3p | NEIL2 | CLASH |
| hsa-mir-193b-3p | GPATCH11 | Microarray |
| hsa-mir-193b-3p | ASPM | Microarray |
| hsa-mir-193b-3p | BCL9L | CLASH |
| hsa-mir-193b-3p | RASSF3 | Microarray |
| hsa-mir-193b-3p | GXYLT1 | HITS-CLIP |
| hsa-mir-193b-3p | WDR62 | Microarray |
| hsa-mir-193b-3p | RPL7L1 | CLASH |
| hsa-mir-193b-3p | HIST2H3A | CLASH |
| hsa-mir-193b-3p | NAT8L | CLASH |

| | | |
|---|---|---|
| hsa-mir-193b-3p | SLC10A6 | HITS-CLIP |
| hsa-mir-193b-3p | RFLNB | Microarray |
| hsa-mir-193b-3p | NHLRC2 | Microarray |
| hsa-mir-193b-3p | KMT5A | Microarray |
| hsa-mir-193b-3p | LOC391247 | Microarray |
| hsa-mir-193b-3p | FAM221B | HITS-CLIP |
| hsa-mir-193b-3p | MEX3D | PAR-CLIP |
| hsa-mir-193b-3p | MYO18A | CLASH |
| hsa-mir-193b-3p | CHCHD10 | CLASH |
| hsa-mir-193b-3p | TOMM5 | CLASH |
| hsa-mir-193b-3p | MXRA7 | CLASH |
| hsa-mir-193b-3p | NRARP | CLASH |
| hsa-mir-193b-3p | PEF1 | CLASH |
| hsa-mir-193b-3p | DHFRP1 | Microarray |
| hsa-mir-193b-3p | TMPPE | PAR-CLIP |
| hsa-mir-193b-3p | SHISA9 | PAR-CLIP |
| hsa-mir-193b-3p | ZNF814 | CLASH |
| hsa-mir-193b-3p | EPOP | CLASH |
| hsa-mir-193b-3p | HSBP1P2 | Microarray |
| hsa-mir-503-5p | AP2B1 | PAR-CLIP |
| hsa-mir-503-5p | ANK3 | HITS-CLIP |
| hsa-mir-503-5p | ATP5G3 | PAR-CLIP |
| hsa-mir-503-5p | ATP6V1B2 | CLASH |
| hsa-mir-503-5p | CCND1 | Immunohistochemistry//Luciferase reporter assay//Northern blot//PAR-CLIP//qRT-PCR//Western blot |
| hsa-mir-503-5p | BCL2 | Flow//Immunohistochemistry//Immunoprecipitaion//Luciferase reporter assay//qRT-PCR//Western blot |
| hsa-mir-503-5p | CA8 | PAR-CLIP |
| hsa-mir-503-5p | CANX | PAR-CLIP |
| hsa-mir-503-5p | CAPZA2 | PAR-CLIP |
| hsa-mir-503-5p | CCND2 | PAR-CLIP |
| hsa-mir-503-5p | CCND3 | Flow//qRT-PCR//Western blot |
| hsa-mir-503-5p | CCNE1 | Luciferase reporter assay//PAR-CLIP |
| hsa-mir-503-5p | CCNF | Luciferase reporter assay |
| hsa-mir-503-5p | CCNG2 | CLASH |
| hsa-mir-503-5p | CCNT1 | PAR-CLIP |
| hsa-mir-503-5p | CD40 | Luciferase reporter assay |
| hsa-mir-503-5p | CDC25A | Luciferase reporter assay//PAR-CLIP |
| hsa-mir-503-5p | CDKN1A | Luciferase reporter assay |
| hsa-mir-503-5p | CHEK1 | HITS-CLIP//Immunohistochemistry//Luciferase reporter assay//qRT-PCR//Western blot |
| hsa-mir-503-5p | CLTC | CLASH |
| hsa-mir-503-5p | COL1A1 | PAR-CLIP |
| hsa-mir-503-5p | CREBL2 | PAR-CLIP |
| hsa-mir-503-5p | CRK | PAR-CLIP |
| hsa-mir-503-5p | CTSD | CLASH |
| hsa-mir-503-5p | DDX3X | PAR-CLIP |
| hsa-mir-503-5p | DECR1 | PAR-CLIP |

| | | |
|---|---|---|
| hsa-mir-503-5p | DHFR | CLASH |
| hsa-mir-503-5p | DLST | CLASH |
| hsa-mir-503-5p | E2F3 | Luciferase reporter assay//Western blot |
| hsa-mir-503-5p | EFNB2 | PAR-CLIP |
| hsa-mir-503-5p | EPOR | HITS-CLIP |
| hsa-mir-503-5p | EXT1 | PAR-CLIP |
| hsa-mir-503-5p | FANCA | LacZ reporter assay//qRT-PCR |
| hsa-mir-503-5p | FARSA | CLASH |
| hsa-mir-503-5p | FGF2 | Immunohistochemistry//Luciferase reporter assay//Microarray//Northern blot//PAR-CLIP//qRT-PCR |
| hsa-mir-503-5p | FGF8 | Luciferase reporter assay//qRT-PCR//Western blot |
| hsa-mir-503-5p | FGFR1 | Immunohistochemistry//Luciferase reporter assay//Microarray//Northern blot//qRT-PCR |
| hsa-mir-503-5p | NR6A1 | PAR-CLIP |
| hsa-mir-503-5p | GNAT1 | HITS-CLIP//PAR-CLIP |
| hsa-mir-503-5p | GPR27 | PAR-CLIP |
| hsa-mir-503-5p | HDGF | CLASH |
| hsa-mir-503-5p | HNRNPA1 | HITS-CLIP |
| hsa-mir-503-5p | HNRNPK | CLASH |
| hsa-mir-503-5p | HSPA8 | PAR-CLIP |
| hsa-mir-503-5p | IGF1R | Immunohistochemistry//Luciferase reporter assay//qRT-PCR//Western blot |
| hsa-mir-503-5p | IKBKB | Immunohistochemistry//In situ hybridization//Luciferase reporter assay//qRT-PCR//Western blot |
| hsa-mir-503-5p | IVD | CLASH |
| hsa-mir-503-5p | JARID2 | PAR-CLIP |
| hsa-mir-503-5p | KIF5B | PAR-CLIP |
| hsa-mir-503-5p | KPNA3 | HITS-CLIP//PAR-CLIP |
| hsa-mir-503-5p | SMAD2 | HITS-CLIP |
| hsa-mir-503-5p | SMAD7 | PAR-CLIP |
| hsa-mir-503-5p | MCM7 | CLASH |
| hsa-mir-503-5p | MT1E | CLASH |
| hsa-mir-503-5p | MYB | Luciferase reporter assay//qRT-PCR//Western blot |
| hsa-mir-503-5p | MYO5A | PAR-CLIP |
| hsa-mir-503-5p | OCRL | HITS-CLIP |
| hsa-mir-503-5p | ORC4 | PAR-CLIP |
| hsa-mir-503-5p | CDK17 | PAR-CLIP |
| hsa-mir-503-5p | PHKA1 | HITS-CLIP//PAR-CLIP |
| hsa-mir-503-5p | PIK3R1 | HITS-CLIP//Immunohistochemistry//In situ hybridization//Luciferase reporter assay//PAR-CLIP//qRT-PCR//Western blot |
| hsa-mir-503-5p | PLAG1 | PAR-CLIP |
| hsa-mir-503-5p | CTSA | CLASH |
| hsa-mir-503-5p | PPM1A | PAR-CLIP |
| hsa-mir-503-5p | PPP2R5C | PAR-CLIP |
| hsa-mir-503-5p | PRKAR2A | PAR-CLIP |
| hsa-mir-503-5p | PTPRD | PAR-CLIP |
| hsa-mir-503-5p | MAP4K2 | PAR-CLIP |

| | | |
|---|---|---|
| hsa-mir-503-5p | REL | PAR-CLIP |
| hsa-mir-503-5p | RPL15 | CLASH |
| hsa-mir-503-5p | RPL18A | CLASH |
| hsa-mir-503-5p | RPS5 | CLASH |
| hsa-mir-503-5p | RPS6KA3 | CLASH |
| hsa-mir-503-5p | RS1 | HITS-CLIP |
| hsa-mir-503-5p | SALL1 | PAR-CLIP |
| hsa-mir-503-5p | SBF1 | CLASH |
| hsa-mir-503-5p | SIAH2 | CLASH |
| hsa-mir-503-5p | SKI | PAR-CLIP |
| hsa-mir-503-5p | SLC2A3 | PAR-CLIP |
| hsa-mir-503-5p | SNRPB2 | PAR-CLIP |
| hsa-mir-503-5p | SNTB2 | PAR-CLIP |
| hsa-mir-503-5p | SRPRA | HITS-CLIP//PAR-CLIP |
| hsa-mir-503-5p | MAP3K7 | PAR-CLIP |
| hsa-mir-503-5p | PPP1R11 | HITS-CLIP |
| hsa-mir-503-5p | TFAP2A | PAR-CLIP |
| hsa-mir-503-5p | TLL1 | PAR-CLIP |
| hsa-mir-503-5p | TRAPPC10 | PAR-CLIP |
| hsa-mir-503-5p | UGT2B4 | PAR-CLIP |
| hsa-mir-503-5p | VEGFA | HITS-CLIP//Luciferase reporter assay//PAR-CLIP |
| hsa-mir-503-5p | EIF4H | PAR-CLIP |
| hsa-mir-503-5p | WEE1 | Luciferase reporter assay//PAR-CLIP |
| hsa-mir-503-5p | ZNF282 | PAR-CLIP |
| hsa-mir-503-5p | RECK | PAR-CLIP |
| hsa-mir-503-5p | CUL3 | HITS-CLIP |
| hsa-mir-503-5p | CBX4 | PAR-CLIP |
| hsa-mir-503-5p | CDC14A | Luciferase reporter assay |
| hsa-mir-503-5p | CASK | HITS-CLIP//PAR-CLIP |
| hsa-mir-503-5p | SLC25A12 | PAR-CLIP |
| hsa-mir-503-5p | DNAH17 | PAR-CLIP |
| hsa-mir-503-5p | NAPG | HITS-CLIP//PAR-CLIP |
| hsa-mir-503-5p | TNFRSF11A | Luciferase reporter assay//Microarray//Northern blot//qRT-PCR//Western blot |
| hsa-mir-503-5p | MTMR3 | PAR-CLIP |
| hsa-mir-503-5p | CCNE2 | Luciferase reporter assay//PAR-CLIP |
| hsa-mir-503-5p | SLC33A1 | HITS-CLIP |
| hsa-mir-503-5p | B4GALT5 | CLASH//PAR-CLIP |
| hsa-mir-503-5p | GLP2R | HITS-CLIP |
| hsa-mir-503-5p | RPL23 | CLASH |
| hsa-mir-503-5p | KIF23 | PAR-CLIP |
| hsa-mir-503-5p | SOCS5 | PAR-CLIP |
| hsa-mir-503-5p | N4BP1 | PAR-CLIP |
| hsa-mir-503-5p | BZW1 | PAR-CLIP |
| hsa-mir-503-5p | TOMM20 | CLASH |
| hsa-mir-503-5p | TSC22D2 | PAR-CLIP |
| hsa-mir-503-5p | TLK1 | PAR-CLIP |

| | | |
|---|---|---|
| hsa-mir-503-5p | SEC16A | PAR-CLIP |
| hsa-mir-503-5p | DCLRE1A | CLASH |
| hsa-mir-503-5p | DMTF1 | PAR-CLIP |
| hsa-mir-503-5p | AKT3 | PAR-CLIP |
| hsa-mir-503-5p | ACTR2 | PAR-CLIP |
| hsa-mir-503-5p | RNF41 | PAR-CLIP |
| hsa-mir-503-5p | CTDSPL | PAR-CLIP |
| hsa-mir-503-5p | LANCL1 | PAR-CLIP |
| hsa-mir-503-5p | BTN3A3 | PAR-CLIP |
| hsa-mir-503-5p | AHSA1 | CLASH |
| hsa-mir-503-5p | HEXIM1 | CLASH |
| hsa-mir-503-5p | NUP50 | PAR-CLIP |
| hsa-mir-503-5p | SEC24A | PAR-CLIP |
| hsa-mir-503-5p | PNPLA6 | PAR-CLIP |
| hsa-mir-503-5p | PRSS21 | PAR-CLIP |
| hsa-mir-503-5p | PRDM4 | PAR-CLIP |
| hsa-mir-503-5p | TRAK1 | HITS-CLIP |
| hsa-mir-503-5p | ATF6 | Luciferase reporter assay |
| hsa-mir-503-5p | SETD1B | PAR-CLIP |
| hsa-mir-503-5p | TRIM35 | PAR-CLIP |
| hsa-mir-503-5p | KANK1 | PAR-CLIP |
| hsa-mir-503-5p | DDHD2 | PAR-CLIP |
| hsa-mir-503-5p | LARP1 | PAR-CLIP |
| hsa-mir-503-5p | SF3B3 | CLASH |
| hsa-mir-503-5p | NNT | PAR-CLIP |
| hsa-mir-503-5p | CD2AP | PAR-CLIP |
| hsa-mir-503-5p | TMEM245 | HITS-CLIP//PAR-CLIP |
| hsa-mir-503-5p | PISD | PAR-CLIP |
| hsa-mir-503-5p | MOB4 | PAR-CLIP |
| hsa-mir-503-5p | SZRD1 | PAR-CLIP |
| hsa-mir-503-5p | WIPI2 | PAR-CLIP |
| hsa-mir-503-5p | ago-01 | Luciferase reporter assay |
| hsa-mir-503-5p | ZBTB44 | PAR-CLIP |
| hsa-mir-503-5p | HCFC2 | PAR-CLIP |
| hsa-mir-503-5p | SEC61A1 | PAR-CLIP |
| hsa-mir-503-5p | PSAT1 | PAR-CLIP |
| hsa-mir-503-5p | PIK3R4 | CLASH |
| hsa-mir-503-5p | CUZD1 | CLASH |
| hsa-mir-503-5p | ASCC1 | PAR-CLIP |
| hsa-mir-503-5p | ZNF691 | PAR-CLIP |
| hsa-mir-503-5p | NT5DC3 | HITS-CLIP |
| hsa-mir-503-5p | MRPS23 | CLASH |
| hsa-mir-503-5p | CDK12 | CLASH |
| hsa-mir-503-5p | DNAJC10 | PAR-CLIP |
| hsa-mir-503-5p | ANLN | Luciferase reporter assay |
| hsa-mir-503-5p | USP53 | PAR-CLIP |
| hsa-mir-503-5p | CNNM2 | PAR-CLIP |

| | | |
|---|---|---|
| hsa-mir-503-5p | CDCA4 | PAR-CLIP |
| hsa-mir-503-5p | RIF1 | PAR-CLIP |
| hsa-mir-503-5p | SBNO1 | PAR-CLIP |
| hsa-mir-503-5p | MRGBP | CLASH |
| hsa-mir-503-5p | TMEM100 | PAR-CLIP |
| hsa-mir-503-5p | FBXW7 | Luciferase reporter assay//Western blot |
| hsa-mir-503-5p | PI4K2B | PAR-CLIP |
| hsa-mir-503-5p | RFK | PAR-CLIP |
| hsa-mir-503-5p | PNRC2 | PAR-CLIP |
| hsa-mir-503-5p | CDC37L1 | HITS-CLIP//PAR-CLIP |
| hsa-mir-503-5p | NPLOC4 | PAR-CLIP |
| hsa-mir-503-5p | ASH1L | PAR-CLIP |
| hsa-mir-503-5p | UBFD1 | CLASH |
| hsa-mir-503-5p | CYP26B1 | PAR-CLIP |
| hsa-mir-503-5p | CDC42SE2 | PAR-CLIP |
| hsa-mir-503-5p | RTN4 | PAR-CLIP |
| hsa-mir-503-5p | RALGAPB | PAR-CLIP |
| hsa-mir-503-5p | ACTR3B | PAR-CLIP |
| hsa-mir-503-5p | ODF2L | PAR-CLIP |
| hsa-mir-503-5p | NUFIP2 | PAR-CLIP |
| hsa-mir-503-5p | TAOK1 | PAR-CLIP |
| hsa-mir-503-5p | KIAA1456 | PAR-CLIP |
| hsa-mir-503-5p | DLGAP3 | PAR-CLIP |
| hsa-mir-503-5p | PLEKHA1 | PAR-CLIP |
| hsa-mir-503-5p | GREM2 | PAR-CLIP |
| hsa-mir-503-5p | ZMAT3 | PAR-CLIP |
| hsa-mir-503-5p | NUCKS1 | CLASH |
| hsa-mir-503-5p | CERK | PAR-CLIP |
| hsa-mir-503-5p | TUT1 | CLASH |
| hsa-mir-503-5p | PAPOLG | PAR-CLIP |
| hsa-mir-503-5p | RAPH1 | PAR-CLIP |
| hsa-mir-503-5p | WNK3 | PAR-CLIP |
| hsa-mir-503-5p | CHAC1 | PAR-CLIP |
| hsa-mir-503-5p | DCAF10 | PAR-CLIP |
| hsa-mir-503-5p | YRDC | PAR-CLIP |
| hsa-mir-503-5p | ZFHX4 | HITS-CLIP//PAR-CLIP |
| hsa-mir-503-5p | ATAD5 | PAR-CLIP |
| hsa-mir-503-5p | L2HGDH | PAR-CLIP |
| hsa-mir-503-5p | FBXL18 | PAR-CLIP |
| hsa-mir-503-5p | DCAF17 | HITS-CLIP |
| hsa-mir-503-5p | VOPP1 | PAR-CLIP |
| hsa-mir-503-5p | RCC1L | CLASH |
| hsa-mir-503-5p | C1orf21 | PAR-CLIP |
| hsa-mir-503-5p | TXNDC5 | PAR-CLIP |
| hsa-mir-503-5p | GSG1 | PAR-CLIP |
| hsa-mir-503-5p | BCL2L12 | PAR-CLIP |
| hsa-mir-503-5p | SPNS1 | CLASH |

| | | |
|---|---|---|
| hsa-mir-503-5p | ZNRF3 | PAR-CLIP |
| hsa-mir-503-5p | USP48 | PAR-CLIP |
| hsa-mir-503-5p | DCTN5 | PAR-CLIP |
| hsa-mir-503-5p | CBX2 | PAR-CLIP |
| hsa-mir-503-5p | KIAA1671 | CLASH |
| hsa-mir-503-5p | ZNF622 | PAR-CLIP |
| hsa-mir-503-5p | SESTD1 | PAR-CLIP |
| hsa-mir-503-5p | YTHDC1 | PAR-CLIP |
| hsa-mir-503-5p | ELMSAN1 | PAR-CLIP |
| hsa-mir-503-5p | SCAMP4 | PAR-CLIP |
| hsa-mir-503-5p | RAB3IP | PAR-CLIP |
| hsa-mir-503-5p | MRPL10 | CLASH |
| hsa-mir-503-5p | OSCAR | PAR-CLIP |
| hsa-mir-503-5p | UHMK1 | CLASH |
| hsa-mir-503-5p | LIX1L | CLASH |
| hsa-mir-503-5p | ARHGEF19 | Luciferase reporter assay//qRT-PCR//Western blot |
| hsa-mir-503-5p | UBR3 | HITS-CLIP//PAR-CLIP |
| hsa-mir-503-5p | MTPN | CLASH |
| hsa-mir-503-5p | DYNLL2 | PAR-CLIP |
| hsa-mir-503-5p | SREK1 | HITS-CLIP//PAR-CLIP |
| hsa-mir-503-5p | CACUL1 | PAR-CLIP |
| hsa-mir-503-5p | HNRNPA1L2 | HITS-CLIP |
| hsa-mir-503-5p | CMTM4 | CLASH |
| hsa-mir-503-5p | CREBRF | PAR-CLIP |
| hsa-mir-503-5p | CNKSR3 | PAR-CLIP |
| hsa-mir-503-5p | SPRED1 | PAR-CLIP |
| hsa-mir-503-5p | ZNF367 | HITS-CLIP//PAR-CLIP |
| hsa-mir-503-5p | ZNF449 | PAR-CLIP |
| hsa-mir-503-5p | CCDC83 | PAR-CLIP |
| hsa-mir-503-5p | FOXK1 | PAR-CLIP |
| hsa-mir-503-5p | LRWD1 | PAR-CLIP |
| hsa-mir-503-5p | ZNRF2 | PAR-CLIP |
| hsa-mir-503-5p | ZNF620 | PAR-CLIP |
| hsa-mir-503-5p | UBN2 | PAR-CLIP |
| hsa-mir-503-5p | RNF149 | HITS-CLIP//PAR-CLIP |
| hsa-mir-503-5p | LURAP1L | PAR-CLIP |
| hsa-mir-503-5p | XKR7 | PAR-CLIP |
| hsa-mir-503-5p | NDUFA4P1 | PAR-CLIP |
| hsa-mir-503-5p | ZBTB34 | PAR-CLIP |
| hsa-mir-503-5p | FAM229B | PAR-CLIP |
| hsa-mir-503-5p | ZNF704 | PAR-CLIP |
| hsa-mir-503-5p | KIAA0895L | CLASH |
| hsa-mir-503-5p | HSPE1-MOB4 | PAR-CLIP |
| hsa-mir-455-5p | APLP2 | PAR-CLIP |
| hsa-mir-455-5p | TRIM23 | CLASH |
| hsa-mir-455-5p | RHOH | HITS-CLIP |
| hsa-mir-455-5p | RUNX1T1 | HITS-CLIP//PAR-CLIP |

| | | |
|---|---|---|
| hsa-mir-455-5p | CD36 | HITS-CLIP |
| hsa-mir-455-5p | CDKN1B | PAR-CLIP |
| hsa-mir-455-5p | CRKL | PAR-CLIP |
| hsa-mir-455-5p | DDX3X | PAR-CLIP |
| hsa-mir-455-5p | DYRK1A | HITS-CLIP |
| hsa-mir-455-5p | ERCC4 | PAR-CLIP |
| hsa-mir-455-5p | ETS2 | PAR-CLIP |
| hsa-mir-455-5p | HOXA1 | PAR-CLIP |
| hsa-mir-455-5p | KPNA3 | HITS-CLIP//PAR-CLIP |
| hsa-mir-455-5p | MAP3K9 | PAR-CLIP |
| hsa-mir-455-5p | MYBL1 | PAR-CLIP |
| hsa-mir-455-5p | DRG1 | CLASH |
| hsa-mir-455-5p | OTX1 | HITS-CLIP//PAR-CLIP |
| hsa-mir-455-5p | PCCA | CLASH |
| hsa-mir-455-5p | PCCB | HITS-CLIP |
| hsa-mir-455-5p | PIK3R1 | HITS-CLIP//PAR-CLIP |
| hsa-mir-455-5p | PTPRB | PAR-CLIP |
| hsa-mir-455-5p | REL | HITS-CLIP |
| hsa-mir-455-5p | RPS6KB1 | PAR-CLIP//qRT-PCR |
| hsa-mir-455-5p | RPS14 | PAR-CLIP |
| hsa-mir-455-5p | SLC1A5 | HITS-CLIP |
| hsa-mir-455-5p | SOX11 | PAR-CLIP |
| hsa-mir-455-5p | UBA7 | Luciferase reporter assay//qRT-PCR//Western blot |
| hsa-mir-455-5p | ZNF134 | PAR-CLIP |
| hsa-mir-455-5p | ZNF138 | PAR-CLIP |
| hsa-mir-455-5p | ZFAND5 | PAR-CLIP |
| hsa-mir-455-5p | SOCS3 | qRT-PCR |
| hsa-mir-455-5p | TXNL1 | PAR-CLIP |
| hsa-mir-455-5p | QKI | PAR-CLIP |
| hsa-mir-455-5p | PCLAF | HITS-CLIP |
| hsa-mir-455-5p | RASSF2 | PAR-CLIP |
| hsa-mir-455-5p | G3BP1 | PAR-CLIP |
| hsa-mir-455-5p | VAV3 | HITS-CLIP//PAR-CLIP |
| hsa-mir-455-5p | IPO7 | PAR-CLIP |
| hsa-mir-455-5p | ZNF460 | PAR-CLIP |
| hsa-mir-455-5p | CCNI | HITS-CLIP |
| hsa-mir-455-5p | POLI | PAR-CLIP |
| hsa-mir-455-5p | FKBP9 | HITS-CLIP |
| hsa-mir-455-5p | PLEKHA6 | PAR-CLIP |
| hsa-mir-455-5p | SEPHS1 | PAR-CLIP |
| hsa-mir-455-5p | RAB18 | Luciferase reporter assay//qRT-PCR//Western blot |
| hsa-mir-455-5p | IGSF9B | HITS-CLIP |
| hsa-mir-455-5p | FBXO28 | PAR-CLIP |
| hsa-mir-455-5p | ARC | PAR-CLIP |
| hsa-mir-455-5p | MGRN1 | PAR-CLIP |
| hsa-mir-455-5p | NCSTN | ELISA//Luciferase reporter assay |
| hsa-mir-455-5p | PRKD2 | HITS-CLIP |

| | | |
|---|---|---|
| hsa-mir-455-5p | LYPD3 | HITS-CLIP |
| hsa-mir-455-5p | ZNF544 | PAR-CLIP |
| hsa-mir-455-5p | MYLIP | PAR-CLIP |
| hsa-mir-455-5p | ZNF354C | PAR-CLIP |
| hsa-mir-455-5p | ZNF117 | PAR-CLIP |
| hsa-mir-455-5p | TRPV2 | PAR-CLIP |
| hsa-mir-455-5p | TRIM33 | PAR-CLIP |
| hsa-mir-455-5p | NUP54 | HITS-CLIP//PAR-CLIP |
| hsa-mir-455-5p | PRR13 | HITS-CLIP |
| hsa-mir-455-5p | DDX4 | PAR-CLIP |
| hsa-mir-455-5p | BNC2 | HITS-CLIP |
| hsa-mir-455-5p | AHI1 | PAR-CLIP |
| hsa-mir-455-5p | PIWIL2 | PAR-CLIP |
| hsa-mir-455-5p | MTPAP | PAR-CLIP |
| hsa-mir-455-5p | DARS2 | HITS-CLIP |
| hsa-mir-455-5p | DEPDC1B | PAR-CLIP |
| hsa-mir-455-5p | NUFIP2 | PAR-CLIP |
| hsa-mir-455-5p | FAM160B1 | PAR-CLIP |
| hsa-mir-455-5p | ATP13A3 | PAR-CLIP |
| hsa-mir-455-5p | MOB3B | PAR-CLIP |
| hsa-mir-455-5p | TMC7 | PAR-CLIP |
| hsa-mir-455-5p | DSN1 | PAR-CLIP |
| hsa-mir-455-5p | WDR26 | PAR-CLIP |
| hsa-mir-455-5p | KLHL15 | PAR-CLIP |
| hsa-mir-455-5p | GSG1 | PAR-CLIP |
| hsa-mir-455-5p | SLC9A7 | PAR-CLIP |
| hsa-mir-455-5p | UBASH3B | PAR-CLIP |
| hsa-mir-455-5p | ZNF625 | PAR-CLIP |
| hsa-mir-455-5p | LYRM7 | HITS-CLIP |
| hsa-mir-455-5p | NT5C1B | PAR-CLIP |
| hsa-mir-455-5p | MOGAT1 | HITS-CLIP |
| hsa-mir-455-5p | C15orf40 | PAR-CLIP |
| hsa-mir-455-5p | TMEM170A | PAR-CLIP |
| hsa-mir-455-5p | TNFAIP8L1 | HITS-CLIP |
| hsa-mir-455-5p | PABPC4L | PAR-CLIP |
| hsa-mir-455-5p | UBXN2B | PAR-CLIP |
| hsa-mir-455-5p | LETM2 | HITS-CLIP |
| hsa-mir-455-5p | TRUB1 | PAR-CLIP |
| hsa-mir-455-5p | BCDIN3D | CLASH |
| hsa-mir-455-5p | TCF23 | PAR-CLIP |
| hsa-mir-455-5p | CAMSAP1 | HITS-CLIP |
| hsa-mir-455-5p | DNAJC18 | HITS-CLIP |
| hsa-mir-455-5p | PATL1 | PAR-CLIP |
| hsa-mir-455-5p | YIPF6 | PAR-CLIP |
| hsa-mir-455-5p | ZNF772 | PAR-CLIP |
| hsa-mir-455-5p | ZBTB34 | HITS-CLIP |
| hsa-mir-455-5p | FAM229B | PAR-CLIP |

| | | |
|---|---|---|
| hsa-mir-455-5p | NT5C1B-RDH14 | PAR-CLIP |
| hsa-mir-31-3p | ACADVL | CLASH |
| hsa-mir-31-3p | RHOA | qRT-PCR//Western blot |
| hsa-mir-31-3p | BACH1 | HITS-CLIP |
| hsa-mir-31-3p | CDH13 | HITS-CLIP |
| hsa-mir-31-3p | CRK | PAR-CLIP |
| hsa-mir-31-3p | CRKL | PAR-CLIP |
| hsa-mir-31-3p | DCK | PAR-CLIP |
| hsa-mir-31-3p | E2F2 | Luciferase reporter assay |
| hsa-mir-31-3p | GABRB1 | HITS-CLIP |
| hsa-mir-31-3p | HSPA6 | HITS-CLIP//PAR-CLIP |
| hsa-mir-31-3p | INHBA | CLASH |
| hsa-mir-31-3p | MCM4 | CLASH |
| hsa-mir-31-3p | NUCB1 | HITS-CLIP |
| hsa-mir-31-3p | PAX6 | HITS-CLIP |
| hsa-mir-31-3p | PDE4D | PAR-CLIP |
| hsa-mir-31-3p | PPIA | CLASH |
| hsa-mir-31-3p | PPIC | PAR-CLIP |
| hsa-mir-31-3p | PPP2R5C | CLASH |
| hsa-mir-31-3p | RAN | CLASH |
| hsa-mir-31-3p | SDHA | Luciferase reporter assay//qRT-PCR//Western blot |
| hsa-mir-31-3p | SP1 | PAR-CLIP |
| hsa-mir-31-3p | TRAF1 | PAR-CLIP |
| hsa-mir-31-3p | XK | CLASH |
| hsa-mir-31-3p | KIAA0391 | PAR-CLIP |
| hsa-mir-31-3p | SUPT7L | PAR-CLIP |
| hsa-mir-31-3p | SLC9A6 | CLASH |
| hsa-mir-31-3p | ARL6IP5 | CLASH |
| hsa-mir-31-3p | TXNIP | PAR-CLIP |
| hsa-mir-31-3p | FRS2 | PAR-CLIP |
| hsa-mir-31-3p | DNAJB4 | PAR-CLIP |
| hsa-mir-31-3p | CBX3 | PAR-CLIP |
| hsa-mir-31-3p | NLGN1 | HITS-CLIP |
| hsa-mir-31-3p | NCBP2 | HITS-CLIP |
| hsa-mir-31-3p | RAB18 | PAR-CLIP |
| hsa-mir-31-3p | FBXL7 | PAR-CLIP |
| hsa-mir-31-3p | PUM2 | PAR-CLIP |
| hsa-mir-31-3p | DICER1 | qRT-PCR |
| hsa-mir-31-3p | MAFF | HITS-CLIP |
| hsa-mir-31-3p | FBXL5 | HITS-CLIP//PAR-CLIP |
| hsa-mir-31-3p | ago-02 | HITS-CLIP//PAR-CLIP |
| hsa-mir-31-3p | DBR1 | CLASH |
| hsa-mir-31-3p | BRWD1 | PAR-CLIP |
| hsa-mir-31-3p | EPB41L4B | PAR-CLIP |
| hsa-mir-31-3p | PLEKHB2 | PAR-CLIP |
| hsa-mir-31-3p | BLOC1S4 | HITS-CLIP |

| | | |
|---|---|---|
| hsa-mir-31-3p | RBM38 | PAR-CLIP |
| hsa-mir-31-3p | POLR3E | PAR-CLIP |
| hsa-mir-31-3p | ZNF71 | PAR-CLIP |
| hsa-mir-31-3p | SLC30A5 | PAR-CLIP |
| hsa-mir-31-3p | ZNF614 | PAR-CLIP |
| hsa-mir-31-3p | TNKS2 | CLASH |
| hsa-mir-31-3p | SLC38A1 | PAR-CLIP |
| hsa-mir-31-3p | NECTIN4 | Luciferase reporter assay//Western blot |
| hsa-mir-31-3p | C9orf64 | HITS-CLIP//PAR-CLIP |
| hsa-mir-31-3p | LMNB2 | PAR-CLIP |
| hsa-mir-31-3p | C1orf198 | CLASH |
| hsa-mir-31-3p | PRPF38A | PAR-CLIP |
| hsa-mir-31-3p | CIPC | CLASH |
| hsa-mir-31-3p | PNPT1 | HITS-CLIP |
| hsa-mir-31-3p | PAPLN | PAR-CLIP |
| hsa-mir-31-3p | NUS1 | CLASH |
| hsa-mir-31-3p | CHMP4B | PAR-CLIP |
| hsa-mir-31-3p | PTPDC1 | PAR-CLIP |
| hsa-mir-31-3p | SPRED1 | PAR-CLIP |
| hsa-mir-31-3p | TET3 | HITS-CLIP |
| hsa-mir-31-3p | SLC16A9 | PAR-CLIP |
| hsa-mir-31-3p | ZNF485 | HITS-CLIP//PAR-CLIP |
| hsa-mir-31-3p | RICTOR | CLASH |
| hsa-mir-31-3p | RPL7L1 | PAR-CLIP |
| hsa-mir-31-3p | NUP43 | PAR-CLIP |
| hsa-mir-193b-5p | ADD1 | PAR-CLIP |
| hsa-mir-193b-5p | GRK3 | HITS-CLIP |
| hsa-mir-193b-5p | ALDOA | PAR-CLIP |
| hsa-mir-193b-5p | ART4 | HITS-CLIP |
| hsa-mir-193b-5p | ATP5G1 | HITS-CLIP |
| hsa-mir-193b-5p | BMP7 | HITS-CLIP |
| hsa-mir-193b-5p | BMP8B | PAR-CLIP |
| hsa-mir-193b-5p | ZFP36L1 | PAR-CLIP |
| hsa-mir-193b-5p | CACNG1 | PAR-CLIP |
| hsa-mir-193b-5p | COL9A2 | PAR-CLIP |
| hsa-mir-193b-5p | CRKL | CLASH |
| hsa-mir-193b-5p | DNASE2 | HITS-CLIP//PAR-CLIP |
| hsa-mir-193b-5p | SLC26A2 | HITS-CLIP |
| hsa-mir-193b-5p | CLN8 | HITS-CLIP |
| hsa-mir-193b-5p | NR2F6 | PAR-CLIP |
| hsa-mir-193b-5p | FGB | PAR-CLIP |
| hsa-mir-193b-5p | FXN | PAR-CLIP |
| hsa-mir-193b-5p | GGCX | PAR-CLIP |
| hsa-mir-193b-5p | GLA | PAR-CLIP |
| hsa-mir-193b-5p | GUCA1B | HITS-CLIP |
| hsa-mir-193b-5p | HOXC8 | PAR-CLIP |
| hsa-mir-193b-5p | IFIT2 | Microarray//qRT-PCR |

| | | |
|---|---|---|
| hsa-mir-193b-5p | IFNAR1 | HITS-CLIP |
| hsa-mir-193b-5p | RBPJ | PAR-CLIP |
| hsa-mir-193b-5p | IPP | HITS-CLIP |
| hsa-mir-193b-5p | KCNA7 | PAR-CLIP |
| hsa-mir-193b-5p | STMN1 | Luciferase reporter assay//qRT-PCR//Western blot |
| hsa-mir-193b-5p | MGAT5 | HITS-CLIP |
| hsa-mir-193b-5p | ATXN3 | HITS-CLIP |
| hsa-mir-193b-5p | MYO1C | PAR-CLIP |
| hsa-mir-193b-5p | NDUFV3 | PAR-CLIP |
| hsa-mir-193b-5p | PAK3 | HITS-CLIP |
| hsa-mir-193b-5p | PRKN | PAR-CLIP |
| hsa-mir-193b-5p | PCYT1A | PAR-CLIP |
| hsa-mir-193b-5p | ATP8B1 | PAR-CLIP |
| hsa-mir-193b-5p | MAP2K2 | PAR-CLIP |
| hsa-mir-193b-5p | PSMB9 | PAR-CLIP |
| hsa-mir-193b-5p | QSOX1 | PAR-CLIP |
| hsa-mir-193b-5p | PURB | HITS-CLIP |
| hsa-mir-193b-5p | RAB3B | PAR-CLIP |
| hsa-mir-193b-5p | MAP4K2 | HITS-CLIP |
| hsa-mir-193b-5p | RAB13 | PAR-CLIP |
| hsa-mir-193b-5p | RAB27A | HITS-CLIP |
| hsa-mir-193b-5p | RPL4 | PAR-CLIP |
| hsa-mir-193b-5p | RPS19 | CLASH |
| hsa-mir-193b-5p | RRAD | PAR-CLIP |
| hsa-mir-193b-5p | SLC4A1 | PAR-CLIP |
| hsa-mir-193b-5p | THBS2 | HITS-CLIP |
| hsa-mir-193b-5p | TIAL1 | HITS-CLIP |
| hsa-mir-193b-5p | TMF1 | PAR-CLIP |
| hsa-mir-193b-5p | UGT2B4 | PAR-CLIP |
| hsa-mir-193b-5p | VHL | HITS-CLIP |
| hsa-mir-193b-5p | ZNF8 | PAR-CLIP |
| hsa-mir-193b-5p | ZNF708 | HITS-CLIP |
| hsa-mir-193b-5p | ZNF24 | PAR-CLIP |
| hsa-mir-193b-5p | ZNF138 | PAR-CLIP |
| hsa-mir-193b-5p | BTG2 | PAR-CLIP |
| hsa-mir-193b-5p | MLF2 | PAR-CLIP |
| hsa-mir-193b-5p | GAN | HITS-CLIP |
| hsa-mir-193b-5p | CLPP | PAR-CLIP |
| hsa-mir-193b-5p | CHAF1B | PAR-CLIP |
| hsa-mir-193b-5p | API5 | HITS-CLIP |
| hsa-mir-193b-5p | DEGS1 | PAR-CLIP |
| hsa-mir-193b-5p | GMPS | PAR-CLIP |
| hsa-mir-193b-5p | UBE4A | PAR-CLIP |
| hsa-mir-193b-5p | GSTO1 | PAR-CLIP |
| hsa-mir-193b-5p | AKAP6 | HITS-CLIP |
| hsa-mir-193b-5p | SOCS5 | PAR-CLIP |
| hsa-mir-193b-5p | NUP93 | HITS-CLIP |

| | | |
|---|---|---|
| hsa-mir-193b-5p | KIAA0586 | HITS-CLIP |
| hsa-mir-193b-5p | SLC35E2 | PAR-CLIP |
| hsa-mir-193b-5p | CCS | HITS-CLIP |
| hsa-mir-193b-5p | GNE | HITS-CLIP |
| hsa-mir-193b-5p | PLIN3 | HITS-CLIP |
| hsa-mir-193b-5p | LYPLA1 | HITS-CLIP |
| hsa-mir-193b-5p | HOXB13 | PAR-CLIP |
| hsa-mir-193b-5p | ANP32B | PAR-CLIP |
| hsa-mir-193b-5p | PDLIM5 | PAR-CLIP |
| hsa-mir-193b-5p | GMEB1 | HITS-CLIP |
| hsa-mir-193b-5p | SRSF10 | PAR-CLIP |
| hsa-mir-193b-5p | FTCD | HITS-CLIP |
| hsa-mir-193b-5p | SLC27A4 | PAR-CLIP |
| hsa-mir-193b-5p | RPP14 | PAR-CLIP |
| hsa-mir-193b-5p | RCAN3 | PAR-CLIP |
| hsa-mir-193b-5p | POLR3A | HITS-CLIP |
| hsa-mir-193b-5p | PHB2 | PAR-CLIP |
| hsa-mir-193b-5p | RHOBTB3 | HITS-CLIP |
| hsa-mir-193b-5p | RTF1 | PAR-CLIP |
| hsa-mir-193b-5p | JADE2 | PAR-CLIP |
| hsa-mir-193b-5p | PUM2 | PAR-CLIP |
| hsa-mir-193b-5p | TMEM245 | HITS-CLIP |
| hsa-mir-193b-5p | TNFAIP8 | HITS-CLIP |
| hsa-mir-193b-5p | BACE2 | HITS-CLIP |
| hsa-mir-193b-5p | HEATR5A | HITS-CLIP |
| hsa-mir-193b-5p | RSL1D1 | PAR-CLIP |
| hsa-mir-193b-5p | HSPB8 | PAR-CLIP |
| hsa-mir-193b-5p | TNRC6A | PAR-CLIP |
| hsa-mir-193b-5p | POLL | HITS-CLIP |
| hsa-mir-193b-5p | TOR2A | PAR-CLIP |
| hsa-mir-193b-5p | FLVCR1 | HITS-CLIP |
| hsa-mir-193b-5p | NTMT1 | HITS-CLIP |
| hsa-mir-193b-5p | ORMDL2 | HITS-CLIP |
| hsa-mir-193b-5p | PYCARD | PAR-CLIP |
| hsa-mir-193b-5p | A1CF | PAR-CLIP |
| hsa-mir-193b-5p | TRAT1 | HITS-CLIP |
| hsa-mir-193b-5p | WDPCP | HITS-CLIP |
| hsa-mir-193b-5p | ZC2HC1A | HITS-CLIP |
| hsa-mir-193b-5p | RDH11 | PAR-CLIP |
| hsa-mir-193b-5p | PHF20 | HITS-CLIP |
| hsa-mir-193b-5p | CRIM1 | Microarray//qRT-PCR |
| hsa-mir-193b-5p | TRPV2 | PAR-CLIP |
| hsa-mir-193b-5p | LARS | HITS-CLIP |
| hsa-mir-193b-5p | GDE1 | HITS-CLIP |
| hsa-mir-193b-5p | FXYD5 | PAR-CLIP |
| hsa-mir-193b-5p | CYCS | HITS-CLIP |
| hsa-mir-193b-5p | DNAJC10 | PAR-CLIP |

| | | |
|---|---|---|
| hsa-mir-193b-5p | SDK2 | HITS-CLIP |
| hsa-mir-193b-5p | GPN2 | PAR-CLIP |
| hsa-mir-193b-5p | PGPEP1 | PAR-CLIP |
| hsa-mir-193b-5p | DPP8 | HITS-CLIP |
| hsa-mir-193b-5p | RBM28 | PAR-CLIP |
| hsa-mir-193b-5p | KIAA1551 | PAR-CLIP |
| hsa-mir-193b-5p | SLC38A7 | PAR-CLIP |
| hsa-mir-193b-5p | MIOX | HITS-CLIP |
| hsa-mir-193b-5p | IPO9 | HITS-CLIP |
| hsa-mir-193b-5p | ZNF701 | HITS-CLIP |
| hsa-mir-193b-5p | MINDY1 | HITS-CLIP |
| hsa-mir-193b-5p | METTL2B | PAR-CLIP |
| hsa-mir-193b-5p | FOXJ2 | PAR-CLIP |
| hsa-mir-193b-5p | UTP6 | HITS-CLIP |
| hsa-mir-193b-5p | LMOD3 | HITS-CLIP//PAR-CLIP |
| hsa-mir-193b-5p | ISY1 | PAR-CLIP |
| hsa-mir-193b-5p | MAVS | HITS-CLIP |
| hsa-mir-193b-5p | TAOK1 | HITS-CLIP |
| hsa-mir-193b-5p | KIAA1456 | HITS-CLIP |
| hsa-mir-193b-5p | ZFP14 | PAR-CLIP |
| hsa-mir-193b-5p | CACNG8 | HITS-CLIP |
| hsa-mir-193b-5p | NPFFR1 | PAR-CLIP |
| hsa-mir-193b-5p | AEN | CLASH |
| hsa-mir-193b-5p | ZNF747 | PAR-CLIP |
| hsa-mir-193b-5p | CHAC1 | PAR-CLIP |
| hsa-mir-193b-5p | PPDPF | HITS-CLIP |
| hsa-mir-193b-5p | TTPAL | HITS-CLIP |
| hsa-mir-193b-5p | NKAP | PAR-CLIP |
| hsa-mir-193b-5p | CXorf36 | PAR-CLIP |
| hsa-mir-193b-5p | C12orf49 | PAR-CLIP |
| hsa-mir-193b-5p | ZMYM1 | HITS-CLIP |
| hsa-mir-193b-5p | ZNF669 | HITS-CLIP |
| hsa-mir-193b-5p | CPSF7 | PAR-CLIP |
| hsa-mir-193b-5p | SYNPO2L | PAR-CLIP |
| hsa-mir-193b-5p | ZNF556 | HITS-CLIP |
| hsa-mir-193b-5p | COQ10B | PAR-CLIP |
| hsa-mir-193b-5p | FAHD1 | HITS-CLIP |
| hsa-mir-193b-5p | KREMEN1 | HITS-CLIP |
| hsa-mir-193b-5p | ZRANB3 | PAR-CLIP |
| hsa-mir-193b-5p | YIPF4 | HITS-CLIP |
| hsa-mir-193b-5p | NOA1 | PAR-CLIP |
| hsa-mir-193b-5p | TNRC18 | PAR-CLIP |
| hsa-mir-193b-5p | MYPN | PAR-CLIP |
| hsa-mir-193b-5p | ZNF347 | PAR-CLIP |
| hsa-mir-193b-5p | RRP36 | HITS-CLIP |
| hsa-mir-193b-5p | KIR3DX1 | HITS-CLIP |
| hsa-mir-193b-5p | LYRM7 | PAR-CLIP |

| | | |
|---|---|---|
| hsa-mir-193b-5p | SLC38A5 | HITS-CLIP |
| hsa-mir-193b-5p | ZBTB47 | PAR-CLIP |
| hsa-mir-193b-5p | ZNF101 | HITS-CLIP |
| hsa-mir-193b-5p | FLYWCH2 | HITS-CLIP |
| hsa-mir-193b-5p | DIS3L | PAR-CLIP |
| hsa-mir-193b-5p | BORCS7 | HITS-CLIP |
| hsa-mir-193b-5p | SPPL3 | HITS-CLIP |
| hsa-mir-193b-5p | GJD3 | PAR-CLIP |
| hsa-mir-193b-5p | ZNF813 | HITS-CLIP |
| hsa-mir-193b-5p | ZNF491 | PAR-CLIP |
| hsa-mir-193b-5p | ZNF573 | PAR-CLIP |
| hsa-mir-193b-5p | C19orf47 | HITS-CLIP |
| hsa-mir-193b-5p | RNF19B | HITS-CLIP |
| hsa-mir-193b-5p | ICA1L | HITS-CLIP |
| hsa-mir-193b-5p | CMBL | PAR-CLIP |
| hsa-mir-193b-5p | OTUD6A | PAR-CLIP |
| hsa-mir-193b-5p | SMCR8 | HITS-CLIP |
| hsa-mir-193b-5p | PTGR2 | PAR-CLIP |
| hsa-mir-193b-5p | MANEAL | PAR-CLIP |
| hsa-mir-193b-5p | TCF23 | PAR-CLIP |
| hsa-mir-193b-5p | GPR155 | HITS-CLIP |
| hsa-mir-193b-5p | PAQR3 | HITS-CLIP//PAR-CLIP |
| hsa-mir-193b-5p | GPR156 | HITS-CLIP |
| hsa-mir-193b-5p | ZNF366 | PAR-CLIP |
| hsa-mir-193b-5p | ZNF384 | HITS-CLIP |
| hsa-mir-193b-5p | THAP8 | PAR-CLIP |
| hsa-mir-193b-5p | DENND6A | HITS-CLIP |
| hsa-mir-193b-5p | TMEM154 | HITS-CLIP//PAR-CLIP |
| hsa-mir-193b-5p | TMEM192 | HITS-CLIP |
| hsa-mir-193b-5p | C6orf89 | HITS-CLIP |
| hsa-mir-193b-5p | RSBN1L | HITS-CLIP |
| hsa-mir-193b-5p | FBXL13 | PAR-CLIP |
| hsa-mir-193b-5p | MSRB3 | HITS-CLIP |
| hsa-mir-193b-5p | SLC25A45 | PAR-CLIP |
| hsa-mir-193b-5p | CAVIN1 | PAR-CLIP |
| hsa-mir-193b-5p | GDPD1 | PAR-CLIP |
| hsa-mir-193b-5p | C5orf51 | PAR-CLIP |
| hsa-mir-193b-5p | ZNF677 | HITS-CLIP |
| hsa-mir-193b-5p | MACC1 | HITS-CLIP |
| hsa-mir-193b-5p | GSTK1 | PAR-CLIP |
| hsa-mir-193b-5p | C3orf62 | PAR-CLIP |
| hsa-mir-193b-5p | ZKSCAN4 | PAR-CLIP |
| hsa-mir-193b-5p | ZNF788 | PAR-CLIP |
| hsa-mir-193b-5p | C2orf68 | HITS-CLIP |
| hsa-mir-193b-5p | ZNF793 | HITS-CLIP |
| hsa-mir-193b-5p | HACD4 | PAR-CLIP |
| hsa-mir-193b-5p | TRIM72 | PAR-CLIP |

| | | |
|---|---|---|
| hsa-mir-193b-5p | C6orf132 | HITS-CLIP |
| hsa-mir-193b-5p | ANKRD33B | HITS-CLIP |
| hsa-mir-193b-5p | ZNF878 | HITS-CLIP |
| hsa-mir-193b-5p | ERVMER34-1 | HITS-CLIP |
| hsa-mir-193b-5p | C8orf17 | HITS-CLIP |
| hsa-mir-2355-5p | ABL2 | HITS-CLIP//PAR-CLIP |
| hsa-mir-2355-5p | XIAP | PAR-CLIP |
| hsa-mir-2355-5p | ZFP36L2 | PAR-CLIP |
| hsa-mir-2355-5p | CACNA1A | HITS-CLIP |
| hsa-mir-2355-5p | CAPZA1 | PAR-CLIP |
| hsa-mir-2355-5p | CCNF | PAR-CLIP |
| hsa-mir-2355-5p | CDKN1A | PAR-CLIP |
| hsa-mir-2355-5p | CDKN1B | PAR-CLIP |
| hsa-mir-2355-5p | CPM | HITS-CLIP |
| hsa-mir-2355-5p | CTNND1 | PAR-CLIP |
| hsa-mir-2355-5p | ECE1 | PAR-CLIP |
| hsa-mir-2355-5p | GCNT1 | HITS-CLIP |
| hsa-mir-2355-5p | GRIN2A | HITS-CLIP |
| hsa-mir-2355-5p | HDGF | PAR-CLIP |
| hsa-mir-2355-5p | HMGA1 | HITS-CLIP |
| hsa-mir-2355-5p | HNRNPH1 | HITS-CLIP |
| hsa-mir-2355-5p | HOXD9 | PAR-CLIP |
| hsa-mir-2355-5p | HSPA4 | HITS-CLIP |
| hsa-mir-2355-5p | HSPA6 | HITS-CLIP |
| hsa-mir-2355-5p | MCC | PAR-CLIP |
| hsa-mir-2355-5p | MVK | HITS-CLIP |
| hsa-mir-2355-5p | OAZ2 | HITS-CLIP |
| hsa-mir-2355-5p | CDK16 | PAR-CLIP |
| hsa-mir-2355-5p | PIGA | PAR-CLIP |
| hsa-mir-2355-5p | POU6F1 | HITS-CLIP |
| hsa-mir-2355-5p | PPP2R5E | HITS-CLIP |
| hsa-mir-2355-5p | PURA | HITS-CLIP |
| hsa-mir-2355-5p | PYCR1 | PAR-CLIP |
| hsa-mir-2355-5p | RPL41 | PAR-CLIP |
| hsa-mir-2355-5p | SRSF2 | PAR-CLIP |
| hsa-mir-2355-5p | SOX12 | HITS-CLIP |
| hsa-mir-2355-5p | SP1 | HITS-CLIP |
| hsa-mir-2355-5p | SPTBN2 | HITS-CLIP |
| hsa-mir-2355-5p | TBX15 | PAR-CLIP |
| hsa-mir-2355-5p | TTR | HITS-CLIP |
| hsa-mir-2355-5p | VASP | HITS-CLIP |
| hsa-mir-2355-5p | WNT2B | HITS-CLIP |
| hsa-mir-2355-5p | ZNF79 | HITS-CLIP |
| hsa-mir-2355-5p | ZNF138 | PAR-CLIP |
| hsa-mir-2355-5p | LUZP1 | PAR-CLIP |
| hsa-mir-2355-5p | PTP4A1 | PAR-CLIP |
| hsa-mir-2355-5p | BSND | HITS-CLIP |

| | | |
|---|---|---|
| hsa-mir-2355-5p | PABPN1 | PAR-CLIP |
| hsa-mir-2355-5p | SMC1A | PAR-CLIP |
| hsa-mir-2355-5p | SUCLG2 | HITS-CLIP |
| hsa-mir-2355-5p | TRIM24 | PAR-CLIP |
| hsa-mir-2355-5p | ALDH1A2 | HITS-CLIP |
| hsa-mir-2355-5p | GYG2 | HITS-CLIP |
| hsa-mir-2355-5p | TM4SF5 | PAR-CLIP |
| hsa-mir-2355-5p | SYT7 | PAR-CLIP |
| hsa-mir-2355-5p | GPR55 | HITS-CLIP |
| hsa-mir-2355-5p | GOSR1 | HITS-CLIP |
| hsa-mir-2355-5p | PTGES | HITS-CLIP |
| hsa-mir-2355-5p | CCDC144A | PAR-CLIP |
| hsa-mir-2355-5p | TMEM63A | PAR-CLIP |
| hsa-mir-2355-5p | RIMS3 | HITS-CLIP |
| hsa-mir-2355-5p | TSC22D2 | HITS-CLIP |
| hsa-mir-2355-5p | UBAP2L | HITS-CLIP |
| hsa-mir-2355-5p | FAM13A | HITS-CLIP |
| hsa-mir-2355-5p | DCAF7 | PAR-CLIP |
| hsa-mir-2355-5p | C1D | PAR-CLIP |
| hsa-mir-2355-5p | IGF2BP1 | HITS-CLIP |
| hsa-mir-2355-5p | PXMP4 | HITS-CLIP |
| hsa-mir-2355-5p | PHB2 | PAR-CLIP |
| hsa-mir-2355-5p | ZFP30 | HITS-CLIP |
| hsa-mir-2355-5p | AAK1 | HITS-CLIP |
| hsa-mir-2355-5p | CLUAP1 | HITS-CLIP |
| hsa-mir-2355-5p | WAPL | PAR-CLIP |
| hsa-mir-2355-5p | MRPS27 | HITS-CLIP |
| hsa-mir-2355-5p | RPRD2 | PAR-CLIP |
| hsa-mir-2355-5p | VPS8 | HITS-CLIP |
| hsa-mir-2355-5p | CBX6 | PAR-CLIP |
| hsa-mir-2355-5p | PISD | PAR-CLIP |
| hsa-mir-2355-5p | GABARAPL3 | HITS-CLIP |
| hsa-mir-2355-5p | THUMPD3 | HITS-CLIP |
| hsa-mir-2355-5p | HINFP | HITS-CLIP |
| hsa-mir-2355-5p | GMEB2 | PAR-CLIP |
| hsa-mir-2355-5p | DKK3 | PAR-CLIP |
| hsa-mir-2355-5p | ZNF638 | HITS-CLIP |
| hsa-mir-2355-5p | PARVB | PAR-CLIP |
| hsa-mir-2355-5p | TFCP2L1 | HITS-CLIP |
| hsa-mir-2355-5p | NOP53 | PAR-CLIP |
| hsa-mir-2355-5p | SOCS7 | PAR-CLIP |
| hsa-mir-2355-5p | MRPL4 | HITS-CLIP |
| hsa-mir-2355-5p | APH1A | PAR-CLIP |
| hsa-mir-2355-5p | MRPL30 | PAR-CLIP |
| hsa-mir-2355-5p | FAM8A1 | PAR-CLIP |
| hsa-mir-2355-5p | UFM1 | HITS-CLIP |
| hsa-mir-2355-5p | ERRFI1 | PAR-CLIP |

| | | |
|---|---|---|
| hsa-mir-2355-5p | FBXL19 | PAR-CLIP |
| hsa-mir-2355-5p | FNBP1L | HITS-CLIP |
| hsa-mir-2355-5p | DNAJC28 | HITS-CLIP |
| hsa-mir-2355-5p | CDCA4 | HITS-CLIP |
| hsa-mir-2355-5p | RCBTB1 | PAR-CLIP |
| hsa-mir-2355-5p | LRRC1 | PAR-CLIP |
| hsa-mir-2355-5p | NAGK | HITS-CLIP |
| hsa-mir-2355-5p | SLC30A6 | HITS-CLIP |
| hsa-mir-2355-5p | TENM3 | HITS-CLIP |
| hsa-mir-2355-5p | NAXD | PAR-CLIP |
| hsa-mir-2355-5p | ZNF415 | PAR-CLIP |
| hsa-mir-2355-5p | FAM212B | HITS-CLIP |
| hsa-mir-2355-5p | SAR1A | HITS-CLIP//PAR-CLIP |
| hsa-mir-2355-5p | EMC7 | HITS-CLIP |
| hsa-mir-2355-5p | ATXN7L3 | HITS-CLIP |
| hsa-mir-2355-5p | TOMM22 | HITS-CLIP |
| hsa-mir-2355-5p | IGSF9 | PAR-CLIP |
| hsa-mir-2355-5p | TAOK1 | PAR-CLIP |
| hsa-mir-2355-5p | WDFY1 | PAR-CLIP |
| hsa-mir-2355-5p | HIVEP3 | PAR-CLIP |
| hsa-mir-2355-5p | BACH2 | HITS-CLIP |
| hsa-mir-2355-5p | FAM217B | HITS-CLIP |
| hsa-mir-2355-5p | SUSD1 | HITS-CLIP |
| hsa-mir-2355-5p | INF2 | HITS-CLIP |
| hsa-mir-2355-5p | GIGYF1 | PAR-CLIP |
| hsa-mir-2355-5p | C16orf58 | PAR-CLIP |
| hsa-mir-2355-5p | ZNF649 | PAR-CLIP |
| hsa-mir-2355-5p | TMEM109 | HITS-CLIP |
| hsa-mir-2355-5p | OGFOD3 | HITS-CLIP |
| hsa-mir-2355-5p | GTDC1 | HITS-CLIP |
| hsa-mir-2355-5p | ZNF385D | HITS-CLIP |
| hsa-mir-2355-5p | ZMYM1 | HITS-CLIP//PAR-CLIP |
| hsa-mir-2355-5p | ZSCAN16 | HITS-CLIP |
| hsa-mir-2355-5p | TTYH3 | HITS-CLIP |
| hsa-mir-2355-5p | KCNH6 | HITS-CLIP |
| hsa-mir-2355-5p | SPRY4 | HITS-CLIP |
| hsa-mir-2355-5p | AMMECR1L | HITS-CLIP |
| hsa-mir-2355-5p | ZNF394 | PAR-CLIP |
| hsa-mir-2355-5p | TMEM246 | HITS-CLIP |
| hsa-mir-2355-5p | CMSS1 | PAR-CLIP |
| hsa-mir-2355-5p | FRMPD3 | HITS-CLIP |
| hsa-mir-2355-5p | NEURL4 | PAR-CLIP |
| hsa-mir-2355-5p | EBPL | HITS-CLIP |
| hsa-mir-2355-5p | PPP1R9B | PAR-CLIP |
| hsa-mir-2355-5p | C9orf3 | PAR-CLIP |
| hsa-mir-2355-5p | ARHGAP19 | HITS-CLIP |
| hsa-mir-2355-5p | DCLK3 | HITS-CLIP |

| | | |
|---|---|---|
| hsa-mir-2355-5p | MIDN | HITS-CLIP |
| hsa-mir-2355-5p | ZNF625 | PAR-CLIP |
| hsa-mir-2355-5p | SLC25A46 | HITS-CLIP |
| hsa-mir-2355-5p | SLC39A13 | HITS-CLIP |
| hsa-mir-2355-5p | R3HDM4 | PAR-CLIP |
| hsa-mir-2355-5p | NLRP12 | HITS-CLIP |
| hsa-mir-2355-5p | MYOZ3 | HITS-CLIP |
| hsa-mir-2355-5p | MEX3A | HITS-CLIP |
| hsa-mir-2355-5p | ASB16 | HITS-CLIP |
| hsa-mir-2355-5p | CHST14 | PAR-CLIP |
| hsa-mir-2355-5p | TOP1MT | PAR-CLIP |
| hsa-mir-2355-5p | GSTO2 | HITS-CLIP |
| hsa-mir-2355-5p | GJD3 | HITS-CLIP |
| hsa-mir-2355-5p | MFSD12 | HITS-CLIP |
| hsa-mir-2355-5p | RNF19B | HITS-CLIP |
| hsa-mir-2355-5p | ARHGEF19 | HITS-CLIP |
| hsa-mir-2355-5p | BBS5 | HITS-CLIP |
| hsa-mir-2355-5p | NDUFAF6 | HITS-CLIP |
| hsa-mir-2355-5p | ASB6 | PAR-CLIP |
| hsa-mir-2355-5p | SIRPA | HITS-CLIP |
| hsa-mir-2355-5p | PRICKLE1 | PAR-CLIP |
| hsa-mir-2355-5p | PTGR2 | HITS-CLIP |
| hsa-mir-2355-5p | KCTD11 | HITS-CLIP |
| hsa-mir-2355-5p | SIX5 | HITS-CLIP |
| hsa-mir-2355-5p | C2orf15 | PAR-CLIP |
| hsa-mir-2355-5p | ROPN1B | PAR-CLIP |
| hsa-mir-2355-5p | CXorf38 | HITS-CLIP |
| hsa-mir-2355-5p | ZFP1 | PAR-CLIP |
| hsa-mir-2355-5p | ZNF791 | PAR-CLIP |
| hsa-mir-2355-5p | ADAMTS17 | HITS-CLIP |
| hsa-mir-2355-5p | ZNF525 | PAR-CLIP |
| hsa-mir-2355-5p | DOK6 | HITS-CLIP |
| hsa-mir-2355-5p | NT5DC1 | HITS-CLIP |
| hsa-mir-2355-5p | RALGAPA1 | HITS-CLIP |
| hsa-mir-2355-5p | ST6GALNAC3 | PAR-CLIP |
| hsa-mir-2355-5p | ZNF740 | HITS-CLIP |
| hsa-mir-2355-5p | NEK8 | HITS-CLIP |
| hsa-mir-2355-5p | ZNF860 | HITS-CLIP |
| hsa-mir-2355-5p | FAM71F2 | HITS-CLIP |
| hsa-mir-2355-5p | ANKRD36 | HITS-CLIP |
| hsa-mir-2355-5p | ZNF788 | PAR-CLIP |
| hsa-mir-2355-5p | ACBD7 | PAR-CLIP |
| hsa-mir-2355-5p | LURAP1 | HITS-CLIP |
| hsa-mir-2355-5p | ZNF704 | PAR-CLIP |
| hsa-mir-2355-5p | SMIM15 | HITS-CLIP |
| hsa-mir-2355-5p | TMEM78 | HITS-CLIP |
| hsa-mir-2355-5p | ZNF286B | HITS-CLIP |

| | | |
|---|---|---|
| hsa-mir-2355-5p | POM121C | PAR-CLIP |
| hsa-mir-2355-5p | PPP5D1 | PAR-CLIP |

**Supplementary table 3**

List of primers for gene expression analysis

| Gene | Sequence |
|---|---|
| TMPRSS2 | Fw: ACACCAGCCATGATCTGTGC |
| | Rv: CAGAGGCCCTCCACTGTCA |
| ACE2 | Fw: GGACCCAGGAAATGTTCAGA |
| | Rv: GGCTGCAGAAAGTGACATGA |
| GAPDH | Fw: GAGTCAACGGATTTGTCGT |
| | Rw: GACAAGCTTCCCGTTCTCAG |

List of Taqman assays

| ID (gene) |
|---|
| Hs02800695_m1 (HPRT1) |
| Hs00154614_m1 (CSTF2) |
| Hs00961622_m1 (IL10) |
| Hs00989291_m1 (IFN$\gamma$) |
| Hs99999905_m1 (GAPDH) |
| 002113 (hsa-miR-31-3p) |
| 001048 (hsa-miR-503-5p) |
| 001048 (hsa-miR-503-5p) |
| 001006 (hsa-rnu48) |

**Supplementary figure 1. A** Box-plot analysis representing ACE2 gene expression levels in tumoral HNSCC TCGA samples according to the gender (female or male). **B** Box-plot analysis representing TMPRSS2 gene expression levels in non-tumorous (N) and tumor (T) tissues of oral cavity from the HNSCC TCGA dataset.

**Supplementary figure 2.** qRT-PCR analysis of ACE2 expression levels in Cal-27 and Detroit-562 cell lines upon silencing of p53 (sip53) or YAP (siYAP) or MYC (siMYC) compared to silencing scramble (value=1).

**Supplementary figure 3. A-B** List of genes for MYC (A) and immune (B) signatures.

**Supplementary table 1.** miRNA\TMPRSS2 predicted interactions by miRWalk.

**Supplementary table 2.** List of experimental validated miRNA-target interactions using miRNet.

**Supplementary table 3**. List of primers and probes used for qRT-PCR.
